# Synthetic dosage-compensating miRNA circuits allow precision gene therapy for Rett syndrome

**DOI:** 10.1101/2024.03.13.584179

**Authors:** Michael J. Flynn, Acacia M.H. Mayfield, Rongrong Du, Viviana Gradinaru, Michael B. Elowitz

## Abstract

A longstanding challenge in gene therapy is expressing a dosage-sensitive gene within a tight therapeutic window. For example, loss of *MECP2* function causes Rett syndrome, while its duplication causes *MECP2* duplication syndrome. Viral gene delivery methods generate variable numbers of gene copies in individual cells, creating a need for gene dosage-invariant expression systems. Here, we introduce a compact miRNA-based, incoherent feed-forward loop circuit that achieves precise control of *Mecp2* expression in cells and brains, and improves outcomes in an AAV-based mouse model of Rett syndrome gene therapy. Single molecule analysis of endogenous and ectopic *Mecp2* mRNA revealed precise, sustained expression across a broad range of gene dosages. Delivered systemically in a brain-targeting AAV capsid, the circuit strongly suppressed Rett behavioral symptoms for over 24 weeks, outperforming an unregulated gene therapy. These results demonstrate that synthetic miRNA-based regulatory circuits can enable precise in vivo expression to improve the safety and efficacy of gene therapy.

**One sentence description:** A synthetic miRNA-based incoherent feed-forward loop circuit embedded in a gene delivery vector overcomes the challenge of overexpression toxicity in a mouse model of Rett syndrome gene therapy.

## Introduction

Gene therapy promises to enable lasting cures for genetic diseases by delivering corrected copies of genes mutated within a patient. Recently, the field has achieved clinical successes addressing spinal muscular atrophy (SMA) (*1*), hemophilia (*2–4*), and inherited retinal dystrophy (*5*), with other applications in the pipeline (*6*). New technologies, such as adeno-associated viral vectors (AAVs) that target specific cell types at high efficiency (*7*), may accelerate progress in the coming decade.

However, many gene therapies face a ‘Goldilocks’ problem: too little expression of the gene leads to the disease phenotype, but too much expression can induce other disease phenotypes. For example, loss of function of the *SYNGAP1* gene results in non-syndromic intellectual disability and epilepsy (*8*) while overexpression of *SYNGAP1* leads to a pronounced depression of excitatory signaling in vitro (*9*). Similarly, *UBE3A* deficiency causes Angelman’s syndrome, but duplication and triplications are associated with autism spectrum disorder (*10*). Overexpression toxicity is also a concern in clinical gene replacement. For example, loss of function mutations in *SMN1* cause spinal muscular atrophy, but overexpression of *SMN1* through gene therapy led to clinically silent but concerning dorsal root ganglion pathology in mouse and non-human primate studies of SMA gene therapy (*11–13*).

Rett syndrome presents the prototypical Goldilocks problem. It is a severe neurodevelopmental disease caused by loss-of-function mutations in the gene encoding MeCP2, a methyl-CpG/A binding protein (*14*). MeCP2 binds to methylated regions of the genome and serves as a binding hub and bridge to the NCoR/SMRT co-repressor complex to repress methylated genes, an essential function for brain maturation (*15*). Since *MECP2* is on the X chromosome, mutations in the gene lead to different symptoms between males and females. Human males cannot survive with a single non-functional copy of *MECP2* (*16*). In human females heterozygous for a mutation in *MECP2*, Rett syndrome is characterized by a severe developmental regression at 7-18 months of age, progressive loss of speech and hand use, ataxia, and acquired microcephaly, among other symptoms (*17*). MeCP2-deficient male mice show reductions in lifespan, brain size, neuron soma size, synapse counts, dendritic spine density, and electrophysiological activity (*18*). However, mild overexpression of MeCP2 also leads to a disease phenotype (*19*) and duplication of the *MECP2* gene causes another disorder, *MECP2* duplication syndrome (*20*). In engineered mice, Cre-based reactivation of *Mecp2* expression from its endogenous genomic context alleviates disease phenotypes, suggesting that the condition is reversible (*21*). However, gene therapies based on AAV-mediated delivery of *Mecp2* to Rett model mice have induced toxicity (*22–27*), specifically from overexpression of the MeCP2 protein (*23*). Thus, a critical requirement for Rett syndrome gene therapy is to express MeCP2 within a narrow therapeutic window.

In this work, we designed a synthetic biological circuit that provides this capability, and demonstrated that it can improve Rett syndrome gene therapy in vivo. This compact miRNA-based incoherent feedforward loop (IFFL) circuit limited *Mecp2* expression and reduced its sensitivity to gene dosage in cell culture. Further, it restricted ectopic *Mecp2* mRNA to levels comparable to, but not exceeding, those of endogenous *Mecp2* in the mouse brain. Finally, a gene therapy vector containing the circuit outperformed unregulated gene therapy, improving behavioral symptoms over a timescale of 24 weeks. These results demonstrate that an integrated miRNA-based gene circuit can improve gene therapies for Rett syndrome and likely for other genetic diseases as well.

## Results

### Modeling predicts incoherent feedforward regulation can tune protein expression to within a therapeutic window

Expression patterns of ectopic genes may deviate significantly from the endogenous distribution of expression for several reasons. First, the number of gene copies delivered to an individual cell can vary by orders of magnitude due to varying uptake efficiency of different organs and cell types (**Figure 1A**, upper right) (*28*, *29*), spatial gradients around a direct injection site (**Figure 1A**, bottom left) and Poisson-distributed stochastic vector uptake by individual cells (**Figure 1A**, bottom right) (*30*). Second, engineered promoters typically used in gene therapy vectors often induce stronger expression than endogenous promoters, potentially leading to toxicity from even a single copy (**Figure 1B**, left panel) (*31*). Finally, for an X-linked disease such as Rett syndrome, cells may express either zero or one copy of the endogenous gene due to X inactivation. Thus, even if an ectopic gene is expressed at the physiological levels this could still lead to overexpression in cells that express the endogenous copy (**Figure 1B**, right panel).

**Figure 1.**
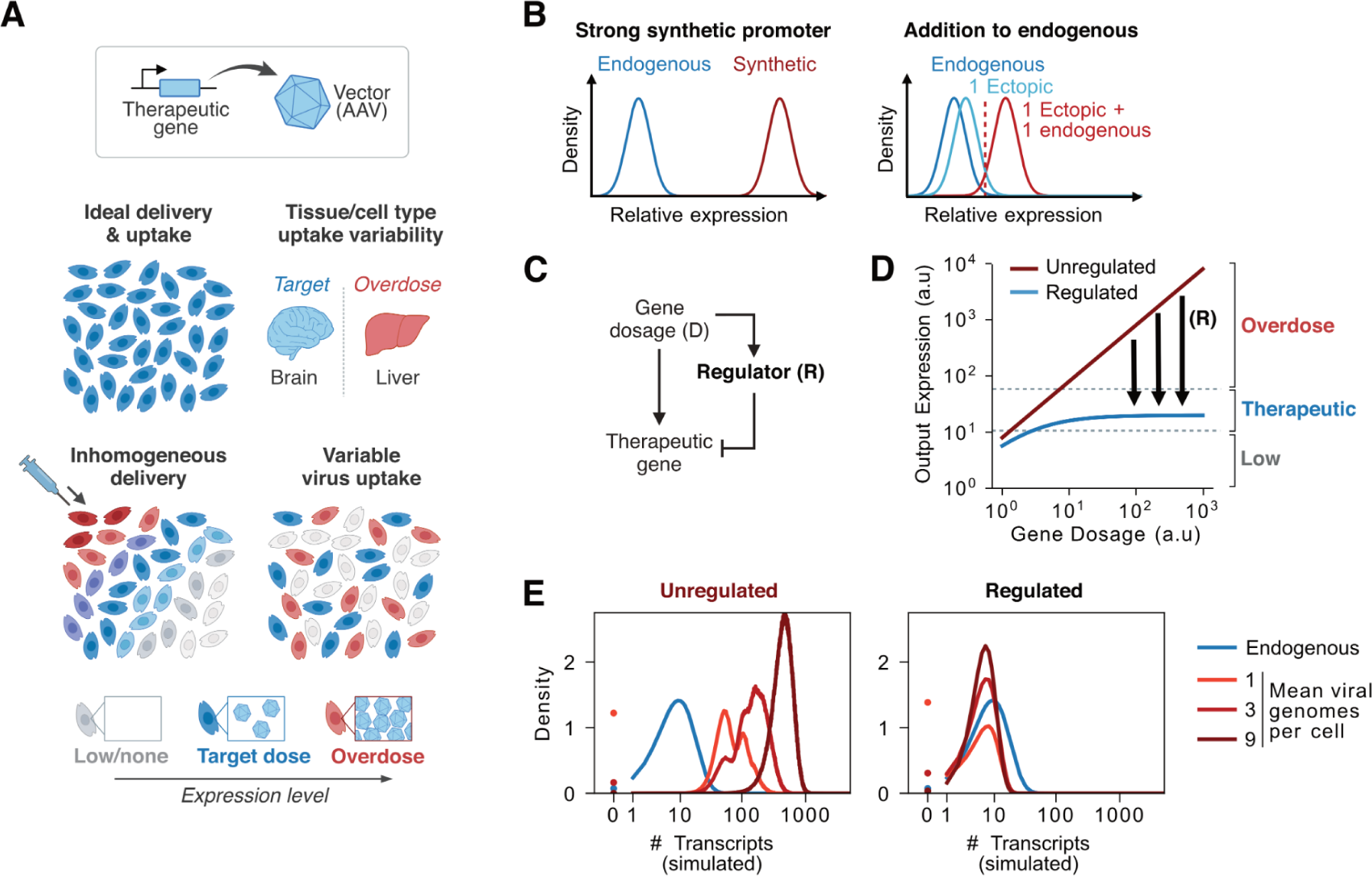
Mathematical modeling predicts that incoherent feedforward loop circuits can maintain gene expression within a therapeutic window. **(A)** Gene therapy contends with multiple sources of variability in expression. Ideally, all cells would receive the same number of viral genome copies, and express the correct amount of the therapeutic gene (upper left). However, viral uptake rates can vary greatly by organ and cell type (upper right), such that a dose that is therapeutic in one organ (e.g. brain, blue) may be toxic in another that takes up viral vectors at a higher rate (e.g. liver, red). With direct injection, cells close to the injection site receive more copies than cells farther away (lower left). Finally, even with correct mean delivery, viral uptake remains subject to stochastic variation (lower right). **(B)** The high level of expression induced by synthetic promoters commonly used in gene therapy may cause toxic overexpression from even a single transgene copy (left). Additionally, for X-linked genes like *MECP2*, approximately half of cells in affected females express a fully functioning endogenous copy. The gene therapy must not overexpress MeCP2 when its expression is added to the wildtype allele (right). **(C)** Schematic of an incoherent feedforward loop motif in which a therapeutic gene is co-expressed with its own negative regulator. **(D)** Therapeutic gene expression as a function of gene dosage, as modeled for an idealized IFFL. The increasingly negative action of the repressor (R, black) compensates for increases in gene dosage, leading to regimes where large changes in gene dosage yield nearly the same output expression of the circuit (blue), preventing overexpression. **(E)** Simulated distributions of therapeutic gene expression at different viral MOI, either unregulated (left) or regulated by an IFFL (right), compared to a target endogenous expression distribution (blue). Simulations incorporate stochastic viral uptake, bursty transcription, and stochastic enzyme kinetics as well as an offset between single-copy expression and the endogenous level of a therapeutic gene. The IFFL circuit compensates for these sources of variation.

Current approaches to limiting ectopic expression include optimizing the promoter (*32*) or incorporation of target sites for endogenous miRNAs (*33*). However, while these approaches generally reduce mean expression relative to unregulated constructs, they cannot actively adapt to variation in gene dosage.

The incoherent feedforward loop (IFFL) is an adaptive biological circuit motif that could address these challenges (*34*, *35*). Previous work has shown that synthetic IFFL circuits can successfully buffer gene expression against variations in gene dosage (*36–39*), noise from upstream regulators (*40*), competition for cellular resources (*39*, *41*), or general perturbations (*42*). Here, we consider IFFLs in which a target gene and its negative regulator are co-transcribed, so that higher gene dosage leads to greater transcription rates of both components (**Figure 1C**) (*34*, *35*, *43*). A simple mathematical model of such an IFFL showed that, above a minimal expression level, the two effects can cancel out to maintain a fixed mean expression level of the target gene across a wide range of gene dosages (**Figure 1D**, **Methods**). We further extended this model to incorporate discrete gene copy numbers, stochastic vector delivery, and bursty gene expression kinetics. We also incorporated a strong promoter so there is overexpression at even a single copy. Simulated IFFLs operating in these regimes successfully regulated the distribution of expression to be similar to a target endogenous distribution, across different MOIs (**Figure 1E, Methods**). These results suggested that a suitably engineered IFFL could generate a more robust gene therapy expression system.

### A synthetic host-targeting miRNA module enables dosage compensation of *Mecp2* expression

In some natural genes, intronic miRNAs downregulate expression of their host gene, forming an IFFL within a single transcript (*44–46*). This circuit architecture has also been demonstrated synthetically (*36*) and could have desirable properties for gene therapy, since intronic miRNA expression cassettes are non-immunogenic and genetically compact. However, it has not been established if synthetic miRNA-based IFFLs can match the expression level of an endogenous mRNA, or whether they can improve the function of an AAV-based gene therapy.

To address these questions, we designed a set of miRNA-based dosage compensating IFFL constructs (**Figure 2A**). We engineered a divergent promoter made up of the CMV enhancer flanked by the MeP229 promoter (*22*) in the forward direction and an intron-free Ef1ɑ promoter in the reverse direction (‘ECM promoter’). The forward promoter drives expression of a previously characterized MeCP2-EGFP protein fusion to facilitate analysis of protein expression (*26*). The reverse promoter drives expression of unregulated mRuby3 as an indicator of gene dosage.

**Figure 2.**
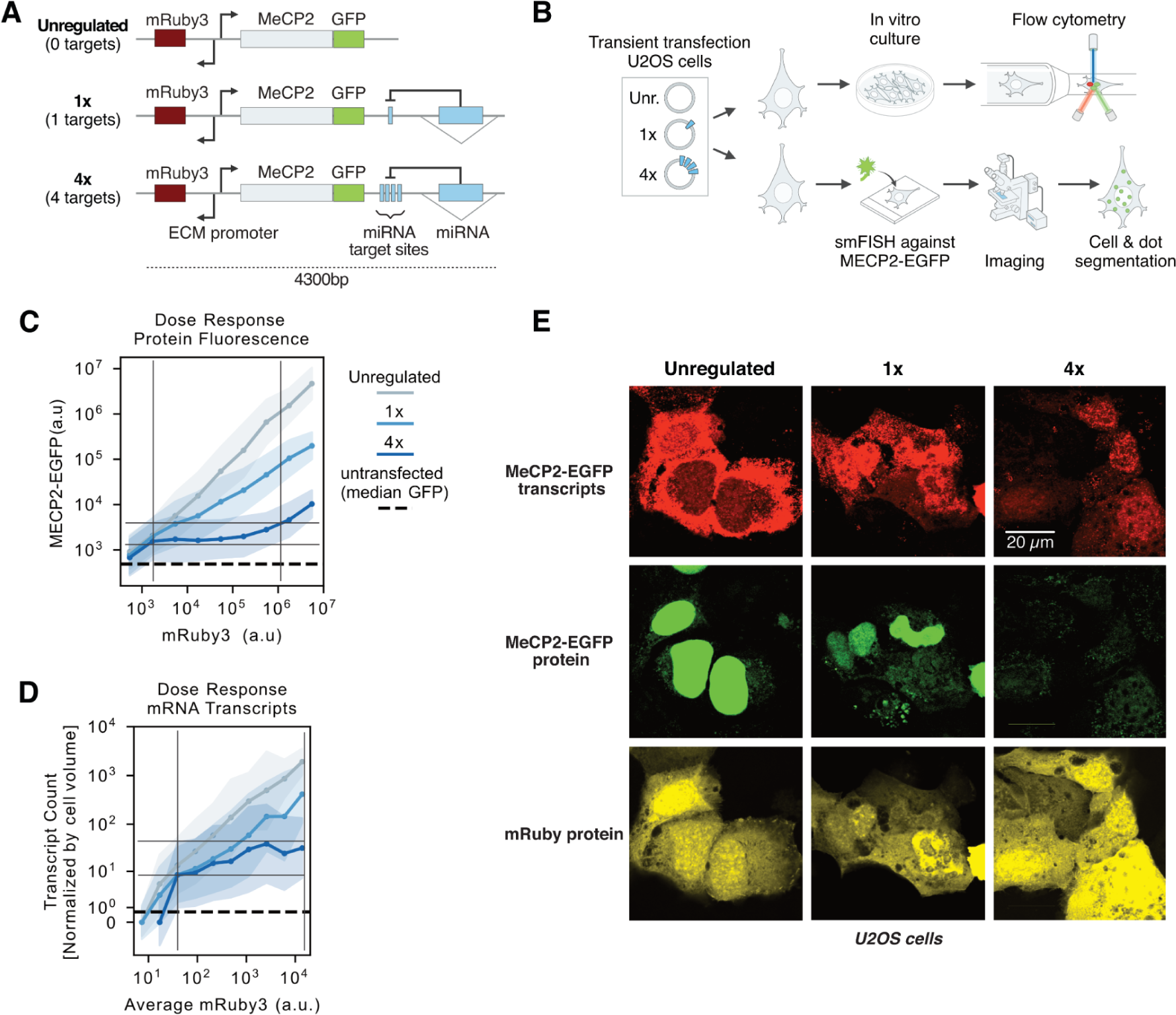
Synthetic miRNA IFFLs can adapt to variations in gene dosage in cell culture. **(A)** Designs of 3 constructs based on a divergent promoter producing MeCP2-EGFP in the forward direction and the mRuby3 dosage indicator in the reverse. The first circuit, labeled “unregulated”, has no miRNA targets and no miRNA cassette and serves as an unregulated control. The second (1x) and third (4x) circuits contain a miRNA cassette located within a synthetic intron in the 3’ UTR of *Mecp2-EGFP*, which respectively targets 1 or 4 fully complementary miRNA target sites upstream of the intron. All constructs are less than 4300 bp and fit inside an AAV. **(B)** Workflow to characterize circuit performance at both the mRNA and protein levels. For protein, U2OS cells were transiently transfected, cultured for 48 hours, and had protein expression measured by flow cytometry (upper path). For mRNA, U2OS cells were transiently transfected, incubated for 48 hours, fixed, and then analyzed with smFISH and confocal microscopy (lower path). **(C)** MeCP2-EGFP protein fluorescence as a function of mRuby3 dosage indicator for the 3 constructs, as measured by flow cytometry. MeCP2-EGFP was proportional to dosage for the unregulated construct (gray), as expected. For the 1x construct (medium blue), the slope was reduced, indicating a decreased responsiveness to dosage. For the 4x construct (dark blue), MeCP2-EGFP expression was nearly independent of dosage over 2.5 orders of magnitude variation in gene dosage (framed region). This stable expression level was approximately 3-fold above the fluorescence of untransfected cells (dashed black line). Here, and in D, shaded regions represent ±1 standard deviation of the logarithmic expression values. **(D)** *Mecp2-EGFP* transcript count as a function of average mRuby3 fluorescence, as measured by smFISH and confocal microscopy. The relationship between *Mecp2-EGFP* transcripts and dosage indicator fluorescence largely agreed with the protein-level results for each construct. The 4x construct produced an expression level that varied less than 4-fold over a greater than 300-fold range of dosage (framed region). **(E)** smFISH imaging of ectopic *Mecp2-EGFP* transcripts (upper row) and protein (middle row), as well as mRuby dosage indicator (lower row). Cells displayed comparable levels of mRuby protein in all conditions (bottom row), while *Mecp2-EGFP* expression decreases with stronger IFFL regulation at both transcript (upper) and protein (middle) levels.

To implement IFFL regulation, we incorporated a miRNA expression cassette in a synthetic intron (*47*) within the 3’UTR of *Mecp2-EGFP*. This miRNA cassette is based on the strong and well-characterized miR-E backbone (*48*), which generates a miRNA complementary to a 21-bp sequence derived from *Renilla luciferase,* which is orthogonal to the human genome. To compare two different strengths of regulation, we inserted either 1 or 4 copies of the target sequence into the 3’UTR, upstream of the miRNA-containing intron, to create “1x” and “4x” circuits (**Figure 2A**). We also constructed an “unregulated” control construct lacking both the miRNA and its target sites. All 3 constructs were less than 4300 base pairs in length, and thus small enough to be efficiently packaged inside an AAV.

To test the ability of these circuits to compensate for variation in gene dosage, we quantified their expression as a function of dosage at the protein and mRNA levels. For protein-level quantification, we transiently transfected U2OS cells and analyzed MeCP2 expression using flow cytometry (**Figure 2B**, upper path, **Methods**). Unregulated MeCP2 was expressed at a level proportional to gene dosage, as expected (**Figure 2C**). The 1x construct reduced MeCP2-EGFP expression and its dependence on dosage (slope of EGFP versus mRuby3) (**Figure 2C**). While it did not achieve complete dosage compensation, it provided a useful intermediate-regulation condition for subsequent studies. The 4x circuit generated behavior closer to that expected from the simplified IFFL model (**Figure 1D**), with relatively constant (<3-fold variation) expression across a broad range (>300-fold) of gene dosage **(Figure 2C**).

To quantify these differences in expression at the mRNA level, we transiently transfected U2OS cells with each construct, performed smFISH (*49*) against the *Mecp2-EGFP* transcript, and imaged both protein fluorescence and transcripts using confocal microscopy (**Figure 2B**, lower path, **Methods**). Consistent with the flow cytometry results, among cells expressing similar levels of the mRuby dosage reporter, MeCP2-EGFP mRNA and protein expression levels decreased with increasing number of miRNA target sites (**Figure 2E**).

To quantitatively measure the relationship between gene dosage and target mRNA levels, we computationally segmented cells in the smFISH images and counted individual transcripts (dots in **Figure 2E**, **Methods**). The unregulated and 1x constructs showed linear and sublinear, but still increasing, dependence on the mRuby3 dosage indicator, respectively (**Figure 2D**). The 4x construct exhibited the lowest level of expression, which also varied less than 4-fold over a greater than 300-fold range of dosage (**Figure 2D**).

To test for potential off-target effects of the miRNA regulation, we performed bulk-RNAseq and compared transcriptome expression of a BFP-miRNA (miR-E) cassette to a negative control transfection of a BFP-only expression vector (**Methods**). Few genes were relatively up- or down-regulated in the miRNA condition and none contained partial sequence matches to the miRNA (**Supplementary Figure 1**). These results suggest any sequence-specific perturbations from the synthetic miRNA itself were minimal.

Taken together, these results indicate that the 4x IFFL circuit can establish dosage-insensitive expression and reduce the magnitude of cell-cell variation in gene expression in vitro, without perturbing endogenous gene expression.

### AAV-delivered IFFL circuits reduce *Mecp2* mRNA to endogenous levels in mouse brains

Maximizing therapeutic efficacy and safety in Rett syndrome gene therapy requires expressing AAV-delivered *Mecp2* at appropriate levels in the brain. While the precise therapeutic window of *Mecp2* expression is unknown, it presumably spans the endogenous expression range. We therefore sought to quantitatively compare expression of AAV-delivered *Mecp2* with endogenous *Mecp2* in mouse brains. To distinguish ectopic and endogenous transcripts, we designed two sets of orthogonal hybridization chain reaction (HCR) probes: one set targeted the EGFP sequence exclusive to the ectopic transcript, while the other set targeted sequences in the 3’ UTR of the major isoforms of *Mecp2* that are exclusive to the endogenous gene (**Figure 3A**).

**Figure 3.**
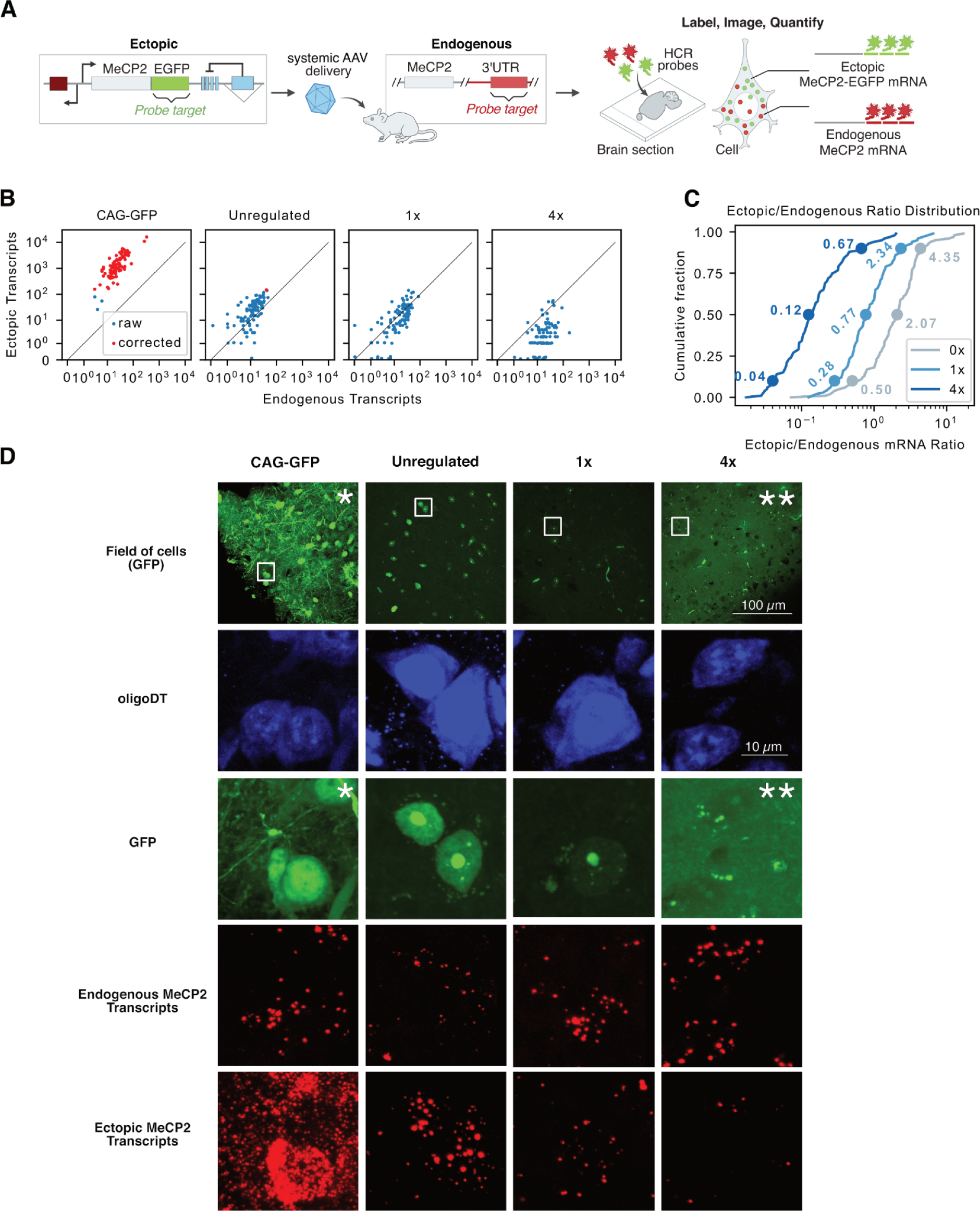
Synthetic miRNA IFFLs regulate expression to near or below endogenous *Mecp2* levels in mouse brains. **(A)** Orthogonal HCR probes were designed to specifically target endogenous or ectopic *Mecp2*. Ectopic *Mecp2-EGFP* was targeted with HCR probes against the EGFP coding sequence. Endogenous *Mecp2* was targeted with HCR probes against sequences in the endogenous 3’ UTR which do not appear in the ectopic construct. Mice were injected with viral constructs and, after 3 weeks of expression, brain slices were analyzed by HCR and confocal microscopy. **(B)** Ectopic (y-axis) vs endogenous (x-axis) transcripts measured in single cells (**Methods**). The diagonal black line denotes equal expression. Red dots denote cells whose counts have been corrected to account for dense dot spacing (**Methods**, **Supplementary Figure 3**). *CAG-EGFP* expressed ectopic transcript at levels an order of magnitude greater than endogenous *Mecp2* transcripts. The unregulated, 1x, and 4x constructs showed progressively reduced ectopic expression, with the 1x construct matching endogenous *Mecp2* transcript levels, and the 4x construct expressing lower levels. **(C)** Distributions of the ectopic to endogenous *Mecp2* transcripts in single cells. Annotations denote the 10th, 50th and 90th percentiles. The median cell receiving the unregulated construct overexpressed ectopic *Mecp2* by a factor of 2.1 relative to endogenous levels. With the 1x circuit, the median cell expressed ectopic *Mecp2* at 0.8 times the endogenous level. However, 10% of cells overexpressed ectopic *Mecp2* by a factor of at least 2.3. The median cell receiving the 4x construct only expressed 0.14 ectopic transcripts per endogenous transcript, but few cells overexpressed *Mecp2*. **(D)** Sample HCR images, focusing on individual cells in a field of cortical neurons (first row, white boxes denote enlarged areas below). All cells exhibited similar endogenous *Mecp2* expression (fourth row), but decreasing ectopic MeCP2-EGFP protein (third row) and ectopic *Mecp2* transcripts (fifth row) from CAG-GFP to unregulated to 1x to 4x constructs. (*) Brightness of CAG-GFP image has been reduced to better distinguish cells. (**) Brightness of the 4x-GFP image has been increased to make the dimmer fluorescence of MeCP2 nuclear puncta more visible.

To achieve brain-specific delivery, we took advantage of a recently developed AAV capsid variant, Cap.B22, which efficiently targets the brain when systemically delivered, while de-targeting the liver and other organs (*50*). We packaged each of the constructs in Cap.B22. To compare circuit-regulated expression to that produced by a typical promoter commonly used in gene therapy (*31*, *51*, *52*), we also packaged a CAG-GFP construct based on the synthetic CAG promoter. We then systemically injected each variant into WT mice at a dose of 5×10^12^ viral genomes (vg) per mouse. 3 weeks post-injection, we collected brain sections and analyzed mRNA levels using HCR and confocal microscopy (**Figure 3A, Methods**). In the resulting images, the highest expression of Mecp2 mRNA and protein was seen in brains that received the CAG-GFP construct, followed by the unregulated, 1x, and 4x constructs in that order (**Figure 3D**).

To quantify the effect of the circuit in single cells, we segmented cells based on oligo-dT fluorescence (**Figure 3D**, second row) and counted individual transcripts within individual cells (**Methods**). The CAG-GFP construct overexpressed the ectopic transcript by a median factor of 54-fold relative to endogenous *Mecp2* (**Figure 3B**, first panel), indicating that standard synthetic promoters can greatly overexpress *Mecp2*.

The unregulated, 1x, and 4x constructs all exhibited distinct behaviors. The unregulated construct overexpressed *Mecp2*, but to a lesser extent than CAG-GFP, with the median cell expressing 2-fold more ectopic than endogenous transcripts (**Figure 3B**, second panel, **Figure 3C**). Since even mild overexpression of MeCP2 was previously found to be harmful (*19*), this could potentially lead to toxicity. Interestingly, the 1x construct matched the expression of *Mecp2* quite well at the median of the distribution, with 0.77 ectopic transcripts expressed per endogenous transcript. However, a tail of ∼10% of cells exhibited at least 2.3-fold more ectopic than endogenous transcripts (**Figure 3B**, third panel, **Figure 3C)**. By contrast, the 4x construct underexpressed *Mecp2* at the median (ectopic:endogenous ratio = 0.12) and did not exhibit a tail of overexpressing cells (**Figure 3B**, fourth panel, **Figure 3C**). The 4x circuit thus ensured that nearly the entire distribution was at or below the endogenous level. Few transcripts were measured in the negative control condition, where primary HCR probes were not added, with 48% of cells expressing one or zero counts of both transcripts (**Supplementary Figure 2**).

These results indicate that the IFFL circuits can regulate *Mecp2* expression within the mouse brain, and can achieve expression levels comparable to or less than that of endogenous *Mecp2*. They also reveal that different construct designs can generate distinct distributions of relative expression levels. Finally, they provoke the critical question of how these different distributions may ultimately impact disease progression in a model organism.

### Circuit-regulated AAV-MeCP2 outperforms unregulated AAV-MeCP2 in a mouse model of Rett syndrome

To assess whether regulating the expression of AAV-delivered *Mecp2* improves behavioral outcomes in Rett model mice compared to unregulated gene therapy, we evaluated the impact of the three constructs in a mouse line carrying a *Mecp2*-null allele (*53*). To quantify outcomes, we used the standardized Rett phenotype score, which is based on several motor phenotypes, including hindlimb clasping and gait analysis (**Methods**) (*21*). Female *Mecp2^-/X^* mice were divided into 4 groups (*n* ≥ 5): treated with the unregulated, 1x, or 4x constructs or left uninjected. At 4 weeks of age, baseline Rett phenotype scores were recorded. Mice were then systemically injected with AAV-CAP.B22-packaged constructs at a dose of 10^14^ vg/kg. The Rett phenotype score was then measured biweekly until the mice reached 28 weeks of age (**Figure 4A**).

**Figure 4.**
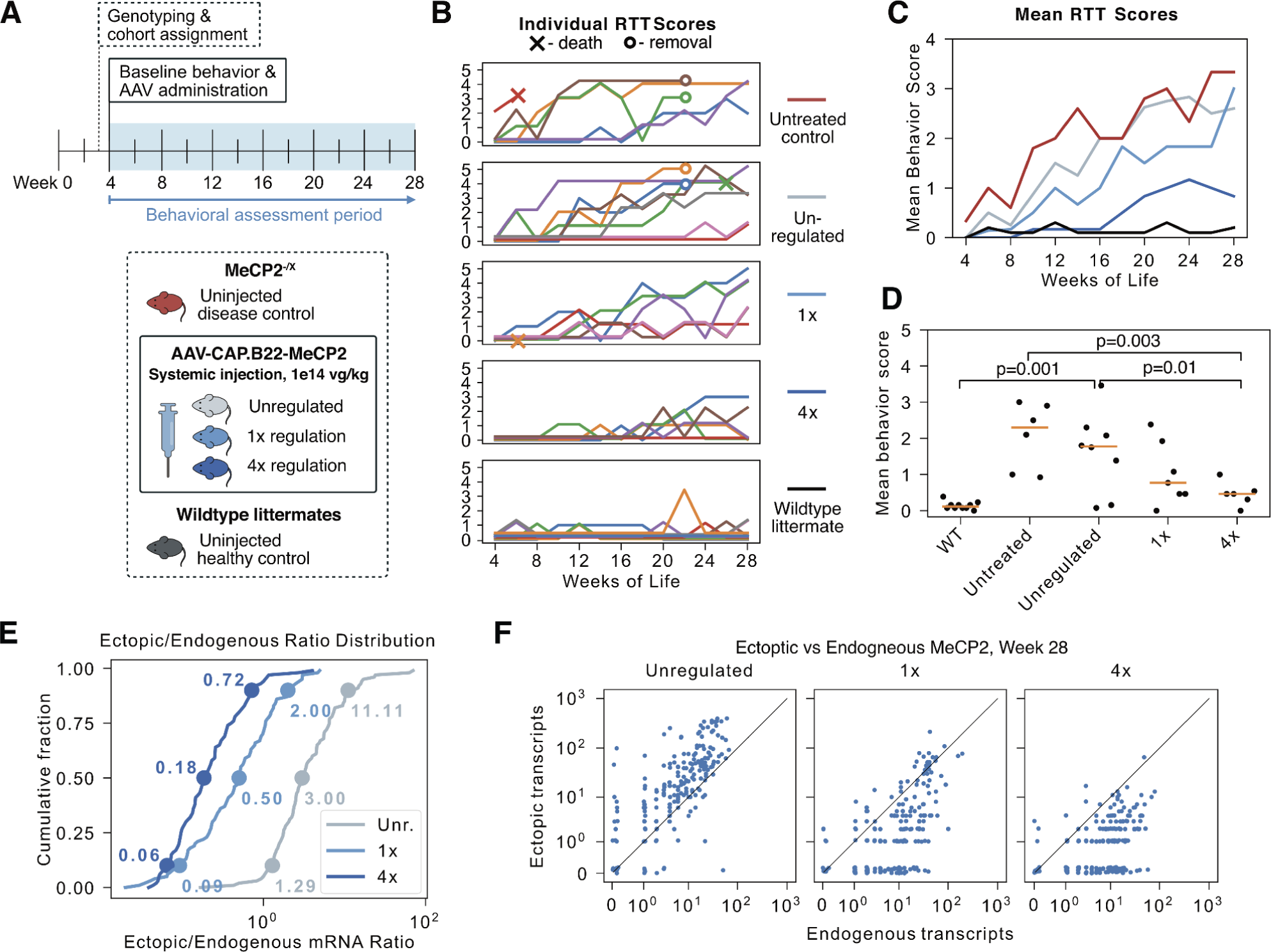
IFFL-regulated gene therapy outperforms unregulated gene therapy in a mouse model of Rett syndrome. **(A)** Female *Mecp2^-/X^* mice were divided into 4 treatment groups: uninjected or injected with one of the 3 constructs (Figure 2A) packaged in AAV-CAP.B22. Female wildtype littermates were included as healthy controls. At 4 weeks of age, baseline Rett behavior scores were recorded and the mice were injected with 1×10^14^ vg/kg. Rett phenotype scores (*21*) were then measured biweekly for 24 weeks. **(B)** Rett behavior (RTT) scores over time for individual mice (colored tracks) in each group. In each of the uninjected control, unregulated, and 1x groups, 1 mouse died during the study (“X” marker). In the 4x and wildtype groups, no mice died. After 22 weeks, 2 mice were removed from both the uninjected control group and the unregulated group for another experiment (“O” marker). **(C)** Mean score trajectories for each group in (B). Legend in left column of (B). **(D)** Rett behavior scores averaged across all timepoints for each mouse. Orange bars denote the median of each group. Mice that received the 4x construct had mean scores significantly lower than both the uninjected mice (bootstrap p=0.003) and mice that received the unregulated construct (bootstrap p=0.01). The standard deviation of the 4x group, 0.30, was also significantly lower than that of the unregulated group, 1.04 (bootstrap p=0.02). The mean scores of the wildtype controls were significantly lower than the uninjected controls (bootstrap p=0.0003) and mice receiving the unregulated (bootstrap p=0.001) and the 4x (bootstrap p=0.01) constructs. **(E)** Ectopic vs endogenous *Mecp2* transcripts quantified in single cells of mouse brains after behavior experiments were completed (week 28). Brain slices were analyzed by HCR as in Figure 3. Despite continuous expression for 24 weeks, ectopic and endogenous levels remained similar for each construct. **(F)** Distribution of the ratios of ectopic to endogenous transcripts in single cells after 24 weeks of expression (cf. Figure 3C). The unregulated construct overexpressed *Mecp2* by a factor of 3 at the median, while the 1x construct roughly agreed at the median, but overexpressed in the tail. The 4x construct was expressed below endogenous levels at the median, with fewer overexpressing cells.

We observed an increase in phenotypic markers (hindlimb clasping and abnormal gait) in untreated female mice, but not in their wildtype littermates (**Methods, Figure 4B,C**). The mice that received unregulated *Mecp2* exhibited a similar motor phenotype to the untreated *Mecp2^-/X^*animals, potentially due to *Mecp2* overexpression-induced toxicity (**Figure 4B,C**) (*19*). The 1x construct produced a mild improvement in symptoms, suggesting that limiting *Mecp2* expression can reduce toxicity (**Figure 4B,C**). Strikingly, the mice that received the tightly regulated (4x) *Mecp2* performed more comparably to WT controls than the other constructs (**Figure 4B,C**), with time-averaged behavior scores significantly lower than those of both the uninjected mice (bootstrap p = 0.003) and the mice that received the unregulated construct (bootstrap p = 0.01) (**Figure 4D**). Similar significant differences were found at individual timepoints, including week 22 (**Supplementary Figure 4A**) and week 28 (**Supplementary Figure 4B**). Notably, the standard deviation of the time-averaged scores for the 4x construct, 0.30, was also significantly lower than that of the unregulated construct, 1.04 (bootstrap, p=0.02), demonstrating that the regulated gene therapy causes a less variable phenotype score. These results demonstrate that tighter regulation improves the efficacy of AAV-delivered *Mecp2* in mitigating progression of the Rett phenotype in female *Mecp2^-/X^* mice.

Not all phenotypes of the Rett mouse model were improved by this gene therapy. Notably, female Rett model mice have an obesity phenotype that was not significantly affected by any of the injected constructs (**Supplementary Figure 5**). Additionally, we assessed whether regulation of *Mecp2* expression could confer any benefit to *Mecp2*-null male mice, which express no MeCP2 in the brain and have a much more severe phenotype, including a 12-week lifespan. However, we found no extension of the lifespan of male *Mecp2*-null male mice in any condition (**Supplementary Figure 6**). This contrasts with modest lifespan extensions reported with other Rett syndrome gene therapies (*23*), and with the strong behavioral results in the female mice in this work. Thus, while the regulated-*Mecp2* construct achieved strong phenotypic improvements in females without toxicity, it does not fully address all Rett phenotypes.

Ectopic constructs are potentially subject to unintended regulation or epigenetic silencing that could impact their operation over time. To check for such effects, we repeated the HCR analysis on the brains of mice that had finished the behavioral testing and had expressed the constructs for more than 24 weeks. All three constructs exhibited similar expression profiles as they did in the initial analysis (**Figure 3**). Specifically, the unregulated construct produced an expression distribution whose median cell overexpressed *Mecp2* 3-fold (**Figure 4E,F**). The 1x construct matched endogenous expression better at the median (0.5-fold), but exhibited a tail of ∼10% of cells overexpressing *Mecp2* at least 2-fold (**Figure 4E,F**). The 4x construct underexpressed *Mecp2*, and did not exhibit a tail of overexpressing cells (**Figure 4E,F**). Thus, the constructs maintained their expression profiles robustly through the 24 weeks of the study.

These results demonstrate that a tightly regulating synthetic miRNA IFFL can significantly augment a gene therapy for Rett syndrome in a Rett mouse model, and maintain the regulated level of expression over periods of at least 24 weeks.

## Discussion

Without regulation, therapeutic genes are expressed at varying degrees across cells, which could range from insufficient to toxic levels (**Figure 1**). Here, we designed and optimized a compact synthetic miRNA-based incoherent feedforward loop circuit that can compensate for variations in gene dosage and other sources of expression variation to ensure that total mRNA and protein expression remains within a therapeutic window (**Figure 2**). When incorporated in an AAV vector, the circuit maintained expression in mouse brains at or below the level of endogenous *Mecp2* (**Figure 3**). In the context of MeCP2 gene therapy, the tightly regulated (“4x”) circuit variant rescued behavioral phenotypes in a Rett female mouse model, outperforming unregulated and more weakly regulated constructs (**Figure 4**). This work benefited from a brain-targeting systemic AAV capsid, Cap.B22 (*50*), which allows higher viral titers because it avoids delivery to the liver, where toxicity has been observed in previous studies (*23*). However, comparison between unregulated and regulated circuit variants shows that the circuit provides additional benefit beyond that achieved by capsid targeting alone. The synthetic miRNA IFFL circuits could thus improve gene therapy for Rett syndrome, and potentially other genetic disorders as well.

In gene therapy for Rett syndrome and other diseases, expression level is critically important. The single-molecule analysis employed here enabled us to quantitatively compare ectopic and endogenous expression levels in the same cell. This showed that a moderately regulated circuit (“1x”) closely matched endogenous mRNA levels in the median cell, but also generated a subpopulation of cells with much higher levels (**Figure 3C**). By contrast, a tightly regulated circuit (“4x”) exhibited expression lower than endogenous *Mecp2* in most cells but also avoided appreciable overexpression. The superior performance of the tightly regulated circuit in rescuing the Rett phenotype (**Figure 4B**) likely reflects this lack of high expression. Additionally, we note that both circuits reduced expression in the brain to levels far lower than that of a CAG-GFP construct, which used a more typical synthetic promoter (**Figure 3B**).

This regulated *Mecp2* gene therapy has some limitations: it did not improve the survival of male *Mecp2*-null mice and did not affect the obesity phenotype of heterozygous females. This may be due to our choice of a central nervous system (CNS)-targeting capsid. MeCP2 expression outside the CNS may be important for development and survival. Several Rett phenotypes in humans, including breathing irregularities, cardiovascular dysfunction, and decreased pain sensitivity (*54*), implicate the peripheral nervous system (PNS). Future studies applying the gene therapy to the PNS and other tissues with targeted AAVs (*55*) could help to improve efficacy and clarify the role of MeCP2 in non-CNS cells. Additionally, the IFFL circuits introduced here do not incorporate post-transcriptional or dynamic regulation that may occur in natural MeCP2 regulation. However, the distribution of endogenous *Mecp2* mRNA appeared relatively uniform and stable over time (**Supplementary Figure 7**), suggesting that such effects may be modest in the brain.

The simplicity and compactness of the IFFL circuit make it potentially broadly useful for AAV-based gene therapies for other genetic diseases, such as SMA, Angelman syndrome, fragile X syndrome, and monogenic autism spectrum disorders (*56*, *57*). The dosage compensation circuit could also extend the durability of gene therapies, by permitting higher initial vector copy numbers, and therefore a longer duration of therapeutic protein expression, while avoiding overexpression, in dividing, diluting cells. The miRNA-based circuits used here could also improve safety and efficacy of other therapies based on ectopic expression, such as gene editing. More generally, these results suggest that synthetic circuits could play a vital role in gene therapies and other emerging therapeutic approaches.

## Methods

### Modeling an idealized dosage-compensating IFFL

To understand the functional behavior of the IFFL circuit, we considered a simple model of IFFL-regulated expression, in which a gene produces both a mRNA gene product, *G*, and its negative regulator (miRNA), *R*, which acts catalytically to degrade the target mRNA. We assume both species are removed through a combination of degradation and dilution with rate constants of γ_G_ and γ_R_, respectively. We denote the expression level of *G*, in units of molecules per cell, produced by a single unregulated gene copy as *P*. This means that at steady state, *P* molecules are produced per cell during the mean life-time, τ = γ _G_^−1^, of the gene product. In other words, the production rate for a single copy of the gene is *P*/τ = *P*γ_G_ molecules per cell per unit time. When there are multiple copies of the gene, we introduce *D* to denote the gene dosage, i.e. the number of gene copies per cell. With *D* gene copies, the production rate is simply *DP*γ_G_. We also assume that *R* is produced proportionally to *G*, with α being the stoichiometric ratio of *R* to *G* production rates. Finally, we assume that the miRNA catalytically degrades *G* at a rate *kGR*. This assumes that the gene product is far from saturation, i.e. *G* ≪ *K_M_*, where *K*_M_ denotes the Michaelis constant for miRNA-mediated mRNA degradation. Together, these reactions can be described by a pair of ordinary differential equations:

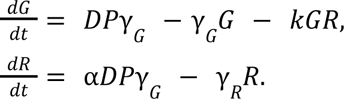

Solving for the steady state of this system, 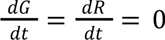, gives

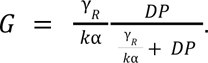

When *DP* is much less than γ_R_/*k*α, the circuit has a negligible effect, and the steady-state expression level, *G*, is approximately equal to *DP*, the unregulated expression level. When *DP* approaches or exceeds γ*_R_*/*k*α, expression approaches a constant limiting value, *G*_*_ = γ_R_/ *k*α, insensitive to both *D* and *P*. The ideal IFFL thus caps expression at the value γ_R_/*k*α. This cap can be decreased by strengthening the catalytic rate of the catalytic repression (increasing *k*), by decreasing the repressor degradation rate γ_R_, or by increasing the stoichiometric ratio α.

### Stochastic simulations

To simulate the miRNA IFFL circuit in the presence of Poissonian viral uptake and high ectopic promoter strength, we assumed the same reactions as the ODE model above. In place of D, we assume a given cell has *N* viral genomes, where *N* is drawn from a Poisson distribution. To accurately simulate bursty transcription, of these *N* genomes, we assume *n*≤*N* copies are in a transcriptionally active state at any given time, while the remaining *N* − *n* copies are off (silent). We assume individual viral copies switch from the off to the on state at a rate of *l* and from on to off at a rate of µ. The total set of reactions is as follows:

| Reaction | Rate |
| --- | --- |
| $(G, R) \rightarrow (G + 1, R + 1)$ | $nD\gamma_G\mu/l$ |
| $G \rightarrow G - 1$ | $\gamma_G$ |
| $R \rightarrow R - 1$ | $\gamma_R$ |
| $G \rightarrow G - 1$ | $kRG$ |
| $n \rightarrow n + 1$ | $(N - n) l$ |
| $n \rightarrow n - 1$ | $n\mu$ |

The code^1^ conducts a stochastic (Gillespie) simulation of this system (*58*). The code was run for 2,000,000 samples for each of the distributions in the following table. All rates are assumed to be per hour.

| Condition | $D$<br>(#) | $\gamma_G$<br>($hour^{-1}$) | $\gamma_R$<br>($hour^{-1}$) | $l(hour^{-1})$ | $\mu(hour^{-1})$ | $k$<br>($\#/hour^{-1}$) | Mean $N(\#)$ |
| --- | --- | --- | --- | --- | --- | --- | --- |
| Endogenous | 10 | 1/8 | 1/24 | 2.5/8 | 20 | 0 | 1 (fixed) |
| Low transduction<br>(unregulated) | 100 | 1/8 | 1/24 | 1 | 1 | 0 | 0.69 |
| Medium transduction<br>(unregulated) | 100 | 1/8 | 1/24 | 1 | 1 | 0 | 3 |
| High Transduction<br>(unregulated) | 100 | 1/8 | 1/24 | 1 | 1 | 0 | 9 |
| Low Transduction<br>(regulated) | 100 | 1/8 | 1/24 | 1 | 1 | 1/160 | 0.69 |
| Medium Transduction<br>(regulated) | 100 | 1/8 | 1/24 | 1 | 1 | 1/160 | 3 |
| High Transduction<br>(regulated) | 100 | 1/8 | 1/24 | 1 | 1 | 1/160 | 9 |

### Transient transfection

For imaging experiments, glass bottom plates were coated overnight by diluting human laminin (ThermoFisher #A29249) to a surface density of 0.5 µg/cm^2^. The next day, U2OS cells were seeded at a density of 50,000 cells in each well of a 24 well plate, either Falcon Polystyrene Microplates (Fisher Scientific #08-772-1) for flow cytometry or MatTek No 1.5 uncoated glass bottom plates (Fisher Scientific #NC9988706) for smFISH imaging experiments, and cultured under standard conditions overnight. The following day, the cells were transiently transfected using Fugene HD (Promega #E2311), according to the manufacturer’s protocol, and incubated for 48 hours before analysis.

### Flow cytometry

Cells in a 24-well plate were trypsinized with 50 μL of 0.25% Trypsin (ThermoFisher #25200056) for 5 minutes at 37 °C. Next, 150 μL of HBSS with 2.5 mg/ml BSA and 1 mM EDTA was used to wash down the cells. Cells were then filtered through a 40 μm cell strainer (Fisher Scientific #08-771-1) into a 96-well round bottom plate (Fisher Scientific #08-772-17). This plate was then analyzed with a CytoFLEX flow cytometer.

### In vitro smFISH

To analyze *Mecp2* expression in brain slices, we used a previously published smFISH protocol (*49*).

*Probe Design*: 27 primary probes were constructed by tiling the EGFP segment of the *Mecp2-EGFP* RNA, extending the length of each probe until the melting temperature reached 42 Celsius in the presence of 50% formaldehyde according to the MeltingTemp utility in the BioPython package. Each probe was then flanked with two copies (one each side) of a readout sequence ‘GCGCAATAAACCCTA’ separated by a ‘TT’ element. Readout probes complementary to the readout sequence were ordered separately, conjugated to Alexa594. For visualizing the cytoplasm, oligoDT (30 ‘T’ nucleotides) was conjugated to Alexa Fluor 405.

*Protocol:* Cells were washed twice with PBS before fixation with 4% formaldehyde for 10 minutes at room temperature. Cells were then washed twice with PBS and then permeabilized with cold 70% ethanol and incubated at 4 °C for 1 hour. The ethanol was then aspirated and the cells were allowed to air dry. The cells were then washed 3 times with 2xSSC and incubated overnight at 37 °C with 50% hybridization buffer (5 mL formamide, 4 mL Ultrapure RNase/DNase free water, 1 mL 20x SSC, and 1 g of high MW dextran sulfate (Sigma-Aldrich #D8906) with 10 nM of smFISH probes. The next day, the cells were washed 4 times with 30% wash buffer (3 mL formamide, 5.9 mL Ultrapure RNase/DNase free water, 1 mL 20xSSC, and 0.1 mL Triton-X) at 37 °C and then once with 4x SSC at room temperature. To add the readout probes, 50 nM readout probes in 10% EC Buffer (1 mL of ethylene carbonate, 7 mL Ultrapure RNase/DNase free water, 2 mL 20x SSC, and 1g Low MW dextran sulfate Sigma #D4911) was added to the cells and incubated for 20 minutes at room temperature. After, the cells were washed 3 times with 10% wash buffer (1 mL formamide, 7.9 mL Ultrapure RNase/DNase free water, 1 mL 20xSSC, and 0.1 mL Triton-X) for 5 minutes at room temperature. Then, the cells were washed briefly with 4xSSC, then 2xSSC, and then the cells were incubated with 500nM oligoDT-Alexa405 in 2xSSC for 1 hour at room temperature. Next, the cells were washed with 2xSSC twice, and then twice with antibody staining buffer (50 mL PBS, 0.5 g Ultrapure BSA, 1.13 g glycine, 50 µL Tween 20). Then, the cells were incubated with antibody staining buffer for 30 minutes for blocking. Then, the cells were incubated with 0.5 µg/mL Alexa-Fluor 647 conjugated Na/K ATPase antibody (Abcam #ab196695) in antibody staining buffer for 2 hours at room temperature. Finally, the cells were washed twice with antibody staining buffer, and twice with 2xSSC, and mounted in 50 µL of Prolong Gold (ThermoFisher #P10144) before imaging.

### Bulk RNA sequencing

#### Sample preparation and sequencing

To assess off-target effect of the miRNA cassette, U2OS cells were plated on 6-well plates with 300,000 cells per well. Cells were transfected the following day with 1,000 ng of either the control plasmid or the BFP-miR-L plasmid using Fugene HD (Promega #E2311) according to the manufacturer’s instructions. Media was replaced with 2 mL of fresh media 24 hours post-transfection. Cells were harvested 48 hours post-transfection by digestion with 0.25% Trypsin-EDTA, centrifugation at 300 g for 5 minutes, and removal of the supernatant by aspiration. The cell pellet was stored in −80 °C prior to the purification. RNA was extracted using the RNeasy kit (Qiagen #74106) according to the manufacturer’s instructions. RNA was treated with Turbo DNase (ThermoFisher #AM2238) and purified using the RNeasy kit RNA cleanup protocol. mRNA sequencing libraries were prepared by Novogene.

### Differential gene expression analysis

To characterize the perturbations that synthetic miRNA brought to the endogenous transcriptome, differential expression analysis was performed using DESeq2 (1.40.1) (*59*) in R (4.3.1), comparing transcript counts in miRNA-transfected cells and BFP-only cells. Preprocessing and further analysis of RNA expression was performed as in ref. (*60*).

### AAV production and purification for in vivo assessment of constructs

Plasmids used for AAV preparation include the single stranded (ss) rAAV genomes containing the cassettes described in the main text, pHelper (Addgene) and plasmids encoding AAV-CAP.B22 ((*50*, *61*). Viruses were prepared through triple-transient transfection in adherent HEK293 cell culture (ATCC) and purified by ultracentrifugation as previously described (*62*). Viruses were concentrated in sterile saline for injection into rodents, and viral titers were measured by qPCR.

### Animals

All experiments involving animals were approved by the Institutional Animal Care and Use Committee (IACUC) and Office of Laboratory Animal Research (OLAR) at the California Institute of Technology.

The animals used in this study are the Rett syndrome model commonly known as *Mecp2*-null (Jackson Laboratories 003890). The colony was maintained on a C57Bl/6J background for compatibility with the engineered AAV capsids used in this study. Mice in behavioral cohorts were group housed when possible at 71-75℉ under a reverse light cycle (12h on, 12h off) with enrichment materials such as blocks and shepherd shacks.

### Intravenous AAV administration and behavioral assessment in *Mecp2*-null mice

To assess the performance of the AAV-MeCP2 candidate constructs against the disease phenotype of the *Mecp2*-null rodent model of Rett Syndrome, animals were genotyped at week 3 of life and randomized into groups controlled for sex, age and breeding pair origin. Mice in treatment groups were injected intravenously through the retro-orbital vein as previously described (*62*). Heterozygous females and hemizygous males were divided into balanced cohorts to receive no injection (n=12; 6 male and 6 female), or were dosed by weight to receive 1e14 viral genomes/kilogram of the AAV-CAP.B22-MeCP2 unregulated (n=13; 5 male and 8 female), 1x (1 target site; n=13; 6 male and 7 female) or 4x (4 target sites; n=11; 5 male and 6 female) *Mecp2* therapeutic constructs. These mice, along with their wildtype littermates, were assessed at baseline for motor and neurological performance using the RTT Phenotype Scoring Scale as previously described (*21*), and re-assessed every two weeks. Testing continued until the study end at 18 or 24 weeks post injection, or until humane endpoint criteria were reached, resulting in withdrawal of an animal from the study. All behavioral scoring was done individually in a large plastic arena outside of the home cage, and was performed during the dark phase under red light. We observed a phenotype increase earlier than some studies (*21*, *53*), but consistent with timing observed in others (*63*).

### *Mecp2* RNA histological analysis

Following 3 weeks of expression, injected rodents were deeply sedated via intraperitoneal injection of Euthasol (pentobarbital sodium and phenytoin sodium solution, Virbac AC) prior to cardiac perfusion using RNAse-free, heparinized saline and 4% paraformaldehyde (PFA) in 0.1M phosphate buffered saline (PBS). Tissues were post-fixed in 4% PFA for 24-48 hours and either sliced on a vibratome at 50 µM for immediate analysis or cryoprotected with RNAse-free 10% and 30% sucrose solution, frozen in OCT and stored at −80°C. Prior to analysis, tissues were sliced to a thickness of 50 µM on a Leica cryostat. Tissue slices collected for FISH analysis were incubated in ice cold RNAse-free 70% ethanol prior to probing.

### Hybridization chain reaction

*Probes and Buffers:* Probe sets were ordered from Molecular Technologies (https://www.moleculartechnologies.org/) against the coding sequence of EGFP and endogenous mus Musculus *Mecp2* isoform 1 (NM_001081979), with specific instructions to only include the 3’ UTR for the latter. Molecular Technologies hybridization buffer, wash buffer, and amplification buffer were included as part of the order.

*Protocol*: Paraformaldehyde-fixed fresh or frozen brain tissue was sliced to a thickness of 50µm, mounted onto glass coverslips and dried in a fume hood. Slices obtained from frozen brain tissue were rinsed with RNAse-free PBS to remove OCT compound prior to drying. Ice cold RNAse-free 70% ethanol was applied to the dried tissue slices. After incubating at 4 °C for 1 hour, the ethanol was removed and samples were dried in a fume hood. Once dry, the coverslips were washed with PBS before incubating in 8% SDS in PBS for 20 minutes at room temperature. The SDS solution was then poured off and the coverslips were rinsed 3 times with PBS for 2 minutes. The area of the coverslips around the tissue slices was then dabbed dry with a Kimwipe before mounting a SecureSeal hybridization chamber (Grade Biolabs #621502) on top of the tissue slices. Using the hybridization chambers, the samples were washed with 5xSSCT (5xSSC with 0.1% Tween 20) and then the samples were pre-incubated with 30% HCR probe hybridization buffer for 30 minutes at 37 °C. After this incubation, the tissue slices were incubated with 0.2 µL of 2 µM stock of each of the odd and even HCR probe mixtures for both ectopic and endogenous target in 100 µL 30% HCR probe hybridization buffer for 3 days at 37 °C. After, the tissue slices were washed 4 times with 30% HCR probe wash buffer at 37 °C for 15 minutes. Then the tissue slices were washed twice with 5xSSCT for 5 minutes at room temperature. Next, the tissue slices were pre-incubated with HCR amplification buffer for 30 minutes at room temperature. While this was happening, for each of the 3 µM stocks of hairpins H1 and H2 for both endogenous and ectopic targets (4 total), 2µL was added to separate PCR tubes (4 total) and heated to 95 °C for 90 seconds before being allowed to cool to room temperature in a dark drawer for 30 minutes. After this was completed, the 2 µL from each of the PCR tubes was added to 100 µL of amplification buffer and the tissue slices were incubated in this solution for 90 minutes at room temperature in the dark. Once complete, the samples were washed 4 times for 15 minutes with 5xSSCT. The samples were then washed once with 2xSSC, and then the slices were incubated with 500nM oligoDT-Alexa405 in 2xSSC for 1 hour at room temperature. The samples were then washed 4 times with 2xSSC, and finally mounted in Prolong Gold (ThermoFisher #P10144).

### Cell segmentation

Cell segmentation was performed using the Cellpose program (https://www.cellpose.org/) (*64*, *65*) applied to the oligoDT signal (**Figure 3**) or a Na/K ATPase membrane marker (**Figure 2**). The basic ‘cyto’ model proved sufficient to accurately segment cell bodies in the vast majority of cases. The masks generated by Cellpose were then passed to the next parts of the program.

### Dot counting algorithm

In Figures 2D, 3B, 3C, 4E, and 4F, dots representing single transcripts were detected using the following algorithm. First, each fluorescence image was thresholded using a minimum intensity parameter (see below). Next, the local maxima of the images were identified. Maxima that overlapped with a mask created by Cellpose were labeled as dots inside that cell.

In brain samples, autofluorescence also exhibited dots. However, the broader spectrum of autofluorescence caused these to be detected in multiple fluorescence channels, unlike dots associated with specific transcripts, which predominantly appeared only in a single fluorescence channel. To avoid miscategorizing these as transcripts, any dots that occurred at the same point in the 594 and 640 channels were labeled as autofluorescence background and discarded.

To select for cells that were positively transduced with the viral construct, we sorted cells by total GFP signal, and took the top 100 cells in each case.

Images of these cells are included as supplemental PDF files (**Supplementary material**), each row of which depicts an individual cell. The oligoDT images depict a maxproject of the oligoDT signal within the z-range of the cell boundaries, with a white outline depicting the segmentation boundary at the z-level of the centroid of the segmented cell. For HCR, images depicting endogenous and ectopic transcripts are next, followed by an overlay of both. In these images, masks are taken at each z-level of the segmented cell, and then a max-project of the masked z-slices is taken. Dots in each channel are denoted by white circles, while background dots are denoted by cyan circles. Autofluorescent puncta are not true dots, and show up in the overlay. Some dots are missed either due to being too dim, or due to having their local maxima outside the segmentation boundary (**Supplementary Figure 8**). The receiver-operating-characteristic (ROC) analysis (below) showed that there will be a small false negative rate.

### Optimizing dot counting threshold

The accuracy of the dot counting method was sensitive to the fluorescence threshold. We therefore sought to optimize the threshold value by comparison to a ground truth. To establish a ground truth data set, we manually analyzed 4 cells expressing the unregulated construct and 10 negative control cells for which no fluorescent secondary probes were hybridized. We then ran the image analysis pipeline on these samples, scanning the threshold value from a pixel intensity of 100 to 1100. The number of dots identified at each threshold value was compared to the ground truth values. After scanning the threshold, we applied a receiver-operating-characteristic (ROC) analysis to identify an optimal threshold value (**Supplementary Figure 9A, 9C**). More specifically, we compared the number of true positives and false positives for each threshold. Taking P to be the ground truth number of positive dot calls, TP as the number of true positive dots identified by the algorithm, and FP as the number of false positive dots, we searched for the threshold value that minimized (P - (TP + FP))^2^, i.e. the squared error in the total dots called (**Supplementary Figure 9B, 9D**).

### Correcting dense dot counts

In some cells, high densities of HCR dots made it impossible to distinguish individual dots. Because of this, the dot counts saturated, so that even if the total dot fluorescence in the cell increased, there was no increase in dot count. To correct for this regime, we corrected dot counts if they were past the threshold of fluorescence beyond which there was little change in dot count, and assigned them a count that was dependent on a linear function of the total dot fluorescence (**Supplementary Figure 3**). To get this linear function, we find the median dot count *n*_0_ of cells with fluorescence close to the threshold *f*_0_, and beyond that threshold assign cells an adjusted count *n* as a linear function of their observed fluorescence, *f*:

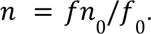

### Bootstrap significance testing

To test the statistical significance of the difference between the means of two sets of samples, we followed a standard bootstrap hypothesis testing procedure. Briefly, samples from both groups were pooled into a single set. Two new sets with the same size as the original sets were then constructed by sampling with replacement from the pooled set and the means of the resampled sets were compared. This was repeated 10^5^ times, and the proportion of resampled set pairs with means that differed by more than the observed difference of the original pair was reported as the p-value.

## Supporting information

CAG-GFP_cell_segmentation

Unregulated_cell_segmentation

1x_cell_segmentation

4x_cell_segmentation

## Declaration of Interests

A patent has been filed by the California Institute of Technology related to this work (US application number 17/100,857). M.B.E. is a scientific advisory board member or consultant at TeraCyte, Primordium, and Spatial Genomics.

## Author Contributions

M.J.F. and M.B.E. conceived the project. M.J.F. identified the incoherent-feedforward-loop motif, performed mathematical modeling, and implemented the circuit using miRNA inside an AAV gene expression cassette. M.J.F. validated the circuit in cell culture using flow cytometry, smFISH, and HCR, wrote the image analysis pipeline, and compared endogenous and ectopic expression of MeCP2 in the mouse brain. M.B.E. provided guidance in circuit design and data analysis. A.M.M. and V.G. selected viral capsids and designed the mouse behavioral study. A.M.M. prepped and injected virus, and performed mouse tissue preparation. A.M.M established Rett model mouse colonies, and performed mouse behavioral studies, including time-course of behavioral phenotyping. R.D. performed mRNA sequencing in response to miRNA expression. M.J.F. and M.B.E. wrote the manuscript with input from all authors. M.B.E. supervised the in vitro work and V.G. supervised the in vivo work. M.B.E. and V.G. funded the project.

## Acknowledgements

We thank Mitch Gutman for informing us about the overexpression problem for Rett syndrome gene therapy, Inna Strazhnik for graphics, Rana Eser for assistance in tissue handling and animal collections, Martin Tran and Yodai Takei for advice on smFISH and HCR experiments and data analysis, and members of the Elowitz and Gradinaru labs for other input and feedback.

## Funding

This work was funded by grants from the Rett Syndrome Research Trust, the Rosen Bioengineering Center, and the Merkin Institute for Translational Research. Research reported in this publication was also supported by the National Institute Of Biomedical Imaging And Bioengineering of the National Institutes of Health under Award Number R01EB030015 and NIH Pioneer DP1OD025535 and U24MH131054. The content is solely the responsibility of the authors and does not necessarily represent the official views of the National Institutes of Health.

---

**Supplementary Figure 1.**
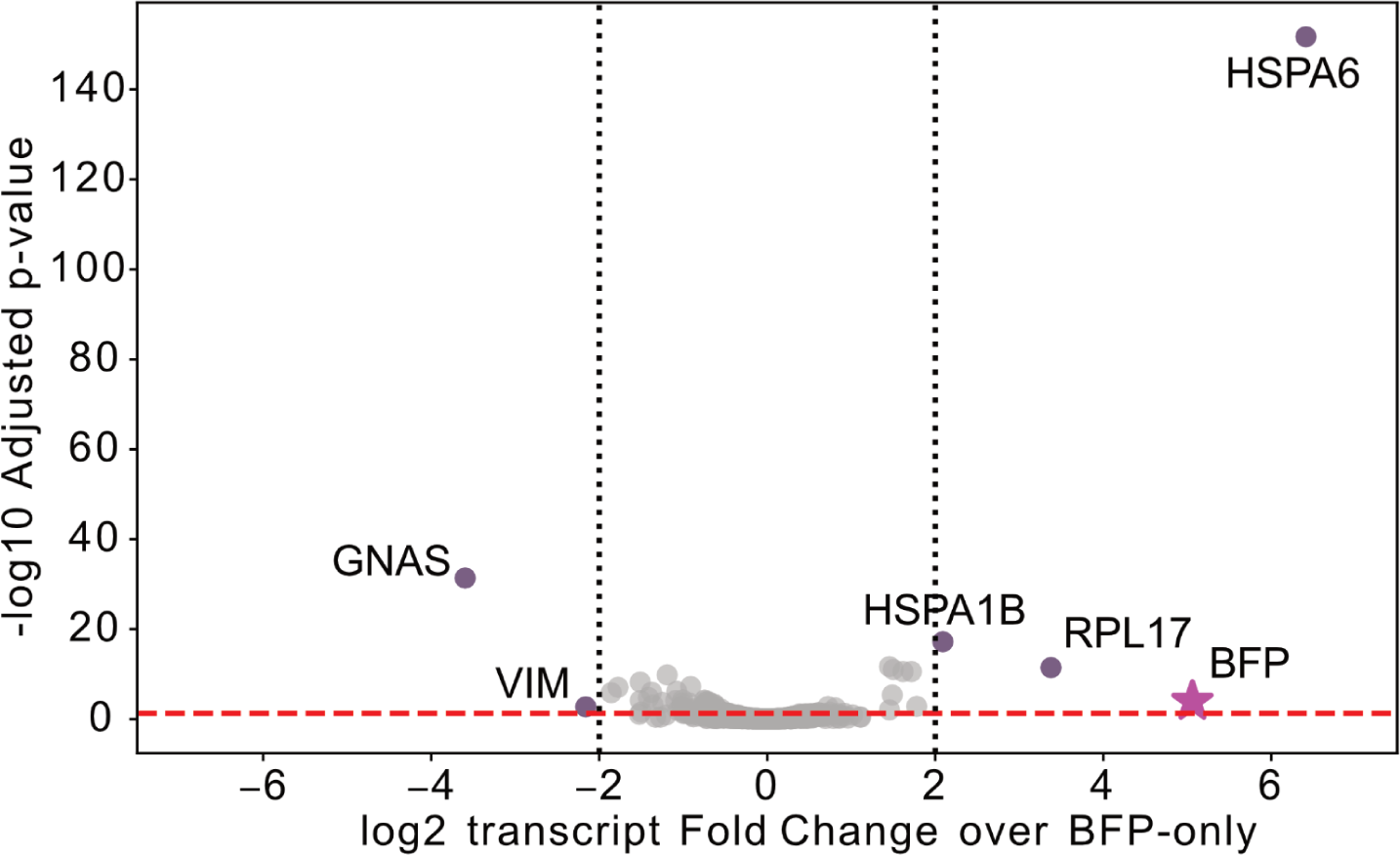
Volcano plot of bulk RNAseq analysis of the effects of miRNA expression on the transcriptome. In order to identify potential off-target effects of the miRNA, we performed bulk RNA sequencing on cells transfected with mTagBFP2-miRNA or with a negative control plasmid lacking the miRNA. The dashed black lines indicate where the absolute value of the log_2_ fold change exceeded a value of 2. The red dashed line indicates where the adjusted p-value is less than 0.05. A handful of genes showed significantly differential expression (labeled purple dots) relative to the construct without miRNA, and are mostly heat shock proteins which may have been induced due to stress from the higher transfection level of the BFP-miRNA cells, as indicated by the higher expression of BFP (star). No upregulated or downregulated transcript contained target sites for the miRNA.

**Supplementary Figure 2.**
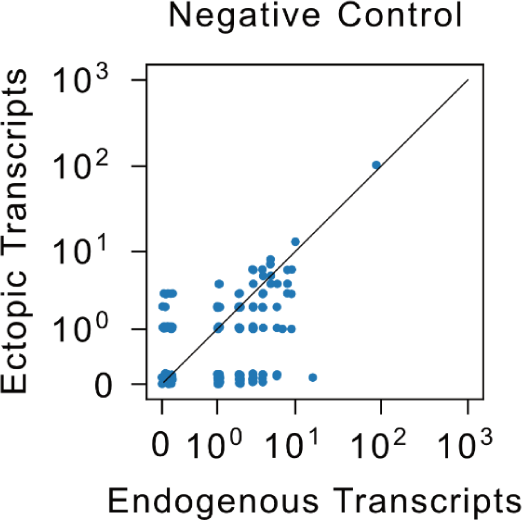
Measured background (negative control) transcript counts without primary probes. The no-primary negative control had no probes against *Mecp2* added and thus no true HCR dots. Ideally, the dot counting pipeline would identify 0 dots in all cells. Many cells are indeed close to the bottom left, with 48% of cells having either 1 or no counts in both the ectopic and endogenous channels.

**Supplementary Figure 3:**
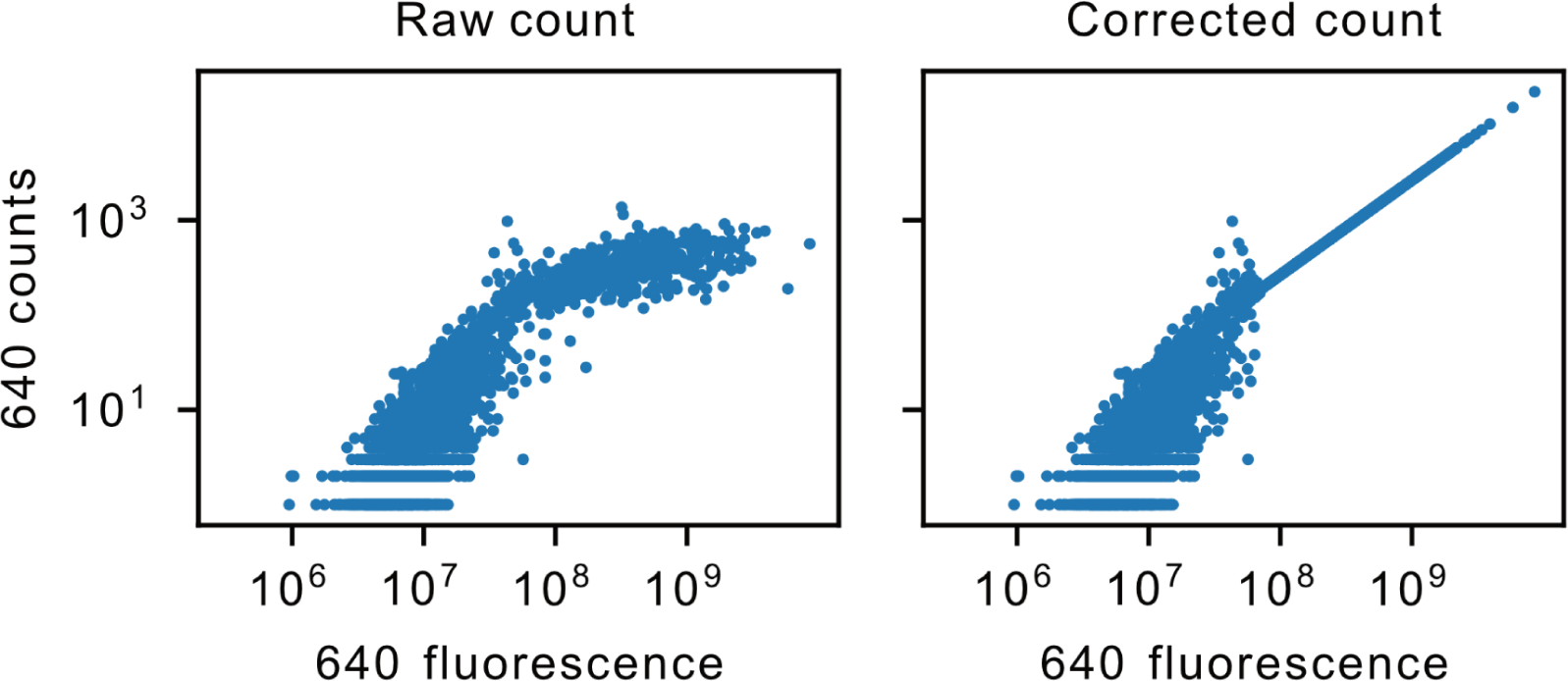
Dense dot count correction. Counted dots in individual cells as a function of total dot fluorescence both before (‘Raw count’) and after (‘Corrected count’) correcting for highly dense cell dots (**Methods**).

**Supplementary Figure 4.**
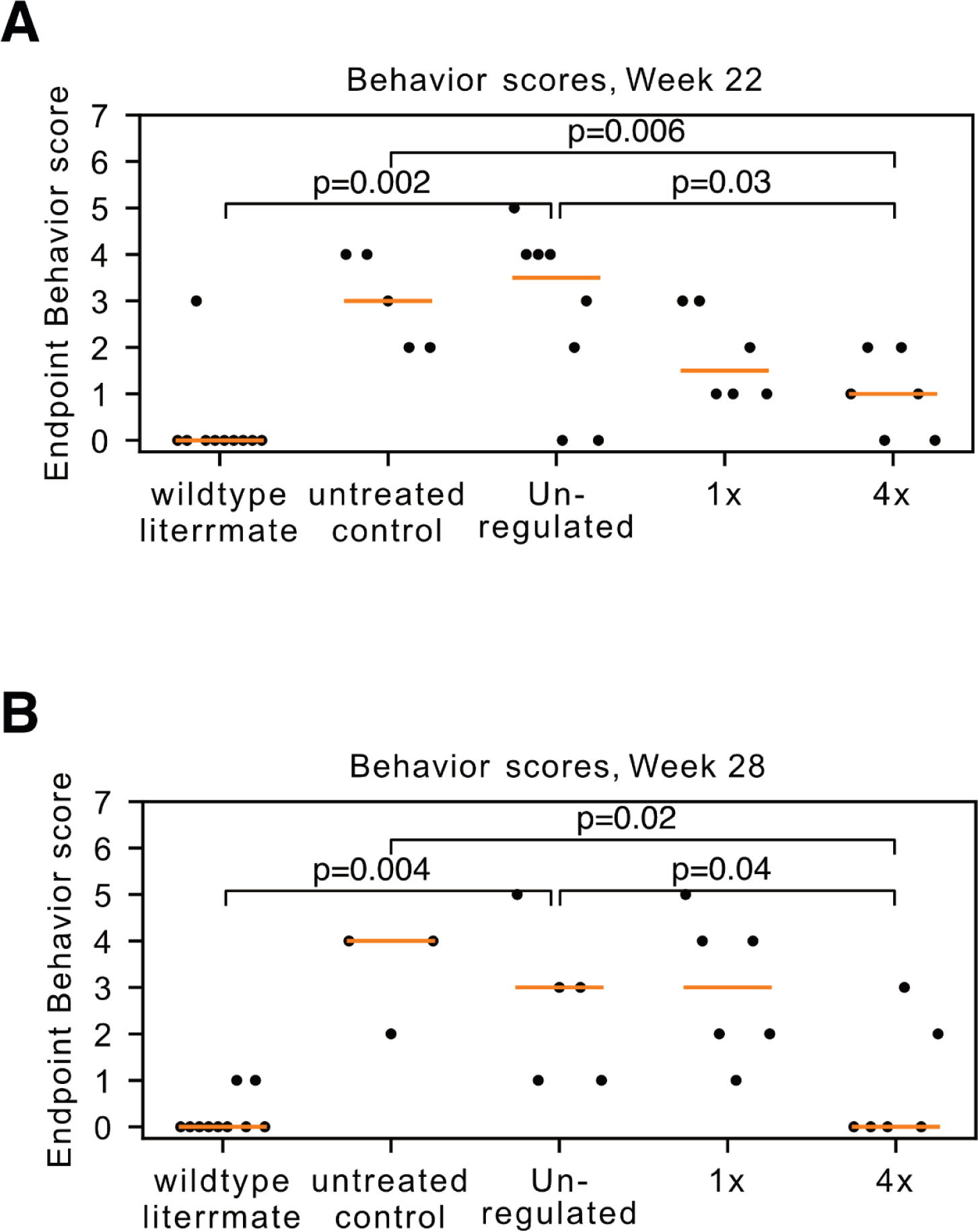
Endpoint behavior score statistics. **(A)** Endpoint Rett behavior scores at week 22. Mice that received the 4x construct scored significantly lower on average than both the uninjected mice (bootstrap p=0.006) and the mice that received the unregulated construct (bootstrap p=0.03). The standard deviation of the 4x group’s scores, 0.82, was significantly lower than the standard deviation of the unregulated group’s scores, 1.79 (bootstrap p=0.03). Wildtype littermates scored significantly lower than the uninjected controls (bootstrap p=0.001) and mice that received the unregulated construct (bootstrap p=0.002). The wildtype and 4x mice were not significantly different (bootstrap p=0.08). **(B)** Endpoint Rett behavior scores at week 28. Mice that received the 4x construct scored significantly lower on average than both the uninjected mice (bootstrap p=0.02) and the mice that received the unregulated construct (bootstrap p=0.04). The standard deviation of the 4x scores, 1.21, was not significantly lower than the standard deviation of the unregulated scores, 1.49 (bootstrap p=0.29). Wildtype littermates scored significantly lower than the uninjected control (bootstrap p=0.002) and mice receiving the unregulated construct (bootstrap p=0.004). The scores of wildtype and 4x mice were not significantly different (bootstrap p=0.08).

**Supplementary Figure 5.**
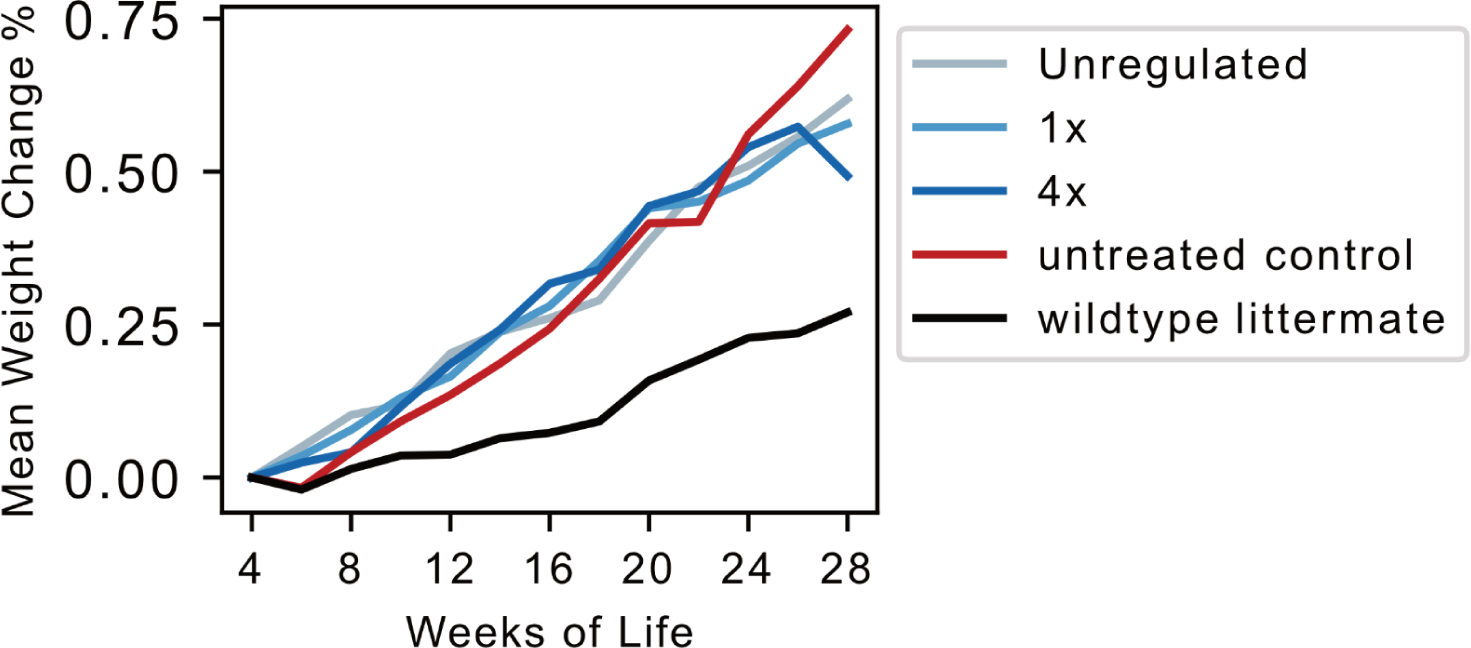
Mouse weight was not affected by AAV-MeCP2 gene therapy. Mouse weight percentage change over time for each cohort. The wildtype mouse cohort had a reduced weight relative to other cohorts, which increased similarly over time, indicating that the obesity phenotype was not affected by AAV-MeCP2 gene therapy.

**Supplementary Figure 6.**
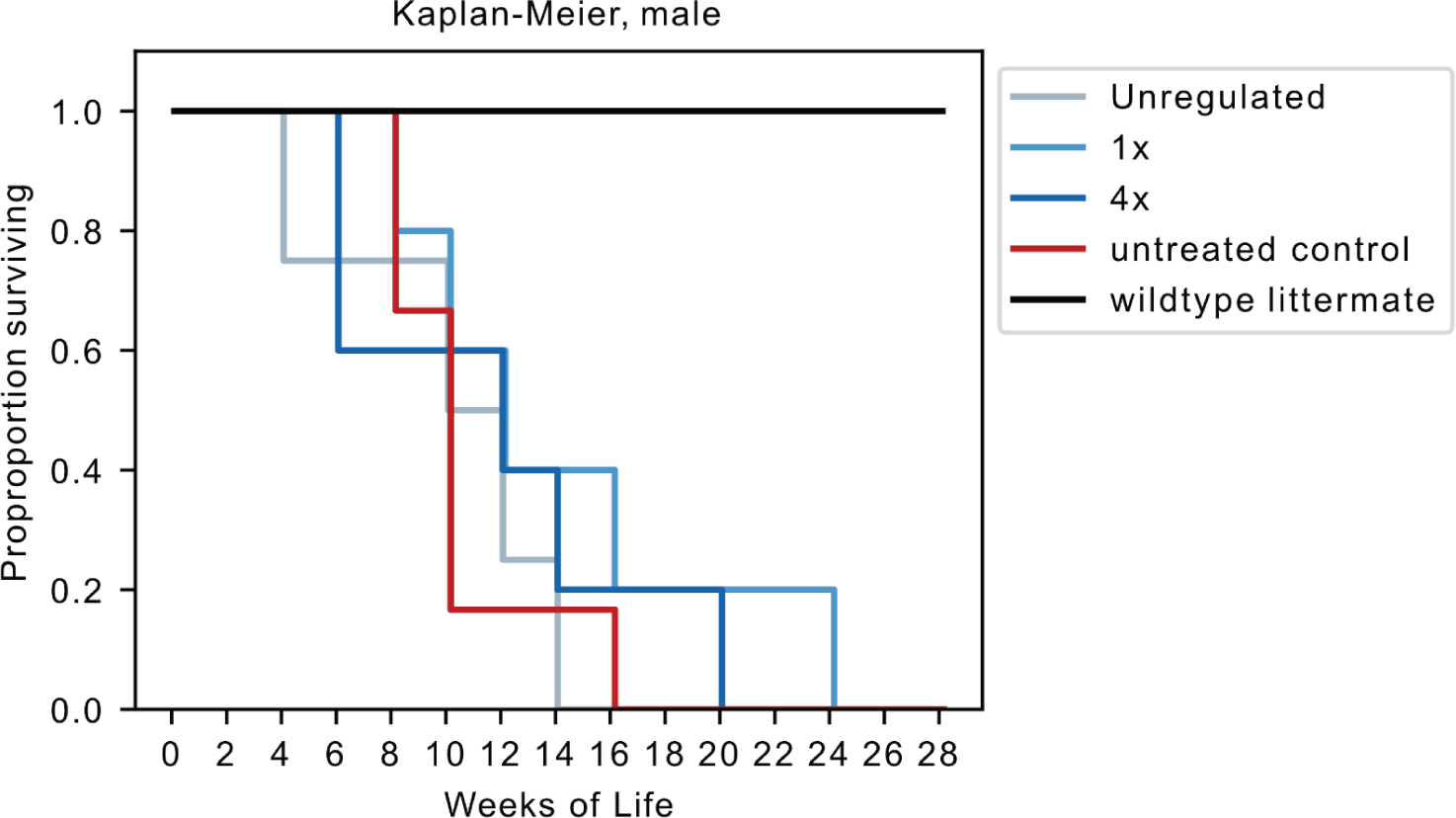
Male survival was not affected by AAV-MeCP2 gene therapy. Kaplan-Meier survival curves show no significant change in lifespan in male Rett mice in any treatment group.

**Supplementary Figure 7.**
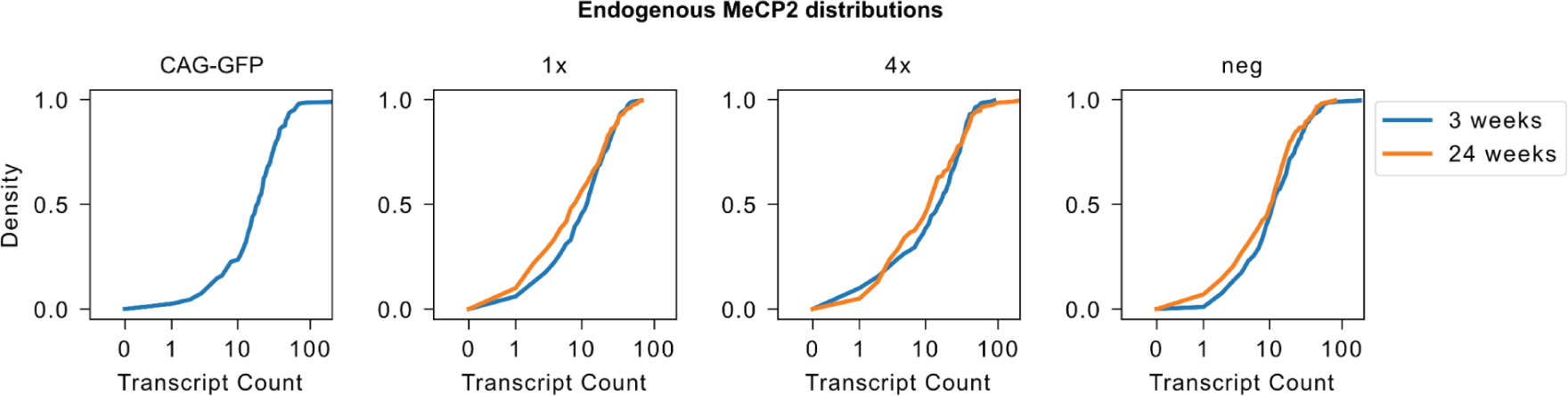
Endogenous MeCP2 distribution after 3 and 24 weeks of viral expression. Empirical cumulative distributions of endogenous *Mecp2* in mice that received different viral constructs were remarkably consistent between constructs and between 3 and 24 weeks of expressing the viral constructs.

**Supplementary Figure 8:**
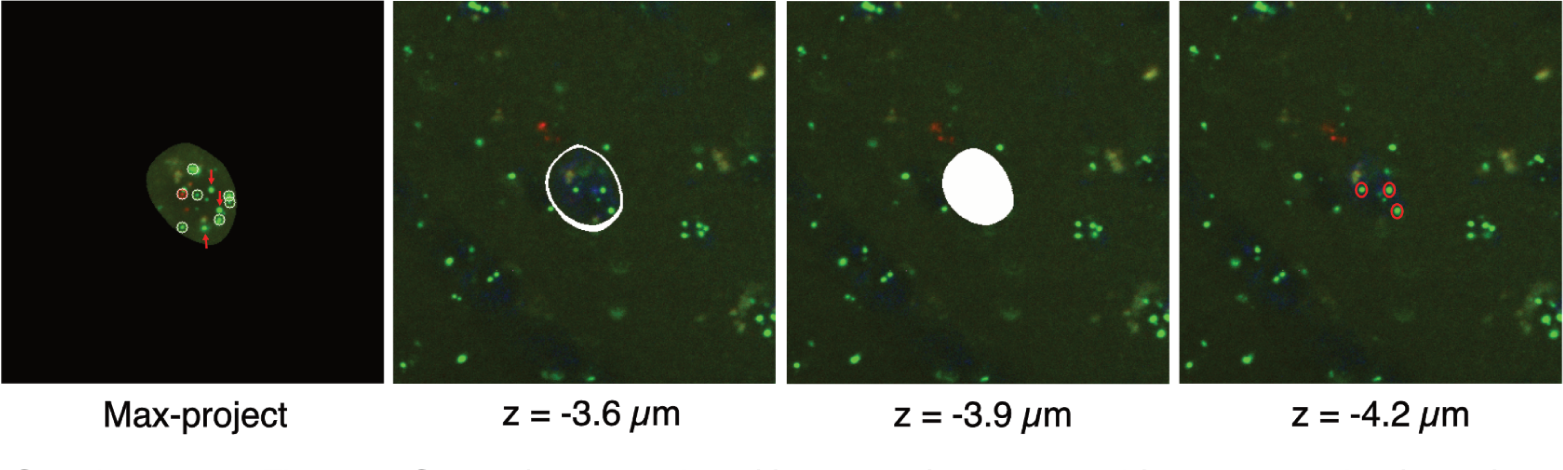
Limitations of dot detection. In the supplementary PDFs, some dots appear to not be counted (max-project, red arrows are missed dots). In some of these cases, this is due to the maxima of the dots being below (in the z direction) the cell, but not inside of it. Z-levels −3.6 µm, −3.9 µm, and −4.2 µm are displayed here. The solid white mass indicates the “bottom” of the detected cell at slice z = −3.9 µm. However the maxima of the missed dots are at z = −4.2 µm or below.

**Supplementary Figure 9.**
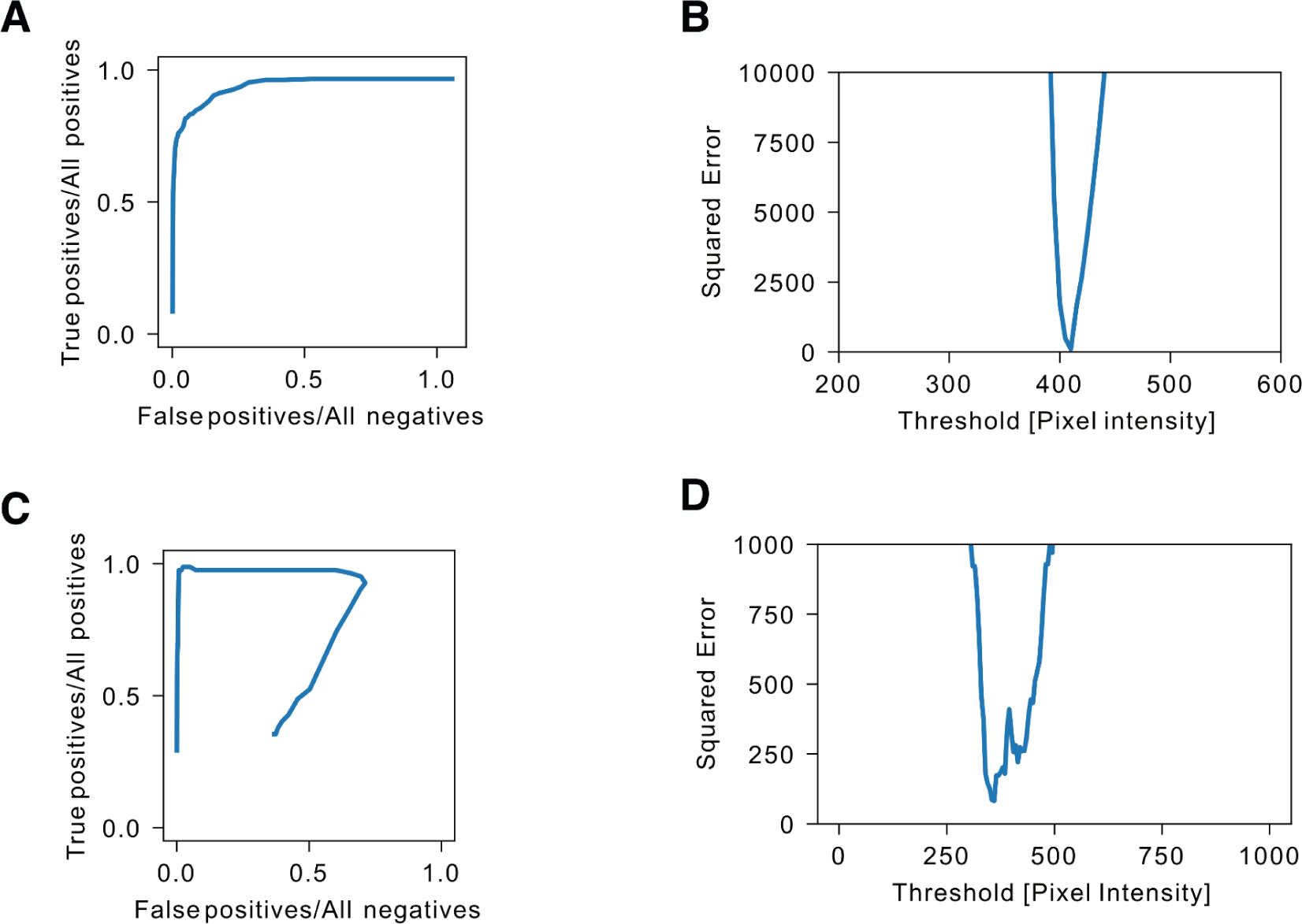
Receiver operating characteristic optimization. **(A)** True positive rate vs false positive rate as a function of the dot counting threshold, which increases from 100 (top right) to 1100 (bottom left), in U2OS cells analyzed with smFISH. True positives and false positives were defined relative to manually annotated ground truth images. An ideal detection algorithm would count all the true positives and no false positives, i.e. would lie in the upper left corner. Some thresholds give results close to this ideal corner, and define the optimal values of the threshold, but perfect classification is not possible. **(B)** The squared difference between the total detected counts and the total true counts as a function of threshold, in data from (A), showed a narrow window of acceptable thresholds that give a low error around a threshold pixel intensity of 410. **(C)** True positive rate vs false positive rate as a function of the dot counting threshold, which increases from 100 (blue curve, top right) to 1100 (blue curve, bottom left), in mouse brain slices analyzed with HCR. The true positive and false positive rate were defined relative to manually annotated ground truth images. In contrast to the smFISH data, a significant component of the background consisted of autofluorescent puncta prevalent in the brain. To correct for this, false positive dots were eliminated by discarding dots that appeared in both channels. This led to a decrease in true positive rate and false positive rate at low thresholds, leading to the curve away from the top right. Some threshold values closely approached a true positive rate of 1 and false positive rate of 0 (upper left corner). **(D)** The squared difference between the detected counts and the total true counts as a function of threshold in data from (C), showed a window of threshold that had minimal error around a pixel intensity of 360.

## Footnotes

1 https://github.com/labowitz/DosageCompensatedGeneTherapy/blob/main/Fig1E%20Gillespie%20simulation.ipynb

