## Supplementary material for "Synthetic dosage-compensating miRNA circuits allow precision gene therapy for Rett syndrome": CAG-GFP_cell_segmentation

mouse20_CAGGFP

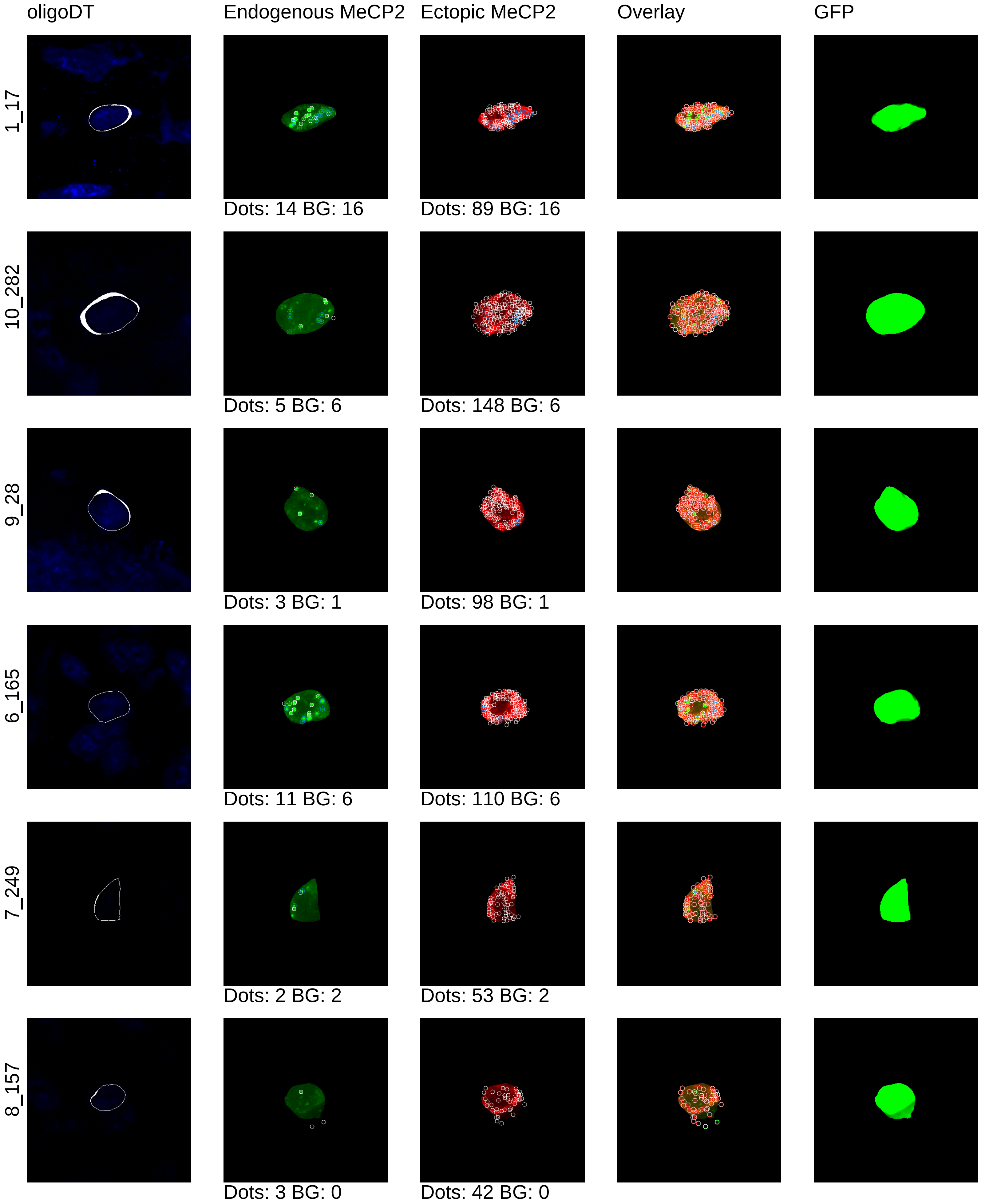

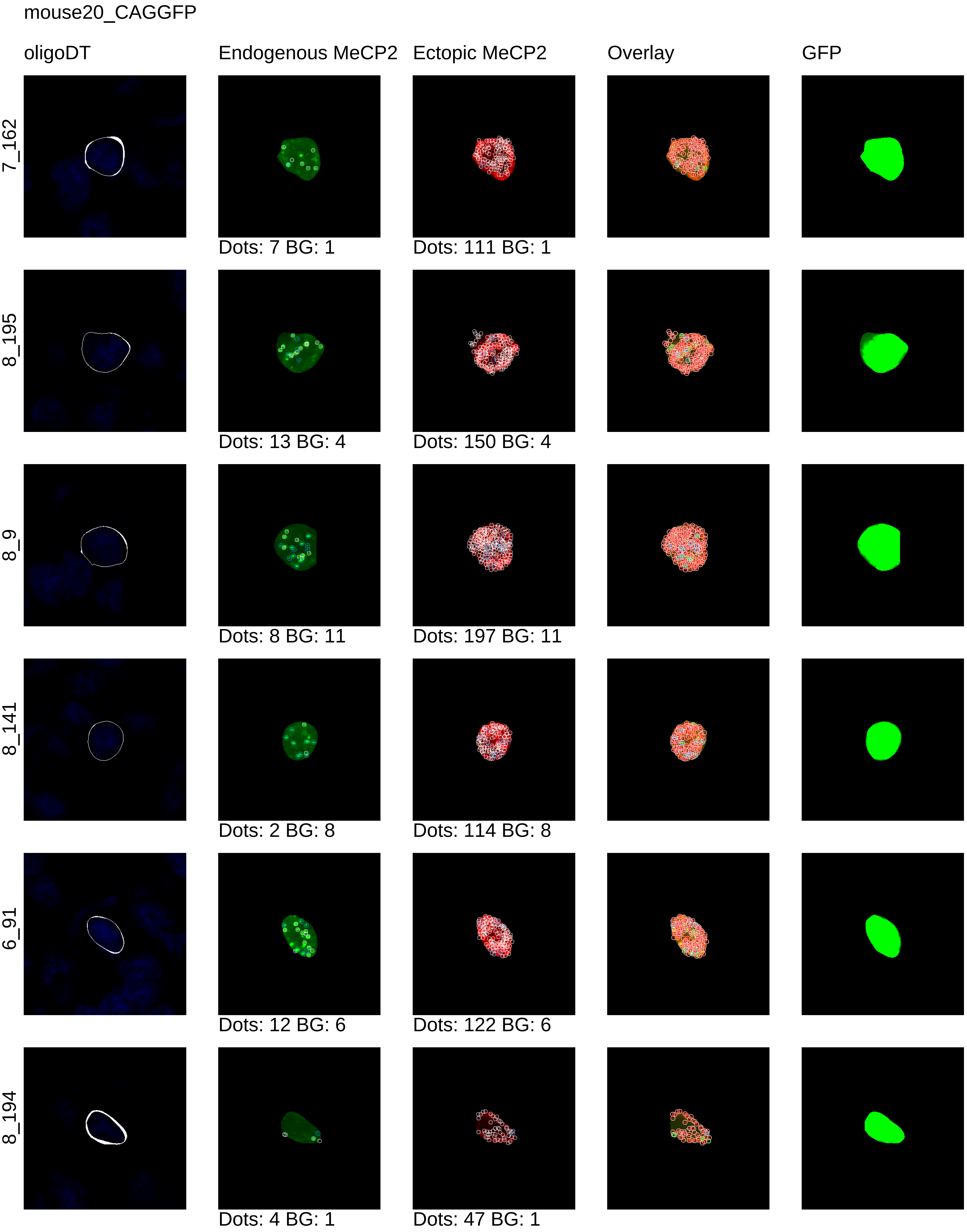
mouse20_CAGGFP
oligoDT
Endogenous MeCP2
Ectopic MeCP2
Overlay
GFP
7_162
Dots: 7 BG: 1
Dots: 111 BG: 1
8_195
Dots: 13 BG: 4
Dots: 150 BG: 4
8_9
Dots: 8 BG: 11
Dots: 197 BG: 11
8_141
Dots: 2 BG: 8
Dots: 114 BG: 8
6_91
Dots: 12 BG: 6
Dots: 122 BG: 6
8_194
Dots: 4 BG: 1
Dots: 47 BG: 1

mouse20_CAGGFP

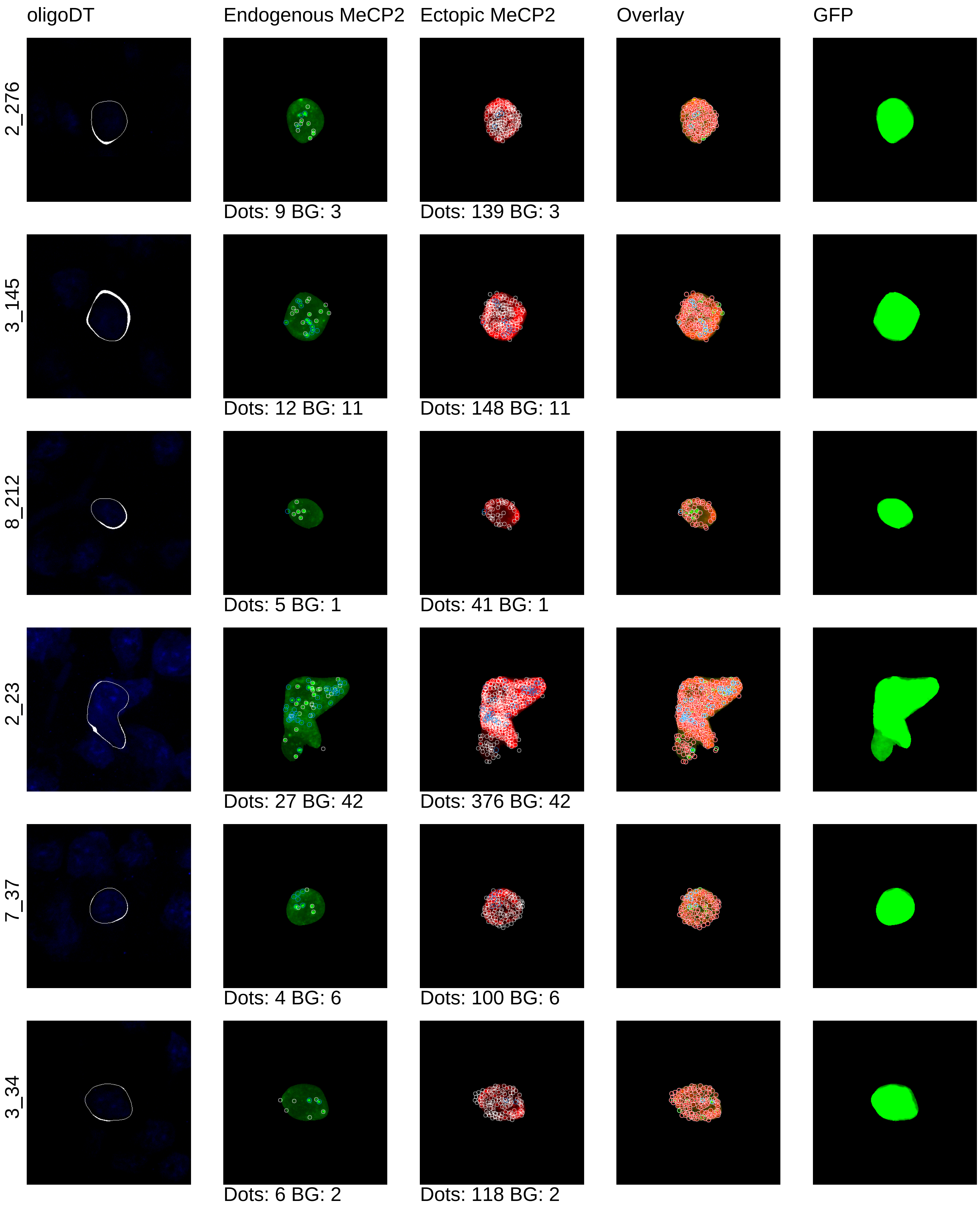

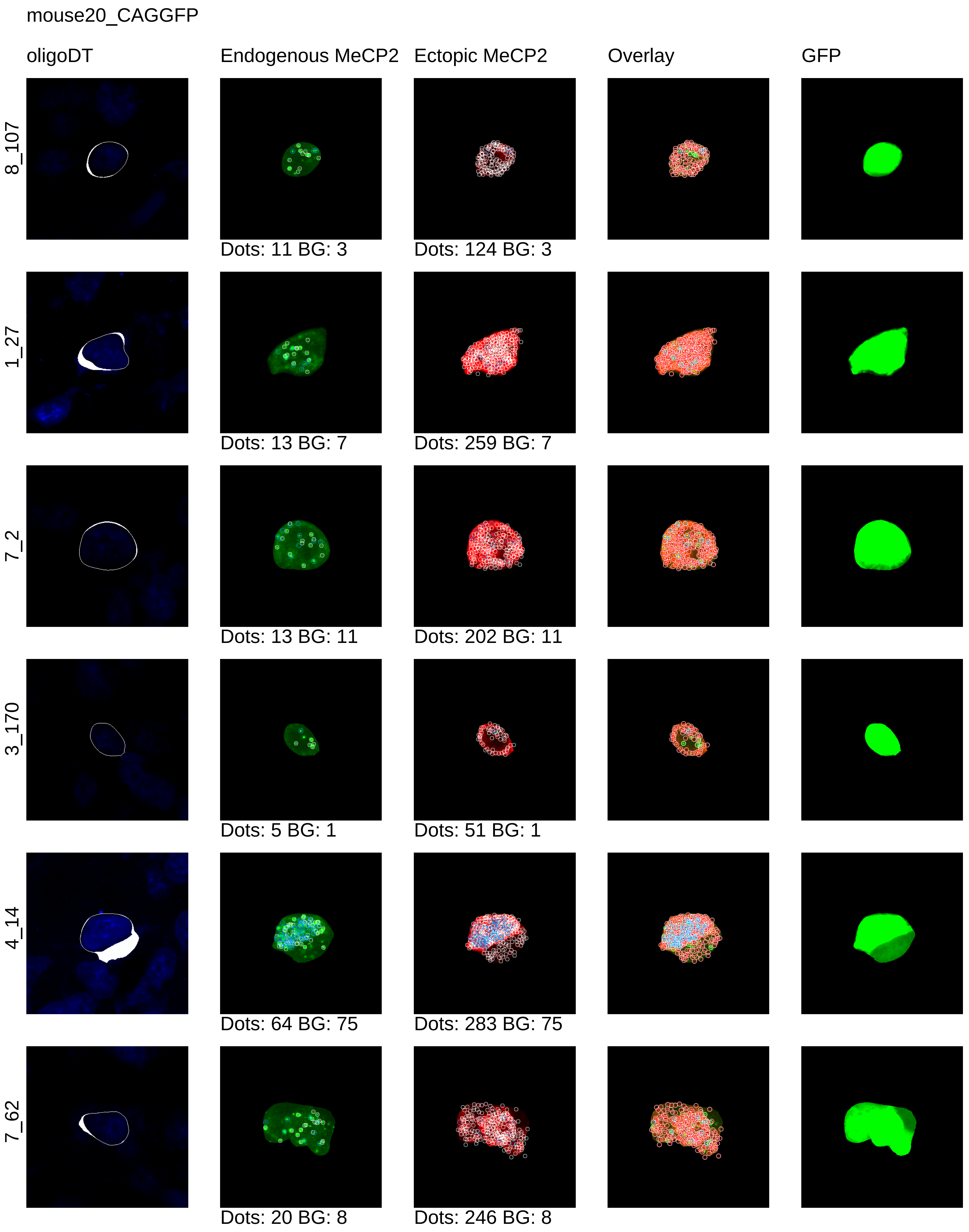
mouse20_CAGGFP
oligoDT
Endogenous MeCP2
Ectopic MeCP2
Overlay
GFP
8_107
Dots: 11 BG: 3
Dots: 124 BG: 3
1_27
Dots: 13 BG: 7
Dots: 259 BG: 7
7_2
Dots: 13 BG: 11
Dots: 202 BG: 11
3_170
Dots: 5 BG: 1
Dots: 51 BG: 1
4_14
Dots: 64 BG: 75
Dots: 283 BG: 75
7_62
Dots: 20 BG: 8
Dots: 246 BG: 8

mouse20_CAGGFP

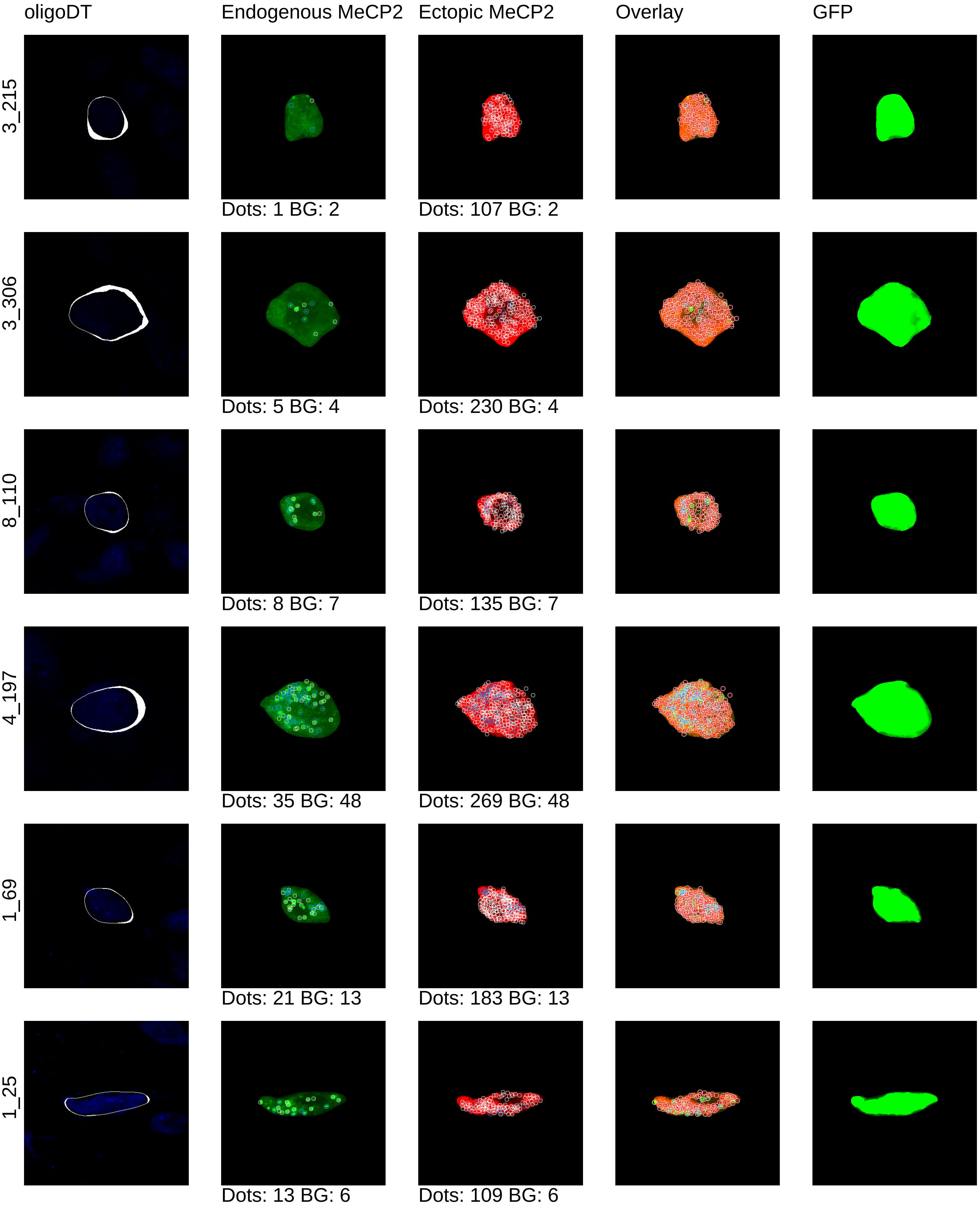

mouse20_CAGGFP

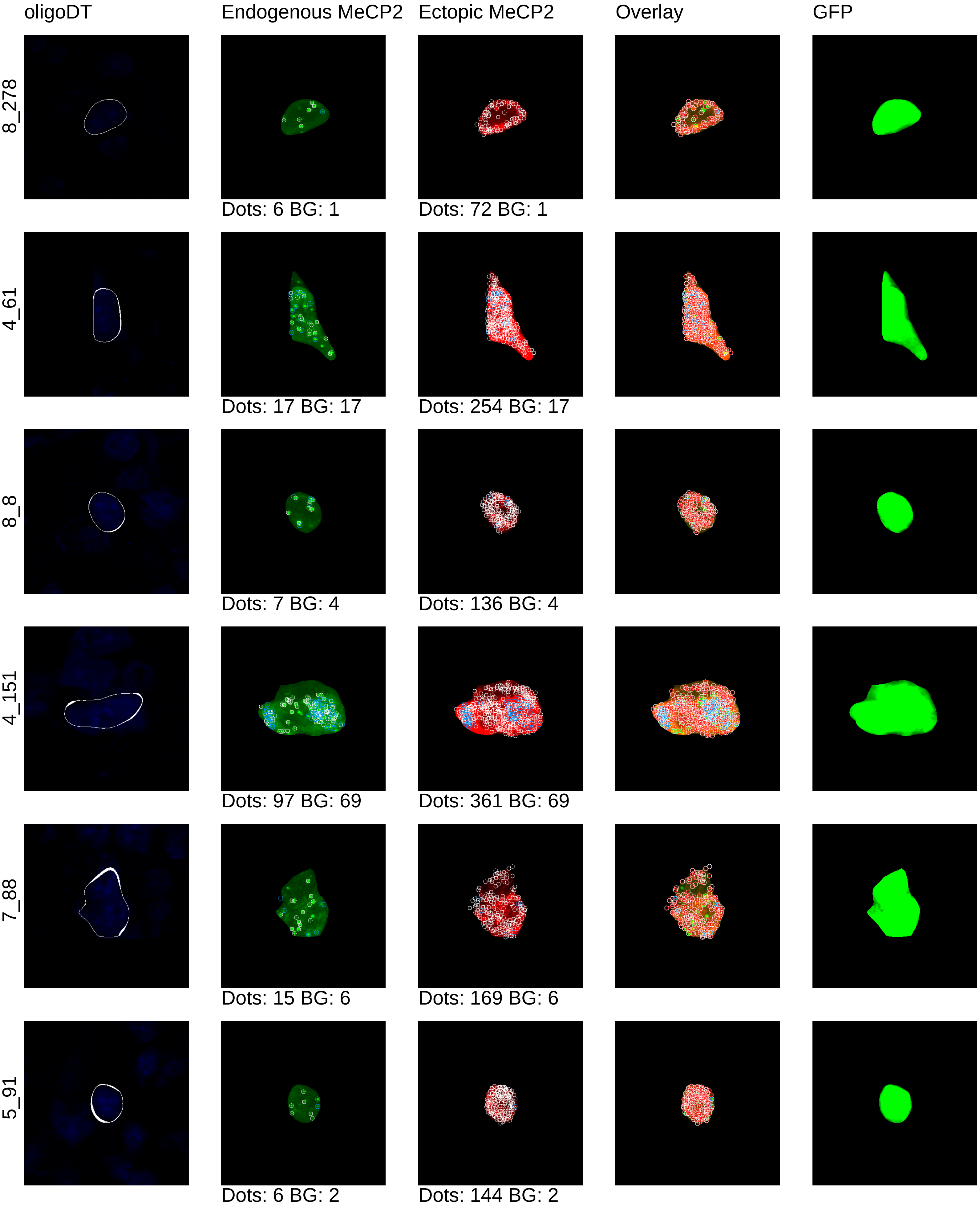

mouse20_CAGGFP

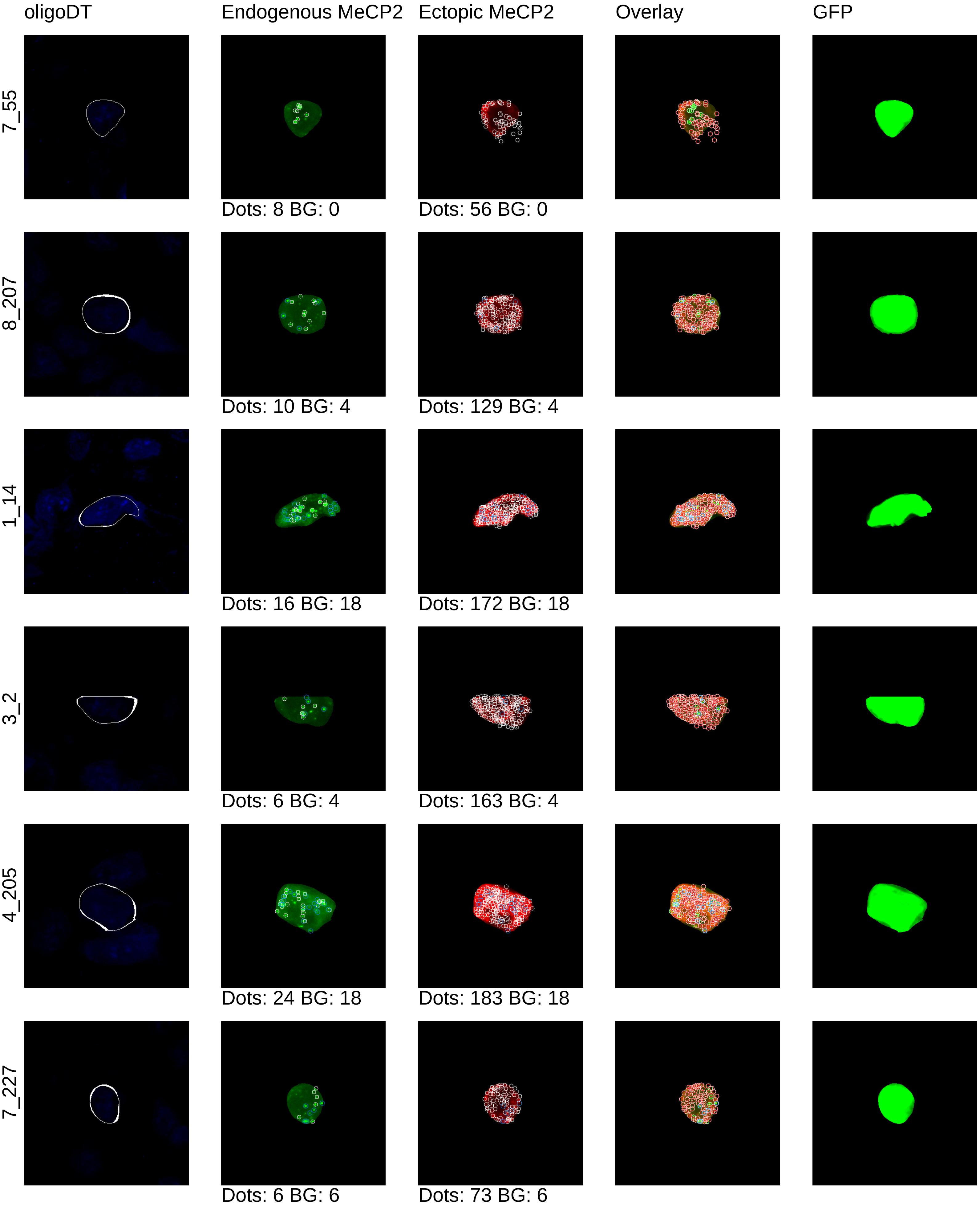

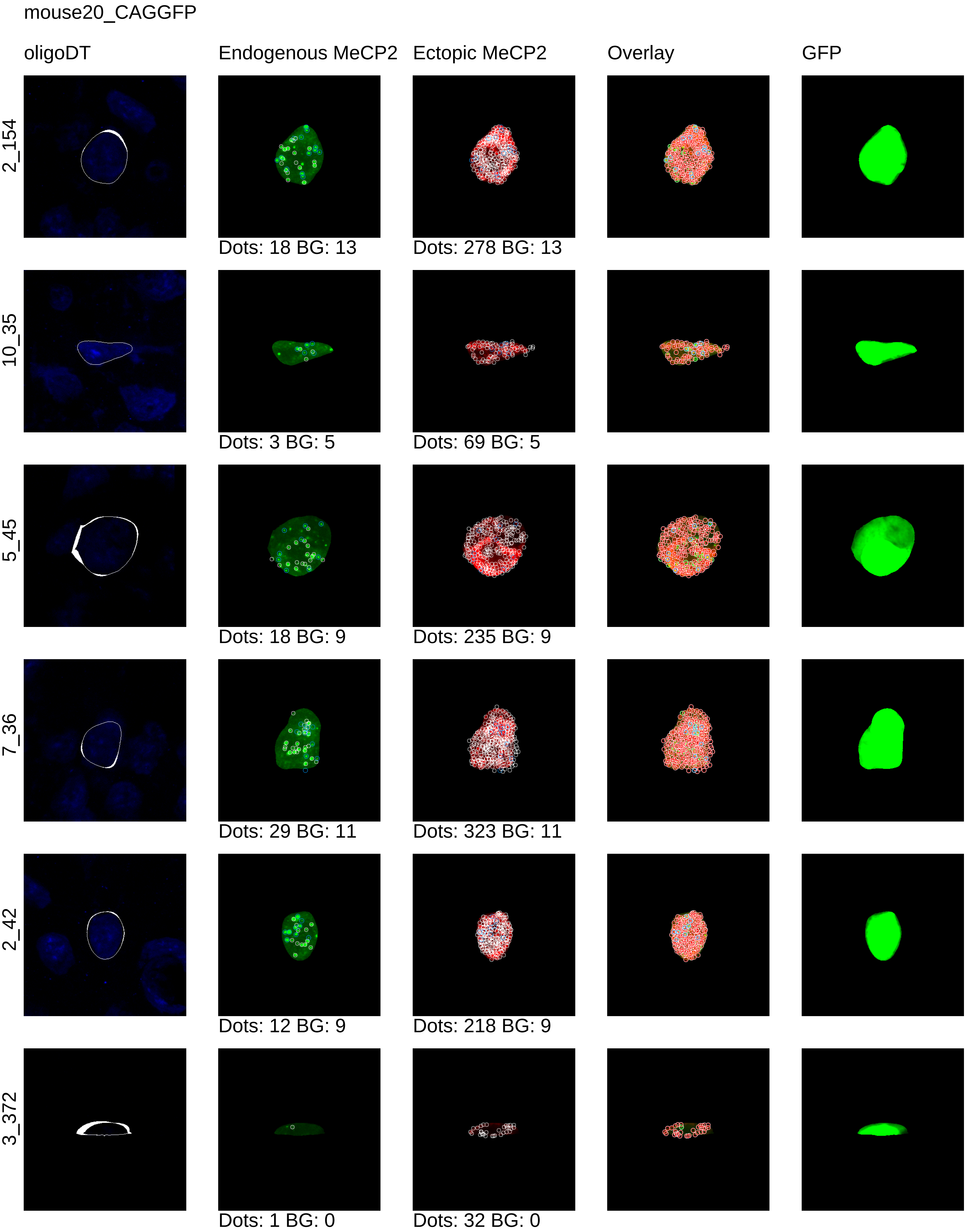
mouse20_CAGGFP
oligoDT
Endogenous MeCP2
Ectopic MeCP2
Overlay
GFP
2_154
Dots: 18 BG: 13
Dots: 278 BG: 13
10_35
Dots: 3 BG: 5
Dots: 69 BG: 5
5_45
Dots: 18 BG: 9
Dots: 235 BG: 9
7_36
Dots: 29 BG: 11
Dots: 323 BG: 11
2_42
Dots: 12 BG: 9
Dots: 218 BG: 9
3_372
Dots: 1 BG: 0
Dots: 32 BG: 0

mouse20_CAGGFP

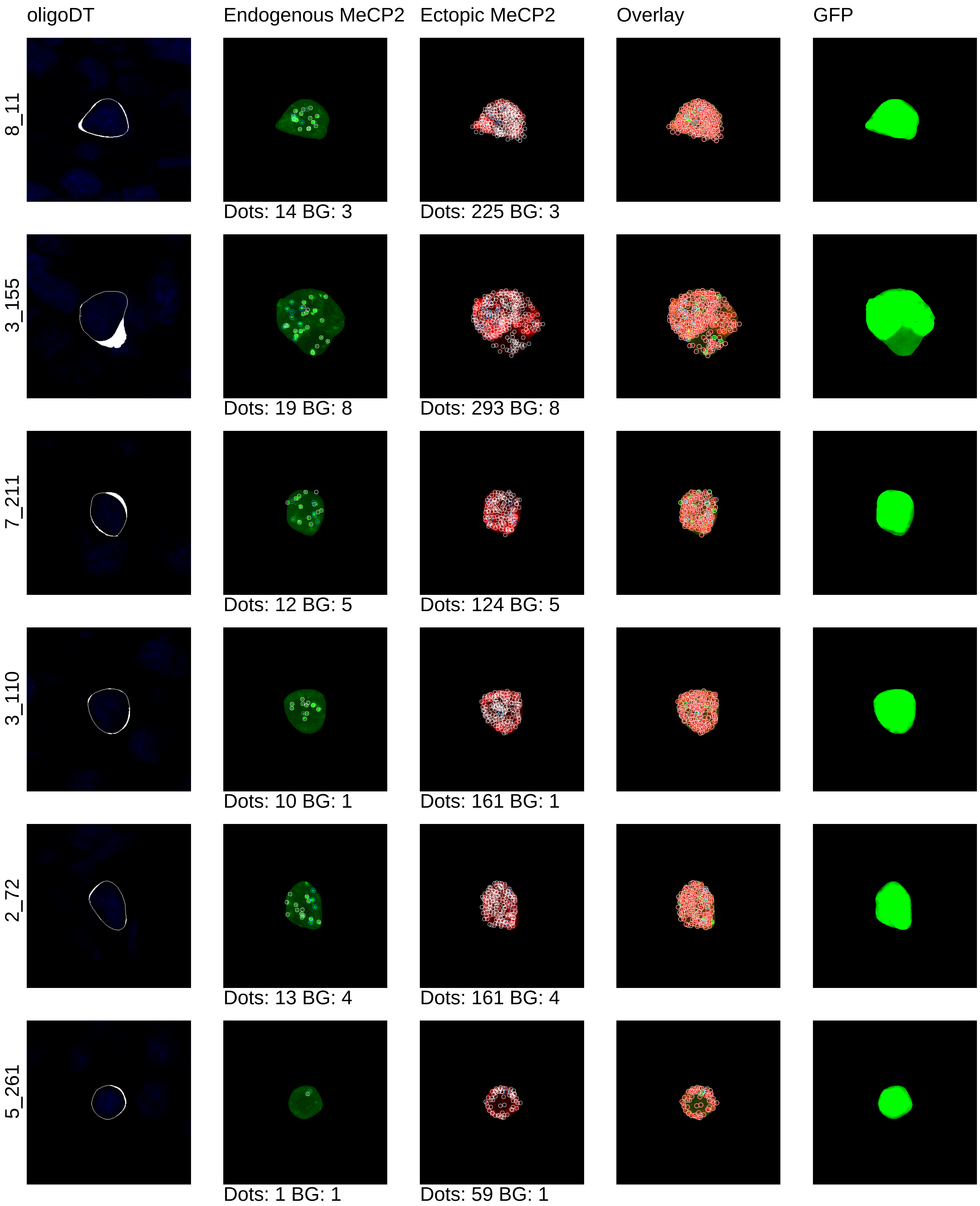

mouse20_CAGGFP

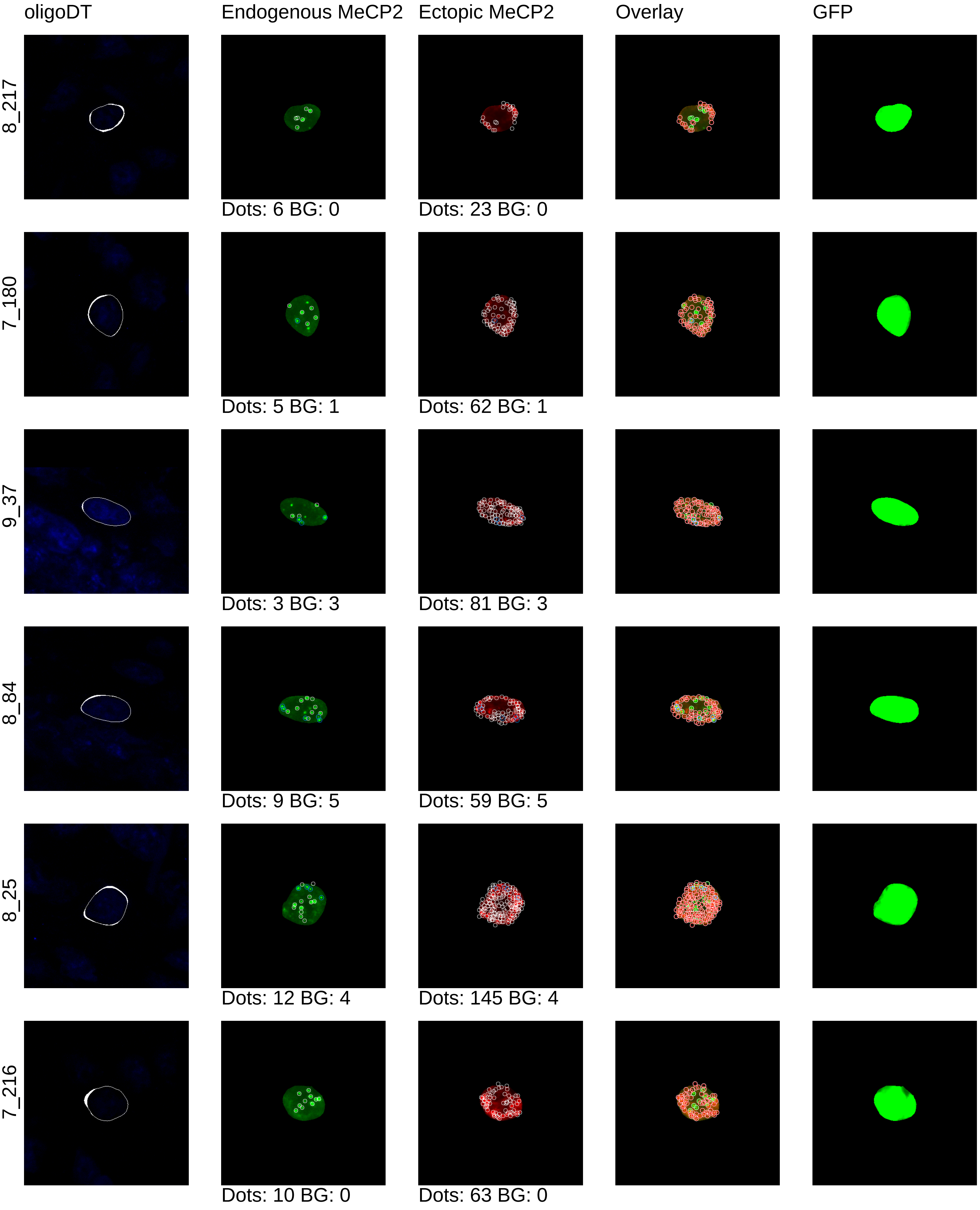

mouse20_CAGGFP

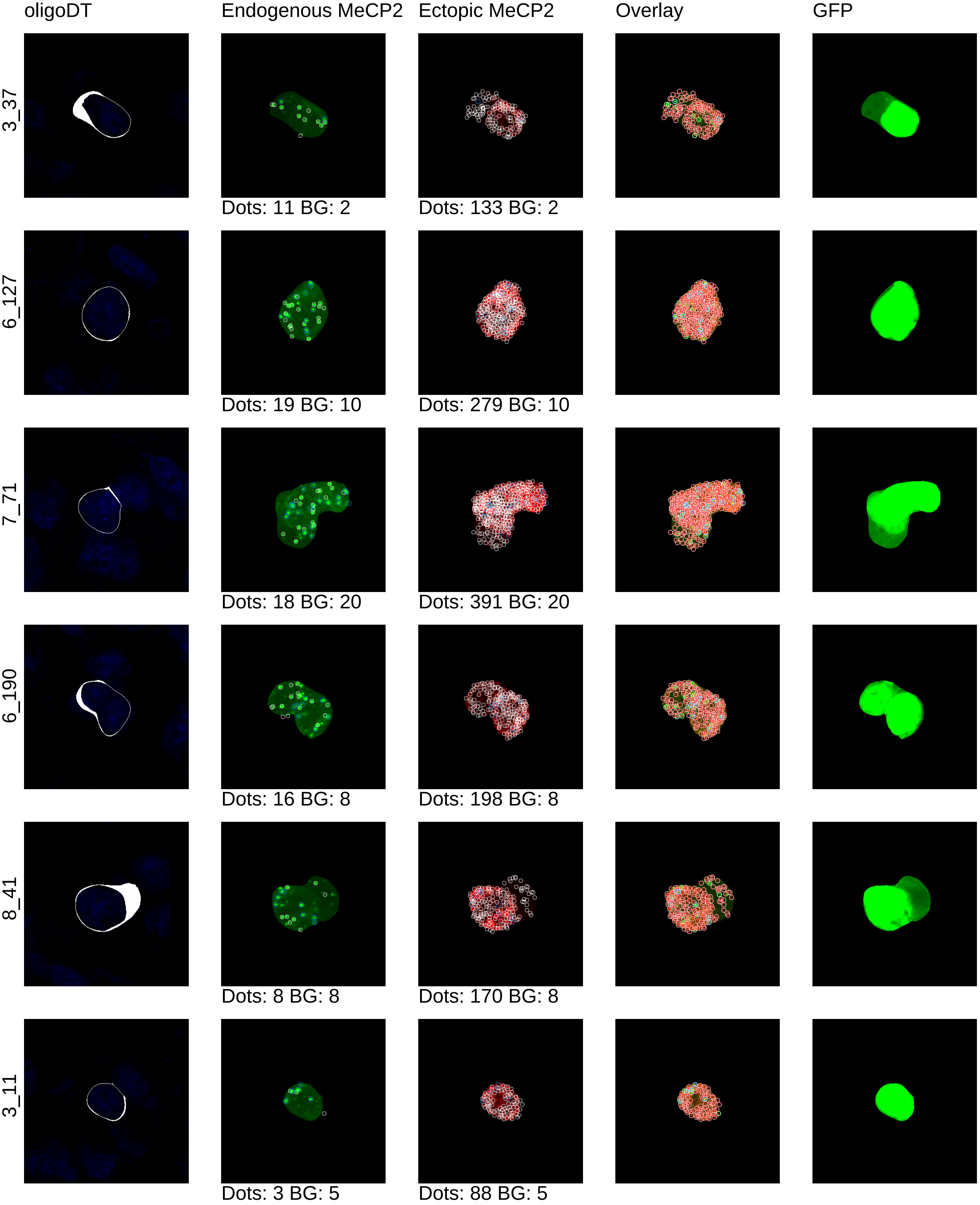

mouse20_CAGGFP

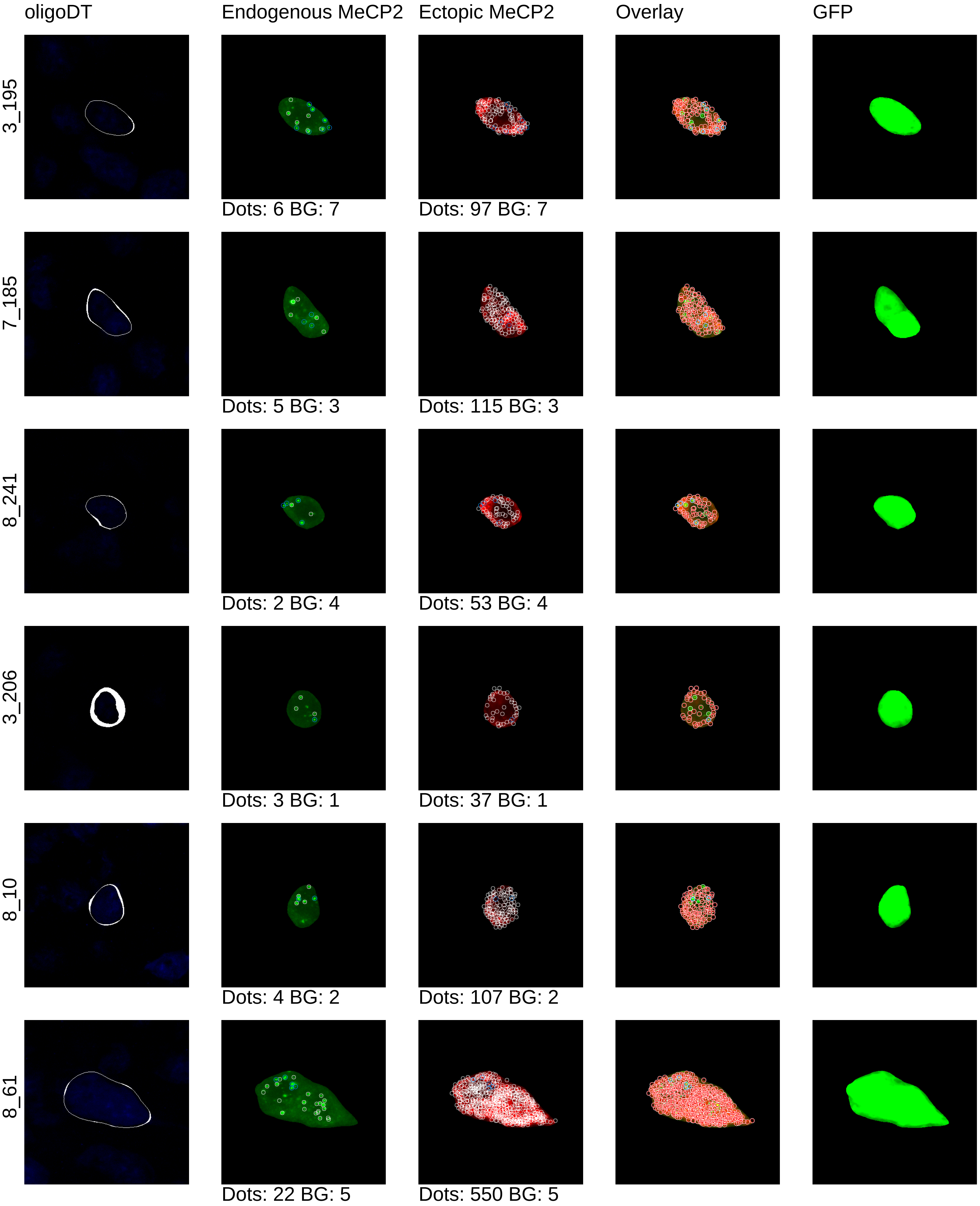

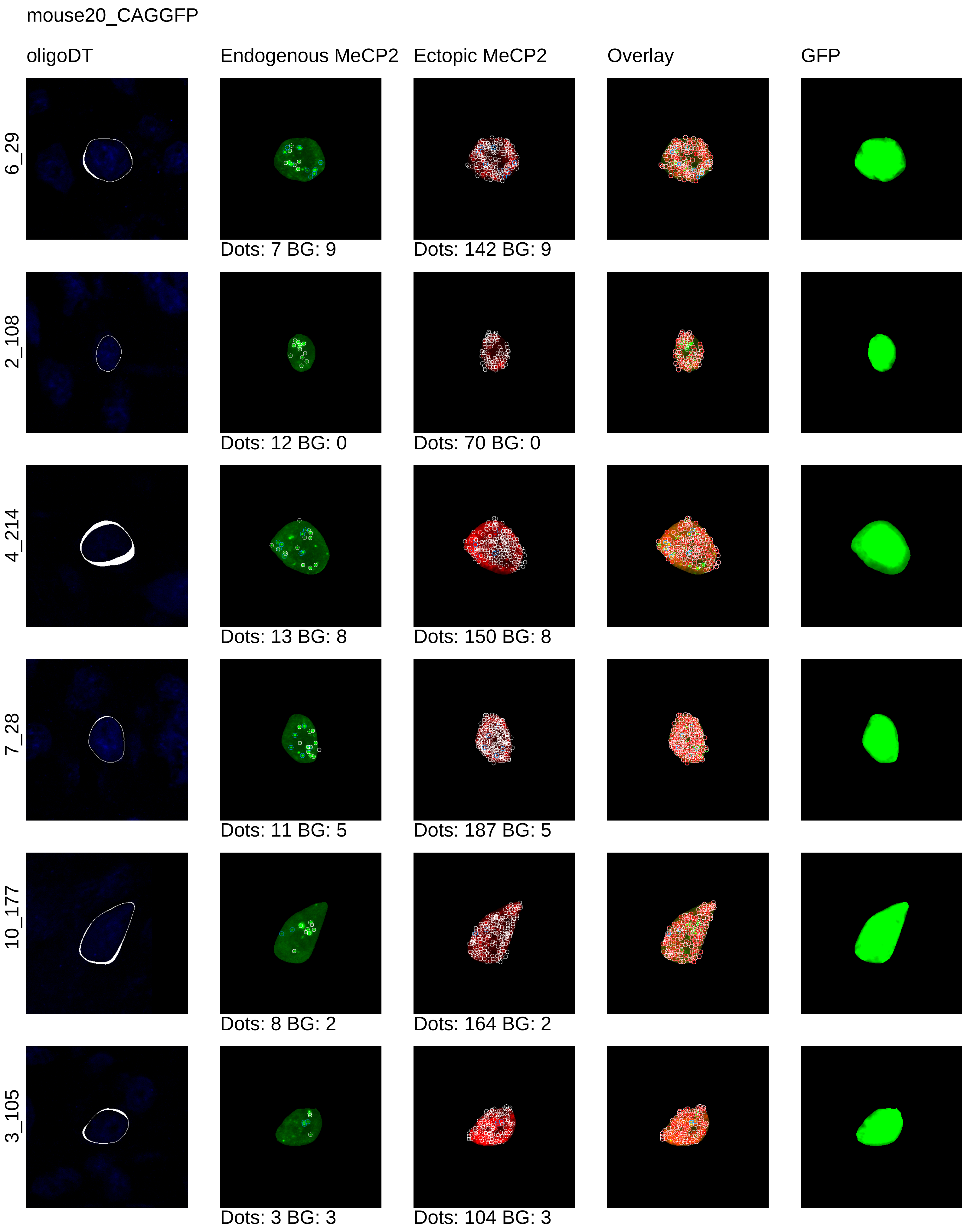

mouse20_CAGGFP
oligoDT
Endogenous MeCP2
Ectopic MeCP2
Overlay
GFP
6_29
Dots: 7 BG: 9
Dots: 142 BG: 9
2_108
Dots: 12 BG: 0
Dots: 70 BG: 0
4_214
Dots: 13 BG: 8
Dots: 150 BG: 8
7_28
Dots: 11 BG: 5
Dots: 187 BG: 5
10_177
Dots: 8 BG: 2
Dots: 164 BG: 2
3_105
Dots: 3 BG: 3
Dots: 104 BG: 3

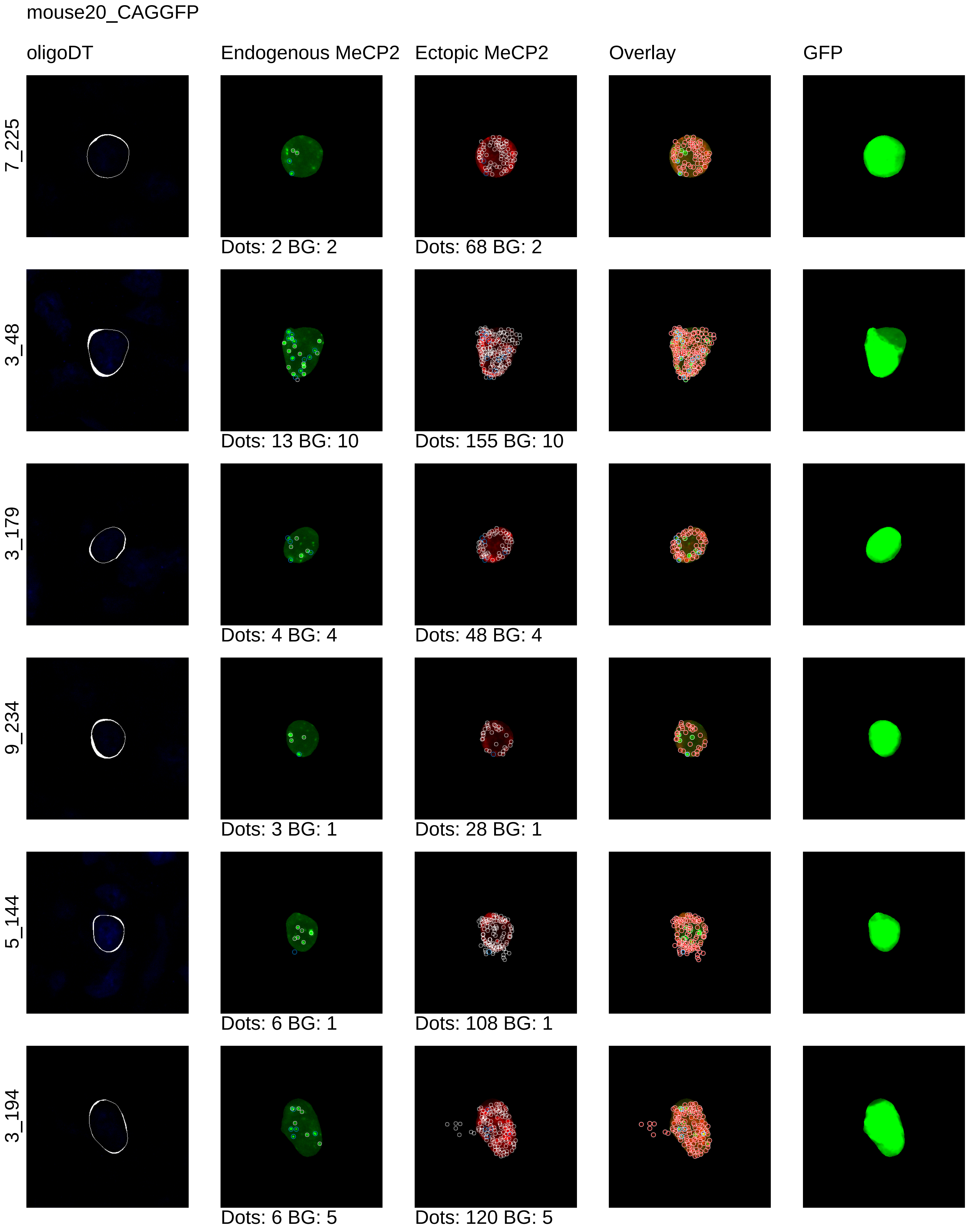
mouse20_CAGGFP
oligoDT
Endogenous MeCP2
Ectopic MeCP2
Overlay
GFP
7_225
Dots: 2 BG: 2
Dots: 68 BG: 2
3_48
Dots: 13 BG: 10
Dots: 155 BG: 10
3_179
Dots: 4 BG: 4
Dots: 48 BG: 4
9_234
Dots: 3 BG: 1
Dots: 28 BG: 1
5_144
Dots: 6 BG: 1
Dots: 108 BG: 1
3_194
Dots: 6 BG: 5
Dots: 120 BG: 5

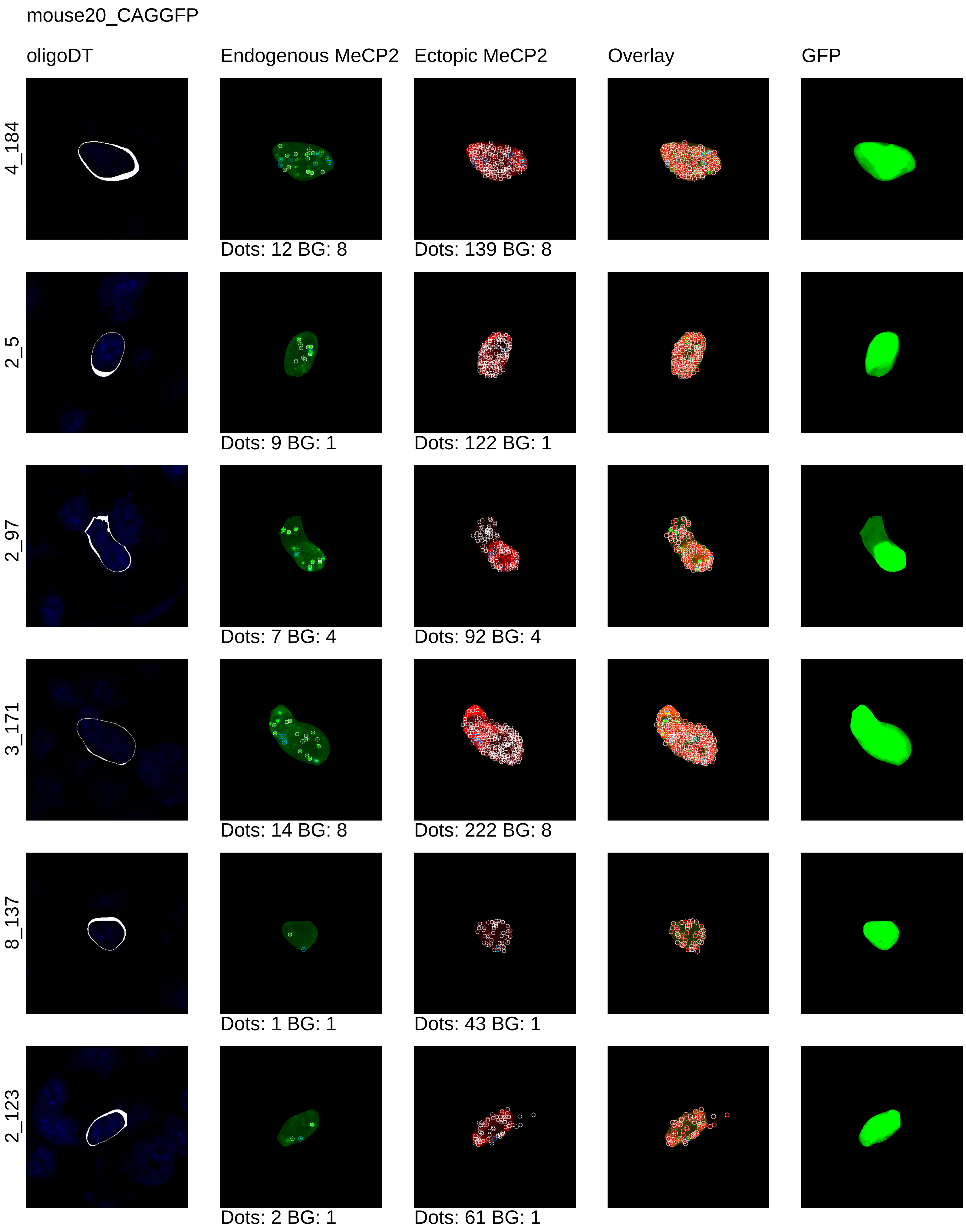
mouse20_CAGGFP
oligoDT
Endogenous MeCP2
Ectopic MeCP2
Overlay
GFP
4_184
Dots: 12 BG: 8
Dots: 139 BG: 8
2_5
Dots: 9 BG: 1
Dots: 122 BG: 1
2_97
Dots: 7 BG: 4
Dots: 92 BG: 4
3_171
Dots: 14 BG: 8
Dots: 222 BG: 8
8_137
Dots: 1 BG: 1
Dots: 43 BG: 1
2_123
Dots: 2 BG: 1
Dots: 61 BG: 1

mouse20_CAGGFP

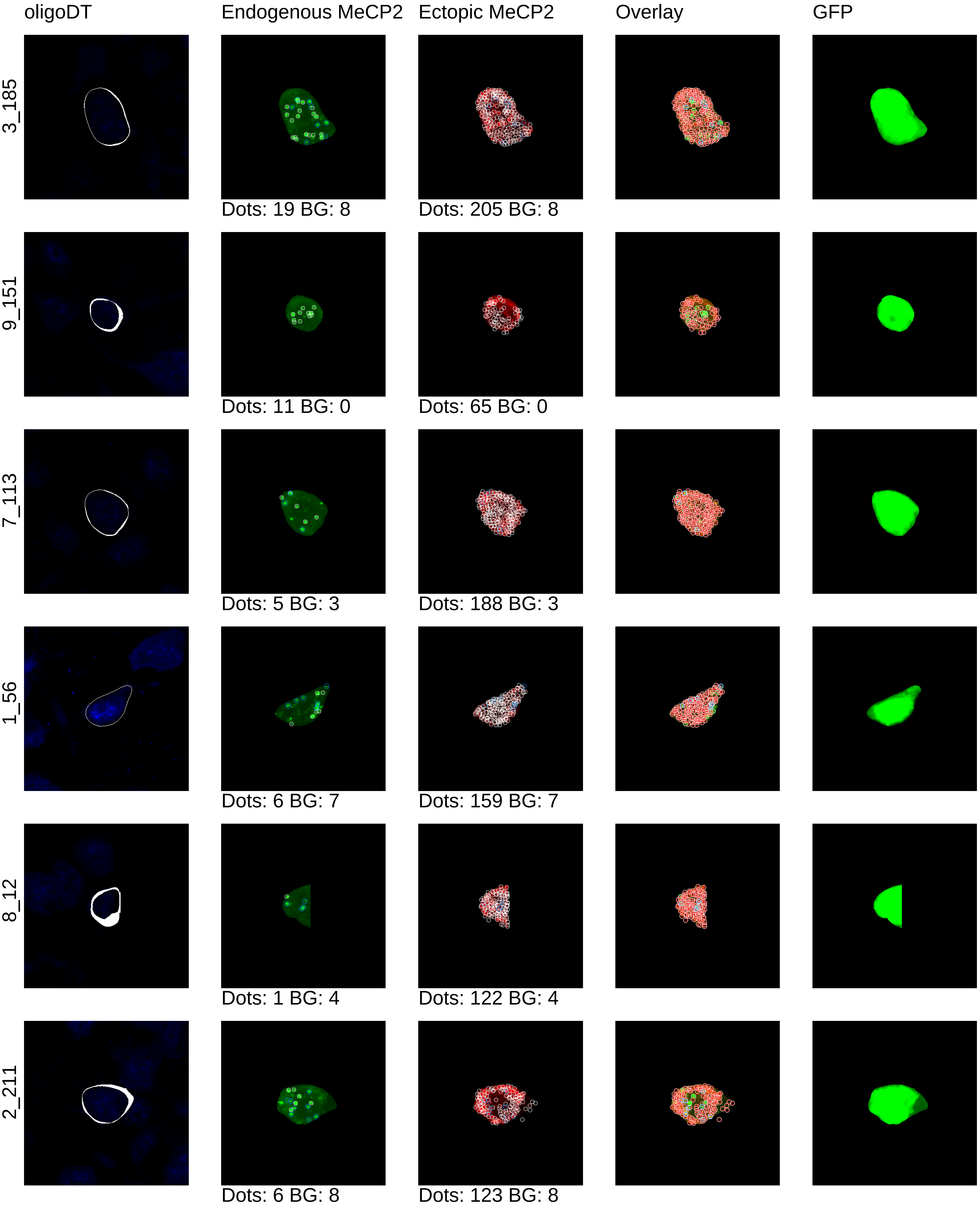

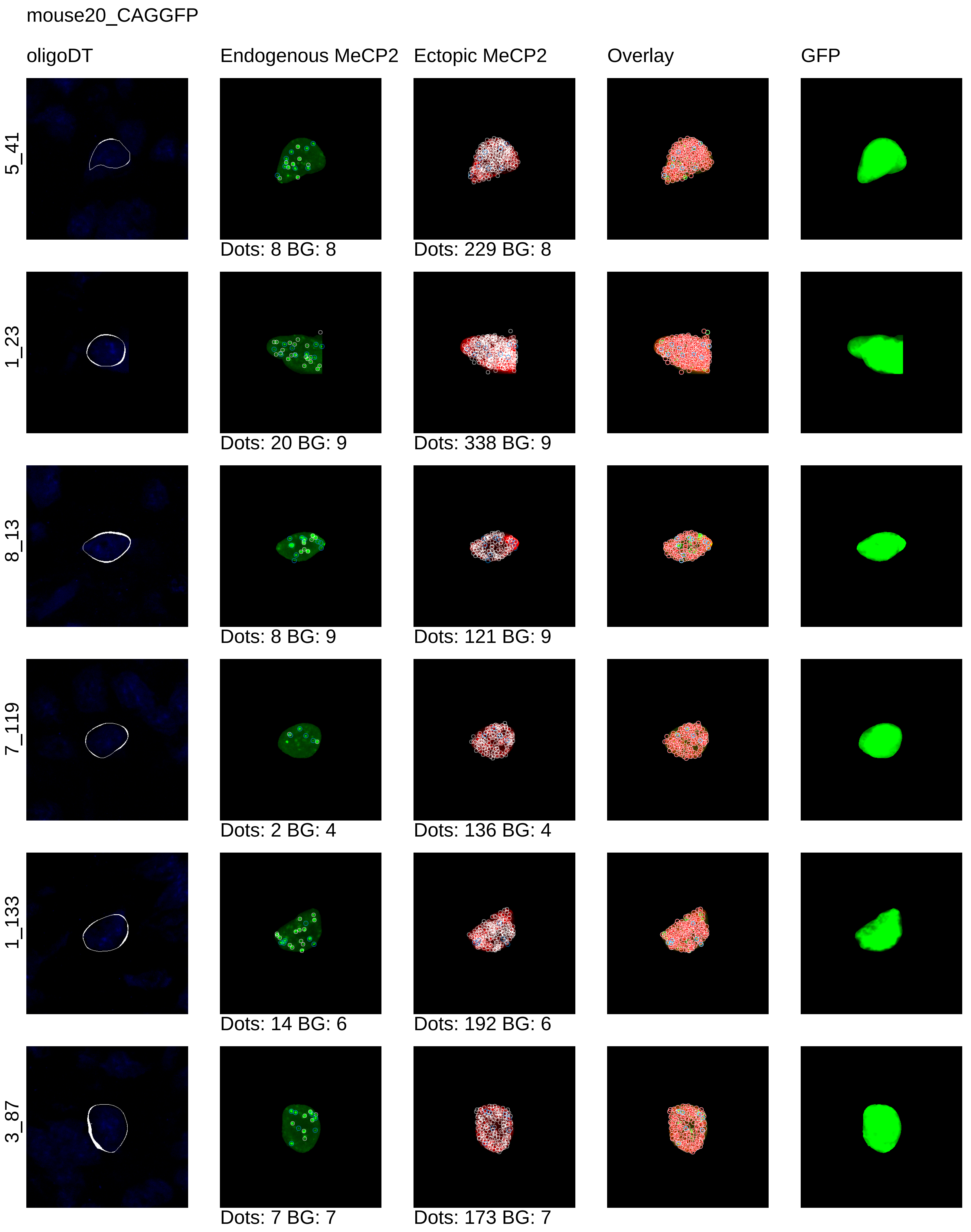
mouse20_CAGGFP
oligoDT
Endogenous MeCP2
Ectopic MeCP2
Overlay
GFP
5_41
Dots: 8 BG: 8
Dots: 229 BG: 8
1_23
Dots: 20 BG: 9
Dots: 338 BG: 9
8_13
Dots: 8 BG: 9
Dots: 121 BG: 9
7_119
Dots: 2 BG: 4
Dots: 136 BG: 4
1_133
Dots: 14 BG: 6
Dots: 192 BG: 6
3_87
Dots: 7 BG: 7
Dots: 173 BG: 7

mouse20_CAGGFP

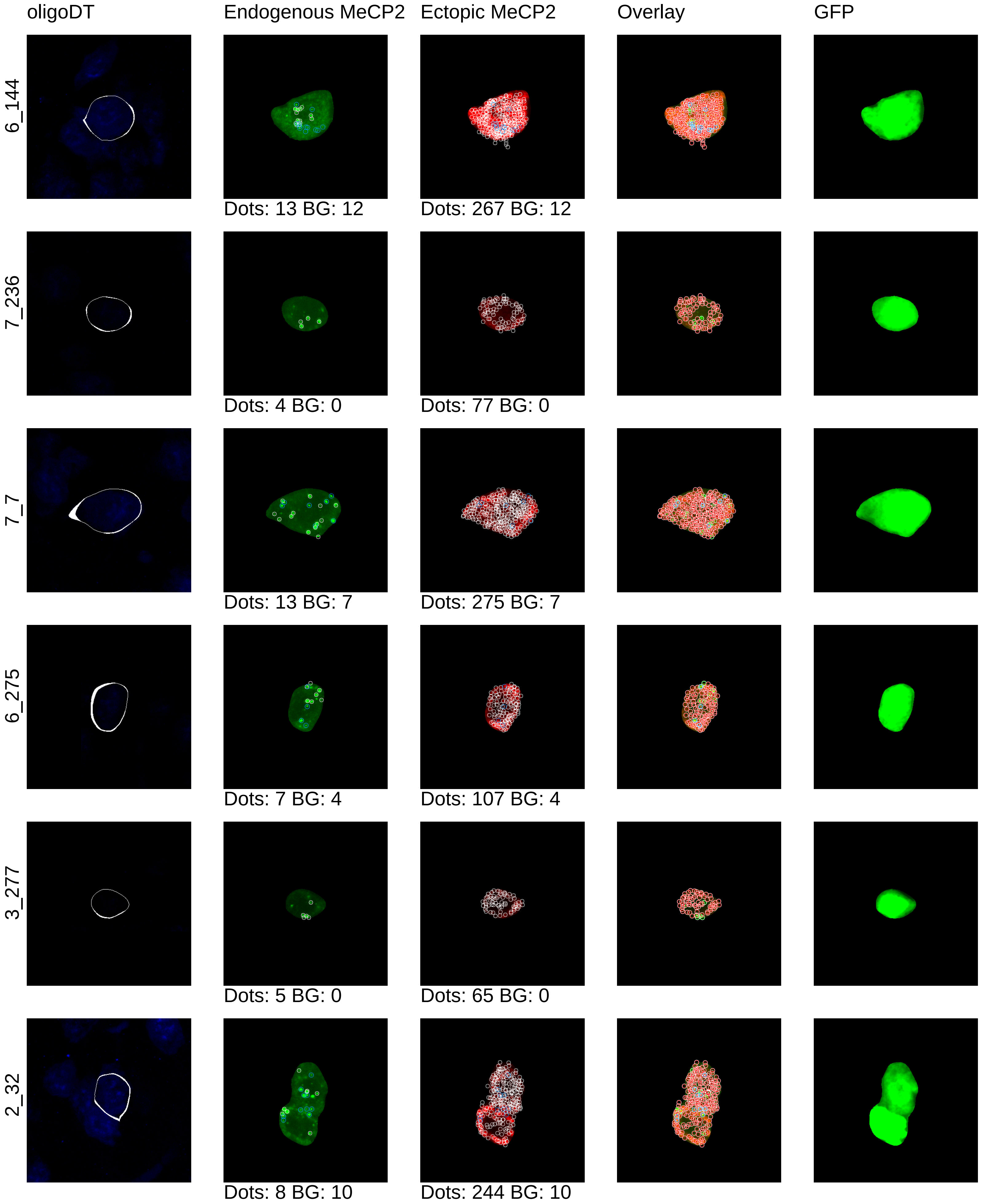

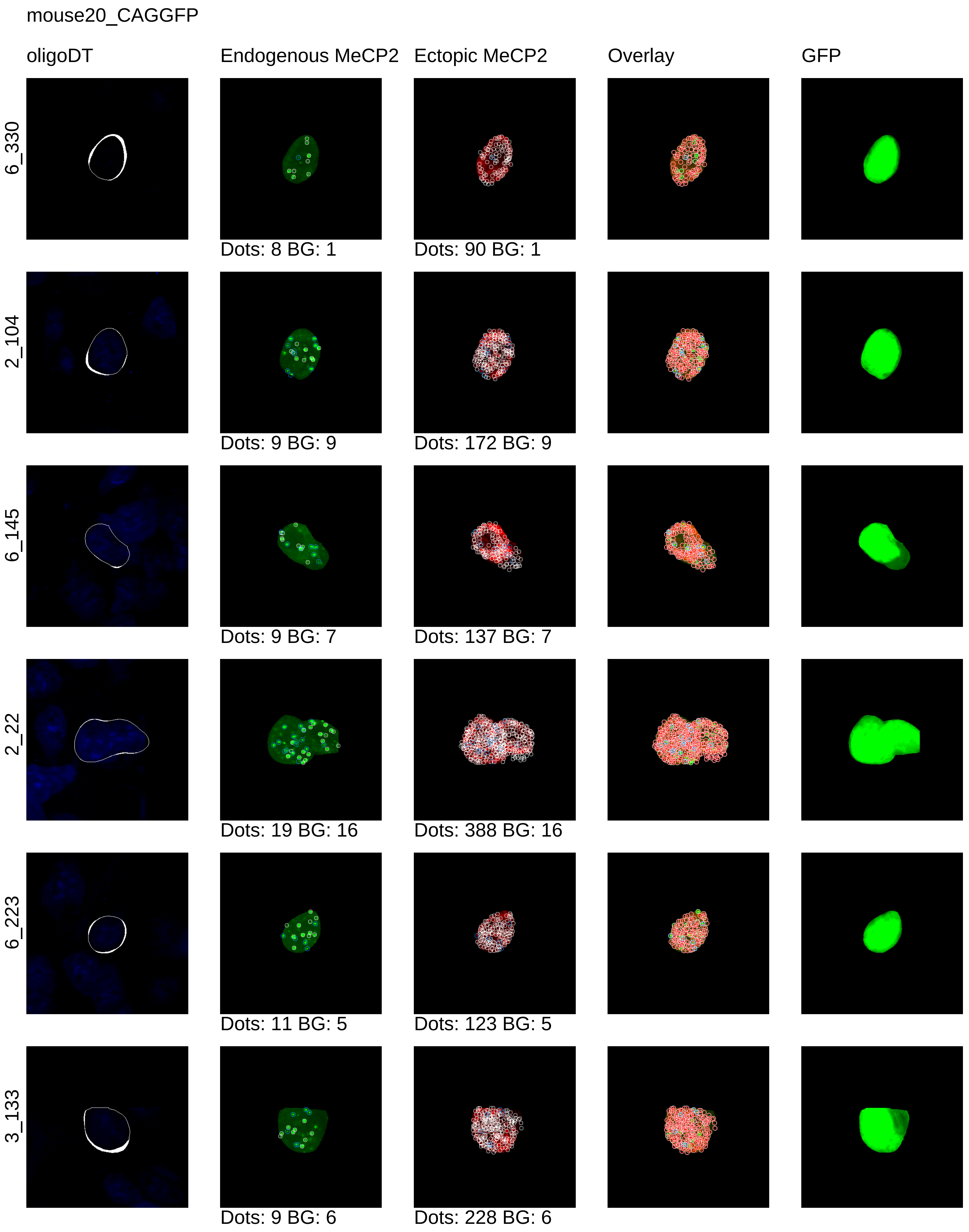
mouse20_CAGGFP
oligoDT
Endogenous MeCP2
Ectopic MeCP2
Overlay
GFP
6_330
Dots: 8 BG: 1
Dots: 90 BG: 1
2_104
Dots: 9 BG: 9
Dots: 172 BG: 9
6_145
Dots: 9 BG: 7
Dots: 137 BG: 7
2_22
Dots: 19 BG: 16
Dots: 388 BG: 16
6_223
Dots: 11 BG: 5
Dots: 123 BG: 5
3_133
Dots: 9 BG: 6
Dots: 228 BG: 6

mouse20_CAGGFP

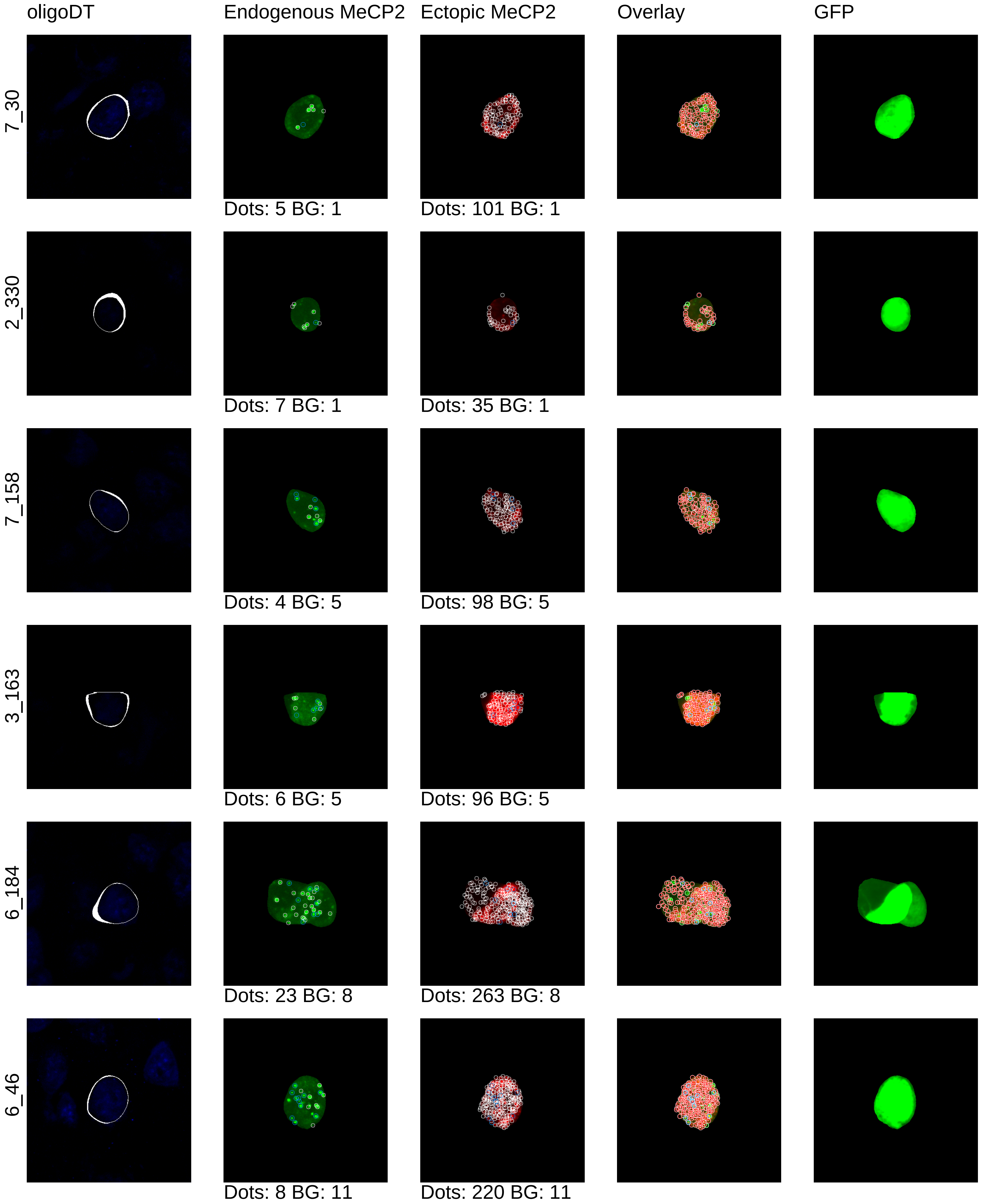

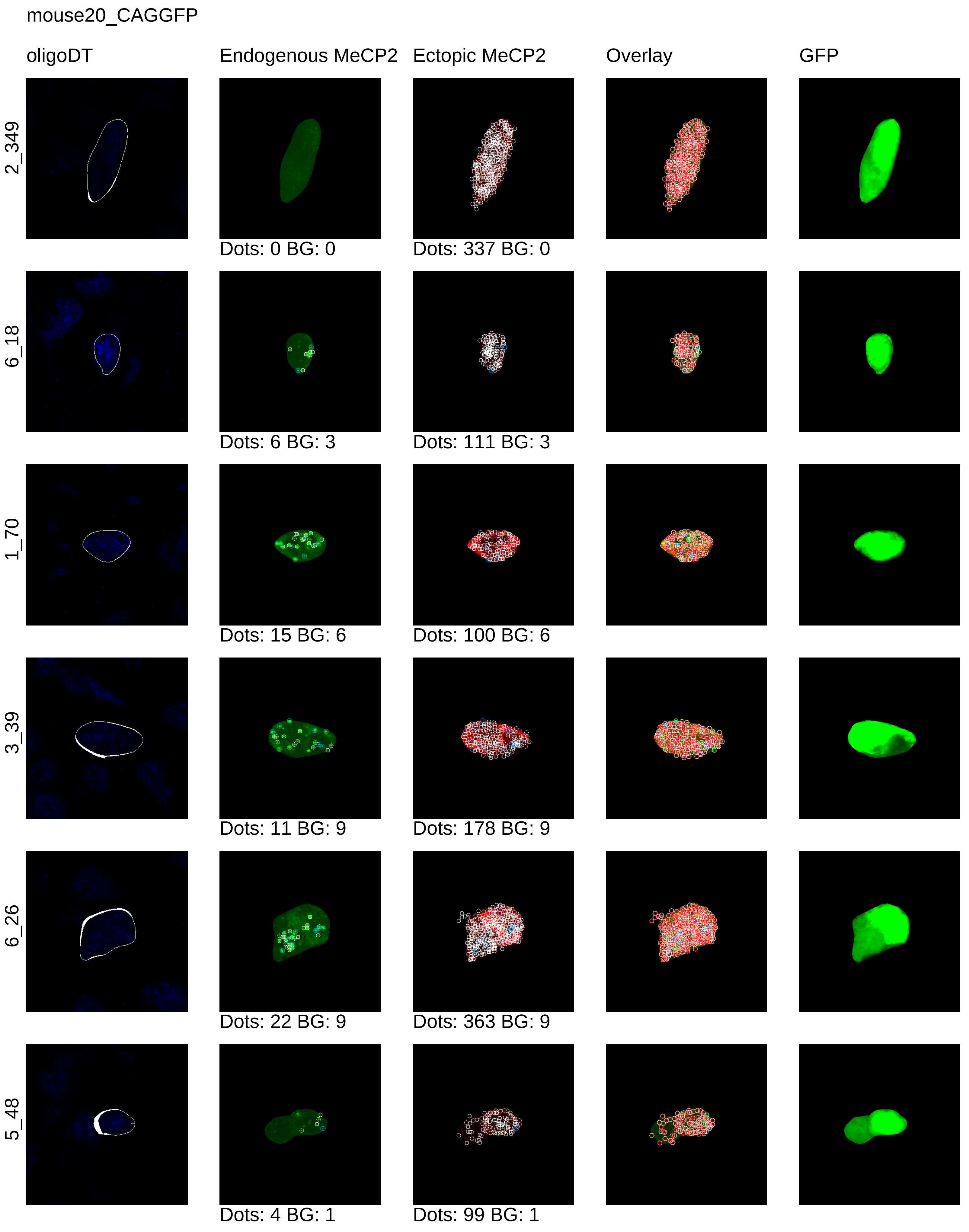
mouse20_CAGGFP
oligoDT
Endogenous MeCP2
Ectopic MeCP2
Overlay
GFP
2_349
Dots: 0 BG: 0
Dots: 337 BG: 0
6_18
Dots: 6 BG: 3
Dots: 111 BG: 3
1_70
Dots: 15 BG: 6
Dots: 100 BG: 6
3_39
Dots: 11 BG: 9
Dots: 178 BG: 9
6_26
Dots: 22 BG: 9
Dots: 363 BG: 9
5_48
Dots: 4 BG: 1
Dots: 99 BG: 1

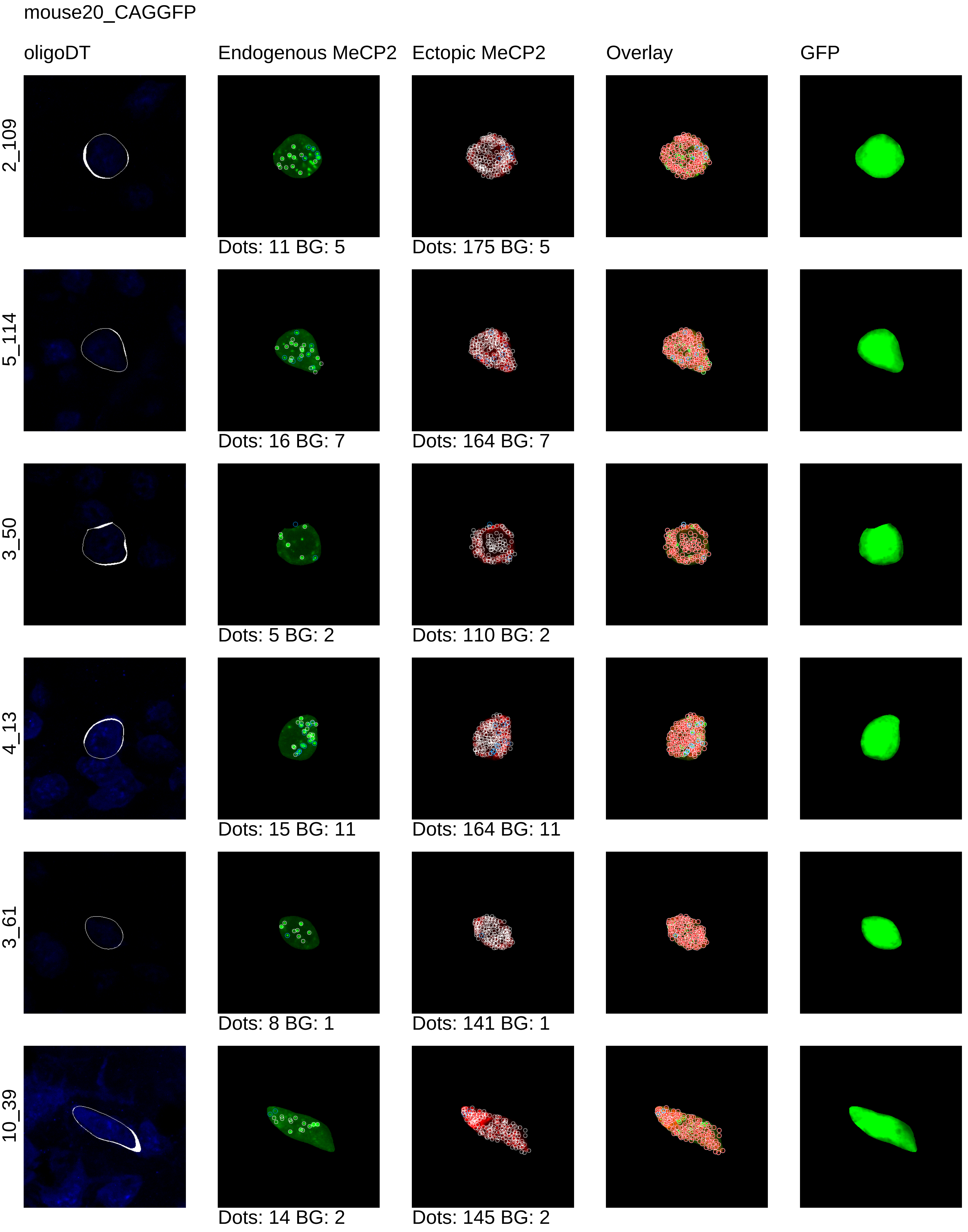
mouse20_CAGGFP
oligoDT
Endogenous MeCP2
Ectopic MeCP2
Overlay
GFP
2_109
Dots: 11 BG: 5
Dots: 175 BG: 5
5_114
Dots: 16 BG: 7
Dots: 164 BG: 7
3_50
Dots: 5 BG: 2
Dots: 110 BG: 2
4_13
Dots: 15 BG: 11
Dots: 164 BG: 11
3_61
Dots: 8 BG: 1
Dots: 141 BG: 1
10_39
Dots: 14 BG: 2
Dots: 145 BG: 2

mouse20_CAGGFP

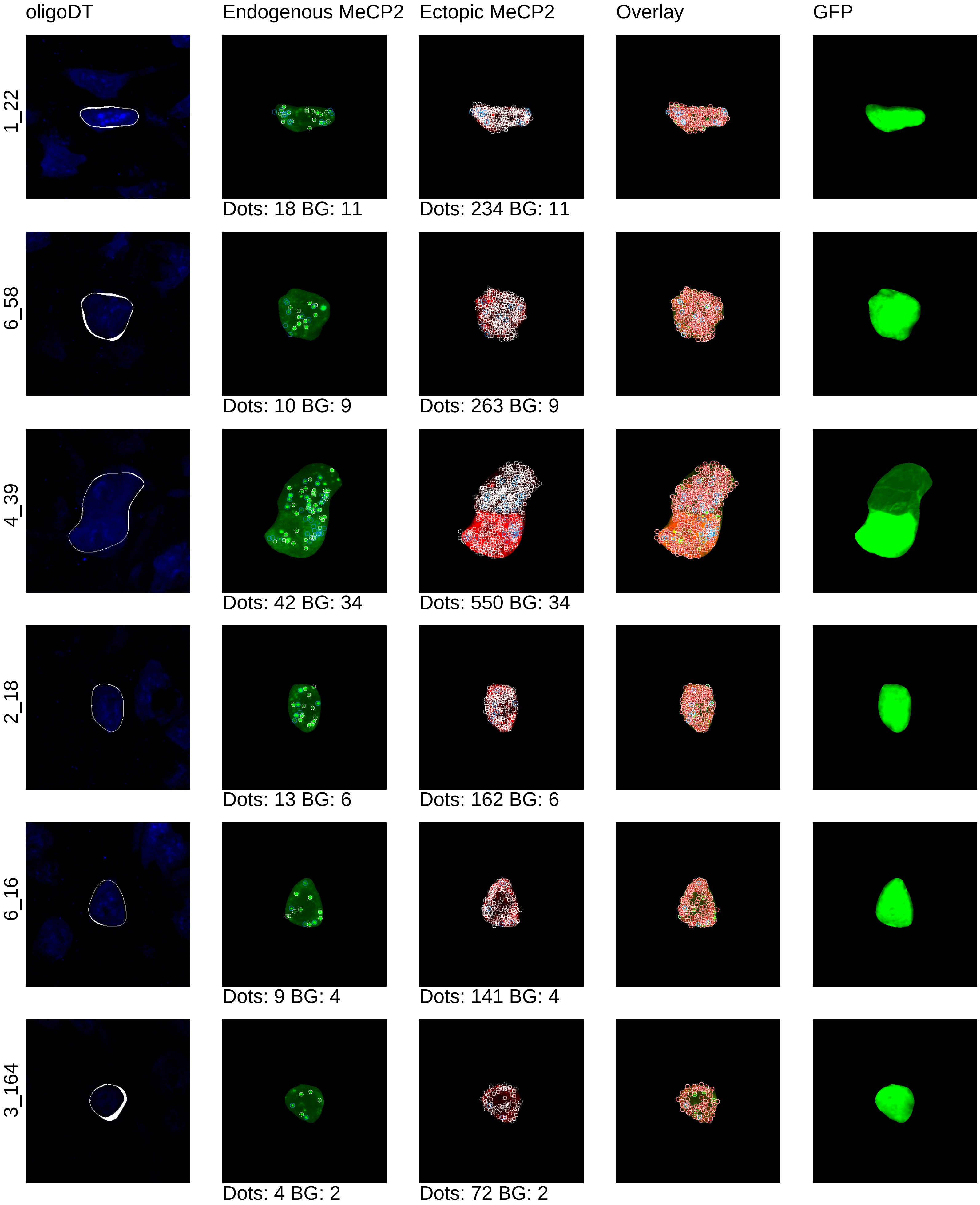

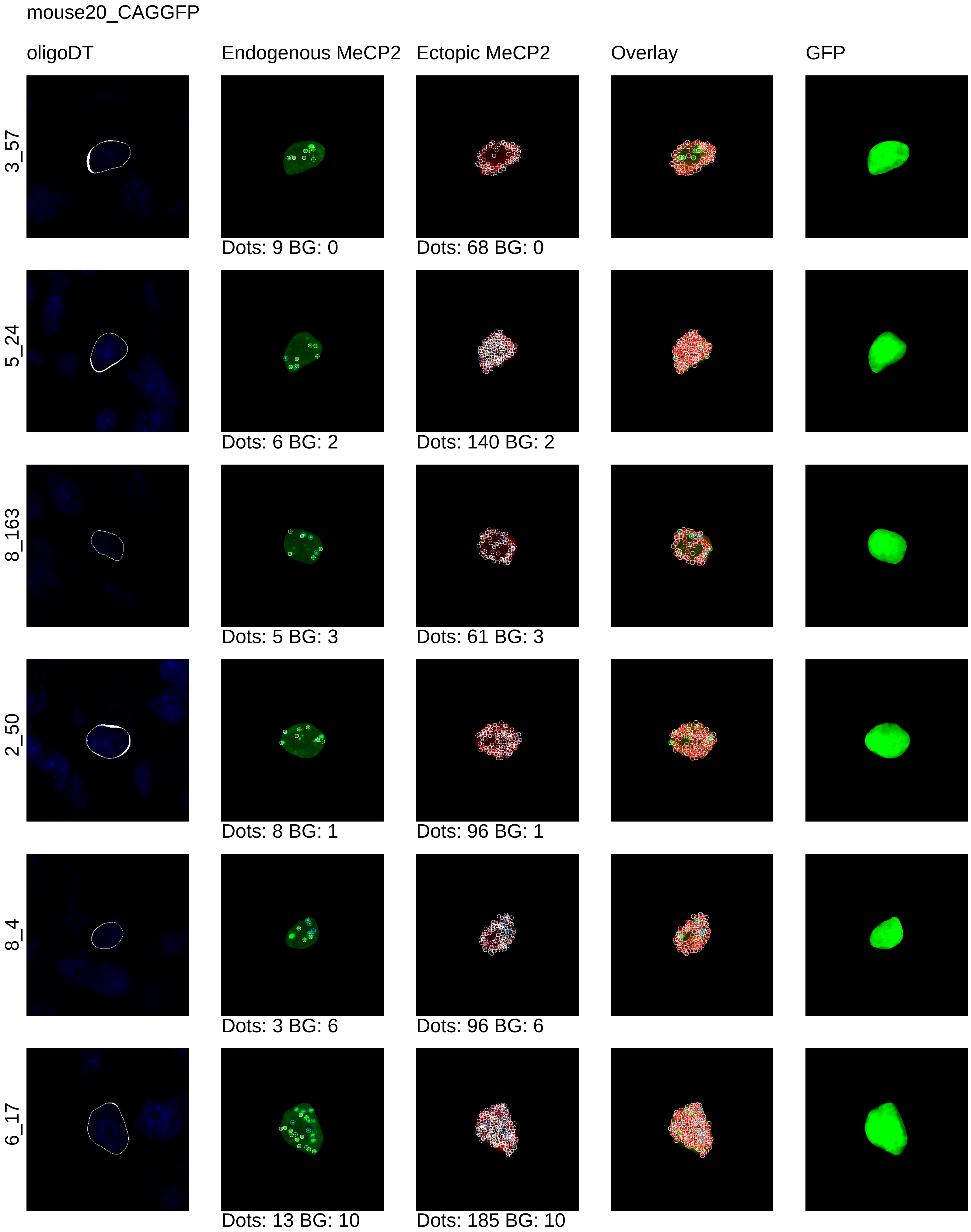

mouse20_CAGGFP
oligoDT
Endogenous MeCP2
Ectopic MeCP2
Overlay
GFP
3_57
Dots: 9 BG: 0
Dots: 68 BG: 0
5_24
Dots: 6 BG: 2
Dots: 140 BG: 2
8_163
Dots: 5 BG: 3
Dots: 61 BG: 3
2_50
Dots: 8 BG: 1
Dots: 96 BG: 1
8_4
Dots: 3 BG: 6
Dots: 96 BG: 6
6_17
Dots: 13 BG: 10
Dots: 185 BG: 10

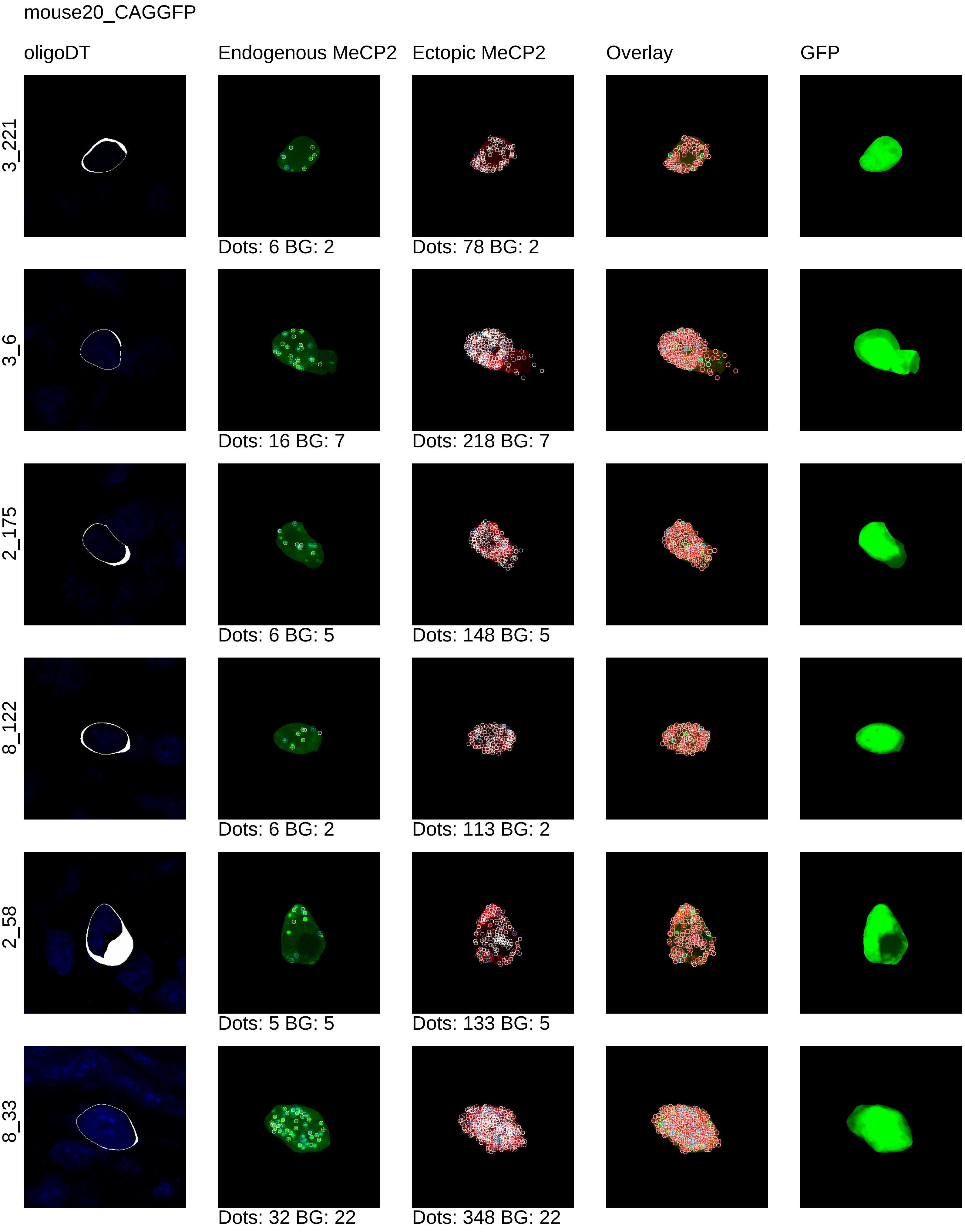
mouse20_CAGGFP
oligoDT
Endogenous MeCP2
Ectopic MeCP2
Overlay
GFP
3_221
Dots: 6 BG: 2
Dots: 78 BG: 2
3_6
Dots: 16 BG: 7
Dots: 218 BG: 7
2_175
Dots: 6 BG: 5
Dots: 148 BG: 5
8_122
Dots: 6 BG: 2
Dots: 113 BG: 2
2_58
Dots: 5 BG: 5
Dots: 133 BG: 5
8_33
Dots: 32 BG: 22
Dots: 348 BG: 22

mouse20_CAGGFP

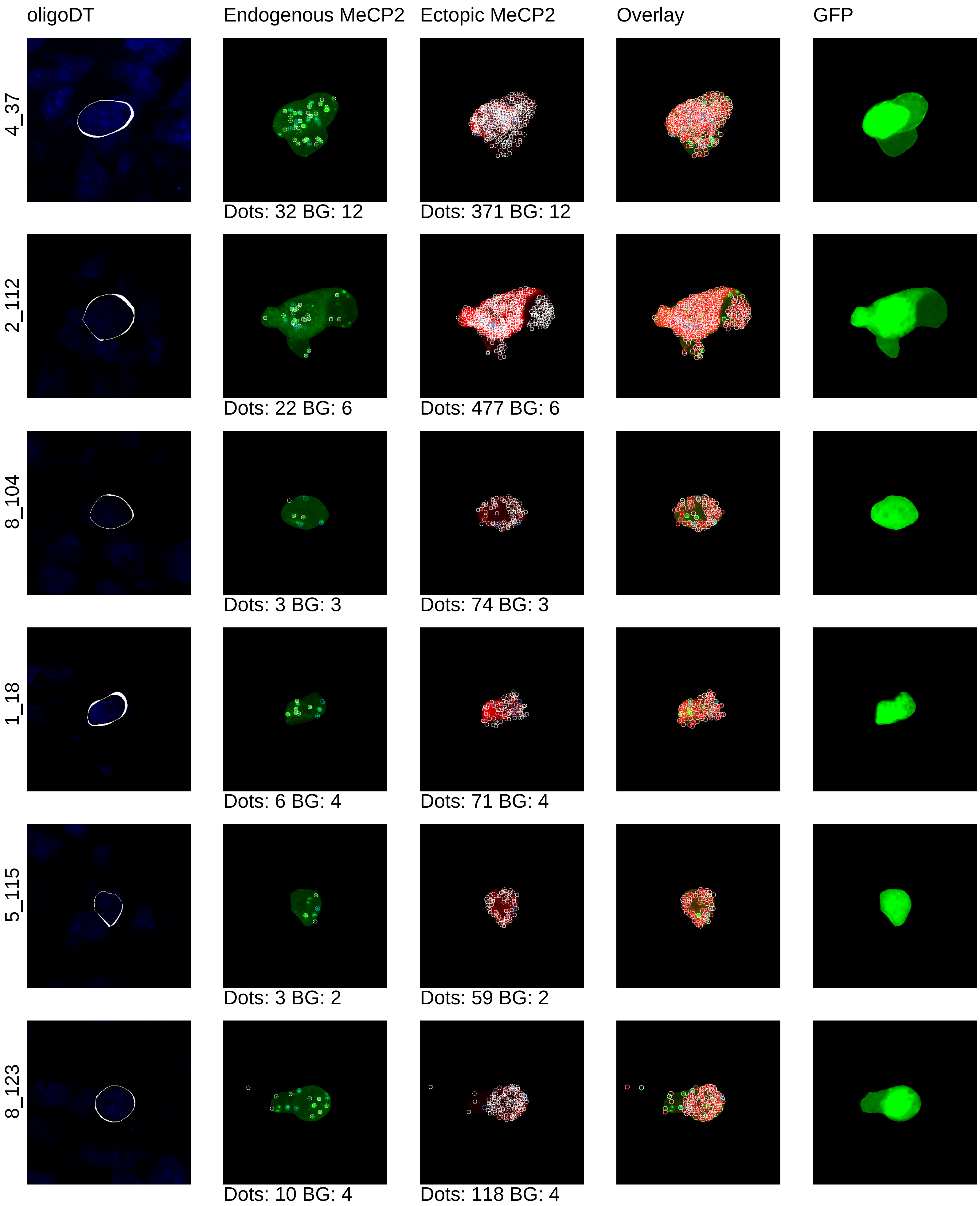

mouse20_CAGGFP

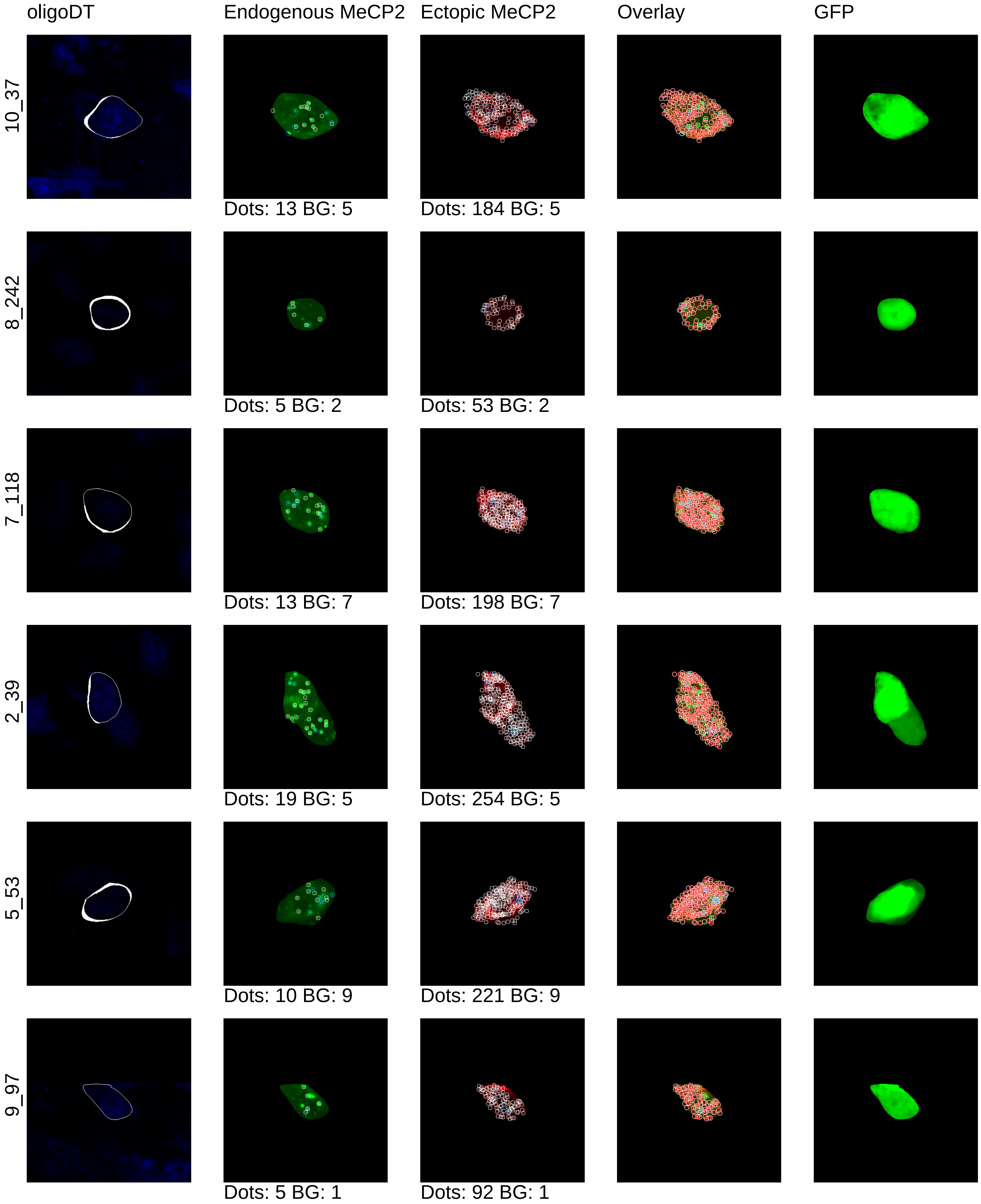

mouse20_CAGGFP

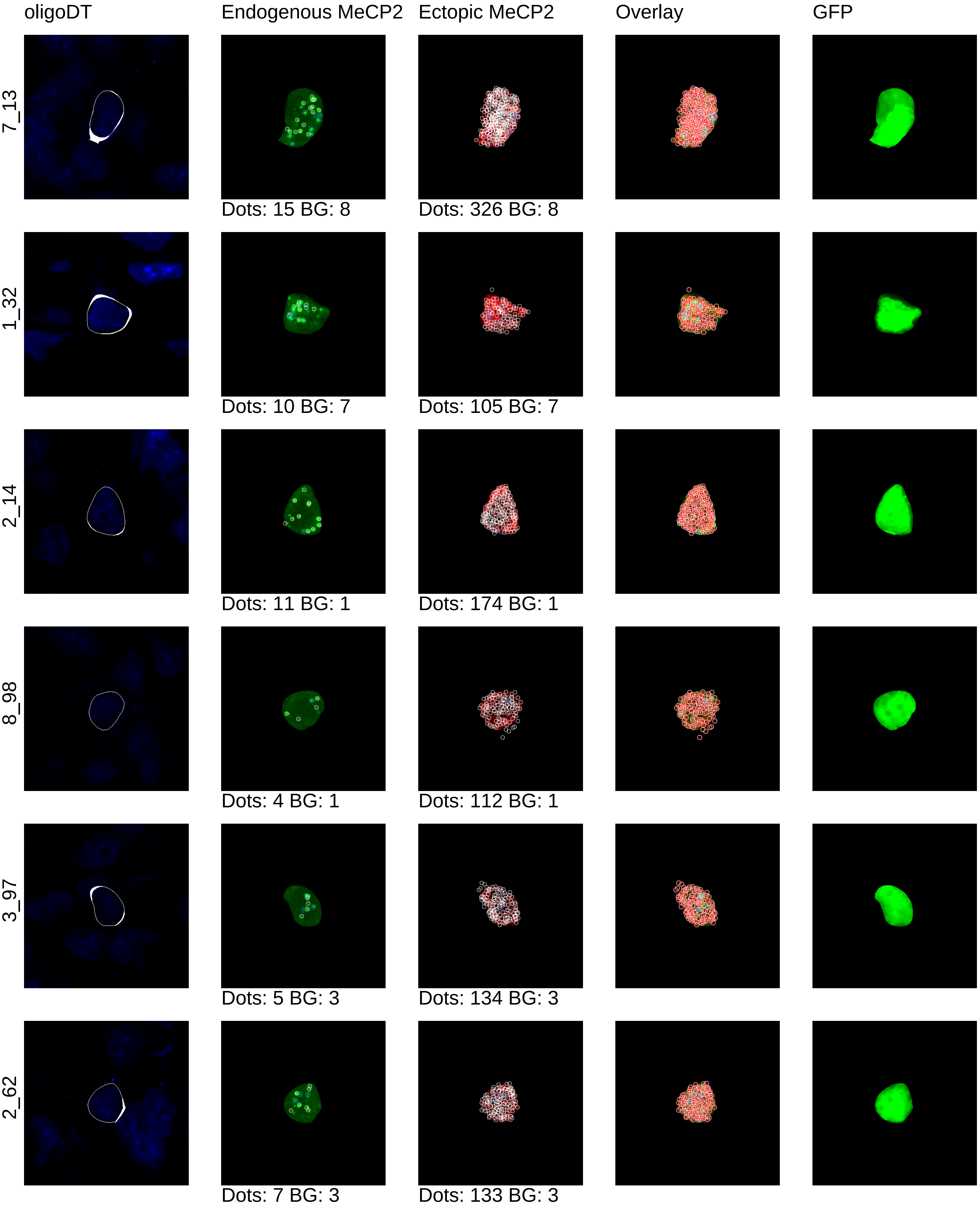

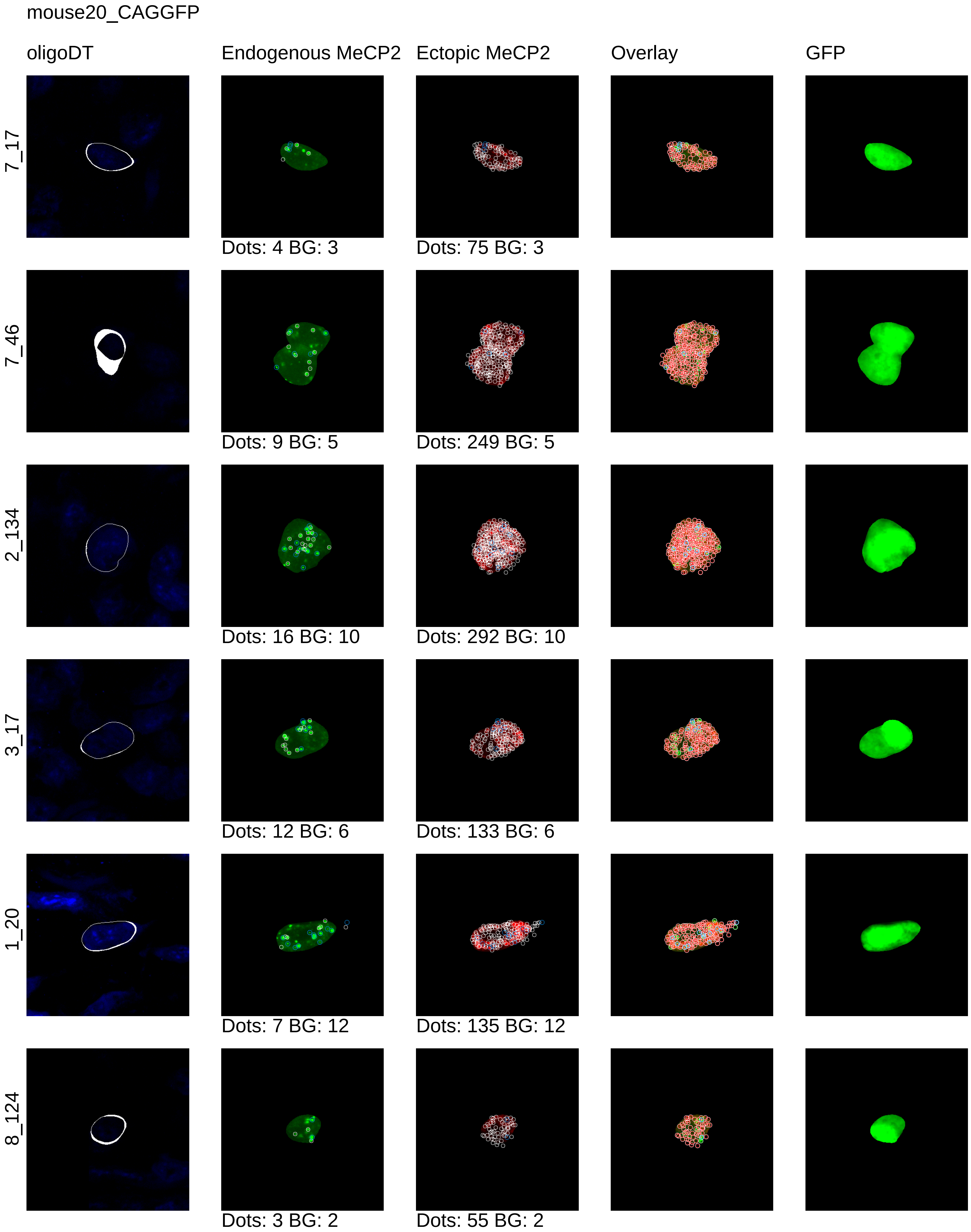
mouse20_CAGGFP
oligoDT
Endogenous MeCP2
Ectopic MeCP2
Overlay
GFP
7_17
Dots: 4 BG: 3
Dots: 75 BG: 3
7_46
Dots: 9 BG: 5
Dots: 249 BG: 5
2_134
Dots: 16 BG: 10
Dots: 292 BG: 10
3_17
Dots: 12 BG: 6
Dots: 133 BG: 6
1_20
Dots: 7 BG: 12
Dots: 135 BG: 12
8_124
Dots: 3 BG: 2
Dots: 55 BG: 2

mouse20_CAGGFP

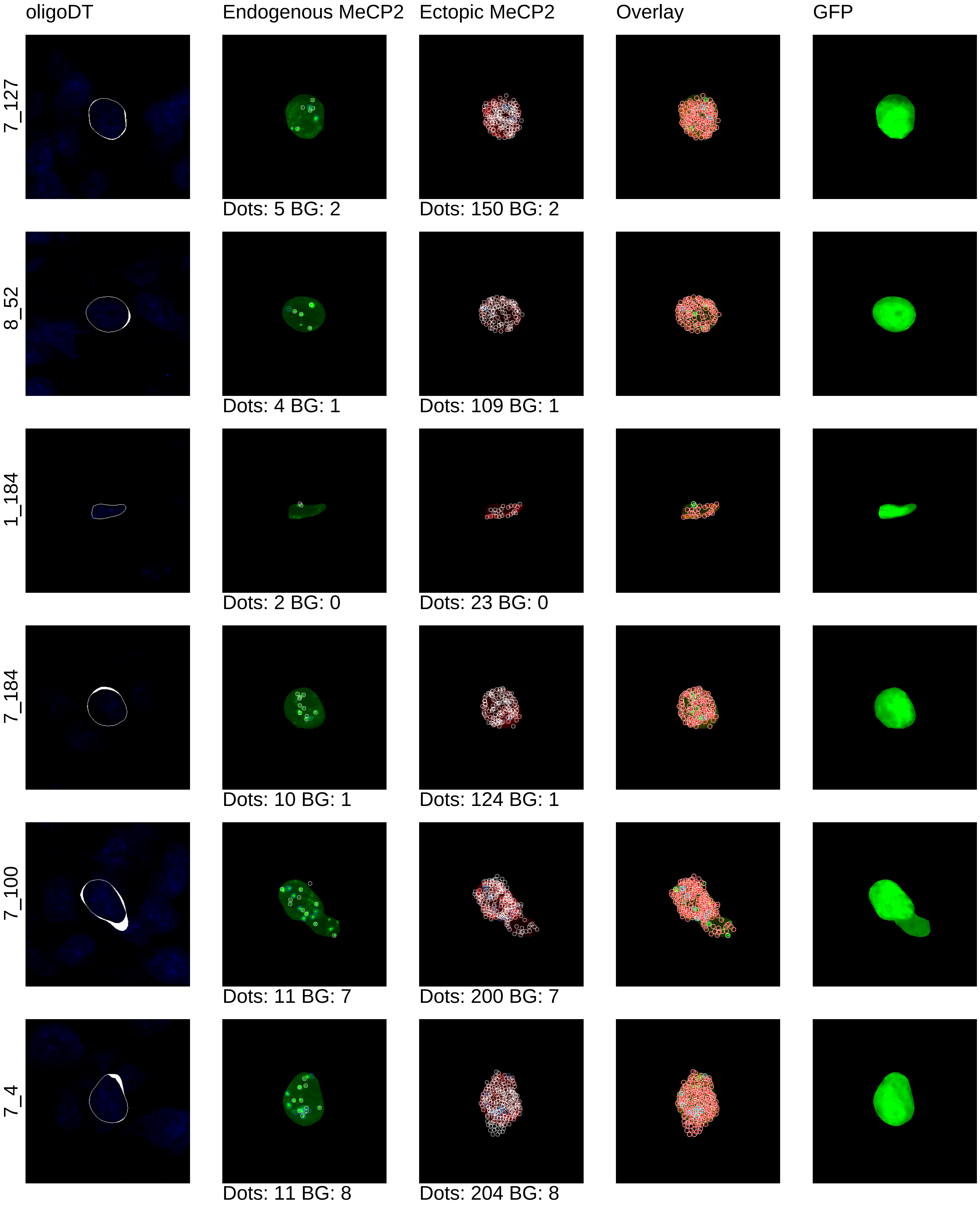

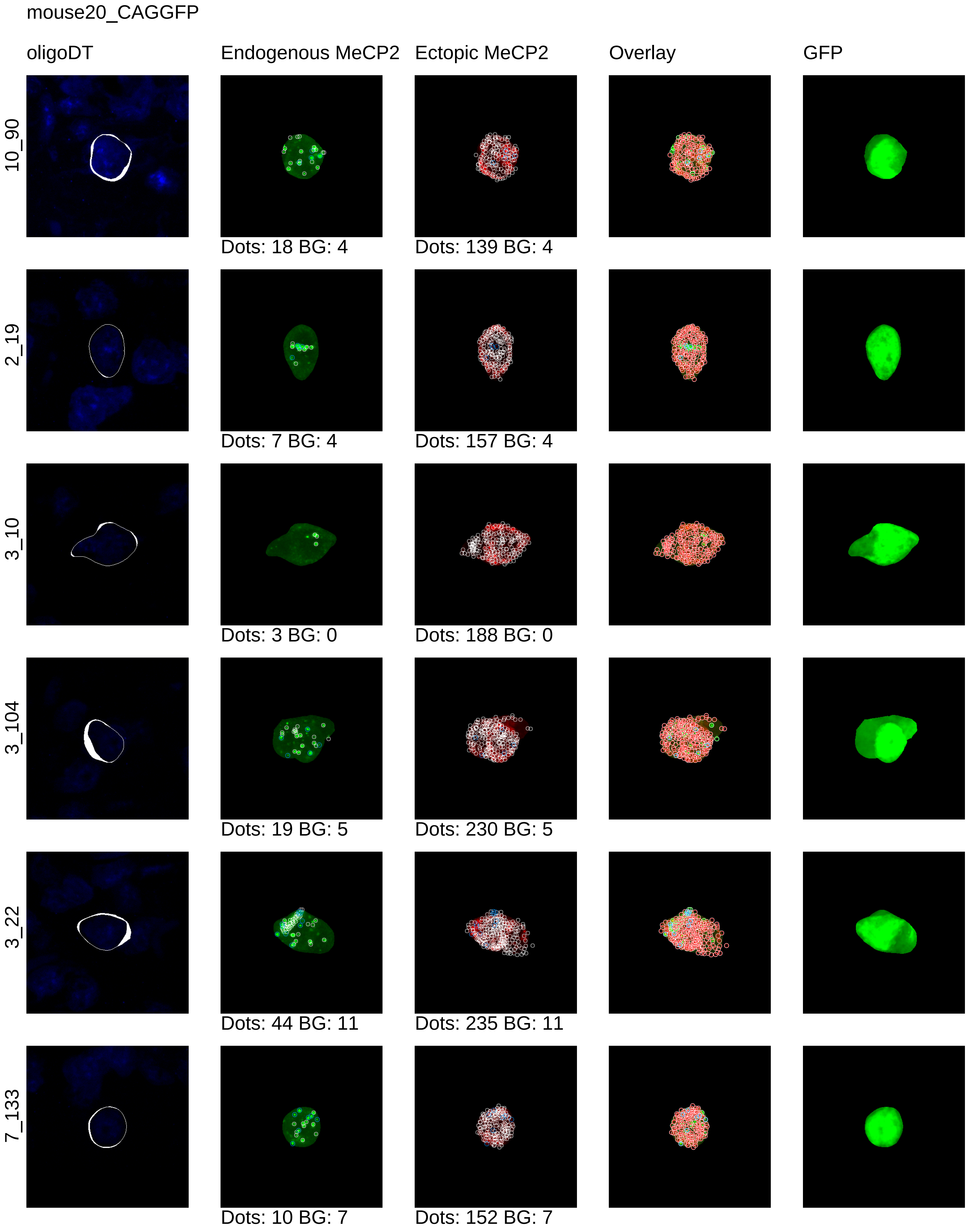
mouse20_CAGGFP
oligoDT
Endogenous MeCP2
Ectopic MeCP2
Overlay
GFP
10_90
Dots: 18 BG: 4
Dots: 139 BG: 4
2_19
Dots: 7 BG: 4
Dots: 157 BG: 4
3_10
Dots: 3 BG: 0
Dots: 188 BG: 0
3_104
Dots: 19 BG: 5
Dots: 230 BG: 5
3_22
Dots: 44 BG: 11
Dots: 235 BG: 11
7_133
Dots: 10 BG: 7
Dots: 152 BG: 7

mouse20_CAGGFP

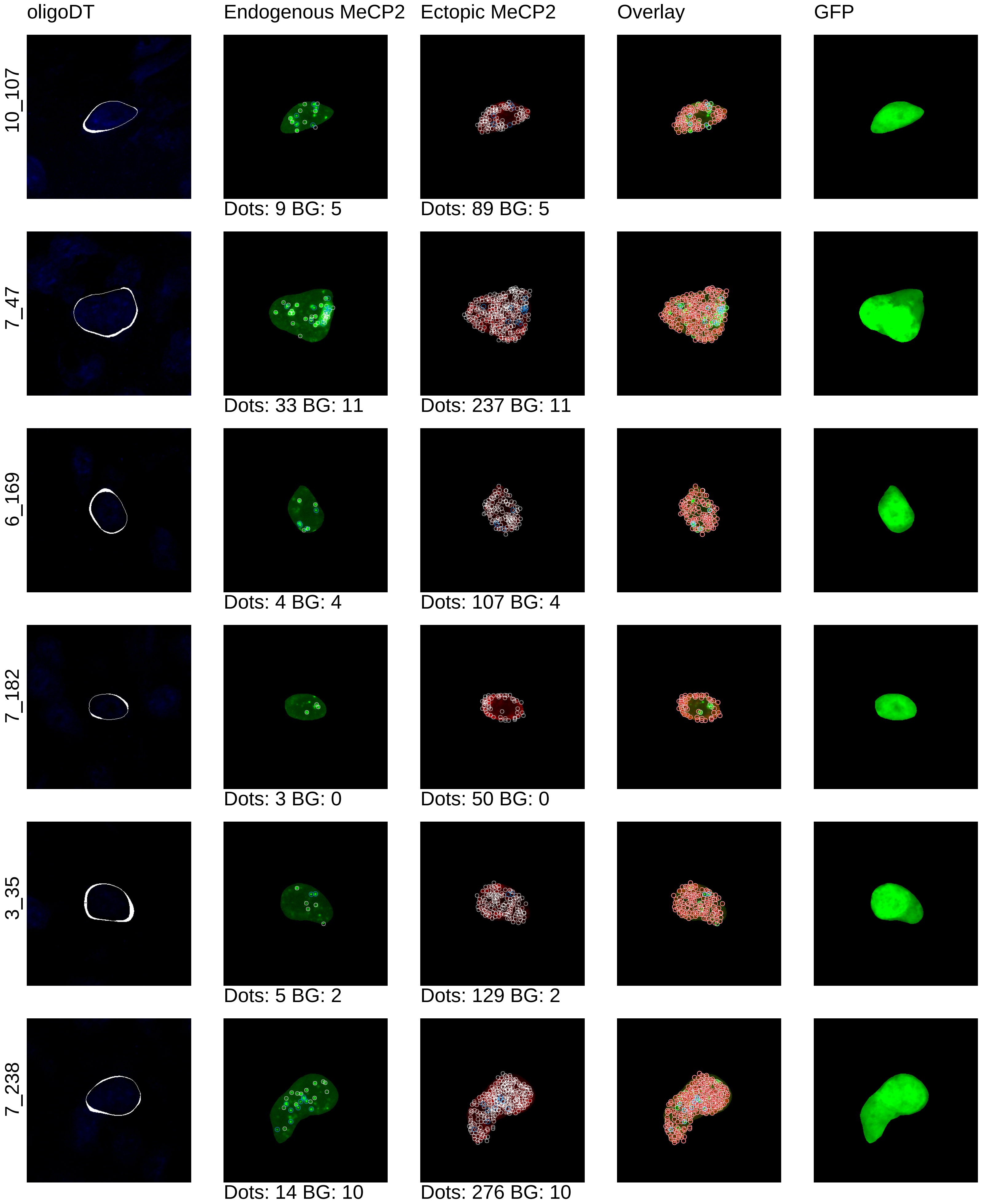

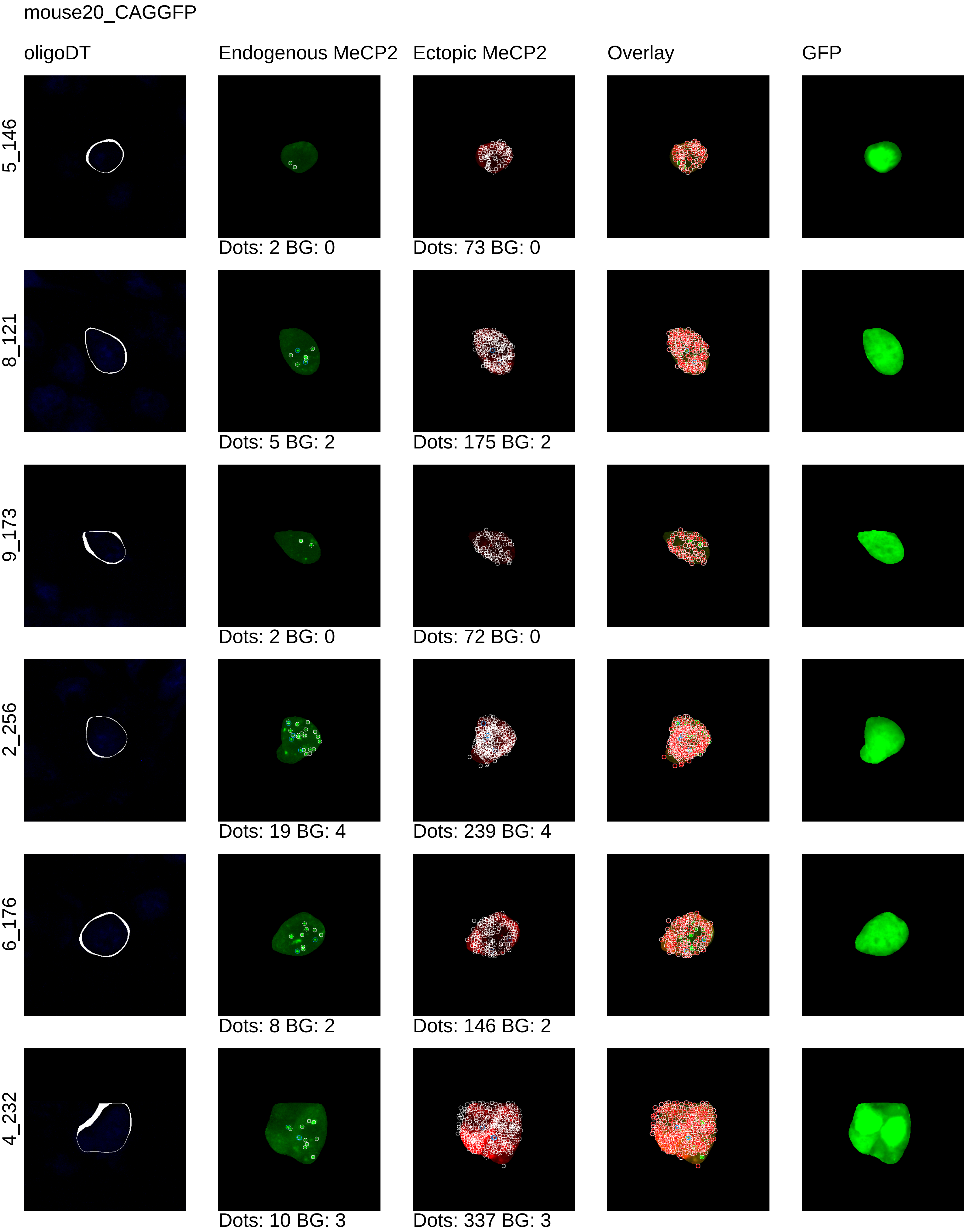
mouse20_CAGGFP
oligoDT
Endogenous MeCP2
Ectopic MeCP2
Overlay
GFP
5_146
Dots: 2 BG: 0
Dots: 73 BG: 0
8_121
Dots: 5 BG: 2
Dots: 175 BG: 2
9_173
Dots: 2 BG: 0
Dots: 72 BG: 0
2_256
Dots: 19 BG: 4
Dots: 239 BG: 4
6_176
Dots: 8 BG: 2
Dots: 146 BG: 2
4_232
Dots: 10 BG: 3
Dots: 337 BG: 3

mouse20_CAGGFP

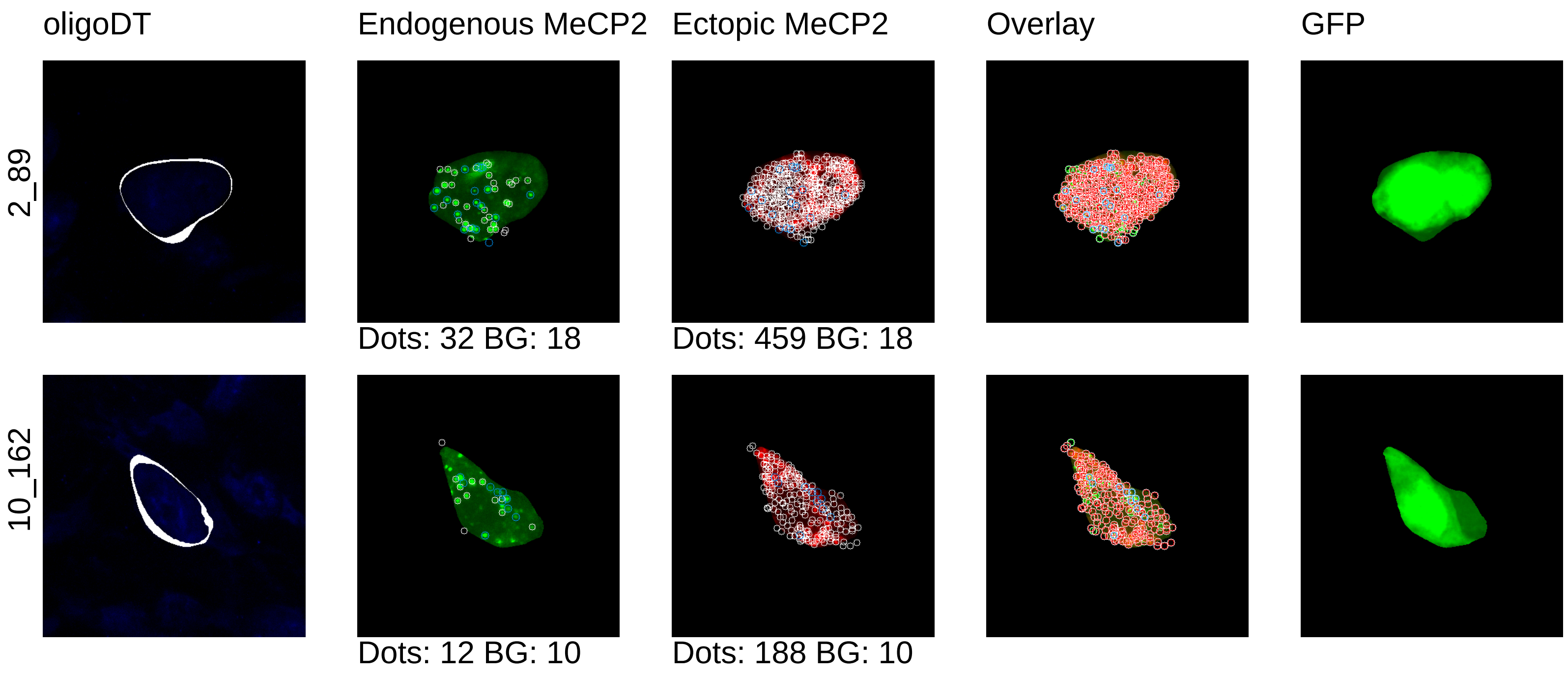
