## Supplementary material for "Synthetic dosage-compensating miRNA circuits allow precision gene therapy for Rett syndrome": Unregulated_cell_segmentation

mouse19_0x

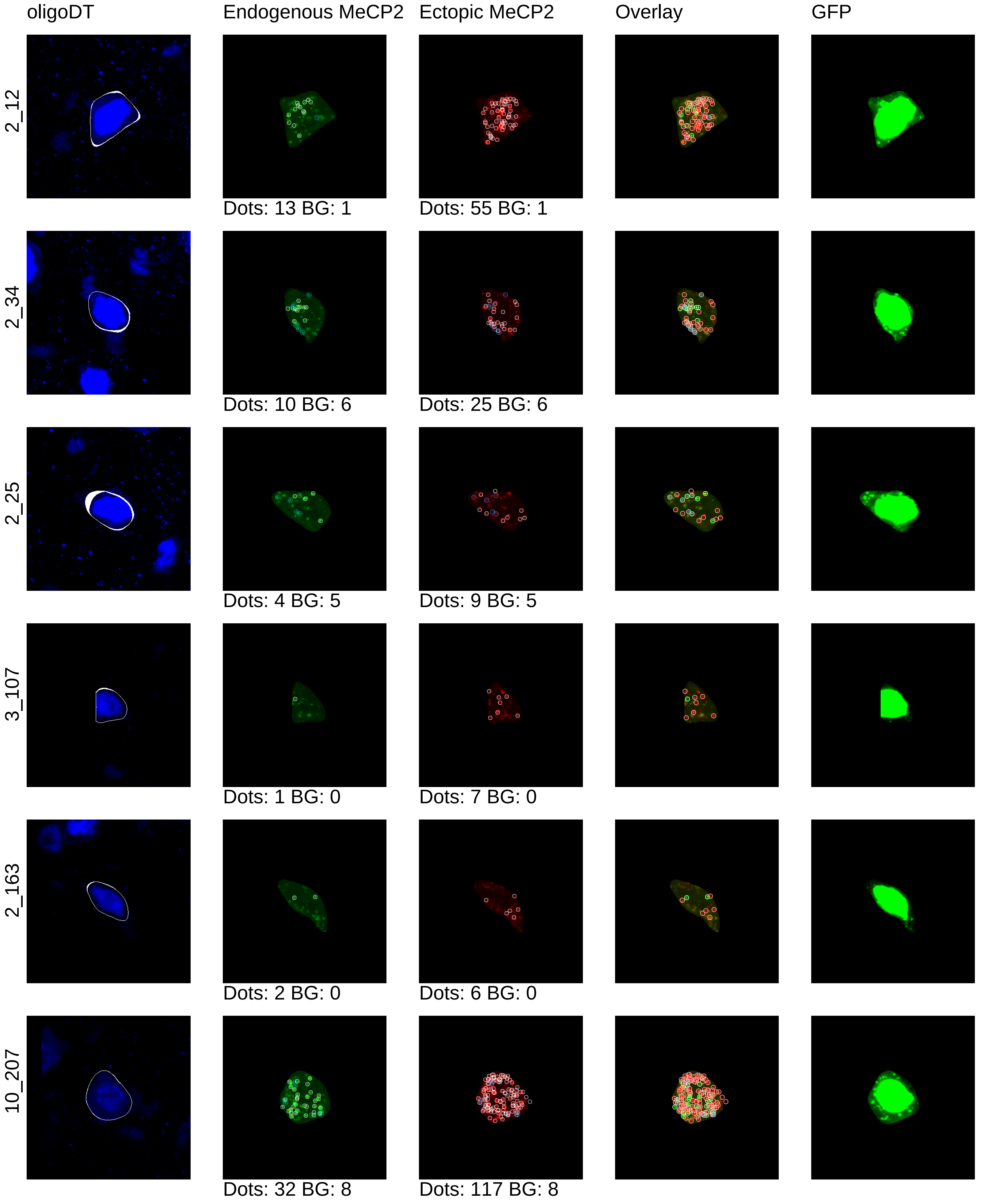

oligoDT
Endogenous MeCP2
Ectopic MeCP2
Overlay
GFP
2_12
Dots: 13 BG: 1
Dots: 55 BG: 1
2_34
Dots: 10 BG: 6
Dots: 25 BG: 6
2_25
Dots: 4 BG: 5
Dots: 9 BG: 5
3_107
Dots: 1 BG: 0
Dots: 7 BG: 0
2_163
Dots: 2 BG: 0
Dots: 6 BG: 0
10_207
Dots: 32 BG: 8
Dots: 117 BG: 8

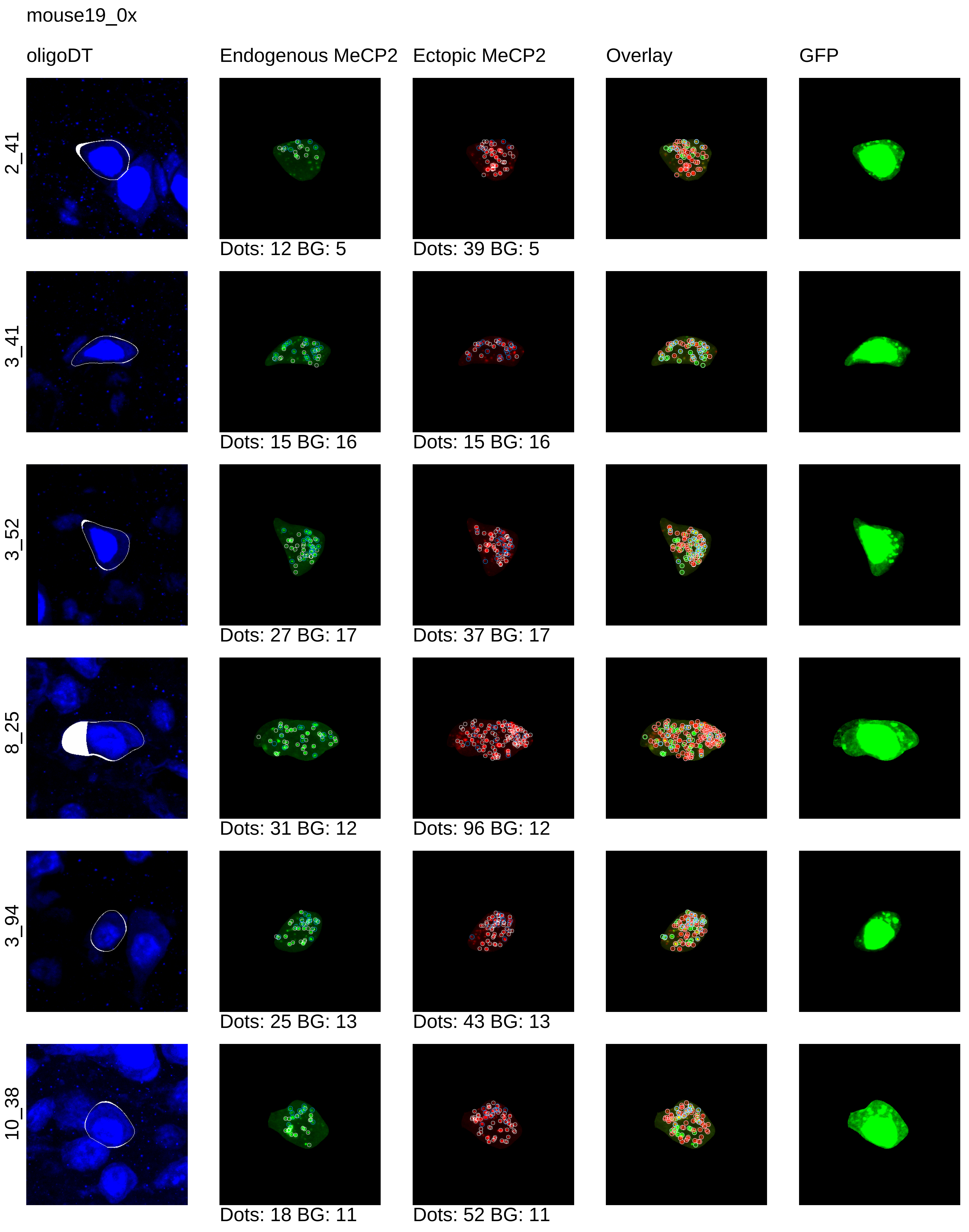
mouse19_0x
oligoDT
Endogenous MeCP2
Ectopic MeCP2
Overlay
GFP
2_41
Dots: 12 BG: 5
Dots: 39 BG: 5
3_41
Dots: 15 BG: 16
Dots: 15 BG: 16
3_52
Dots: 27 BG: 17
Dots: 37 BG: 17
8_25
Dots: 31 BG: 12
Dots: 96 BG: 12
3_94
Dots: 25 BG: 13
Dots: 43 BG: 13
10_38
Dots: 18 BG: 11
Dots: 52 BG: 11

mouse19_0x

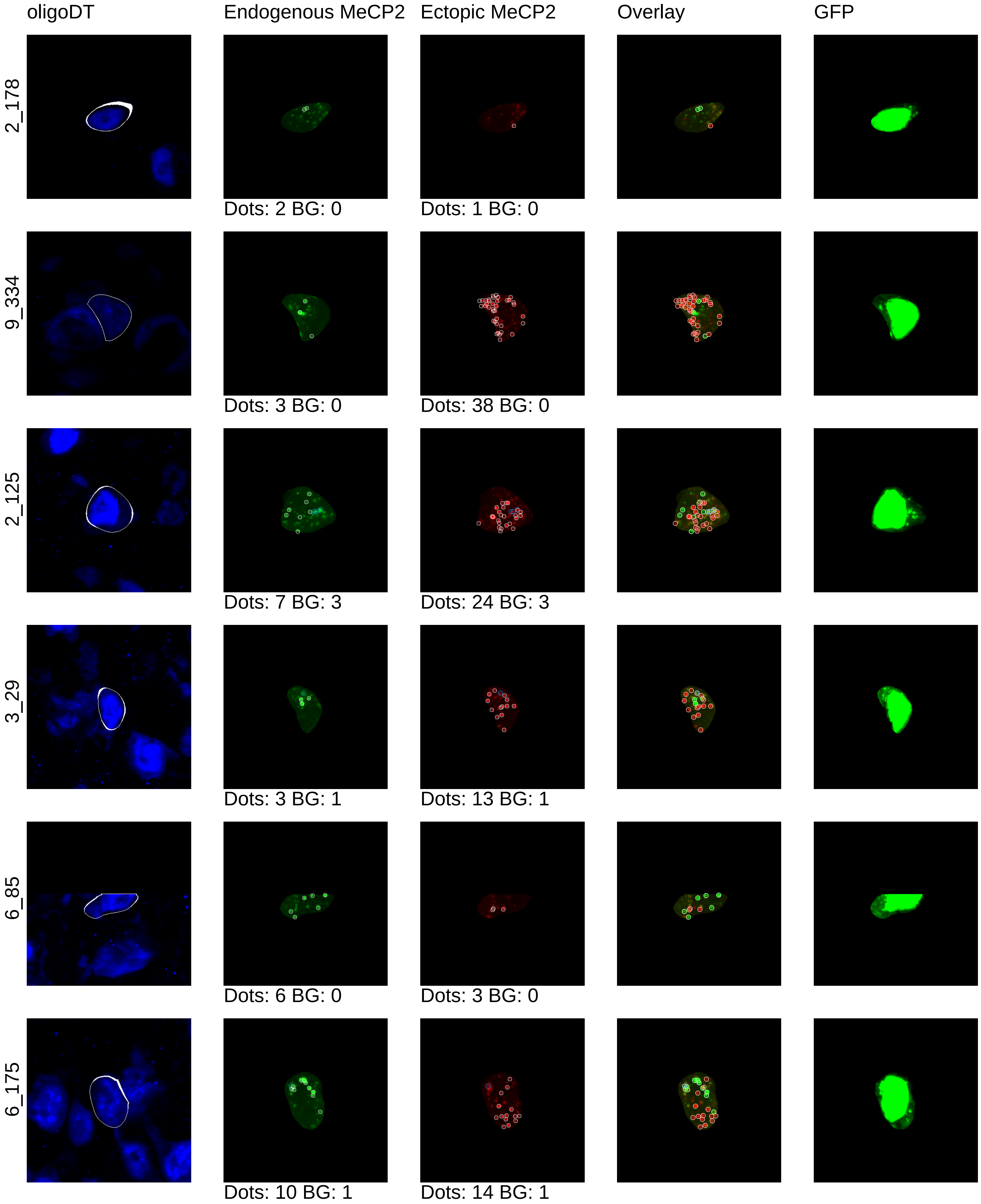

oligoDT
Endogenous MeCP2
Ectopic MeCP2
Overlay
GFP
2_178
Dots: 2 BG: 0
Dots: 1 BG: 0
9_334
Dots: 3 BG: 0
Dots: 38 BG: 0
2_125
Dots: 7 BG: 3
Dots: 24 BG: 3
3_29
Dots: 3 BG: 1
Dots: 13 BG: 1
6_85
Dots: 6 BG: 0
Dots: 3 BG: 0
6_175
Dots: 10 BG: 1
Dots: 14 BG: 1

mouse19_0x

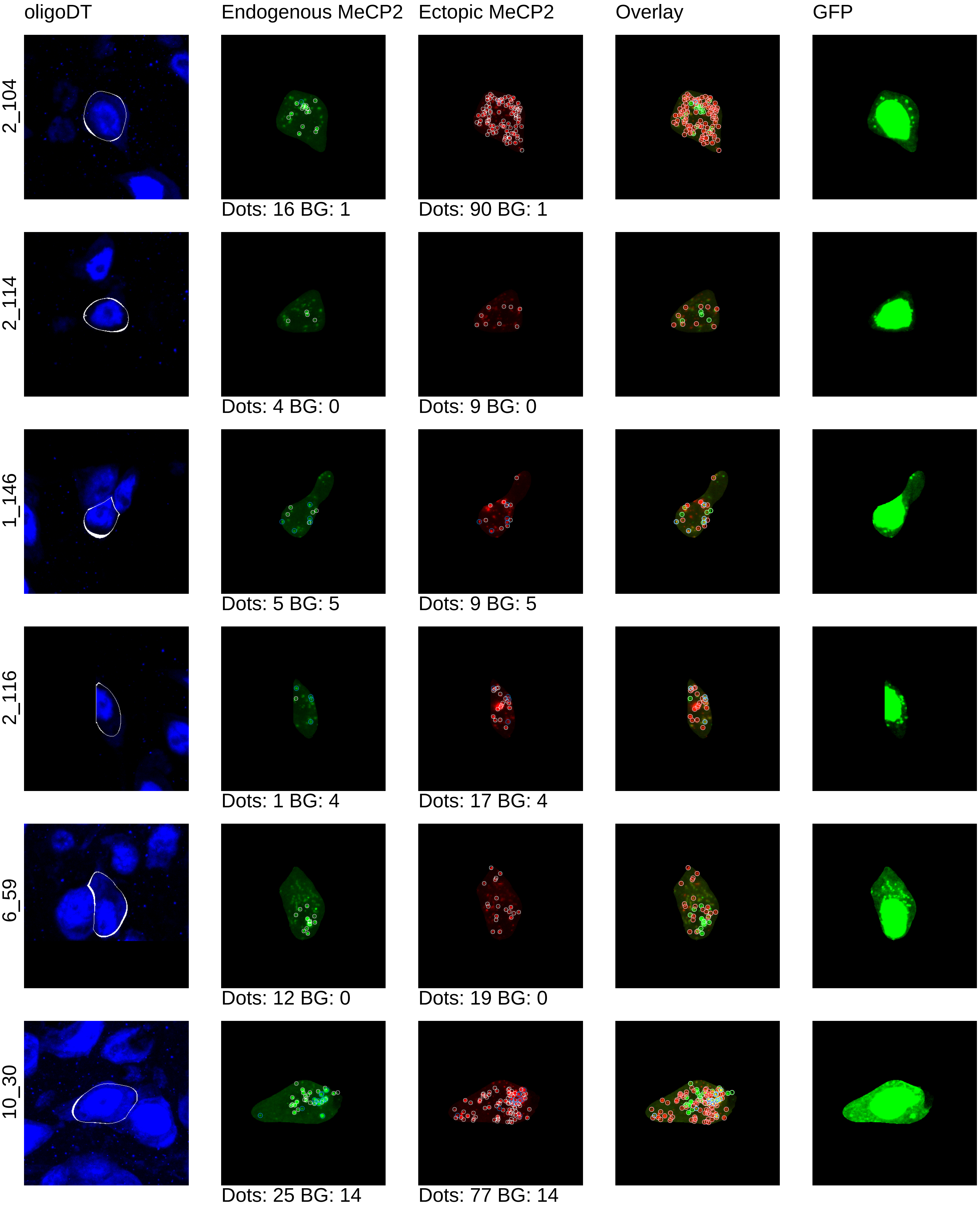
oligoDT
Endogenous MeCP2
Ectopic MeCP2
Overlay
GFP
2_104
Dots: 16 BG: 1
Dots: 90 BG: 1
2_114
Dots: 4 BG: 0
Dots: 9 BG: 0
1_146
Dots: 5 BG: 5
Dots: 9 BG: 5
2_116
Dots: 1 BG: 4
Dots: 17 BG: 4
6_59
Dots: 12 BG: 0
Dots: 19 BG: 0
10_30
Dots: 25 BG: 14
Dots: 77 BG: 14

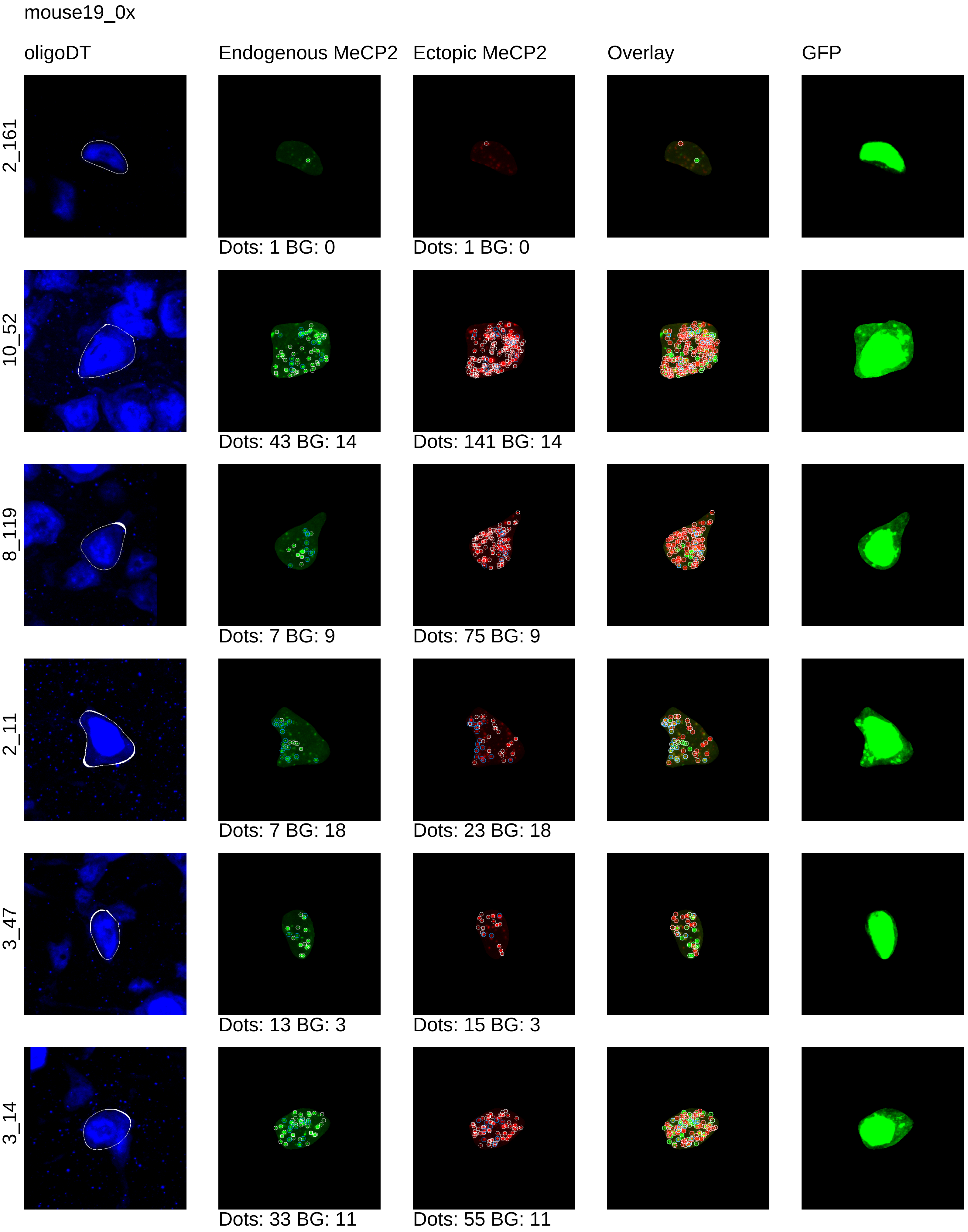
mouse19_0x
oligoDT
Endogenous MeCP2
Ectopic MeCP2
Overlay
GFP
2_161
Dots: 1 BG: 0
Dots: 1 BG: 0
10_52
Dots: 43 BG: 14
Dots: 141 BG: 14
8_119
Dots: 7 BG: 9
Dots: 75 BG: 9
2_11
Dots: 7 BG: 18
Dots: 23 BG: 18
3_47
Dots: 13 BG: 3
Dots: 15 BG: 3
3_14
Dots: 33 BG: 11
Dots: 55 BG: 11

mouse19_0x

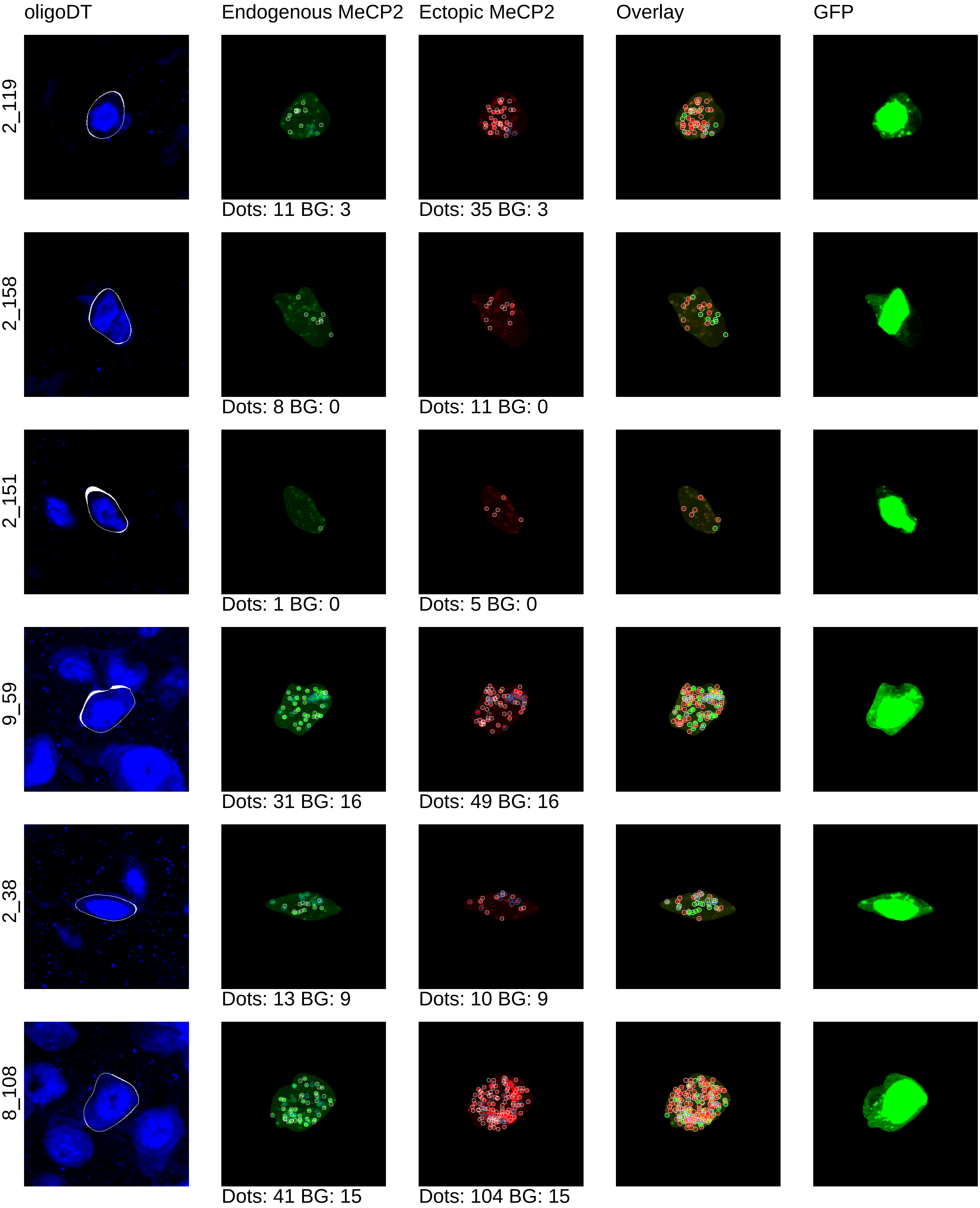

oligoDT
Endogenous MeCP2
Ectopic MeCP2
Overlay
GFP
2_119
Dots: 11 BG: 3
Dots: 35 BG: 3
2_158
Dots: 8 BG: 0
Dots: 11 BG: 0
2_151
Dots: 1 BG: 0
Dots: 5 BG: 0
9_59
Dots: 31 BG: 16
Dots: 49 BG: 16
2_38
Dots: 13 BG: 9
Dots: 10 BG: 9
8_108
Dots: 41 BG: 15
Dots: 104 BG: 15

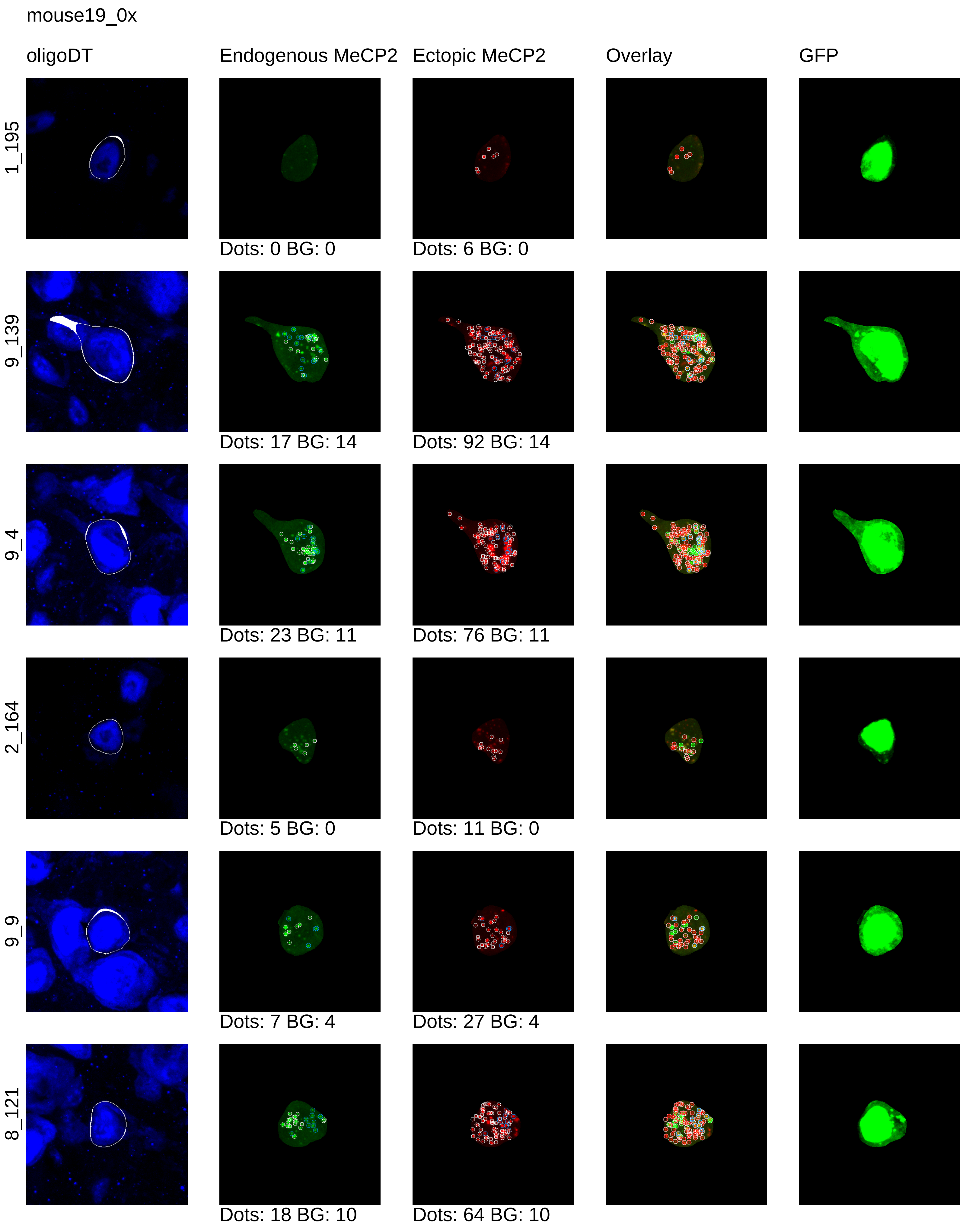
mouse19_0x
oligoDT
Endogenous MeCP2
Ectopic MeCP2
Overlay
GFP
1_195
Dots: 0 BG: 0
Dots: 6 BG: 0
9_139
Dots: 17 BG: 14
Dots: 92 BG: 14
9_4
Dots: 23 BG: 11
Dots: 76 BG: 11
2_164
Dots: 5 BG: 0
Dots: 11 BG: 0
9_9
Dots: 7 BG: 4
Dots: 27 BG: 4
8_121
Dots: 18 BG: 10
Dots: 64 BG: 10

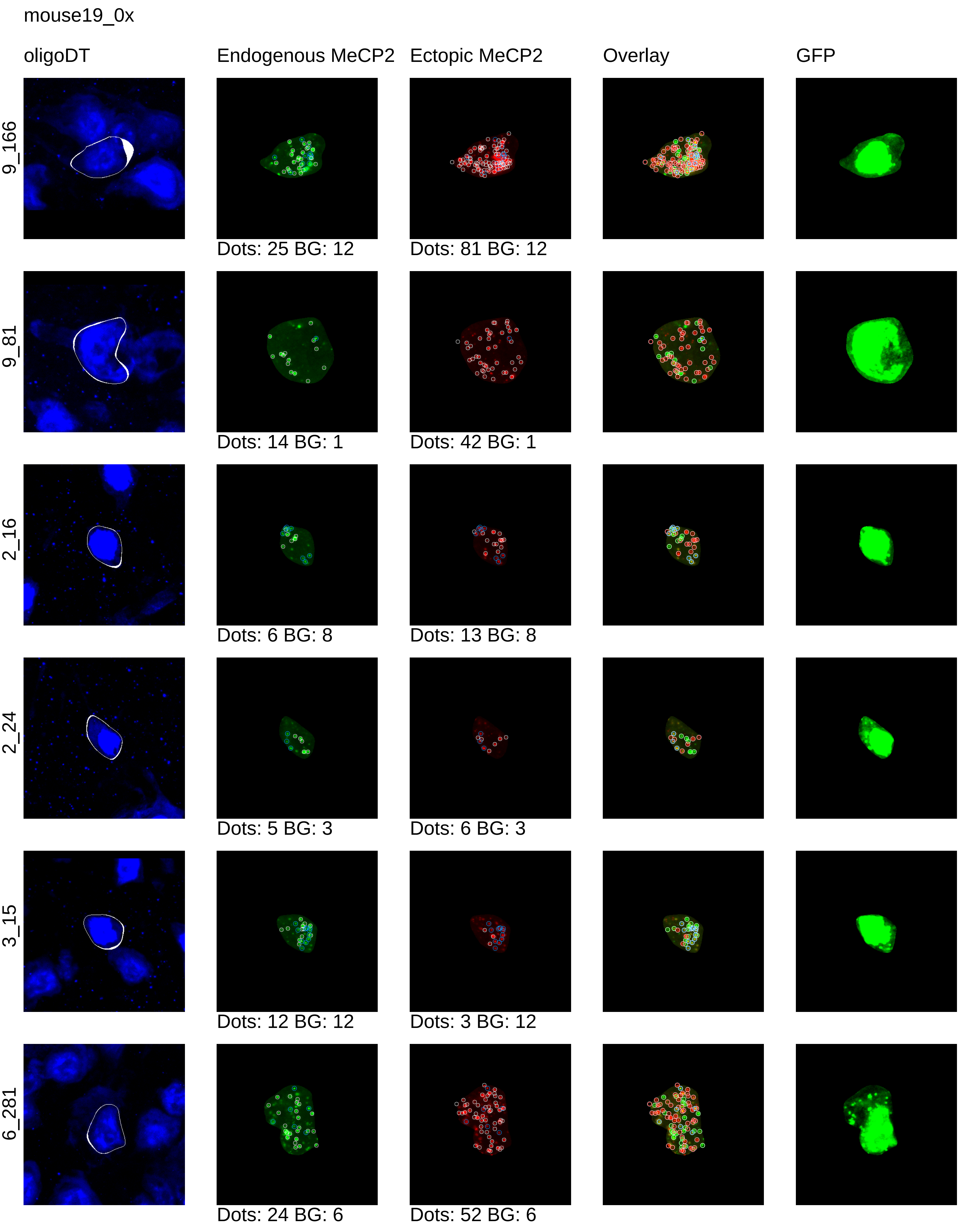
mouse19_0x
oligoDT
Endogenous MeCP2
Ectopic MeCP2
Overlay
GFP
9_166
Dots: 25 BG: 12
Dots: 81 BG: 12
9_81
Dots: 14 BG: 1
Dots: 42 BG: 1
2_16
Dots: 6 BG: 8
Dots: 13 BG: 8
2_24
Dots: 5 BG: 3
Dots: 6 BG: 3
3_15
Dots: 12 BG: 12
Dots: 3 BG: 12
6_281
Dots: 24 BG: 6
Dots: 52 BG: 6

mouse19_0x

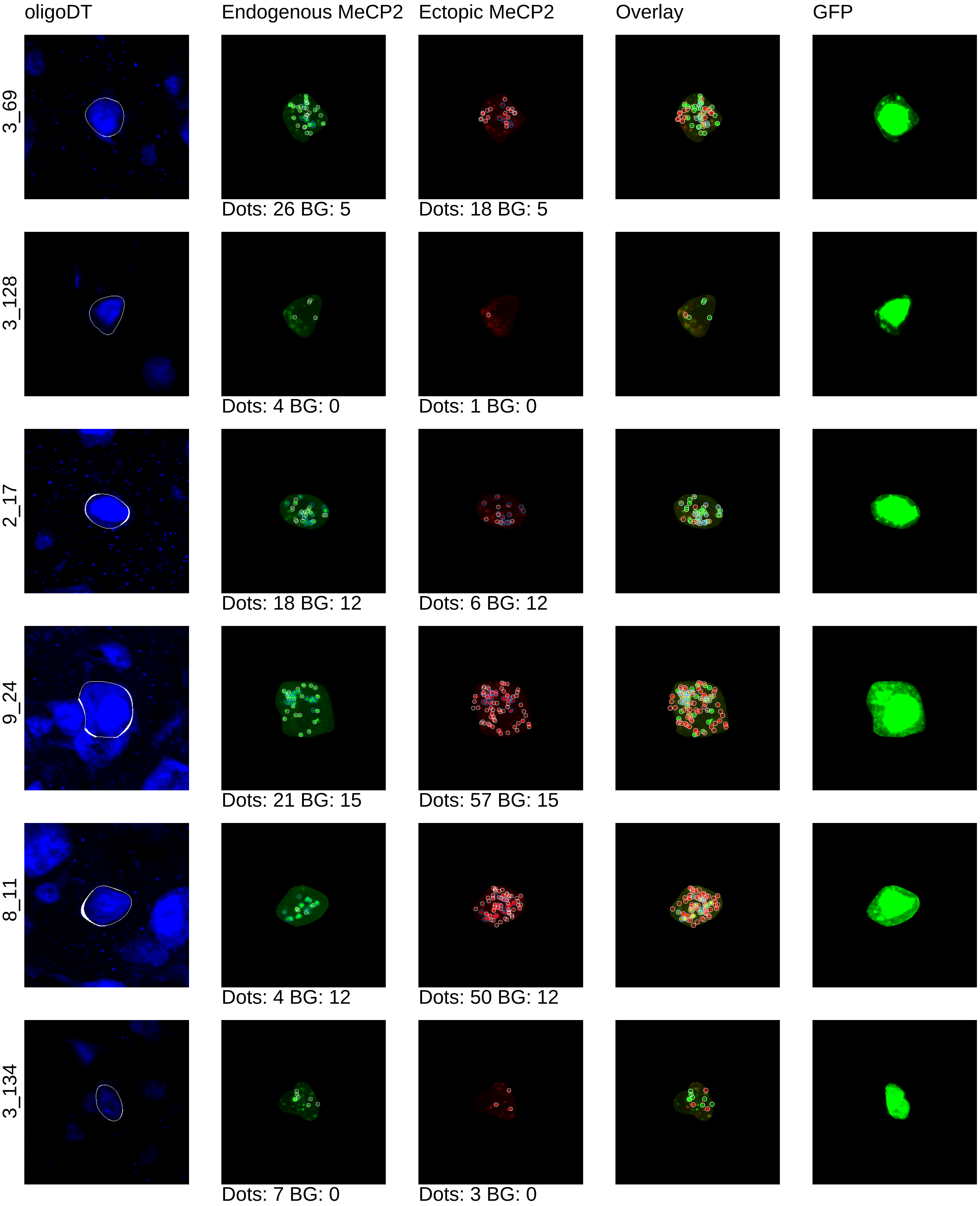
oligoDT
Endogenous MeCP2
Ectopic MeCP2
Overlay
GFP
3_69
Dots: 26 BG: 5
Dots: 18 BG: 5
3_128
Dots: 4 BG: 0
Dots: 1 BG: 0
2_17
Dots: 18 BG: 12
Dots: 6 BG: 12
9_24
Dots: 21 BG: 15
Dots: 57 BG: 15
8_11
Dots: 4 BG: 12
Dots: 50 BG: 12
3_134
Dots: 7 BG: 0
Dots: 3 BG: 0

mouse19_0x

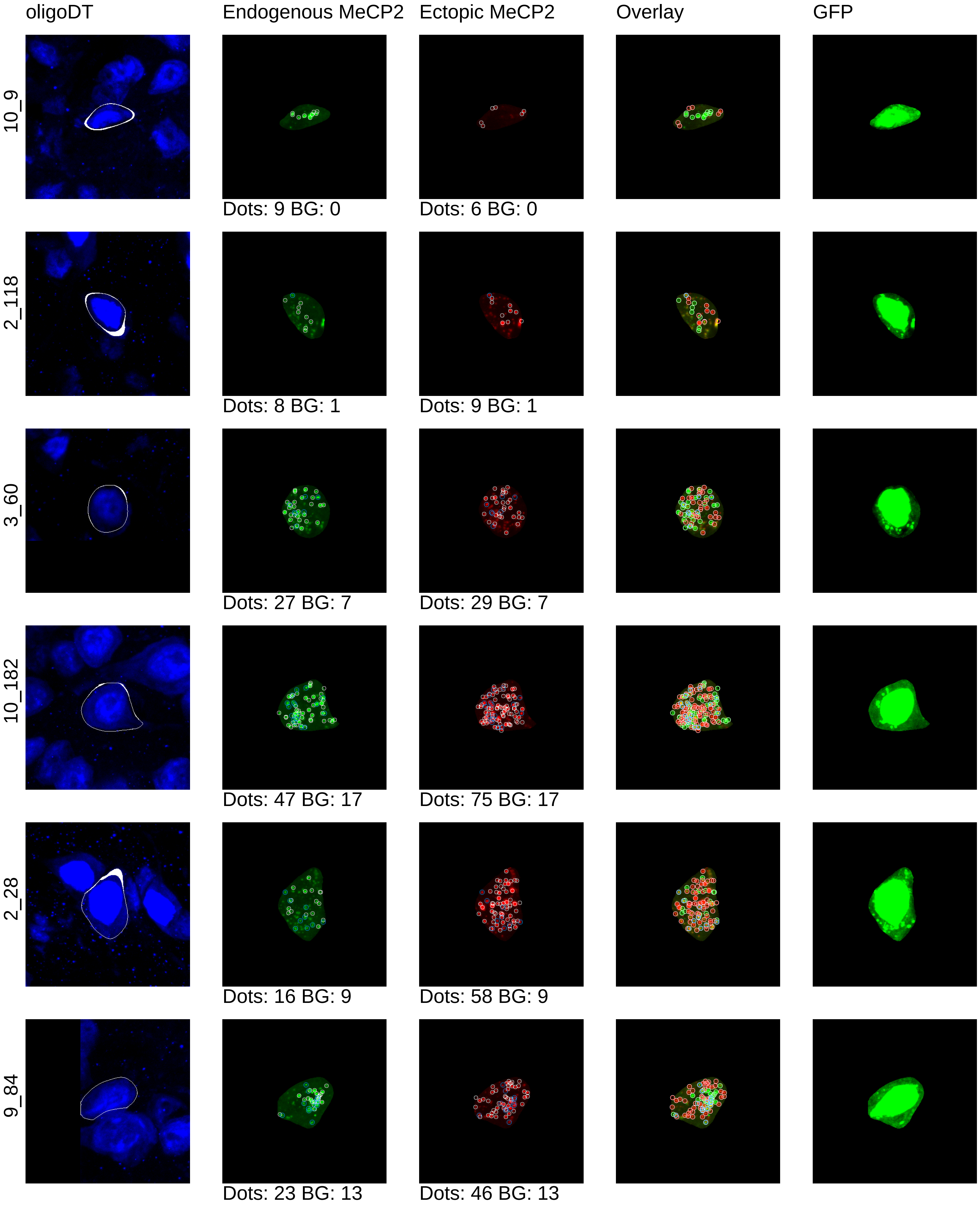

oligoDT
Endogenous MeCP2
Ectopic MeCP2
Overlay
GFP
10_9
Dots: 9 BG: 0
Dots: 6 BG: 0
2_118
Dots: 8 BG: 1
Dots: 9 BG: 1
3_60
Dots: 27 BG: 7
Dots: 29 BG: 7
10_182
Dots: 47 BG: 17
Dots: 75 BG: 17
2_28
Dots: 16 BG: 9
Dots: 58 BG: 9
9_84
Dots: 23 BG: 13
Dots: 46 BG: 13

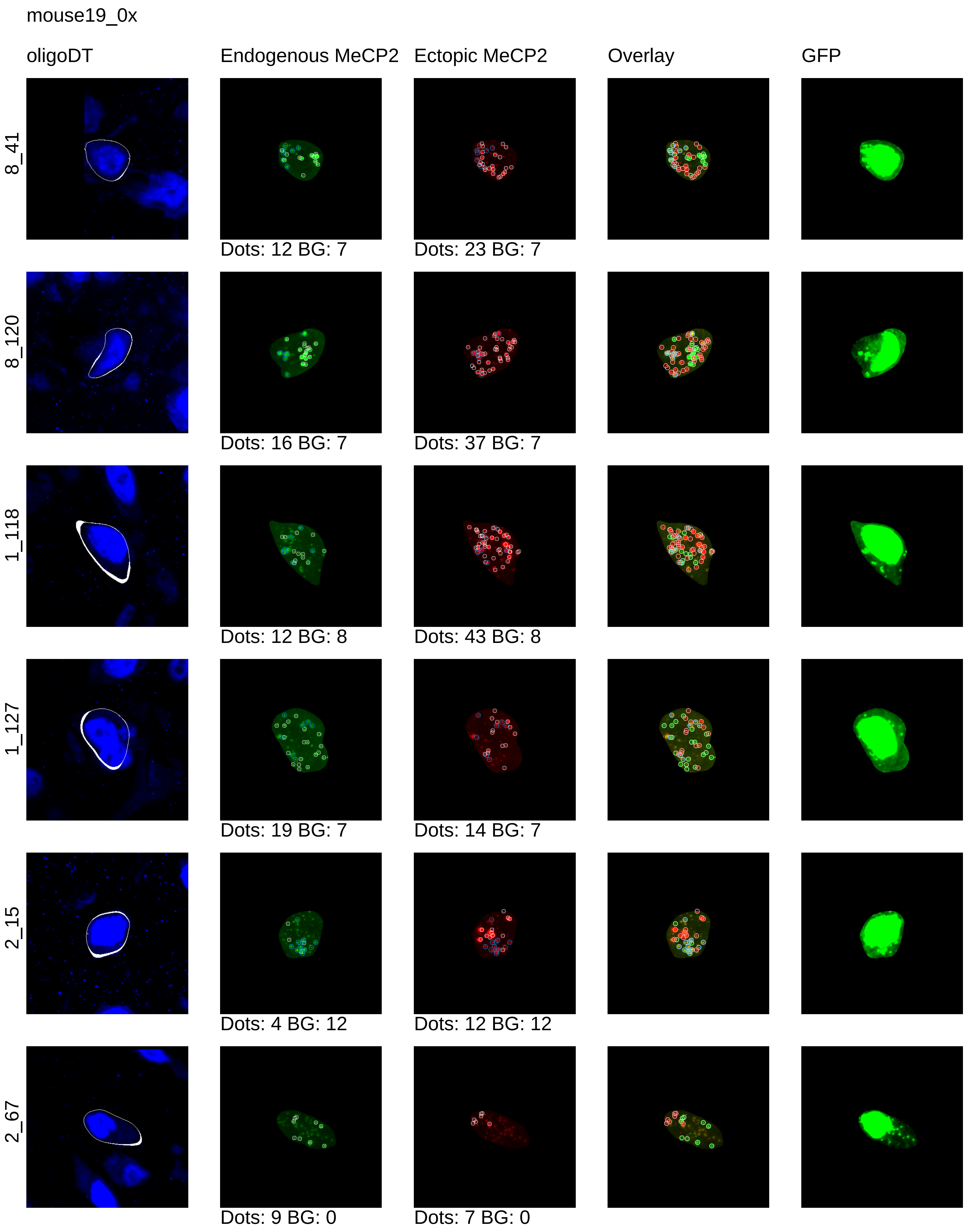
mouse19_0x
oligoDT
Endogenous MeCP2
Ectopic MeCP2
Overlay
GFP
8_41
Dots: 12 BG: 7
Dots: 23 BG: 7
8_120
Dots: 16 BG: 7
Dots: 37 BG: 7
1_118
Dots: 12 BG: 8
Dots: 43 BG: 8
1_127
Dots: 19 BG: 7
Dots: 14 BG: 7
2_15
Dots: 4 BG: 12
Dots: 12 BG: 12
2_67
Dots: 9 BG: 0
Dots: 7 BG: 0

mouse19_0x

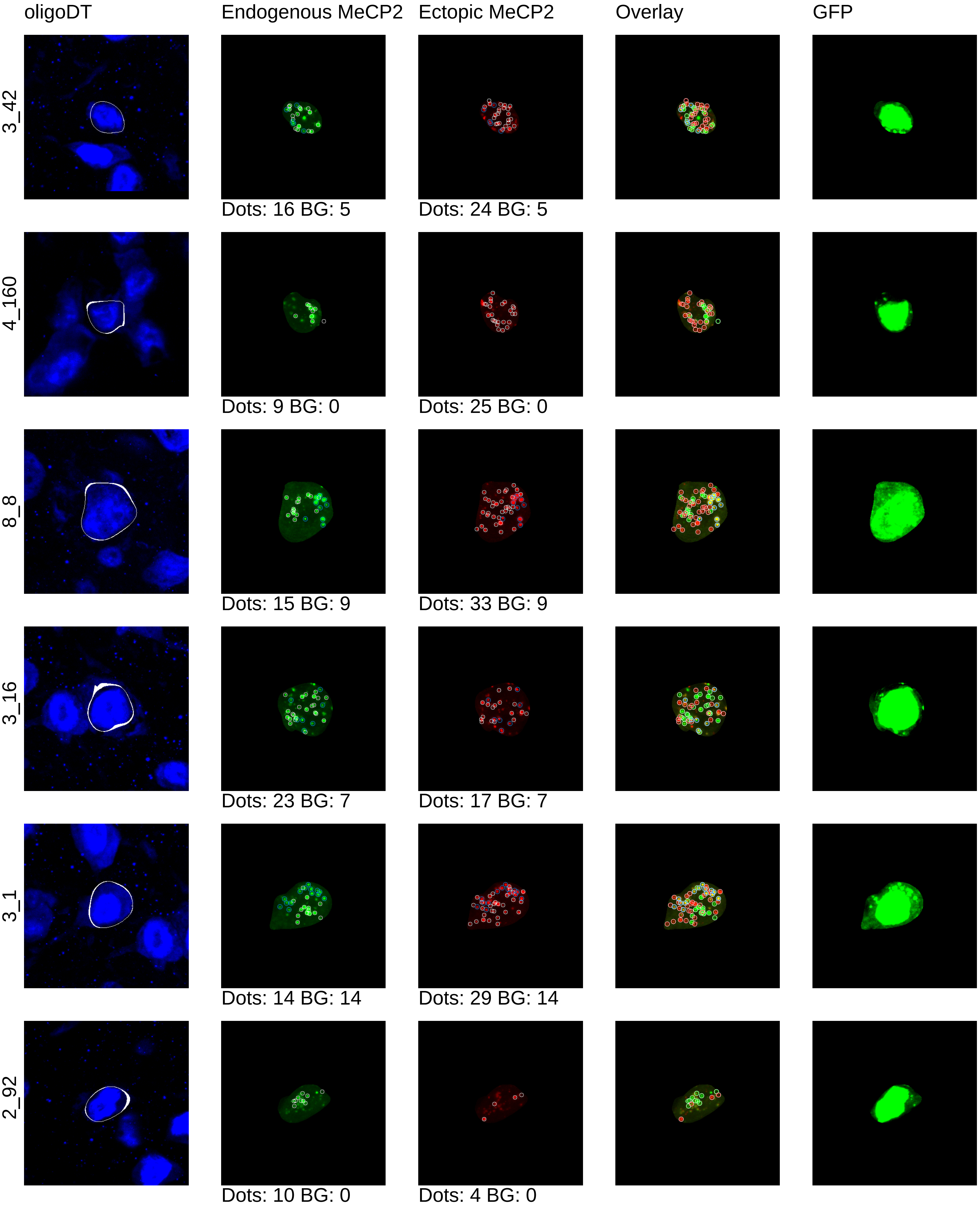

oligoDT
Endogenous MeCP2
Ectopic MeCP2
Overlay
GFP
3_42
Dots: 16 BG: 5
Dots: 24 BG: 5
4_160
Dots: 9 BG: 0
Dots: 25 BG: 0
8_8
Dots: 15 BG: 9
Dots: 33 BG: 9
3_16
Dots: 23 BG: 7
Dots: 17 BG: 7
3_1
Dots: 14 BG: 14
Dots: 29 BG: 14
2_92
Dots: 10 BG: 0
Dots: 4 BG: 0

mouse19_0x

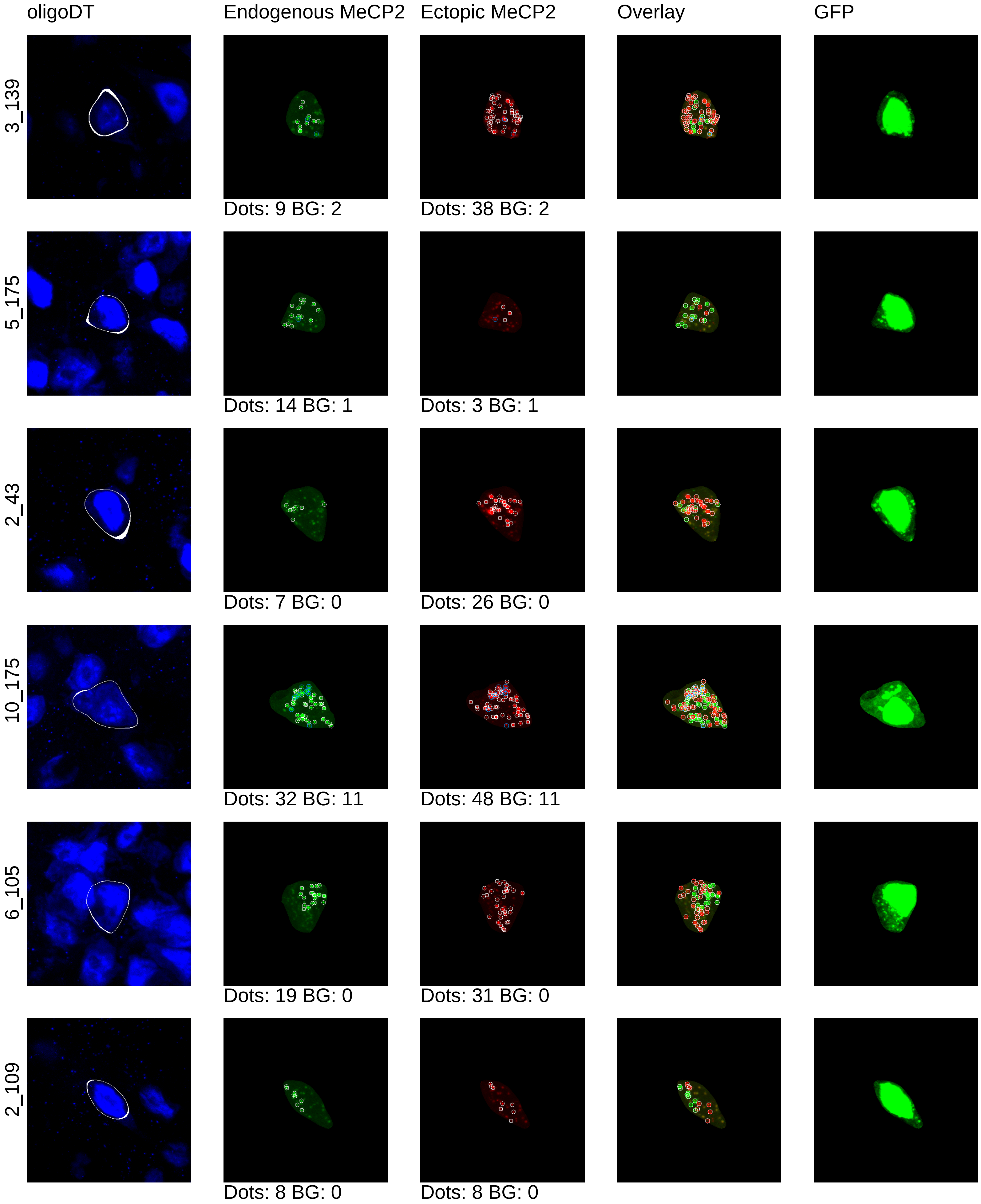
oligoDT
Endogenous MeCP2
Ectopic MeCP2
Overlay
GFP
3_139
Dots: 9 BG: 2
Dots: 38 BG: 2
5_175
Dots: 14 BG: 1
Dots: 3 BG: 1
2_43
Dots: 7 BG: 0
Dots: 26 BG: 0
10_175
Dots: 32 BG: 11
Dots: 48 BG: 11
6_105
Dots: 19 BG: 0
Dots: 31 BG: 0
2_109
Dots: 8 BG: 0
Dots: 8 BG: 0

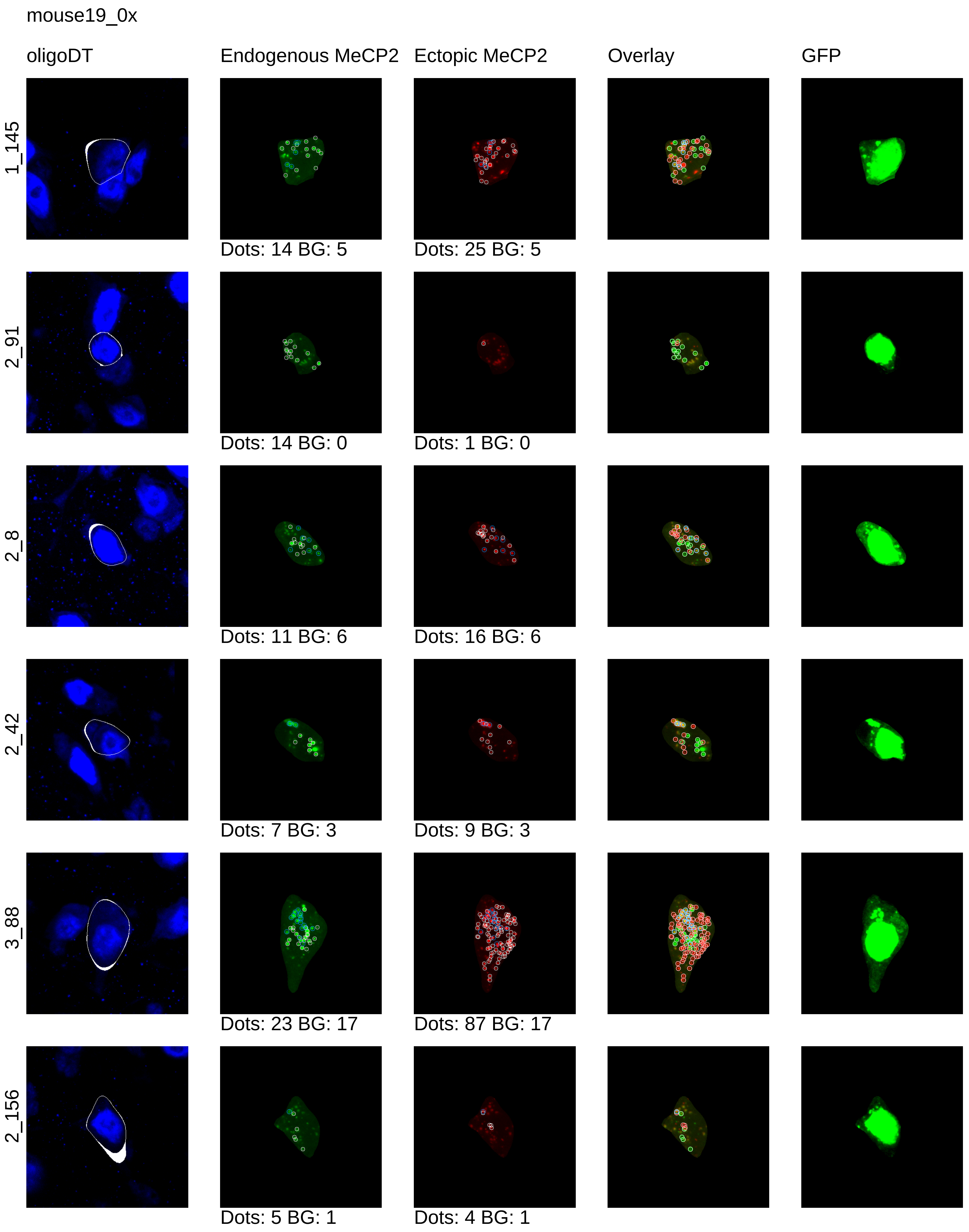
mouse19_0x
oligoDT
Endogenous MeCP2
Ectopic MeCP2
Overlay
GFP
1_145
Dots: 14 BG: 5
Dots: 25 BG: 5
2_91
Dots: 14 BG: 0
Dots: 1 BG: 0
2_8
Dots: 11 BG: 6
Dots: 16 BG: 6
2_42
Dots: 7 BG: 3
Dots: 9 BG: 3
3_88
Dots: 23 BG: 17
Dots: 87 BG: 17
2_156
Dots: 5 BG: 1
Dots: 4 BG: 1

mouse19_0x

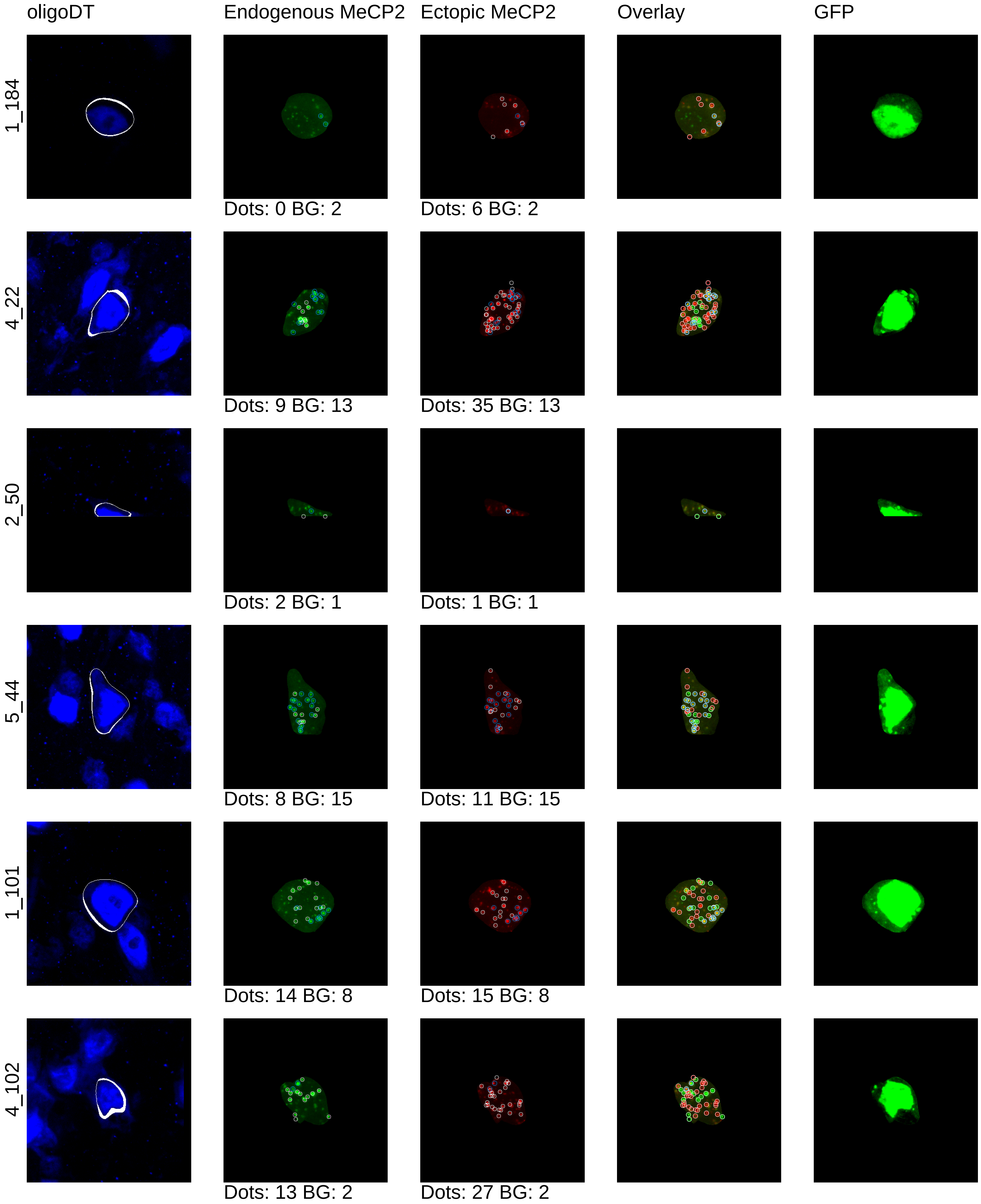
oligoDT
Endogenous MeCP2
Ectopic MeCP2
Overlay
GFP
1_184
Dots: 0 BG: 2
Dots: 6 BG: 2
4_22
Dots: 9 BG: 13
Dots: 35 BG: 13
2_50
Dots: 2 BG: 1
Dots: 1 BG: 1
5_44
Dots: 8 BG: 15
Dots: 11 BG: 15
1_101
Dots: 14 BG: 8
Dots: 15 BG: 8
4_102
Dots: 13 BG: 2
Dots: 27 BG: 2

mouse19_0x

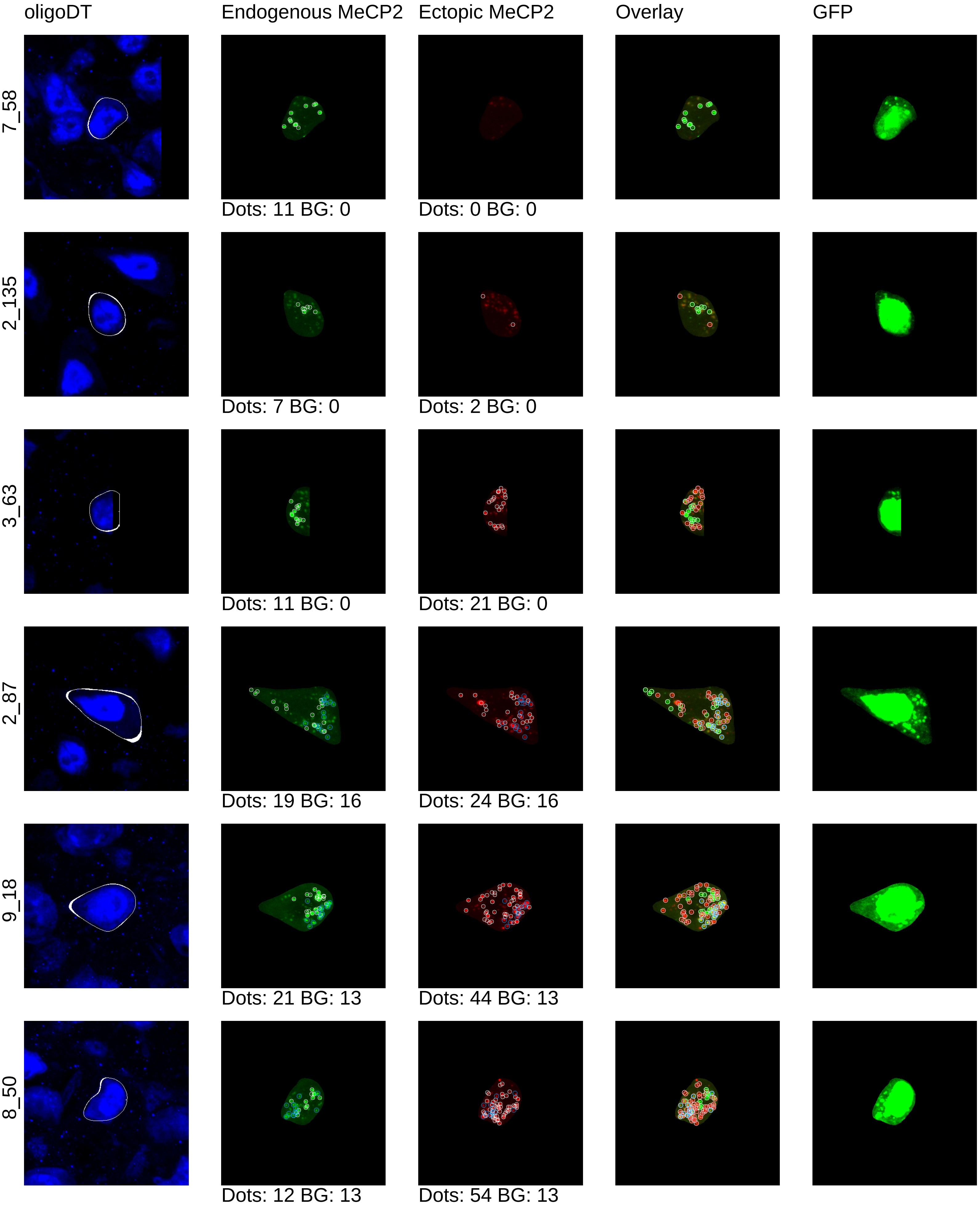
oligoDT
Endogenous MeCP2
Ectopic MeCP2
Overlay
GFP
7_58
Dots: 11 BG: 0
Dots: 0 BG: 0
2_135
Dots: 7 BG: 0
Dots: 2 BG: 0
3_63
Dots: 11 BG: 0
Dots: 21 BG: 0
2_87
Dots: 19 BG: 16
Dots: 24 BG: 16
9_18
Dots: 21 BG: 13
Dots: 44 BG: 13
8_50
Dots: 12 BG: 13
Dots: 54 BG: 13

mouse19_0x

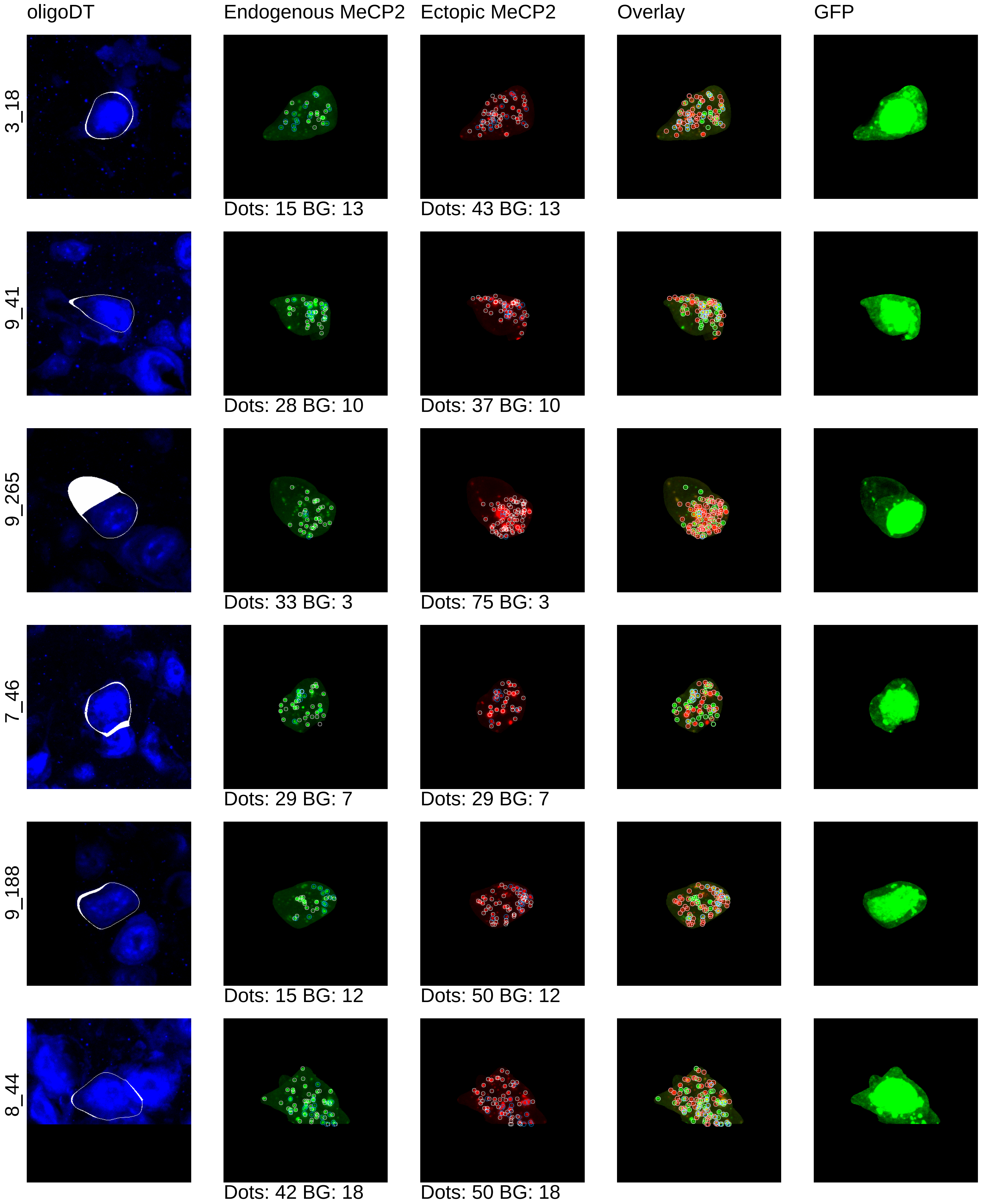
oligoDT
Endogenous MeCP2
Ectopic MeCP2
Overlay
GFP
3_18
Dots: 15 BG: 13
Dots: 43 BG: 13
9_41
Dots: 28 BG: 10
Dots: 37 BG: 10
9_265
Dots: 33 BG: 3
Dots: 75 BG: 3
7_46
Dots: 29 BG: 7
Dots: 29 BG: 7
9_188
Dots: 15 BG: 12
Dots: 50 BG: 12
8_44
Dots: 42 BG: 18
Dots: 50 BG: 18

mouse19_0x

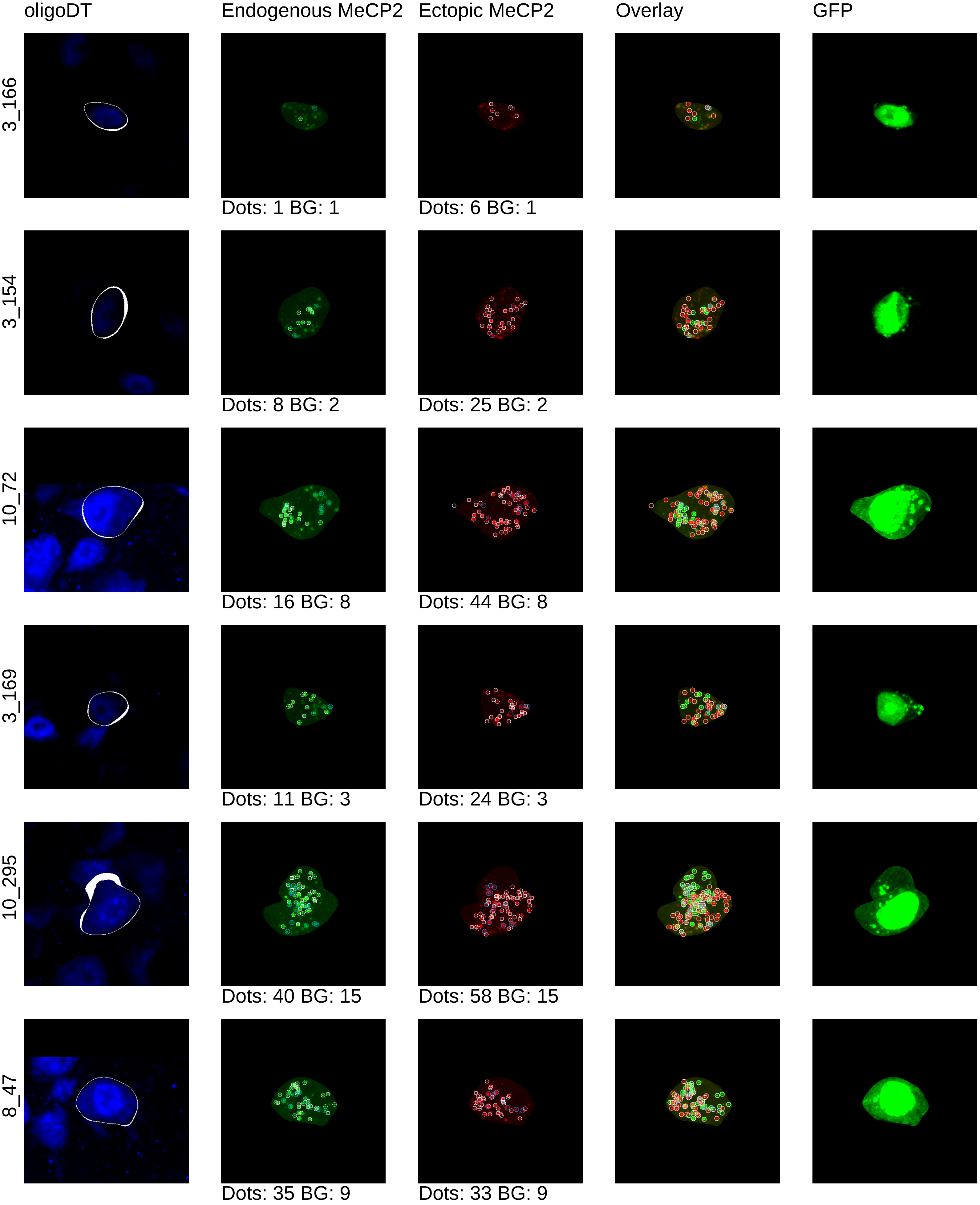
oligoDT
Endogenous MeCP2
Ectopic MeCP2
Overlay
GFP
3_166
Dots: 1 BG: 1
Dots: 6 BG: 1
3_154
Dots: 8 BG: 2
Dots: 25 BG: 2
10_72
Dots: 16 BG: 8
Dots: 44 BG: 8
3_169
Dots: 11 BG: 3
Dots: 24 BG: 3
10_295
Dots: 40 BG: 15
Dots: 58 BG: 15
8_47
Dots: 35 BG: 9
Dots: 33 BG: 9

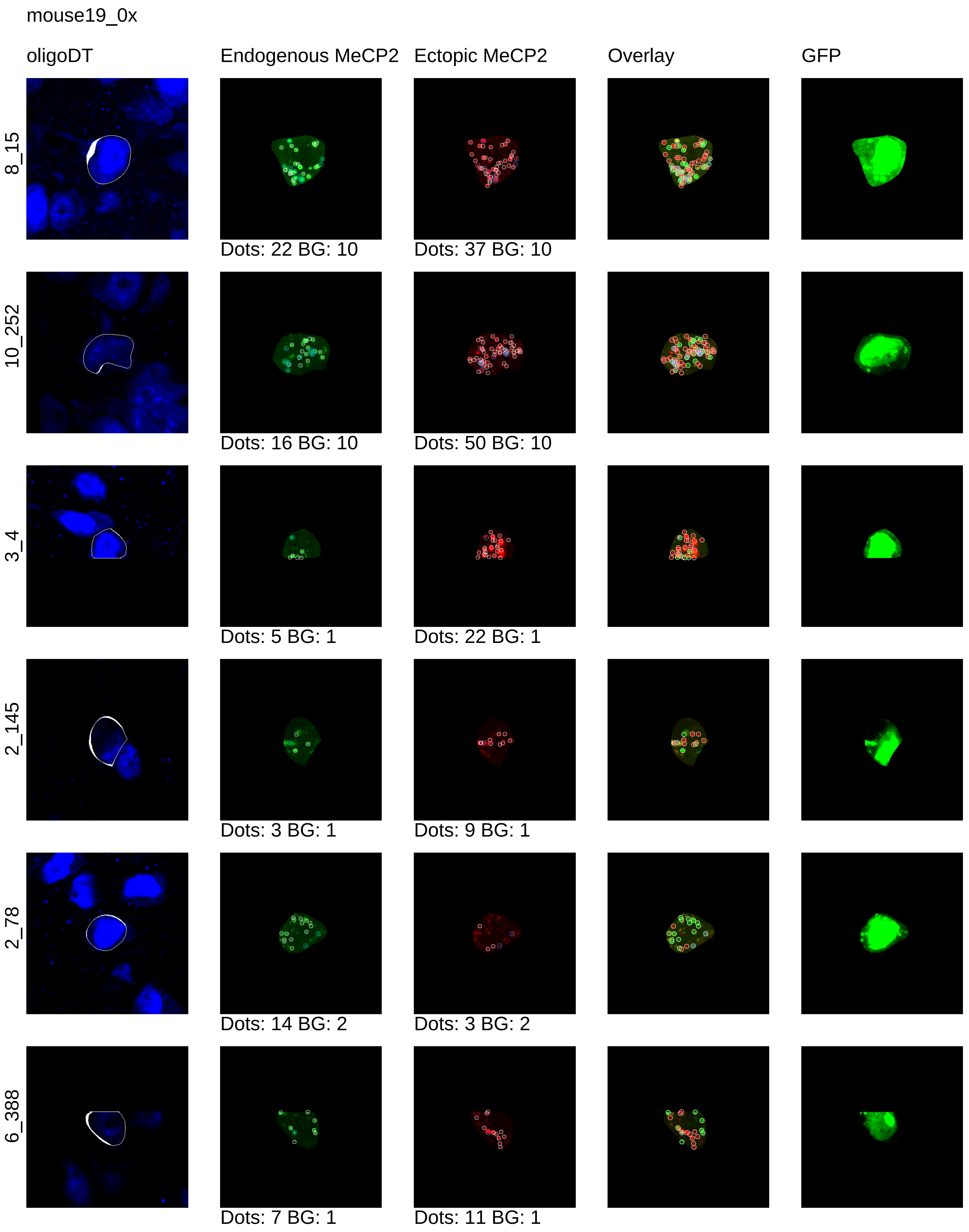
mouse19_0x
oligoDT
Endogenous MeCP2
Ectopic MeCP2
Overlay
GFP
8_15
Dots: 22 BG: 10
Dots: 37 BG: 10
10_252
Dots: 16 BG: 10
Dots: 50 BG: 10
3_4
Dots: 5 BG: 1
Dots: 22 BG: 1
2_145
Dots: 3 BG: 1
Dots: 9 BG: 1
2_78
Dots: 14 BG: 2
Dots: 3 BG: 2
6_388
Dots: 7 BG: 1
Dots: 11 BG: 1

mouse19_0x

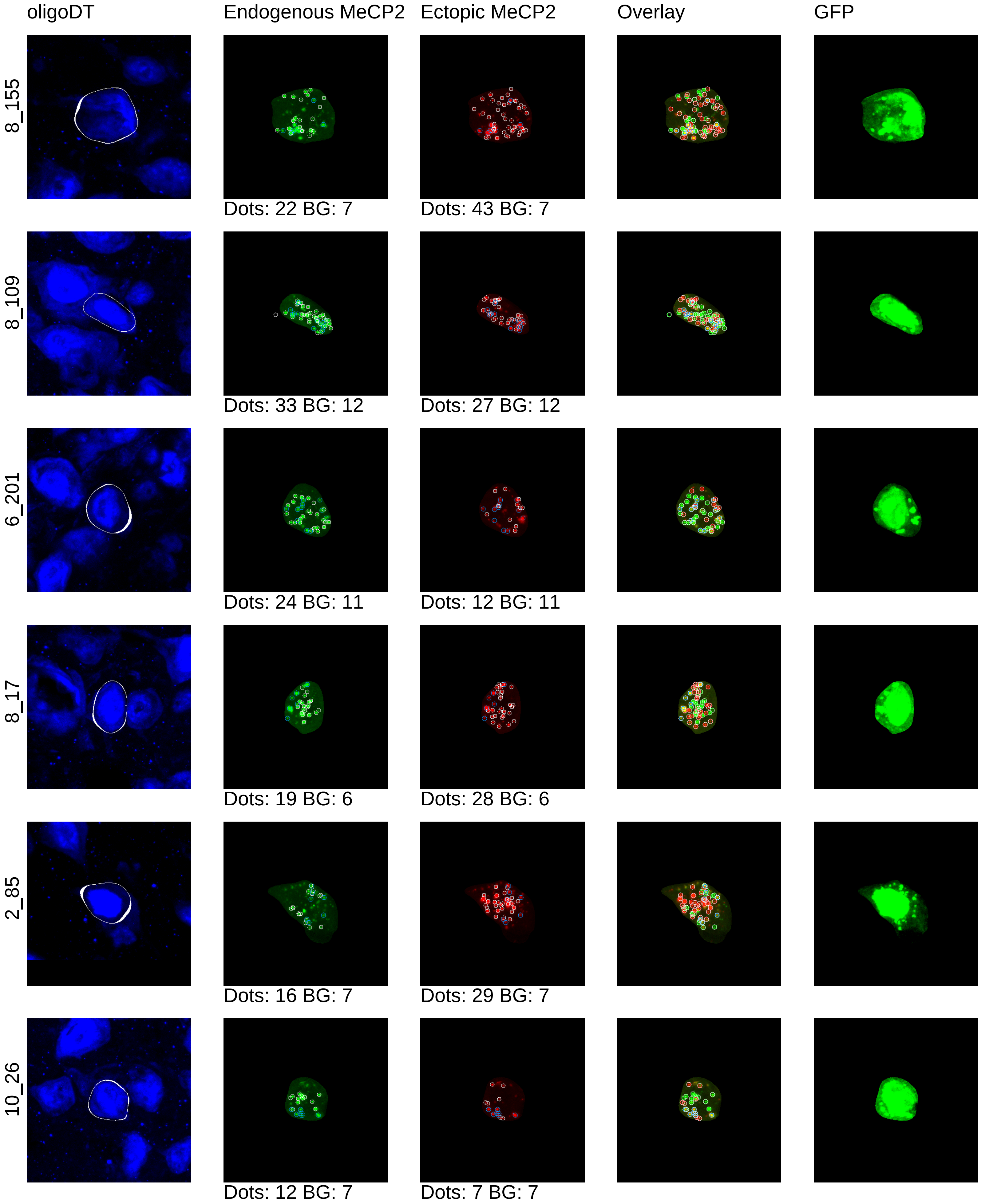

oligoDT
Endogenous MeCP2
Ectopic MeCP2
Overlay
GFP
8_155
Dots: 22 BG: 7
Dots: 43 BG: 7
8_109
Dots: 33 BG: 12
Dots: 27 BG: 12
6_201
Dots: 24 BG: 11
Dots: 12 BG: 11
8_17
Dots: 19 BG: 6
Dots: 28 BG: 6
2_85
Dots: 16 BG: 7
Dots: 29 BG: 7
10_26
Dots: 12 BG: 7
Dots: 7 BG: 7

mouse19_0x

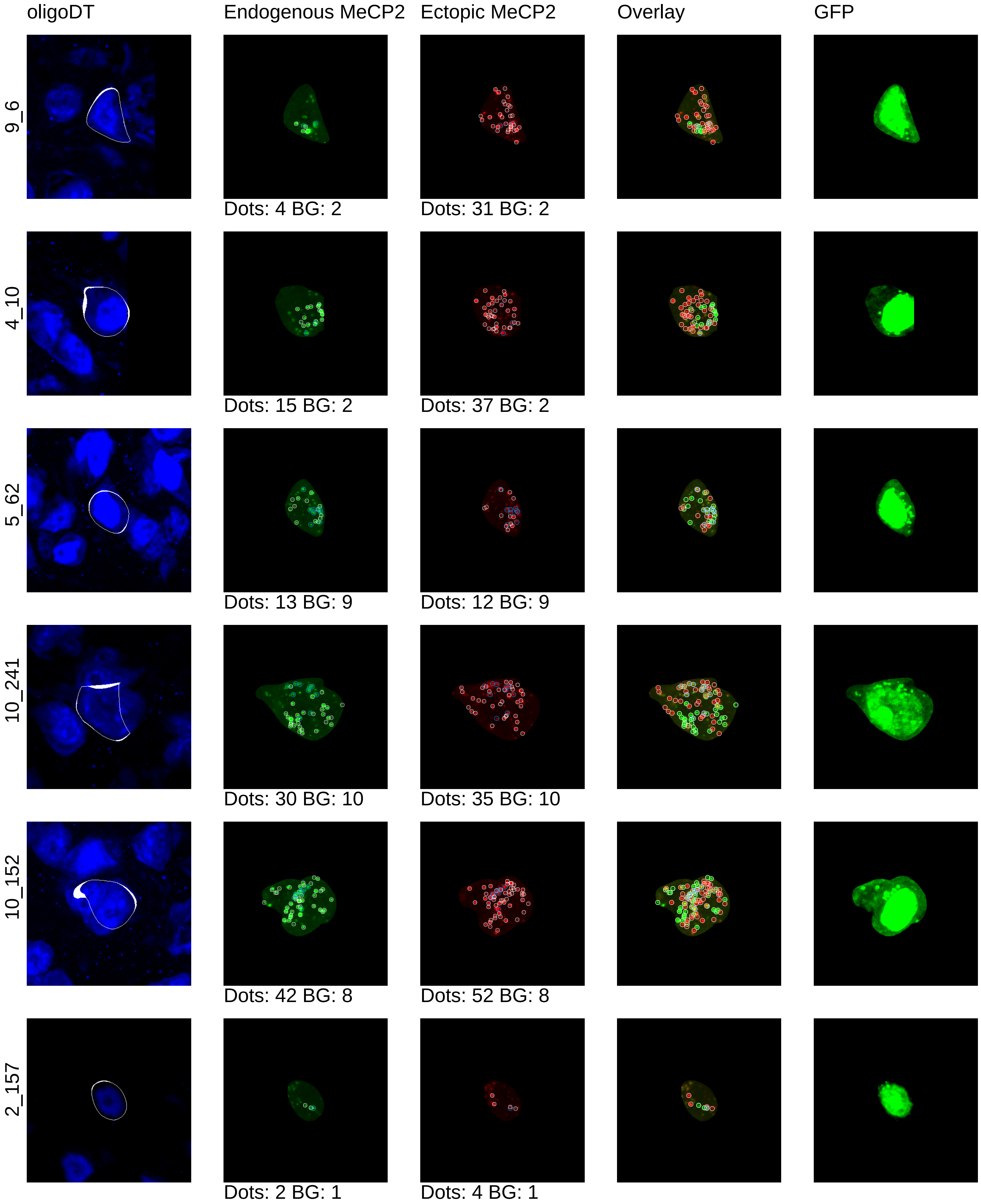

oligoDT
Endogenous MeCP2
Ectopic MeCP2
Overlay
GFP
9_6
Dots: 4 BG: 2
Dots: 31 BG: 2
4_10
Dots: 15 BG: 2
Dots: 37 BG: 2
5_62
Dots: 13 BG: 9
Dots: 12 BG: 9
10_241
Dots: 30 BG: 10
Dots: 35 BG: 10
10_152
Dots: 42 BG: 8
Dots: 52 BG: 8
2_157
Dots: 2 BG: 1
Dots: 4 BG: 1

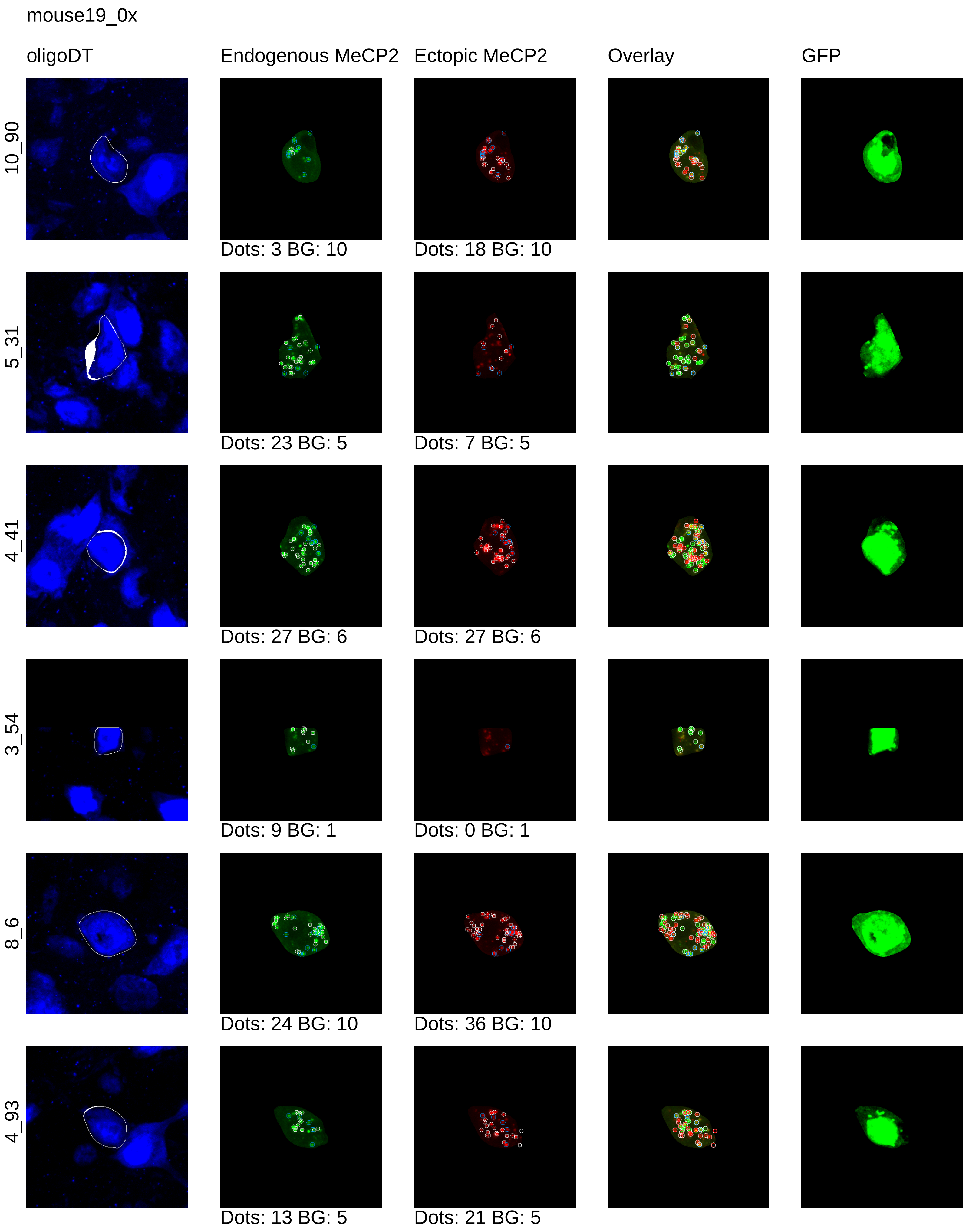
mouse19_0x
oligoDT
Endogenous MeCP2
Ectopic MeCP2
Overlay
GFP
10_90
Dots: 3 BG: 10
Dots: 18 BG: 10
5_31
Dots: 23 BG: 5
Dots: 7 BG: 5
4_41
Dots: 27 BG: 6
Dots: 27 BG: 6
3_54
Dots: 9 BG: 1
Dots: 0 BG: 1
8_6
Dots: 24 BG: 10
Dots: 36 BG: 10
4_93
Dots: 13 BG: 5
Dots: 21 BG: 5

mouse19_0x

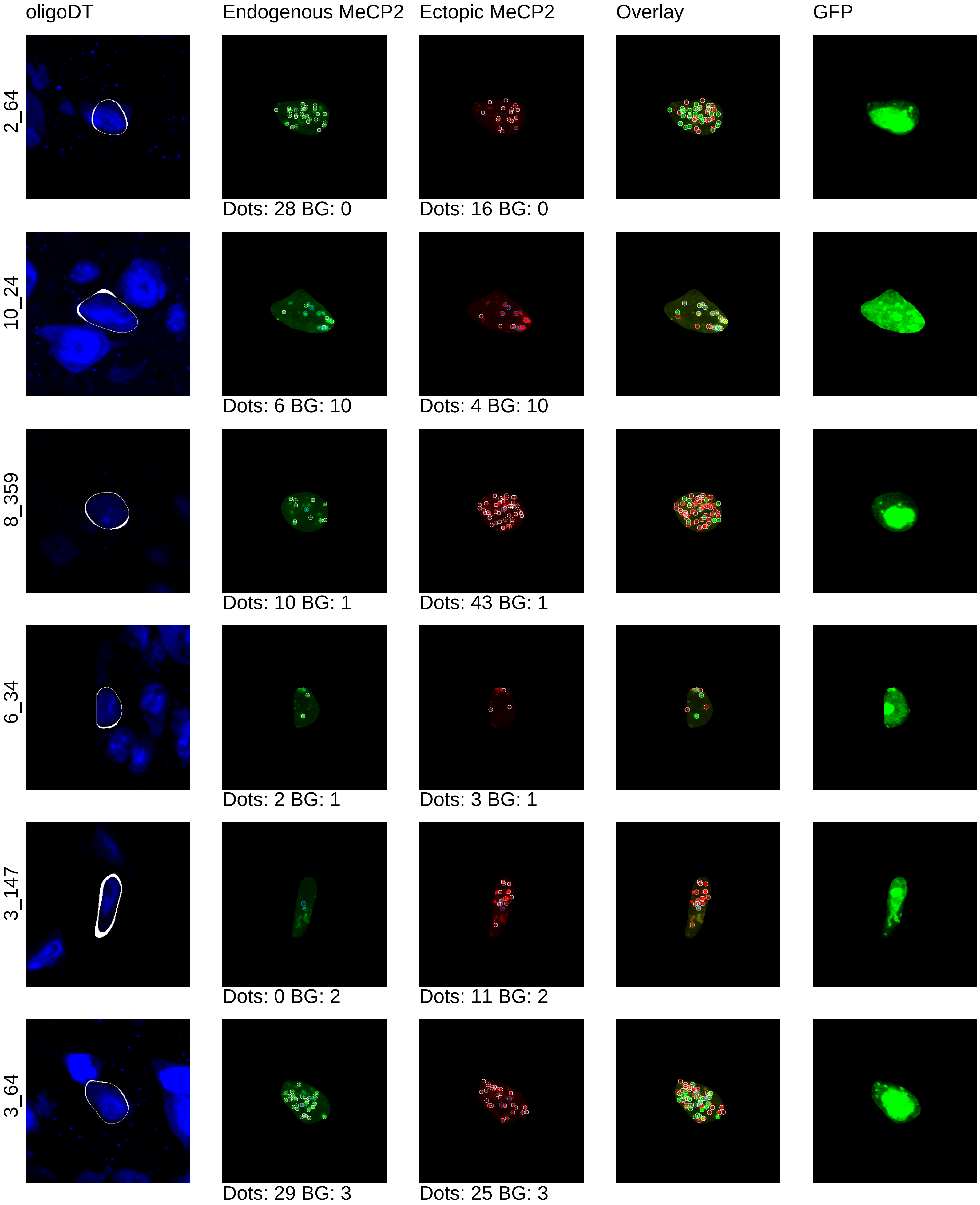
oligoDT
Endogenous MeCP2
Ectopic MeCP2
Overlay
GFP
2_64
Dots: 28 BG: 0
Dots: 16 BG: 0
10_24
Dots: 6 BG: 10
Dots: 4 BG: 10
8_359
Dots: 10 BG: 1
Dots: 43 BG: 1
6_34
Dots: 2 BG: 1
Dots: 3 BG: 1
3_147
Dots: 0 BG: 2
Dots: 11 BG: 2
3_64
Dots: 29 BG: 3
Dots: 25 BG: 3

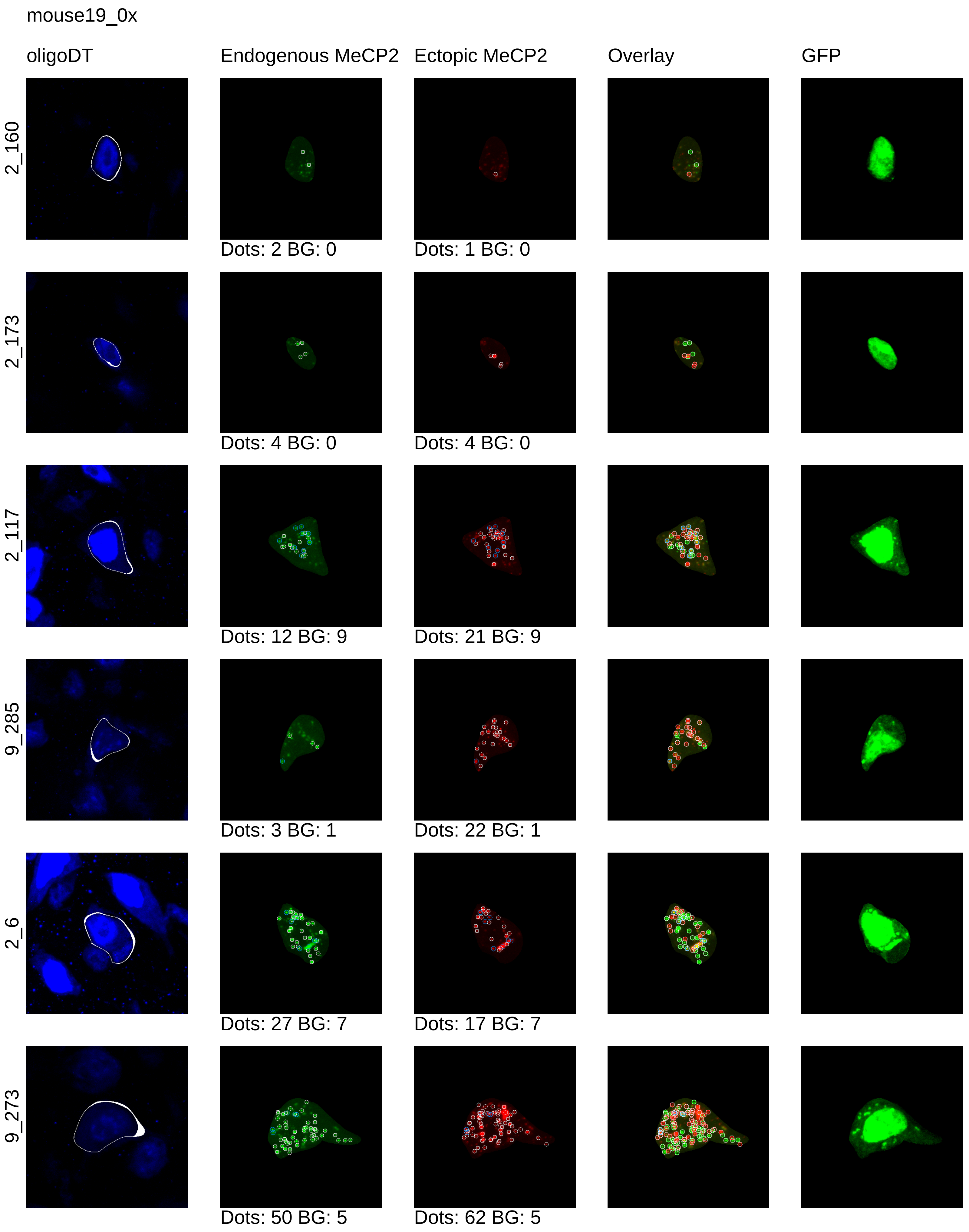
mouse19_0x
oligoDT
Endogenous MeCP2
Ectopic MeCP2
Overlay
GFP
2_160
Dots: 2 BG: 0
Dots: 1 BG: 0
2_173
Dots: 4 BG: 0
Dots: 4 BG: 0
2_117
Dots: 12 BG: 9
Dots: 21 BG: 9
9_285
Dots: 3 BG: 1
Dots: 22 BG: 1
2_6
Dots: 27 BG: 7
Dots: 17 BG: 7
9_273
Dots: 50 BG: 5
Dots: 62 BG: 5

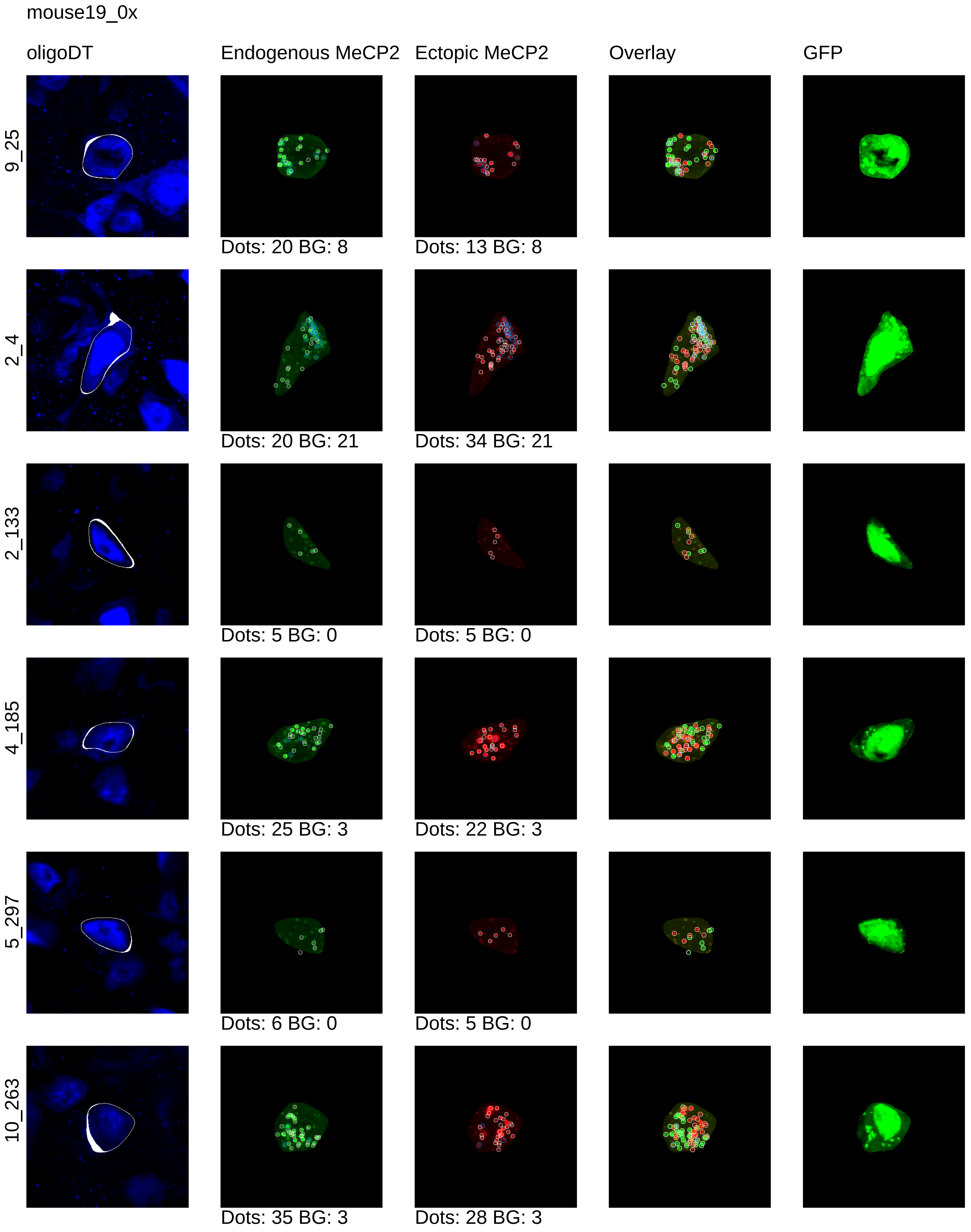
mouse19_0x
oligoDT
Endogenous MeCP2
Ectopic MeCP2
Overlay
GFP
9_25
Dots: 20 BG: 8
Dots: 13 BG: 8
2_4
Dots: 20 BG: 21
Dots: 34 BG: 21
2_133
Dots: 5 BG: 0
Dots: 5 BG: 0
4_185
Dots: 25 BG: 3
Dots: 22 BG: 3
5_297
Dots: 6 BG: 0
Dots: 5 BG: 0
10_263
Dots: 35 BG: 3
Dots: 28 BG: 3

mouse19_0x

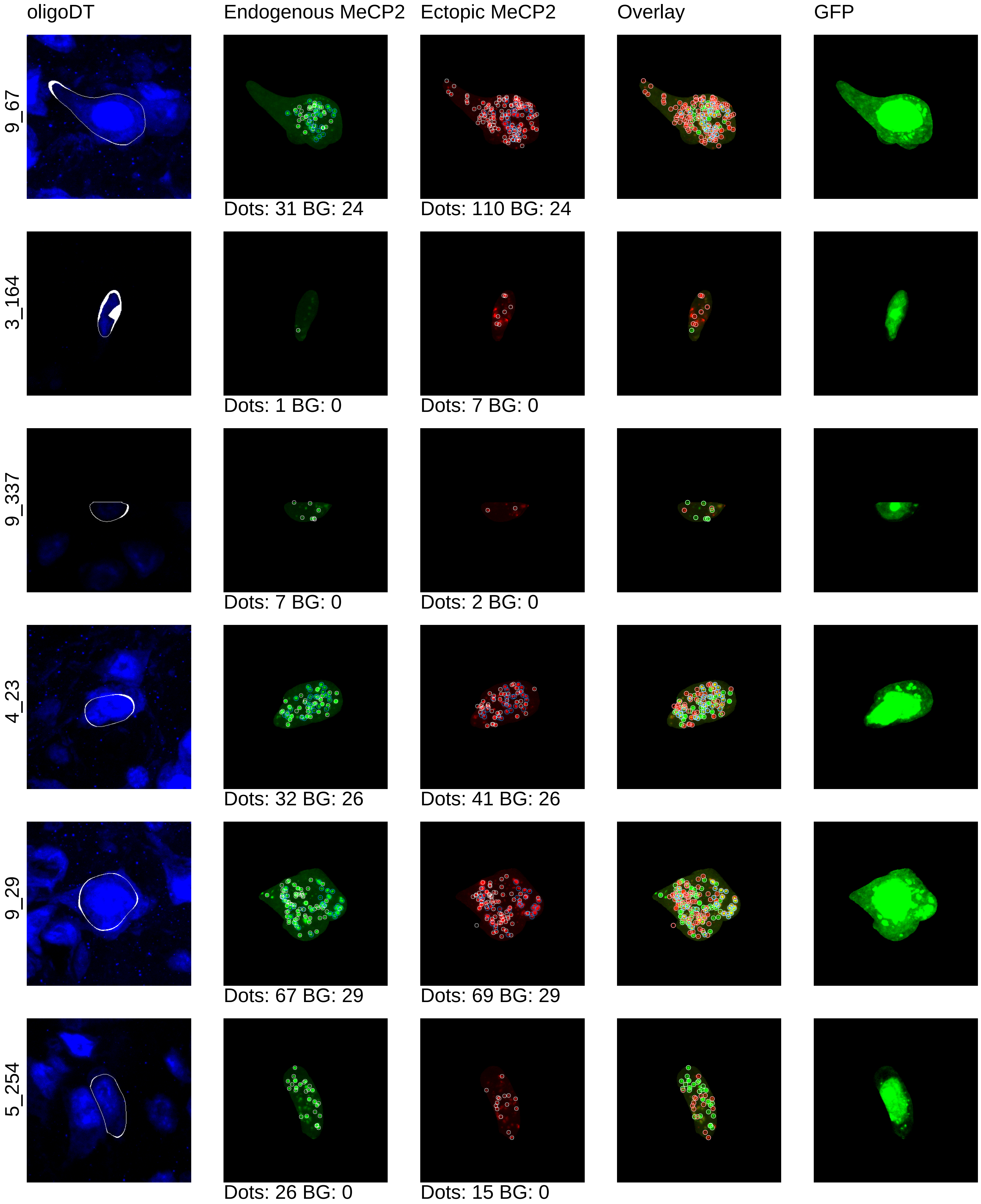
oligoDT
Endogenous MeCP2
Ectopic MeCP2
Overlay
GFP
9_67
Dots: 31 BG: 24
Dots: 110 BG: 24
3_164
Dots: 1 BG: 0
Dots: 7 BG: 0
9_337
Dots: 7 BG: 0
Dots: 2 BG: 0
4_23
Dots: 32 BG: 26
Dots: 41 BG: 26
9_29
Dots: 67 BG: 29
Dots: 69 BG: 29
5_254
Dots: 26 BG: 0
Dots: 15 BG: 0

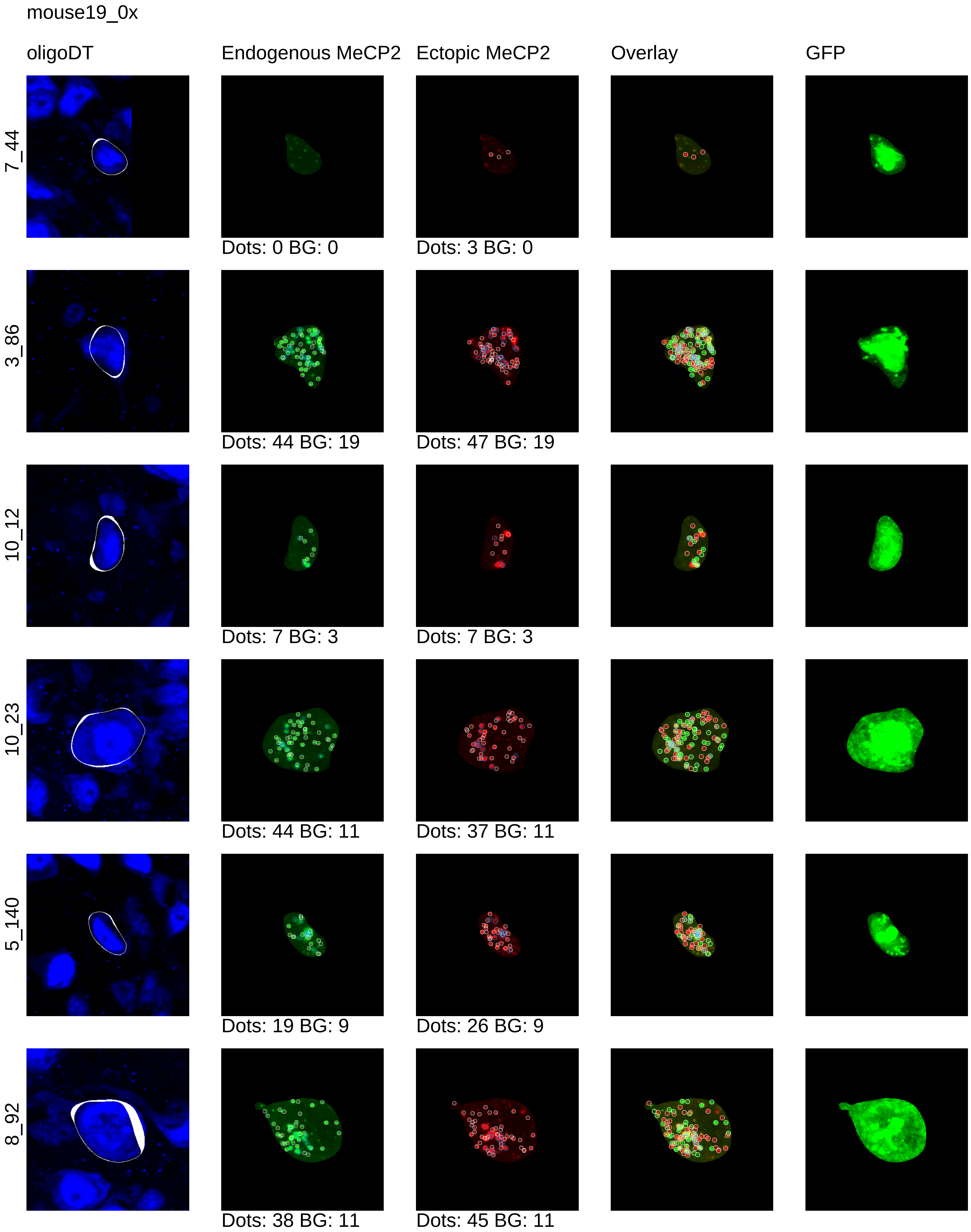
mouse19_0x
oligoDT
Endogenous MeCP2
Ectopic MeCP2
Overlay
GFP
7_44
Dots: 0 BG: 0
Dots: 3 BG: 0
3_86
Dots: 44 BG: 19
Dots: 47 BG: 19
10_12
Dots: 7 BG: 3
Dots: 7 BG: 3
10_23
Dots: 44 BG: 11
Dots: 37 BG: 11
5_140
Dots: 19 BG: 9
Dots: 26 BG: 9
8_92
Dots: 38 BG: 11
Dots: 45 BG: 11

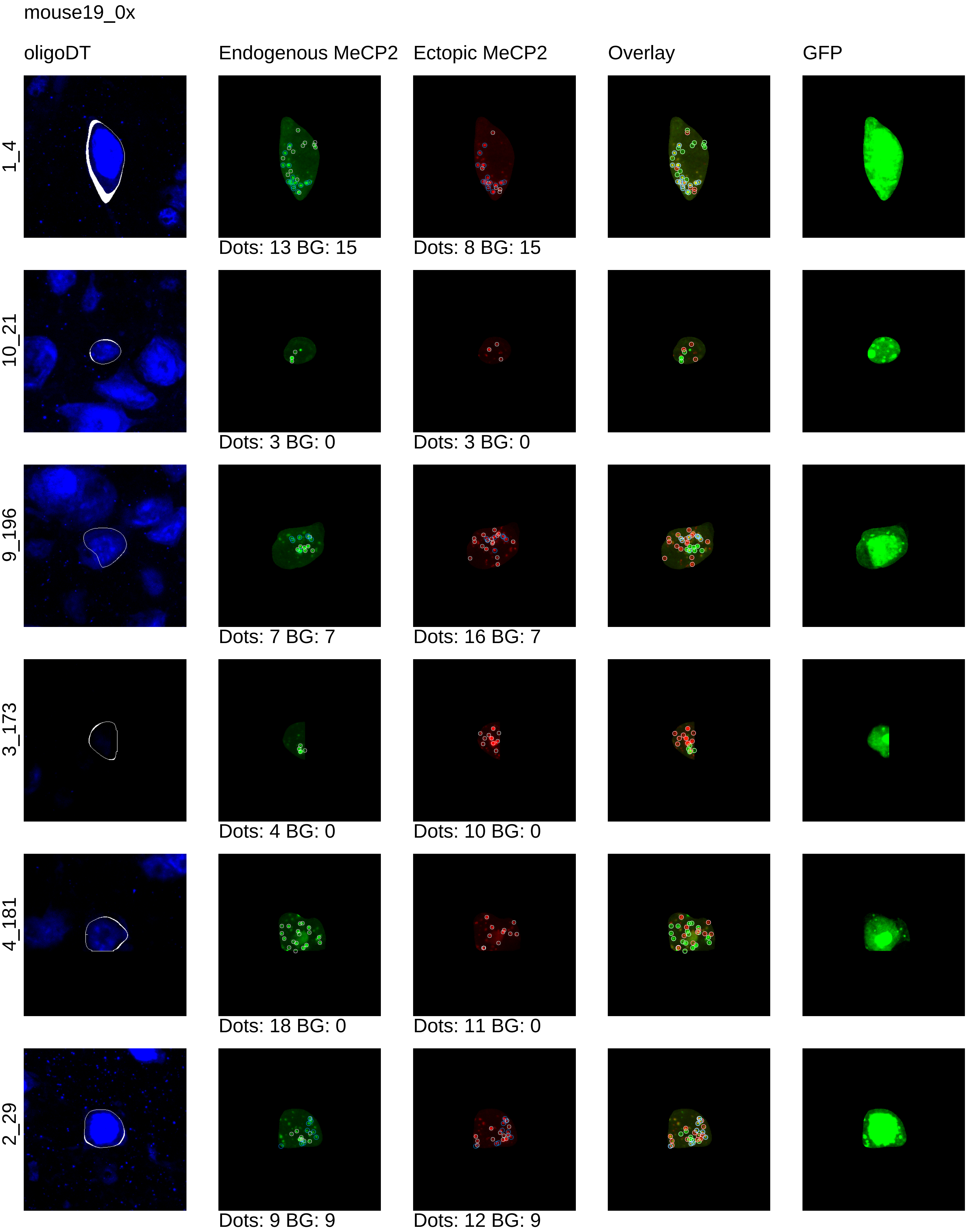
mouse19_0x
oligoDT
Endogenous MeCP2
Ectopic MeCP2
Overlay
GFP
1_4
Dots: 13 BG: 15
Dots: 8 BG: 15
10_21
Dots: 3 BG: 0
Dots: 3 BG: 0
9_196
Dots: 7 BG: 7
Dots: 16 BG: 7
3_173
Dots: 4 BG: 0
Dots: 10 BG: 0
4_181
Dots: 18 BG: 0
Dots: 11 BG: 0
2_29
Dots: 9 BG: 9
Dots: 12 BG: 9

mouse19_0x

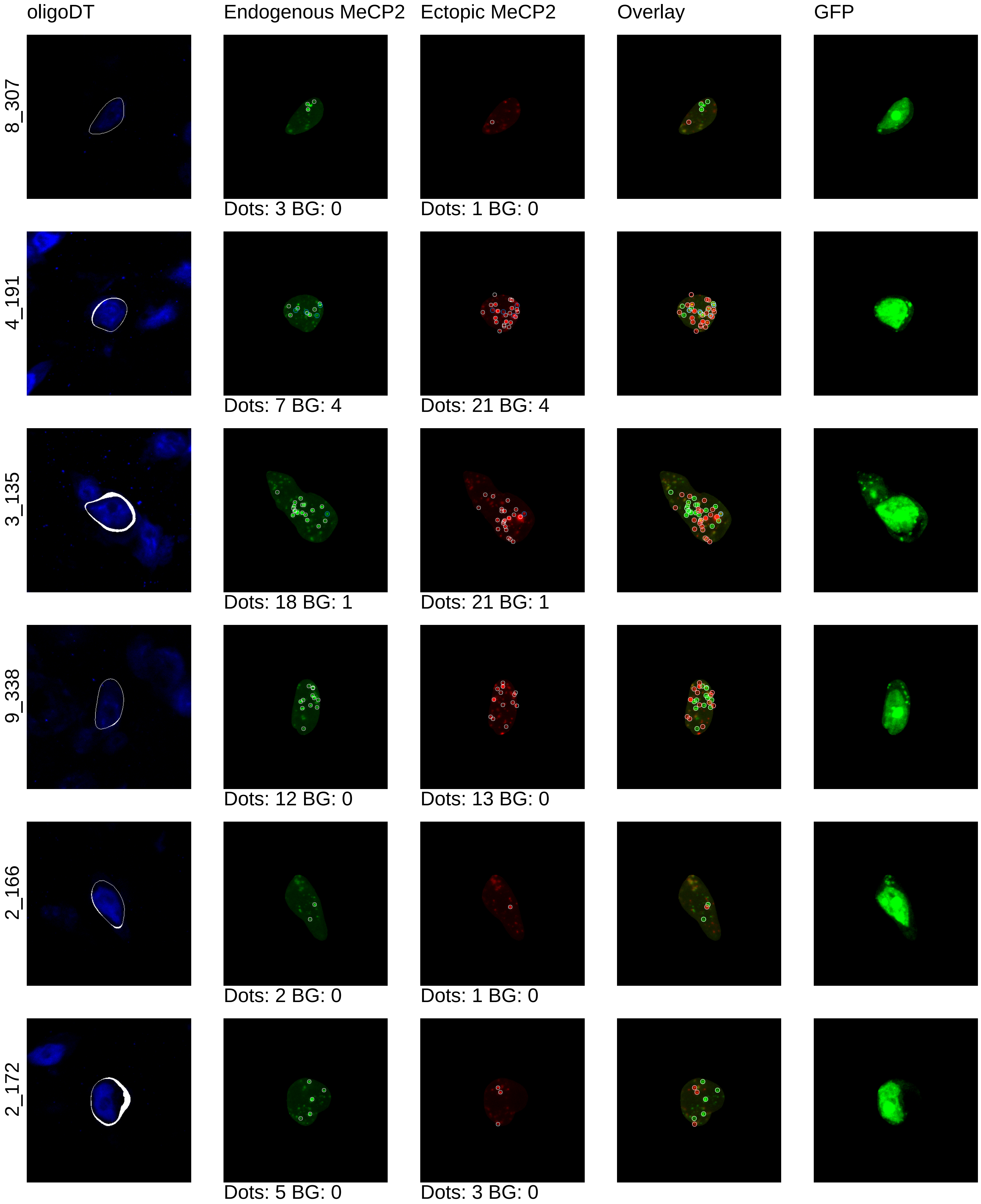

oligoDT
Endogenous MeCP2
Ectopic MeCP2
Overlay
GFP
8_307
Dots: 3 BG: 0
Dots: 1 BG: 0
4_191
Dots: 7 BG: 4
Dots: 21 BG: 4
3_135
Dots: 18 BG: 1
Dots: 21 BG: 1
9_338
Dots: 12 BG: 0
Dots: 13 BG: 0
2_166
Dots: 2 BG: 0
Dots: 1 BG: 0
2_172
Dots: 5 BG: 0
Dots: 3 BG: 0

mouse19_0x

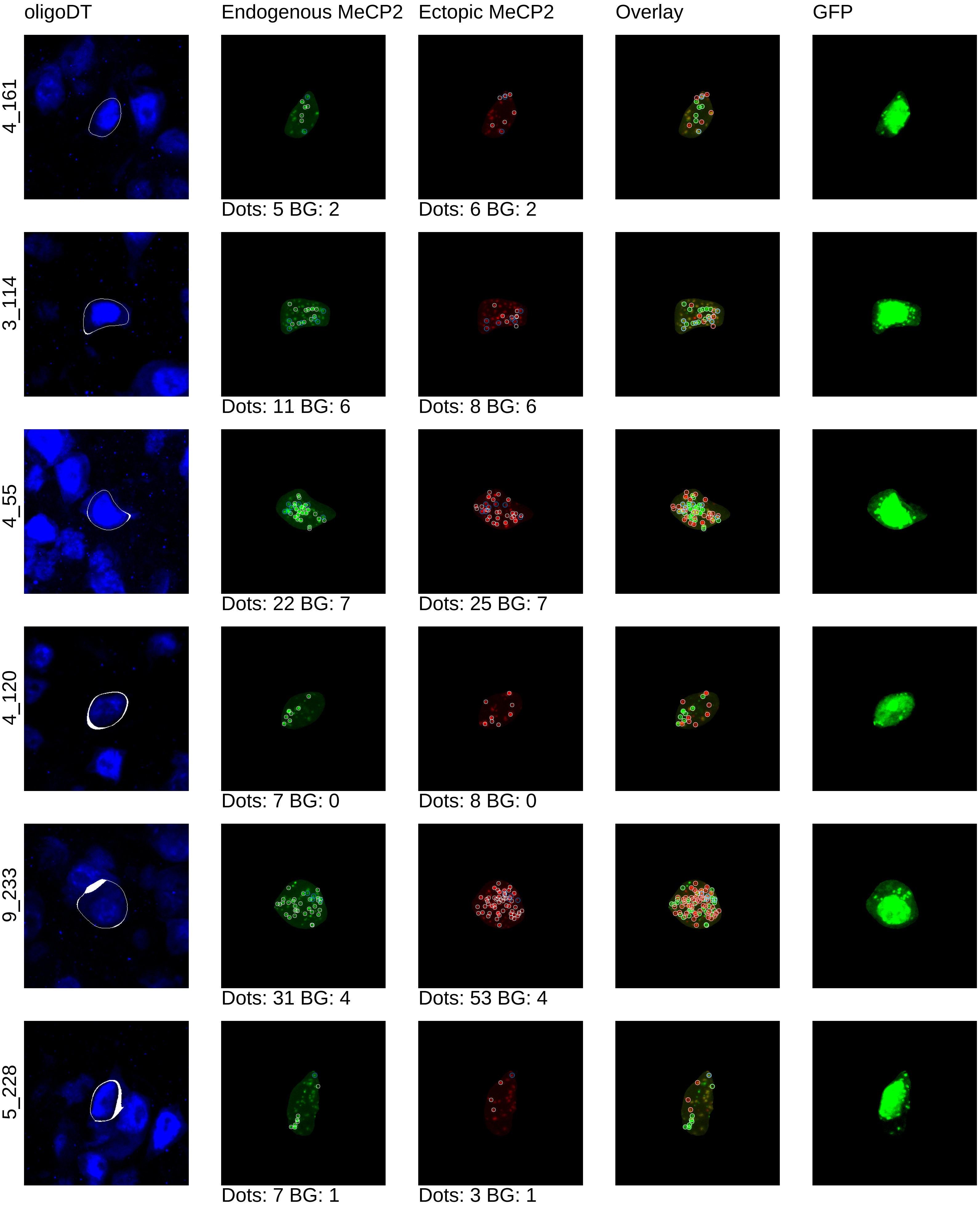
oligoDT
Endogenous MeCP2
Ectopic MeCP2
Overlay
GFP
4_161
Dots: 5 BG: 2
Dots: 6 BG: 2
3_114
Dots: 11 BG: 6
Dots: 8 BG: 6
4_55
Dots: 22 BG: 7
Dots: 25 BG: 7
4_120
Dots: 7 BG: 0
Dots: 8 BG: 0
9_233
Dots: 31 BG: 4
Dots: 53 BG: 4
5_228
Dots: 7 BG: 1
Dots: 3 BG: 1

mouse19_0x

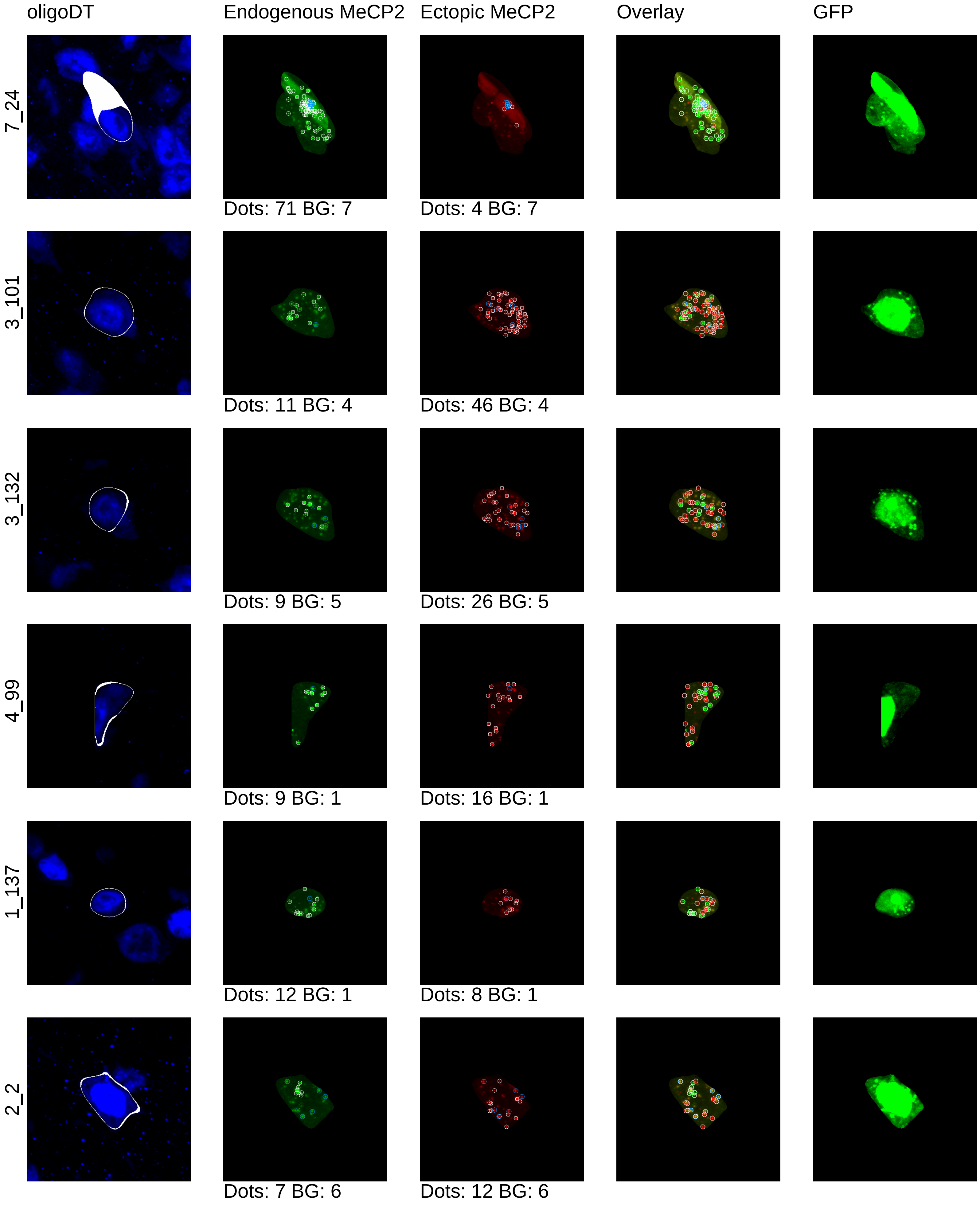
oligoDT
Endogenous MeCP2
Ectopic MeCP2
Overlay
GFP
7_24
Dots: 71 BG: 7
Dots: 4 BG: 7
3_101
Dots: 11 BG: 4
Dots: 46 BG: 4
3_132
Dots: 9 BG: 5
Dots: 26 BG: 5
4_99
Dots: 9 BG: 1
Dots: 16 BG: 1
1_137
Dots: 12 BG: 1
Dots: 8 BG: 1
2_2
Dots: 7 BG: 6
Dots: 12 BG: 6

mouse19_0x

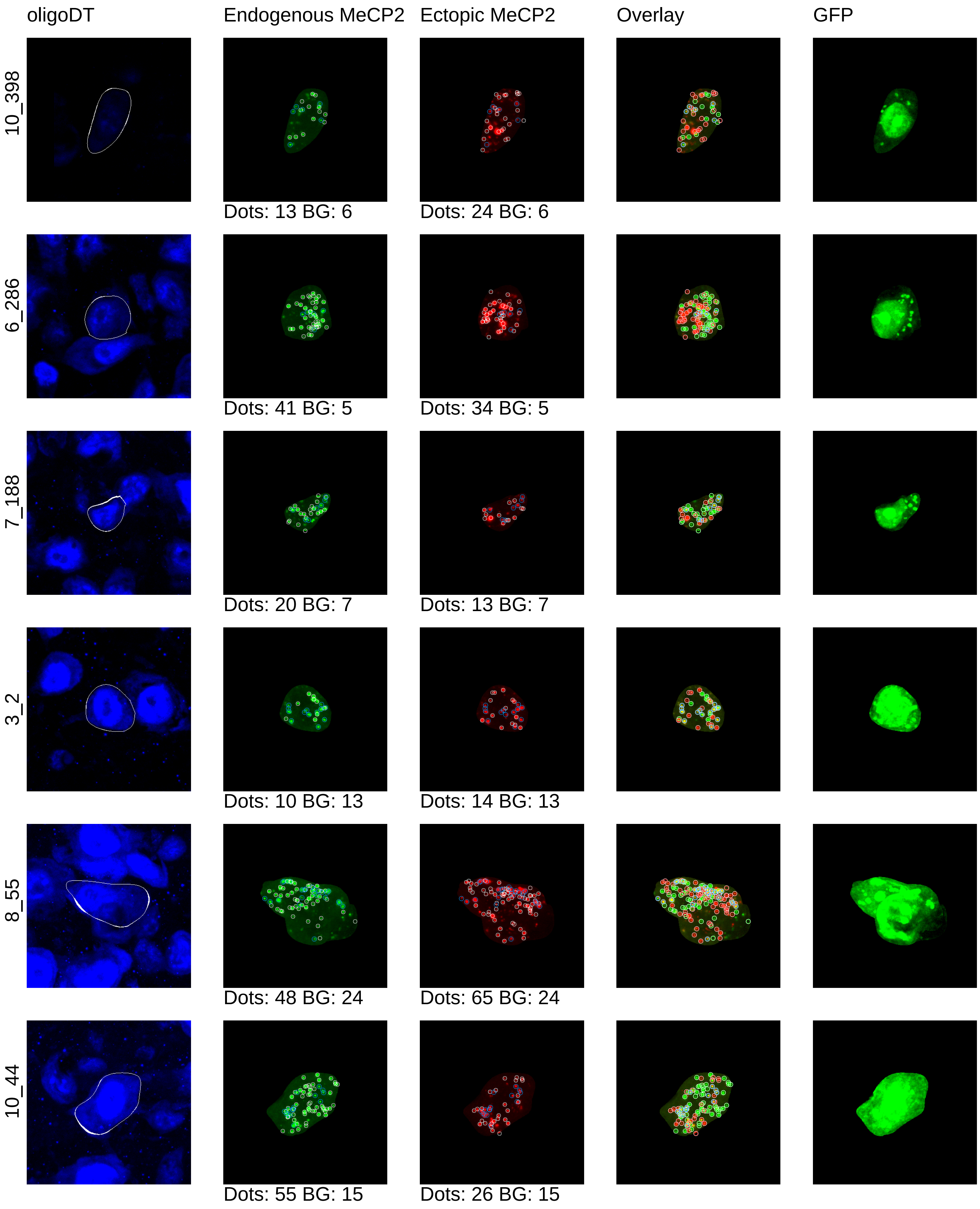
oligoDT
Endogenous MeCP2
Ectopic MeCP2
Overlay
GFP
10_398
Dots: 13 BG: 6
Dots: 24 BG: 6
6_286
Dots: 41 BG: 5
Dots: 34 BG: 5
7_188
Dots: 20 BG: 7
Dots: 13 BG: 7
3_2
Dots: 10 BG: 13
Dots: 14 BG: 13
8_55
Dots: 48 BG: 24
Dots: 65 BG: 24
10_44
Dots: 55 BG: 15
Dots: 26 BG: 15

mouse19_0x

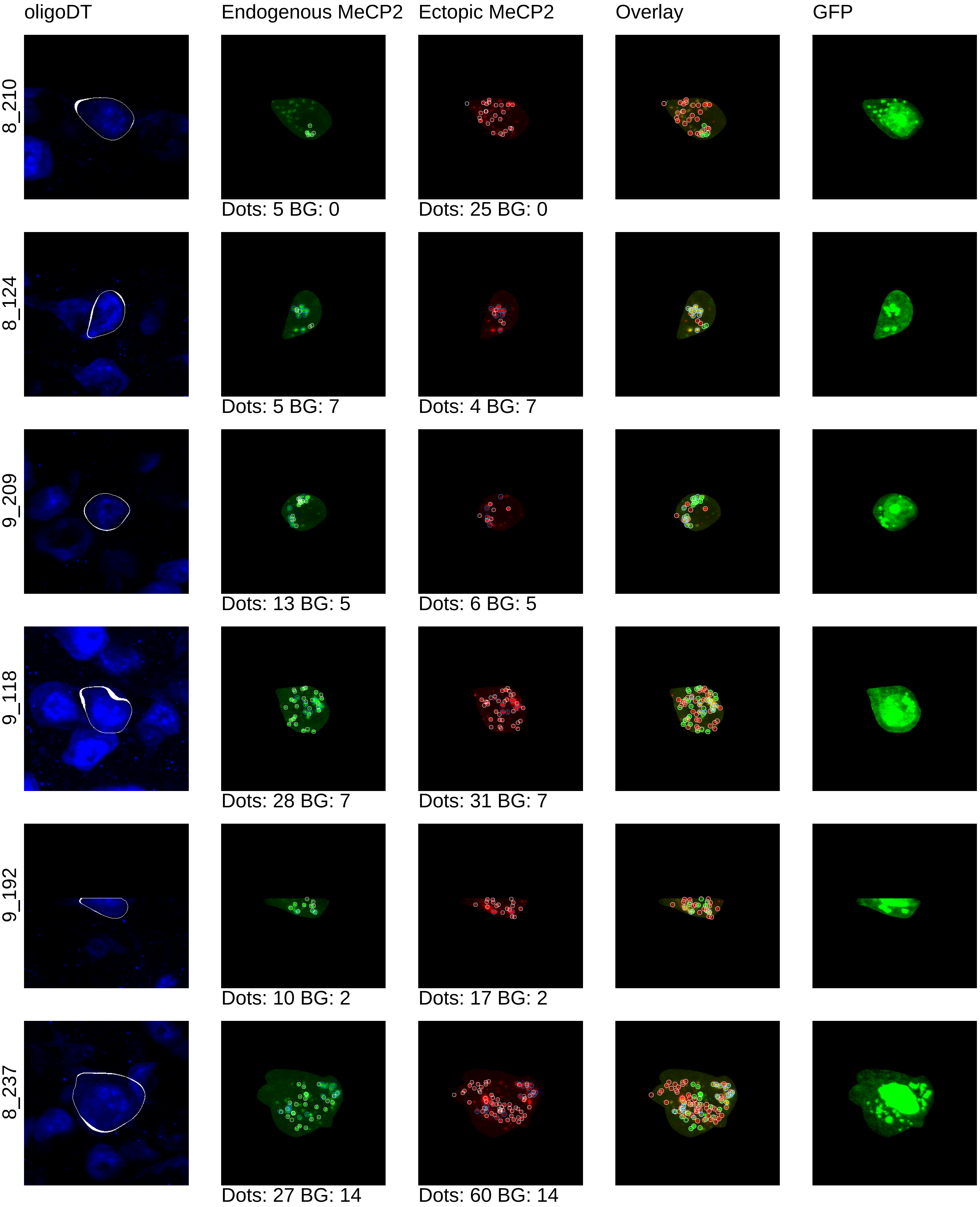
oligoDT
Endogenous MeCP2
Ectopic MeCP2
Overlay
GFP
8_210
Dots: 5 BG: 0
Dots: 25 BG: 0
8_124
Dots: 5 BG: 7
Dots: 4 BG: 7
9_209
Dots: 13 BG: 5
Dots: 6 BG: 5
9_118
Dots: 28 BG: 7
Dots: 31 BG: 7
9_192
Dots: 10 BG: 2
Dots: 17 BG: 2
8_237
Dots: 27 BG: 14
Dots: 60 BG: 14

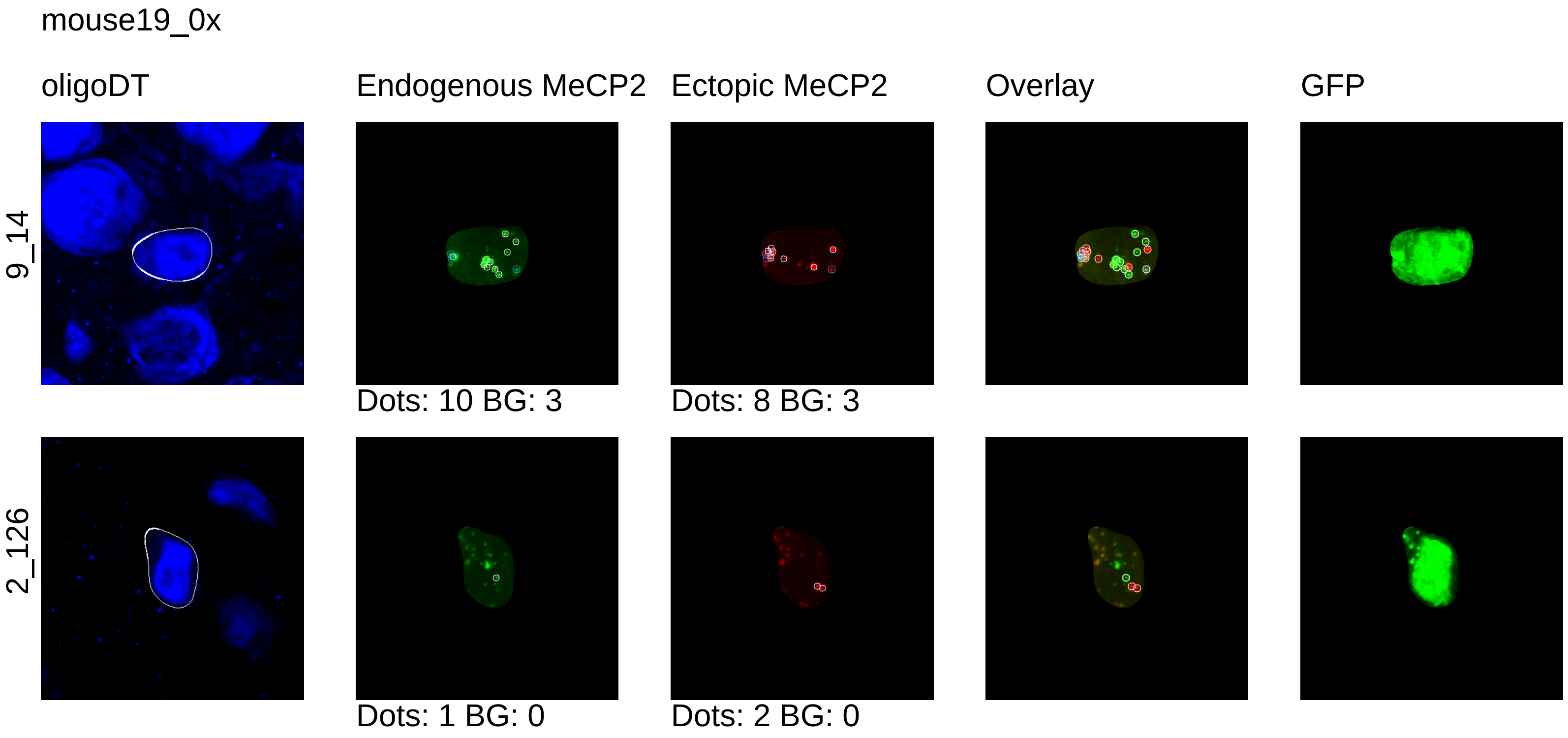
mouse19_0x
oligoDT
Endogenous MeCP2
Ectopic MeCP2
Overlay
GFP
9_14
Dots: 10 BG: 3
Dots: 8 BG: 3
2_126
Dots: 1 BG: 0
Dots: 2 BG: 0
