## Supplementary material for "Synthetic dosage-compensating miRNA circuits allow precision gene therapy for Rett syndrome": 1x_cell_segmentation

mouse12_1x

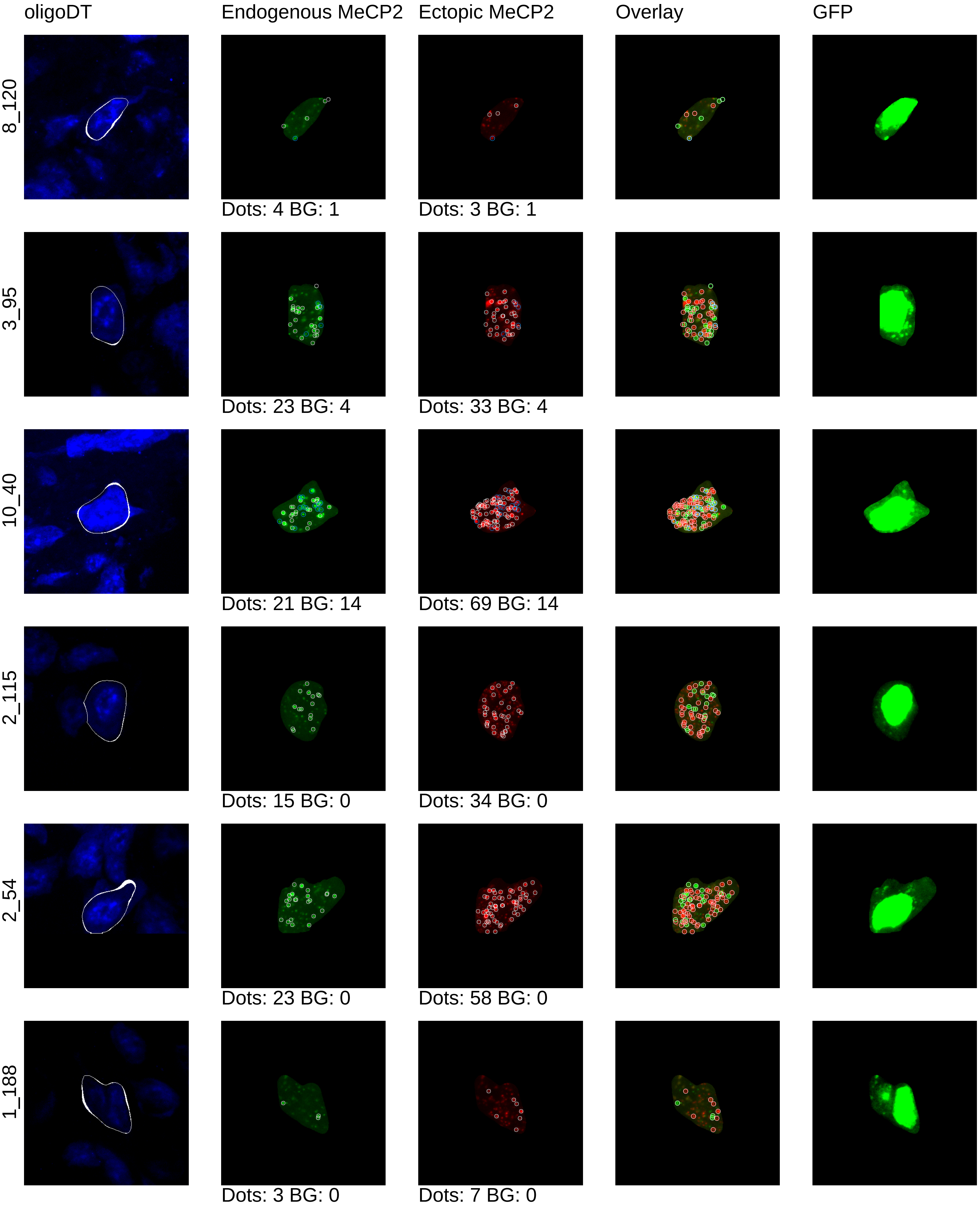
oligoDT
Endogenous MeCP2
Ectopic MeCP2
Overlay
GFP
8_120
Dots: 4 BG: 1
Dots: 3 BG: 1
3_95
Dots: 23 BG: 4
Dots: 33 BG: 4
10_40
Dots: 21 BG: 14
Dots: 69 BG: 14
2_115
Dots: 15 BG: 0
Dots: 34 BG: 0
2_54
Dots: 23 BG: 0
Dots: 58 BG: 0
1_188
Dots: 3 BG: 0
Dots: 7 BG: 0

mouse12_1x

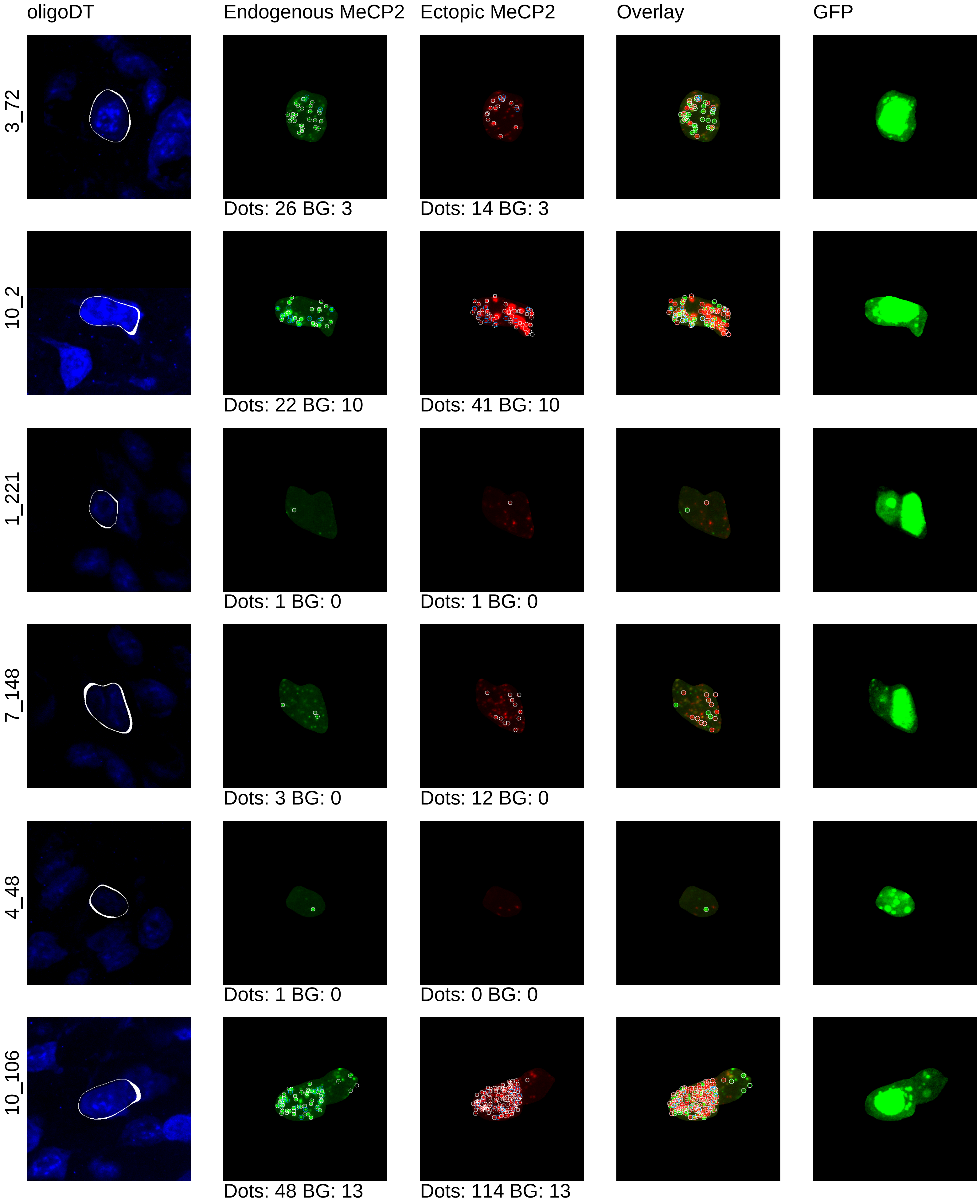
oligoDT
Endogenous MeCP2
Ectopic MeCP2
Overlay
GFP
3_72
Dots: 26 BG: 3
Dots: 14 BG: 3
10_2
Dots: 22 BG: 10
Dots: 41 BG: 10
1_221
Dots: 1 BG: 0
Dots: 1 BG: 0
7_148
Dots: 3 BG: 0
Dots: 12 BG: 0
4_48
Dots: 1 BG: 0
Dots: 0 BG: 0
10_106
Dots: 48 BG: 13
Dots: 114 BG: 13

mouse12_1x

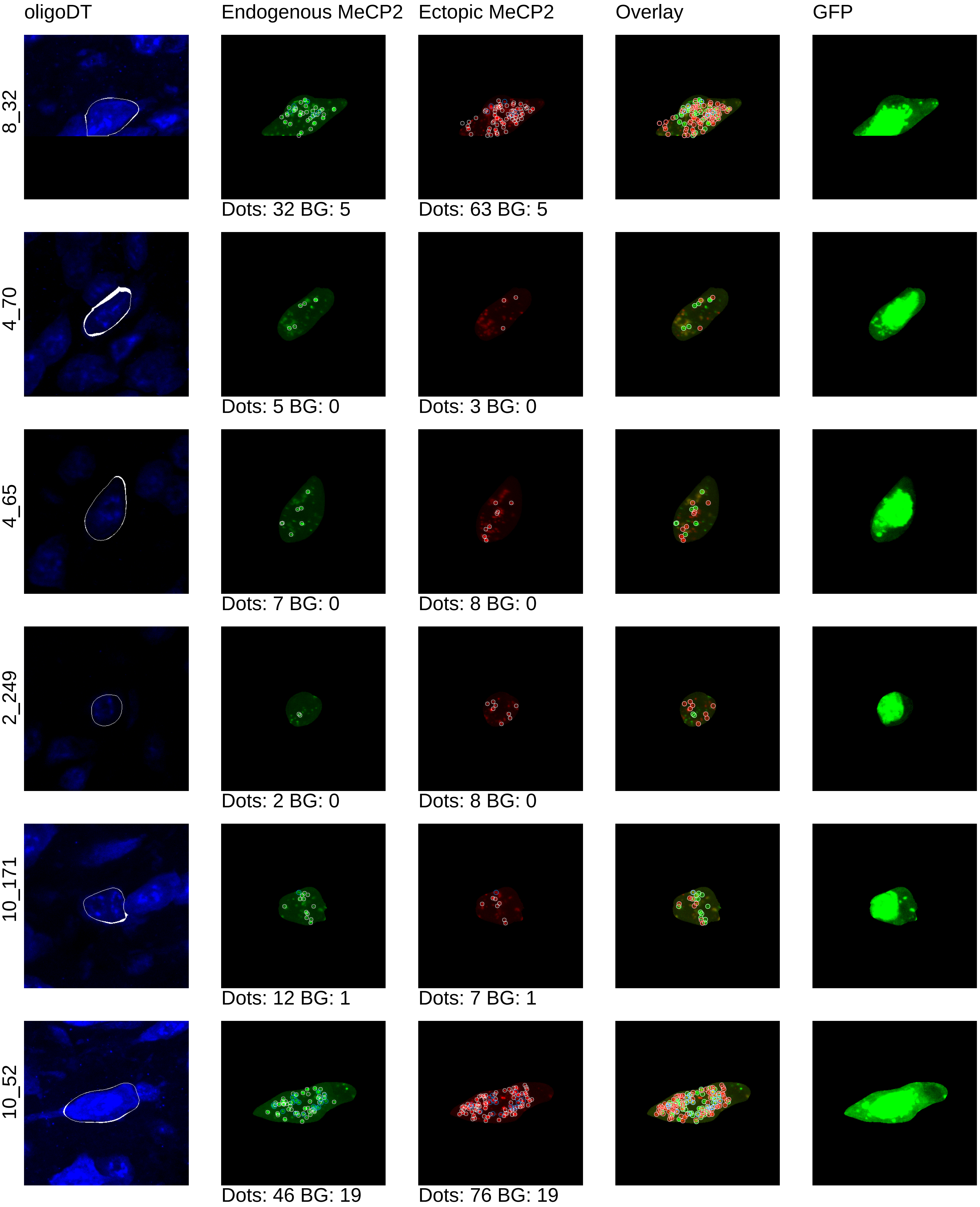

oligoDT
Endogenous MeCP2
Ectopic MeCP2
Overlay
GFP
8_32
Dots: 32 BG: 5
Dots: 63 BG: 5
4_70
Dots: 5 BG: 0
Dots: 3 BG: 0
4_65
Dots: 7 BG: 0
Dots: 8 BG: 0
2_249
Dots: 2 BG: 0
Dots: 8 BG: 0
10_171
Dots: 12 BG: 1
Dots: 7 BG: 1
10_52
Dots: 46 BG: 19
Dots: 76 BG: 19

mouse12_1x

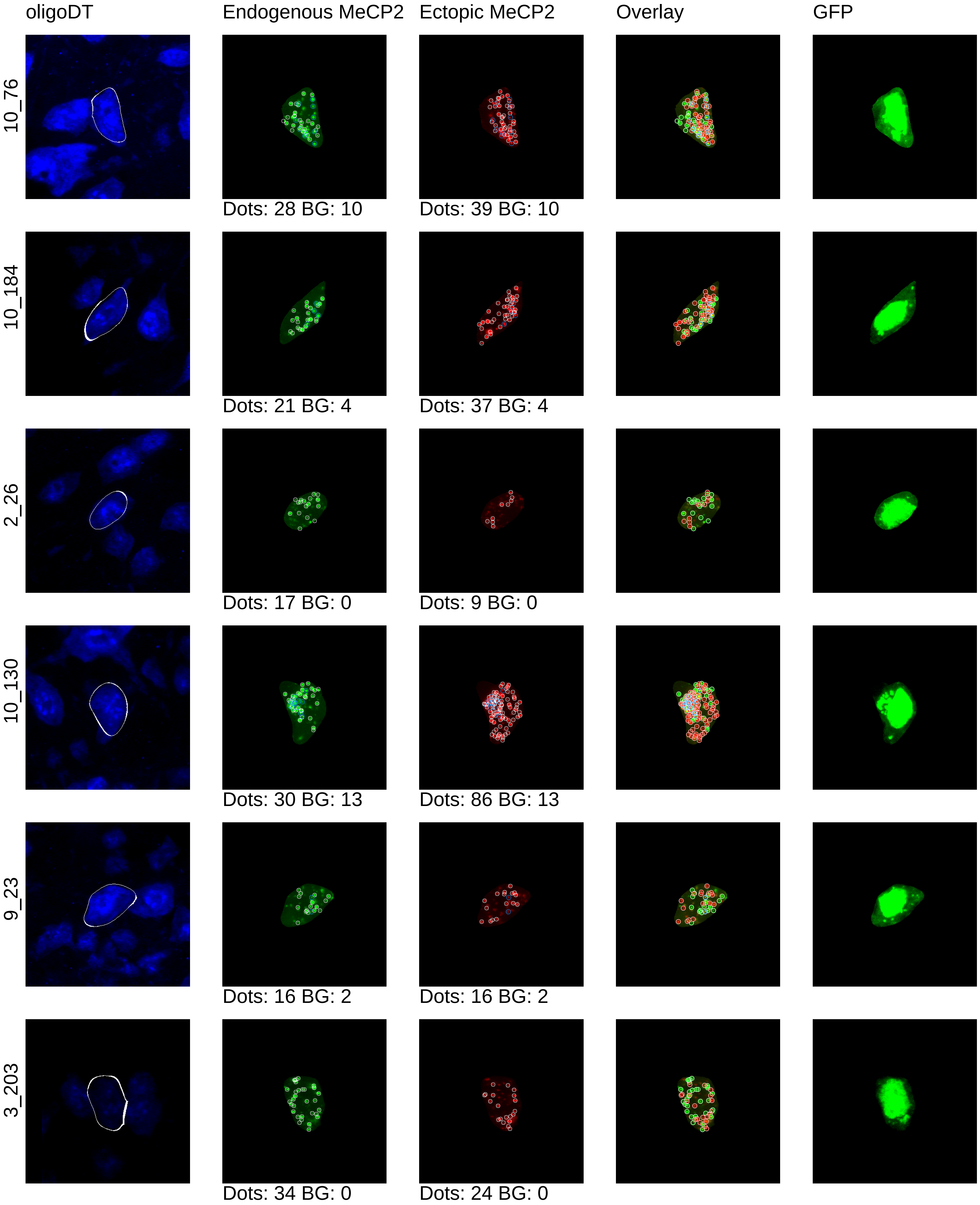
oligoDT
Endogenous MeCP2
Ectopic MeCP2
Overlay
GFP
10_76
Dots: 28 BG: 10
Dots: 39 BG: 10
10_184
Dots: 21 BG: 4
Dots: 37 BG: 4
2_26
Dots: 17 BG: 0
Dots: 9 BG: 0
10_130
Dots: 30 BG: 13
Dots: 86 BG: 13
9_23
Dots: 16 BG: 2
Dots: 16 BG: 2
3_203
Dots: 34 BG: 0
Dots: 24 BG: 0

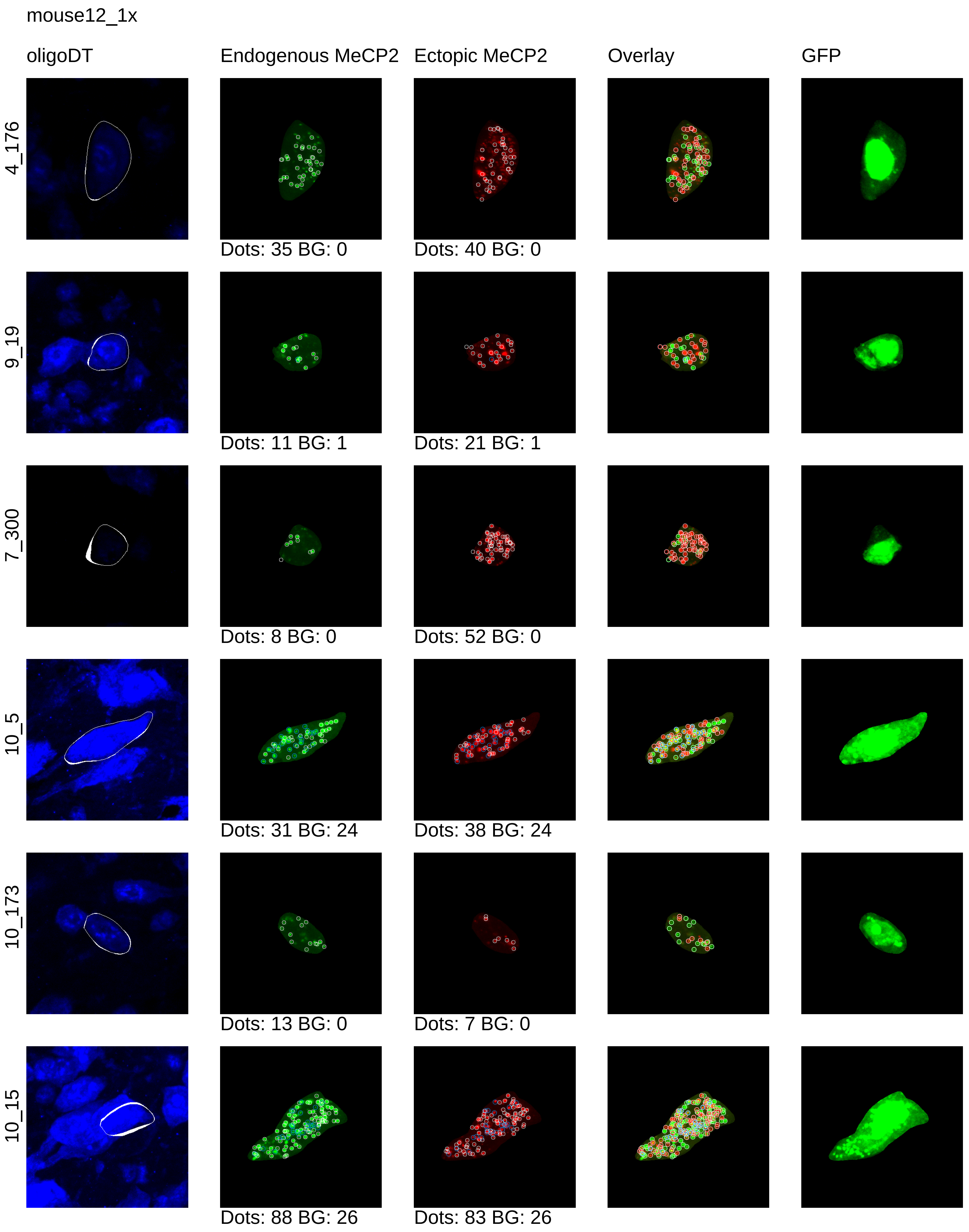
mouse12_1x
oligoDT
Endogenous MeCP2
Ectopic MeCP2
Overlay
GFP
4_176
Dots: 35 BG: 0
Dots: 40 BG: 0
9_19
Dots: 11 BG: 1
Dots: 21 BG: 1
7_300
Dots: 8 BG: 0
Dots: 52 BG: 0
10_5
Dots: 31 BG: 24
Dots: 38 BG: 24
10_173
Dots: 13 BG: 0
Dots: 7 BG: 0
10_15
Dots: 88 BG: 26
Dots: 83 BG: 26

mouse12_1x

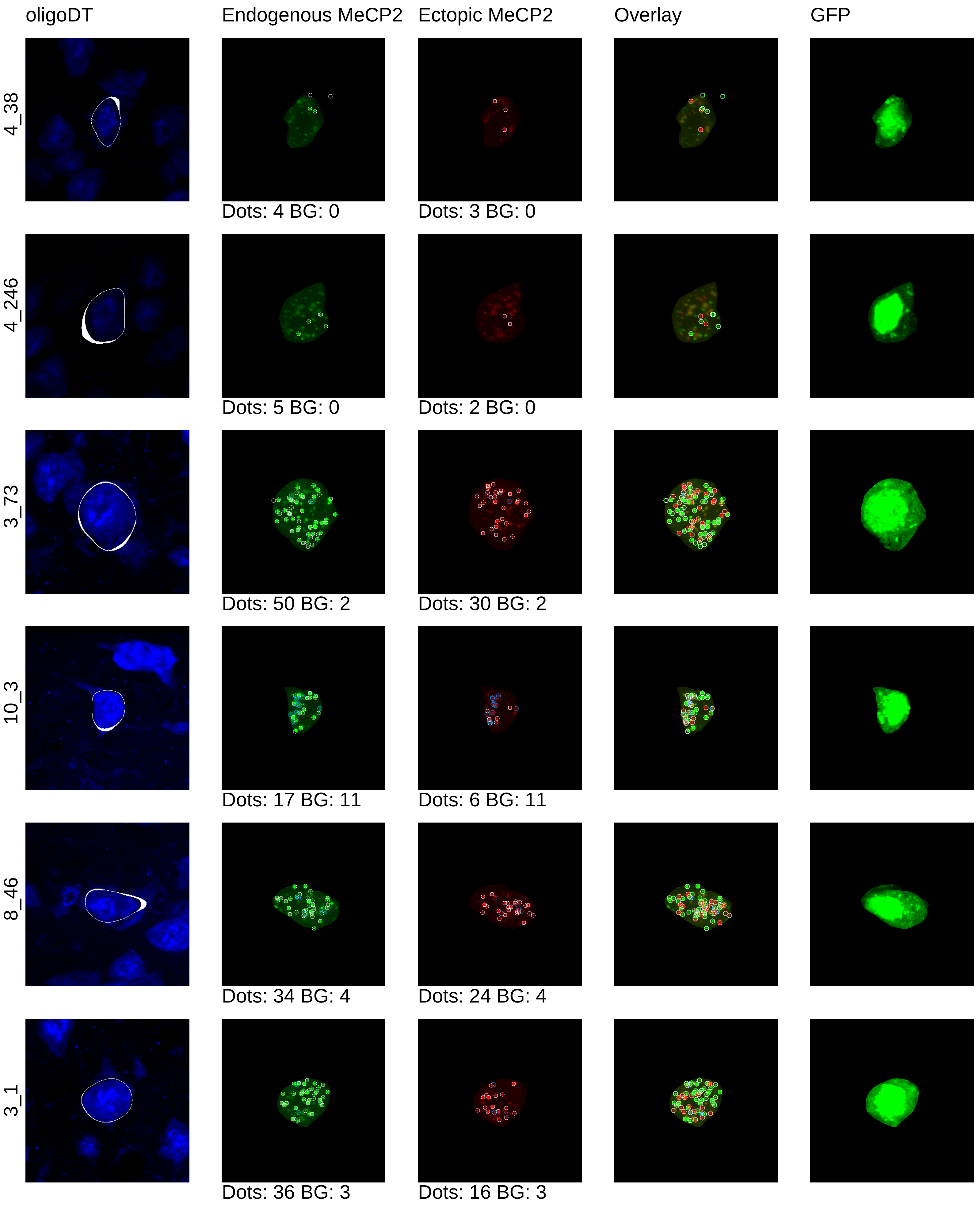
oligoDT
Endogenous MeCP2
Ectopic MeCP2
Overlay
GFP
4_38
Dots: 4 BG: 0
Dots: 3 BG: 0
4_246
Dots: 5 BG: 0
Dots: 2 BG: 0
3_73
Dots: 50 BG: 2
Dots: 30 BG: 2
10_3
Dots: 17 BG: 11
Dots: 6 BG: 11
8_46
Dots: 34 BG: 4
Dots: 24 BG: 4
3_1
Dots: 36 BG: 3
Dots: 16 BG: 3

mouse12_1x

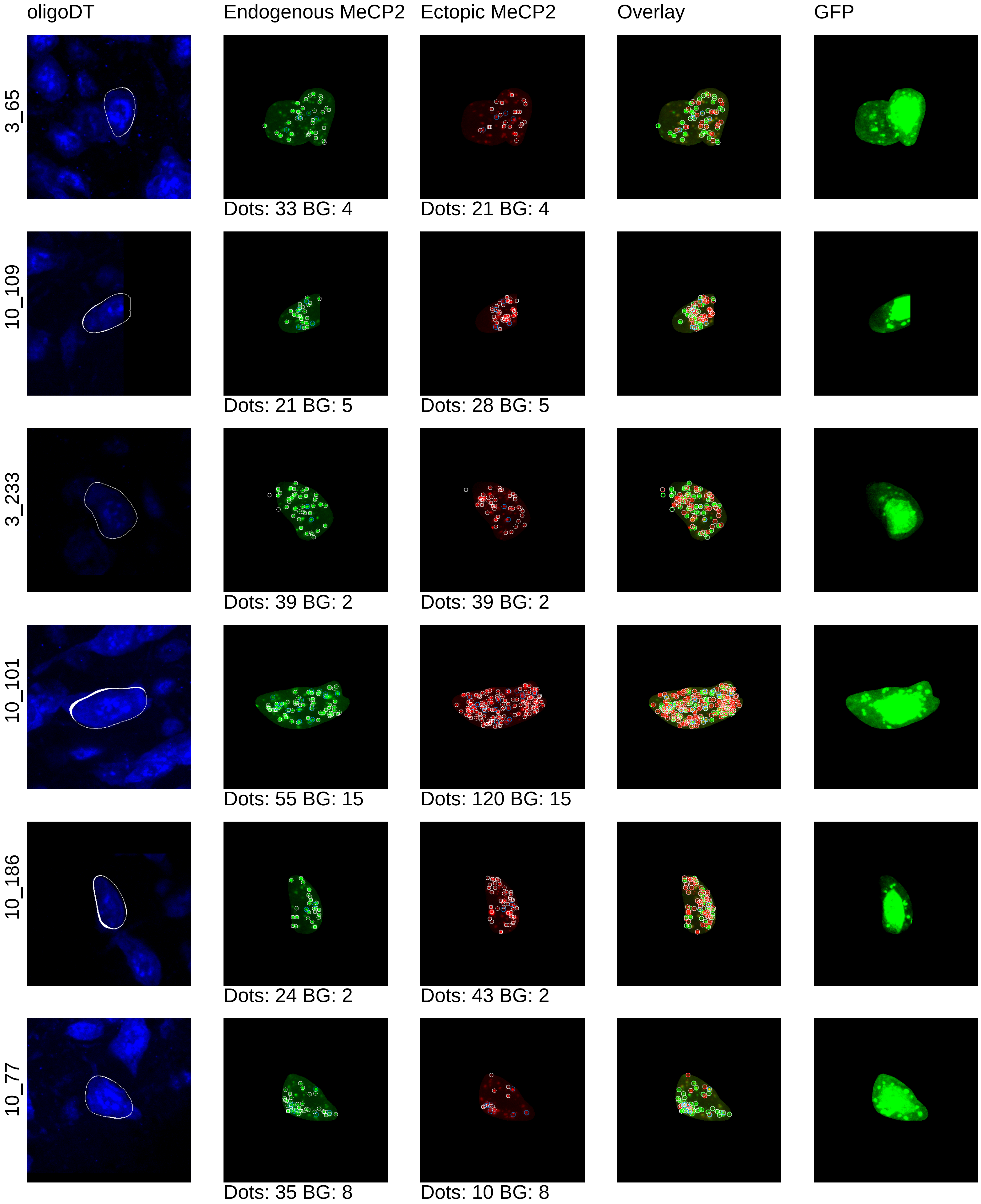
oligoDT
Endogenous MeCP2
Ectopic MeCP2
Overlay
GFP
3_65
Dots: 33 BG: 4
Dots: 21 BG: 4
10_109
Dots: 21 BG: 5
Dots: 28 BG: 5
3_233
Dots: 39 BG: 2
Dots: 39 BG: 2
10_101
Dots: 55 BG: 15
Dots: 120 BG: 15
10_186
Dots: 24 BG: 2
Dots: 43 BG: 2
10_77
Dots: 35 BG: 8
Dots: 10 BG: 8

mouse12_1x

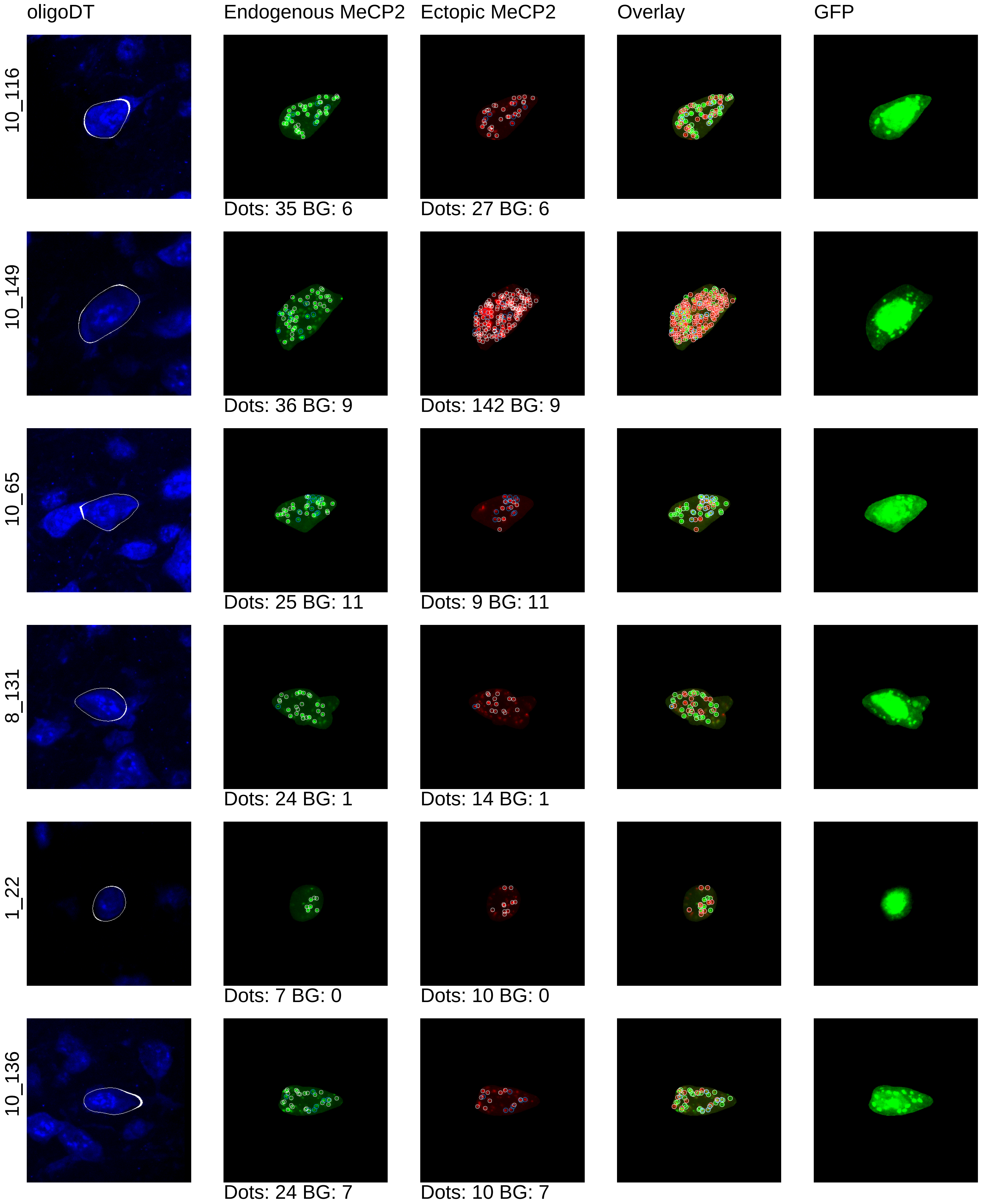
oligoDT
Endogenous MeCP2
Ectopic MeCP2
Overlay
GFP
10_116
Dots: 35 BG: 6
Dots: 27 BG: 6
10_149
Dots: 36 BG: 9
Dots: 142 BG: 9
10_65
Dots: 25 BG: 11
Dots: 9 BG: 11
8_131
Dots: 24 BG: 1
Dots: 14 BG: 1
1_22
Dots: 7 BG: 0
Dots: 10 BG: 0
10_136
Dots: 24 BG: 7
Dots: 10 BG: 7

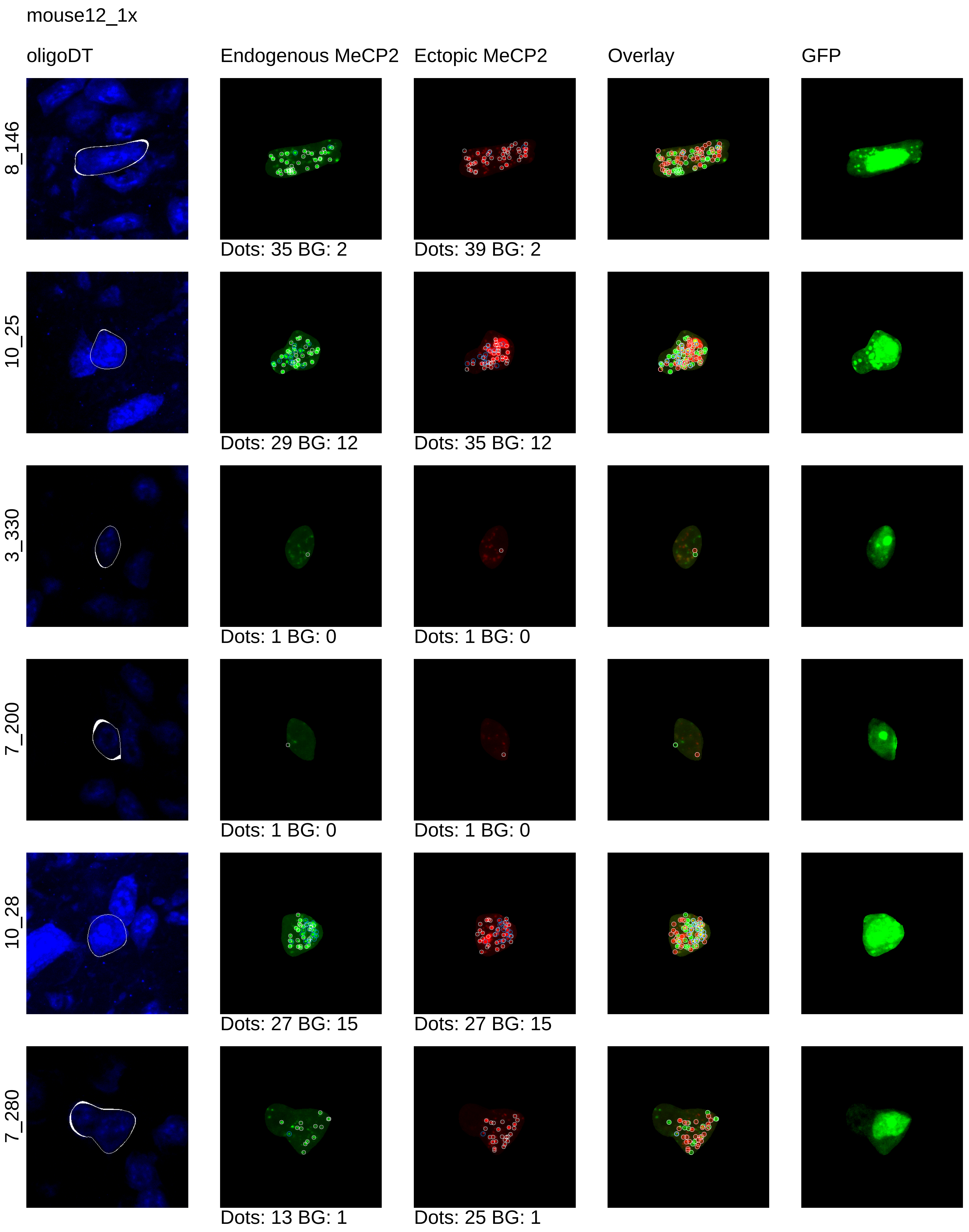
mouse12_1x
oligoDT
Endogenous MeCP2
Ectopic MeCP2
Overlay
GFP
8_146
Dots: 35 BG: 2
Dots: 39 BG: 2
10_25
Dots: 29 BG: 12
Dots: 35 BG: 12
3_330
Dots: 1 BG: 0
Dots: 1 BG: 0
7_200
Dots: 1 BG: 0
Dots: 1 BG: 0
10_28
Dots: 27 BG: 15
Dots: 27 BG: 15
7_280
Dots: 13 BG: 1
Dots: 25 BG: 1

mouse12_1x

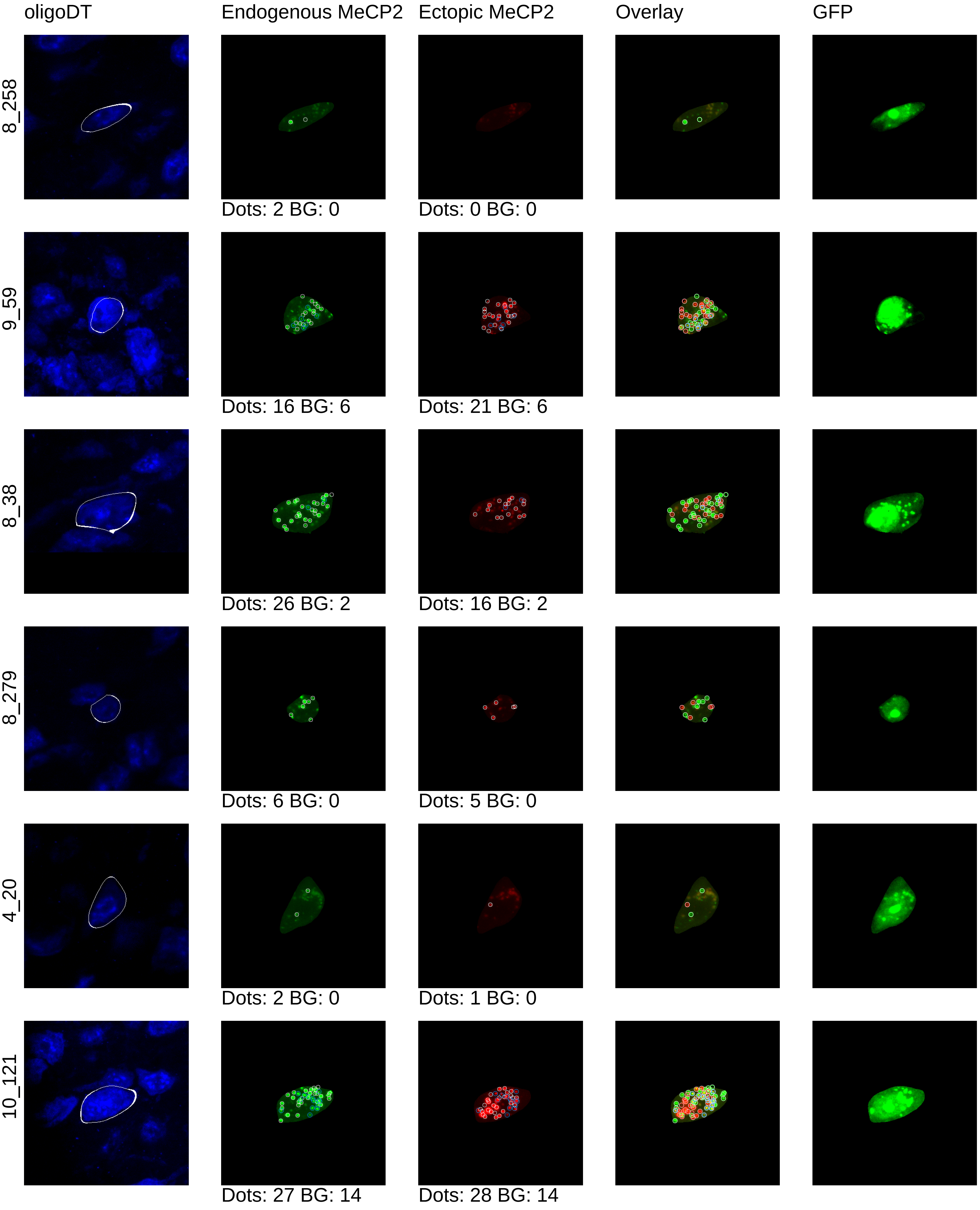

oligoDT
Endogenous MeCP2
Ectopic MeCP2
Overlay
GFP
8_258
Dots: 2 BG: 0
Dots: 0 BG: 0
9_59
Dots: 16 BG: 6
Dots: 21 BG: 6
8_38
Dots: 26 BG: 2
Dots: 16 BG: 2
8_279
Dots: 6 BG: 0
Dots: 5 BG: 0
4_20
Dots: 2 BG: 0
Dots: 1 BG: 0
10_121
Dots: 27 BG: 14
Dots: 28 BG: 14

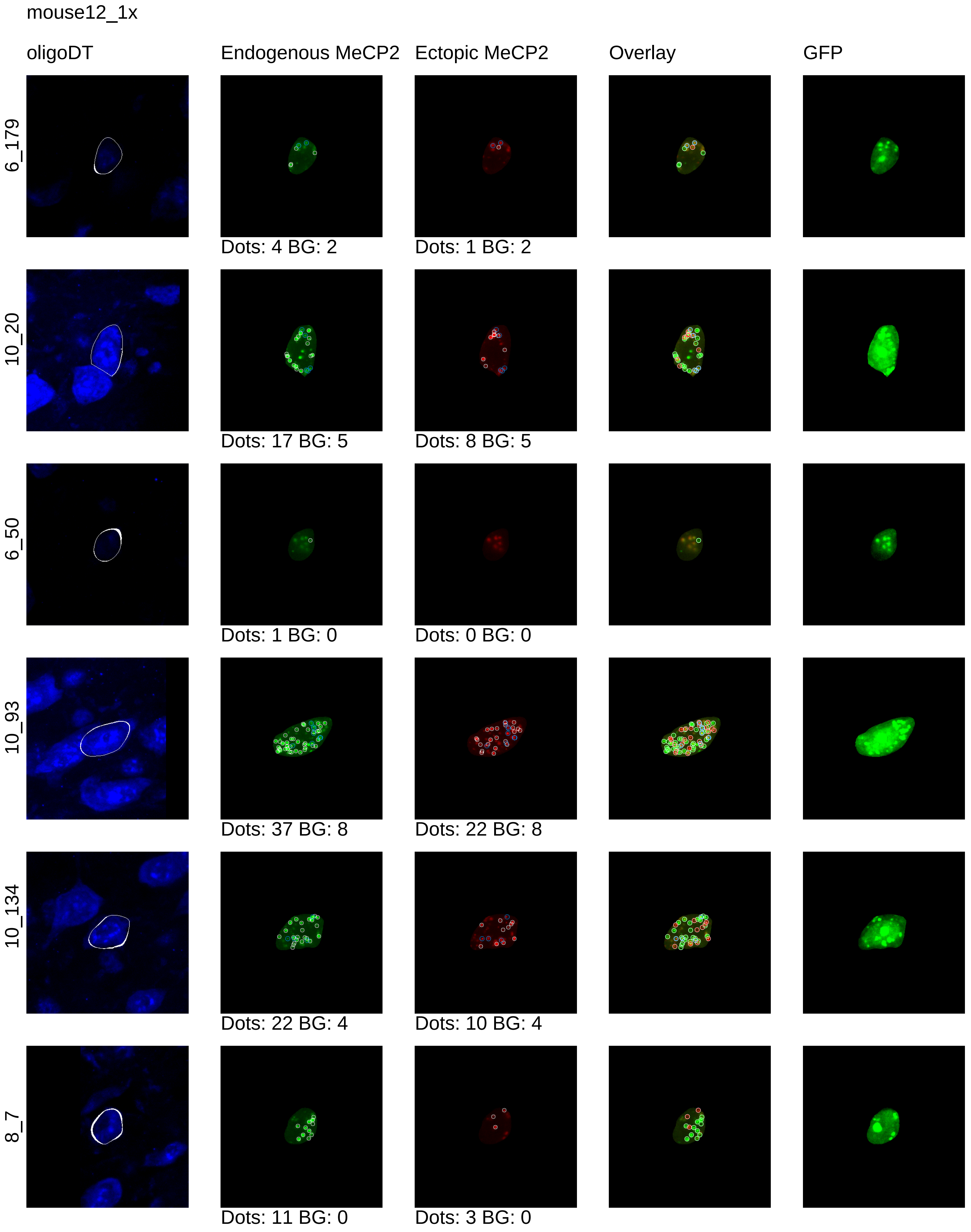
mouse12_1x
oligoDT
Endogenous MeCP2
Ectopic MeCP2
Overlay
GFP
6_179
Dots: 4 BG: 2
Dots: 1 BG: 2
10_20
Dots: 17 BG: 5
Dots: 8 BG: 5
6_50
Dots: 1 BG: 0
Dots: 0 BG: 0
10_93
Dots: 37 BG: 8
Dots: 22 BG: 8
10_134
Dots: 22 BG: 4
Dots: 10 BG: 4
8_7
Dots: 11 BG: 0
Dots: 3 BG: 0

mouse12_1x

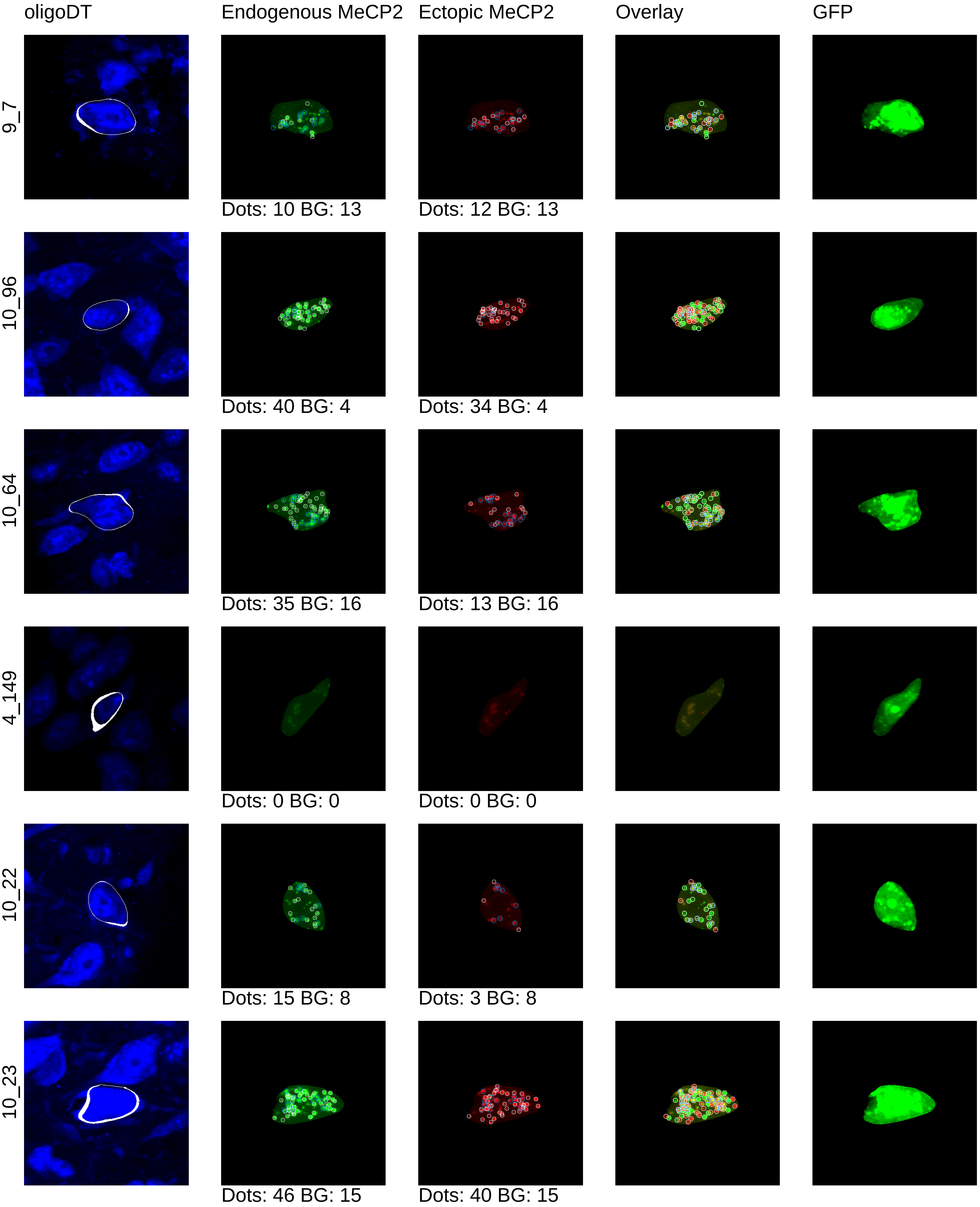
oligoDT
Endogenous MeCP2
Ectopic MeCP2
Overlay
GFP
9_7
Dots: 10 BG: 13
Dots: 12 BG: 13
10_96
Dots: 40 BG: 4
Dots: 34 BG: 4
10_64
Dots: 35 BG: 16
Dots: 13 BG: 16
4_149
Dots: 0 BG: 0
Dots: 0 BG: 0
10_22
Dots: 15 BG: 8
Dots: 3 BG: 8
10_23
Dots: 46 BG: 15
Dots: 40 BG: 15

mouse12_1x

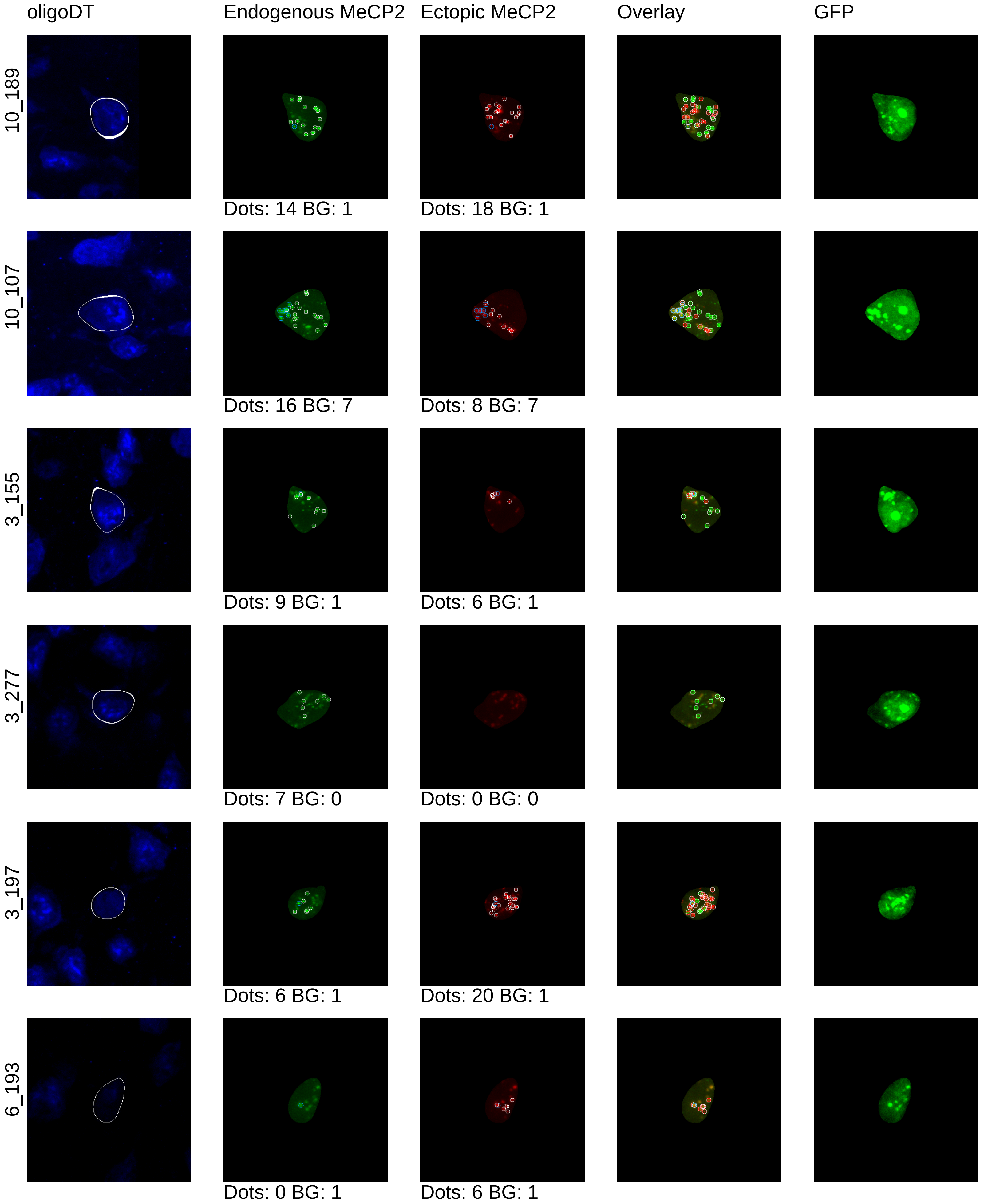

oligoDT
Endogenous MeCP2
Ectopic MeCP2
Overlay
GFP
10_189
Dots: 14 BG: 1
Dots: 18 BG: 1
10_107
Dots: 16 BG: 7
Dots: 8 BG: 7
3_155
Dots: 9 BG: 1
Dots: 6 BG: 1
3_277
Dots: 7 BG: 0
Dots: 0 BG: 0
3_197
Dots: 6 BG: 1
Dots: 20 BG: 1
6_193
Dots: 0 BG: 1
Dots: 6 BG: 1

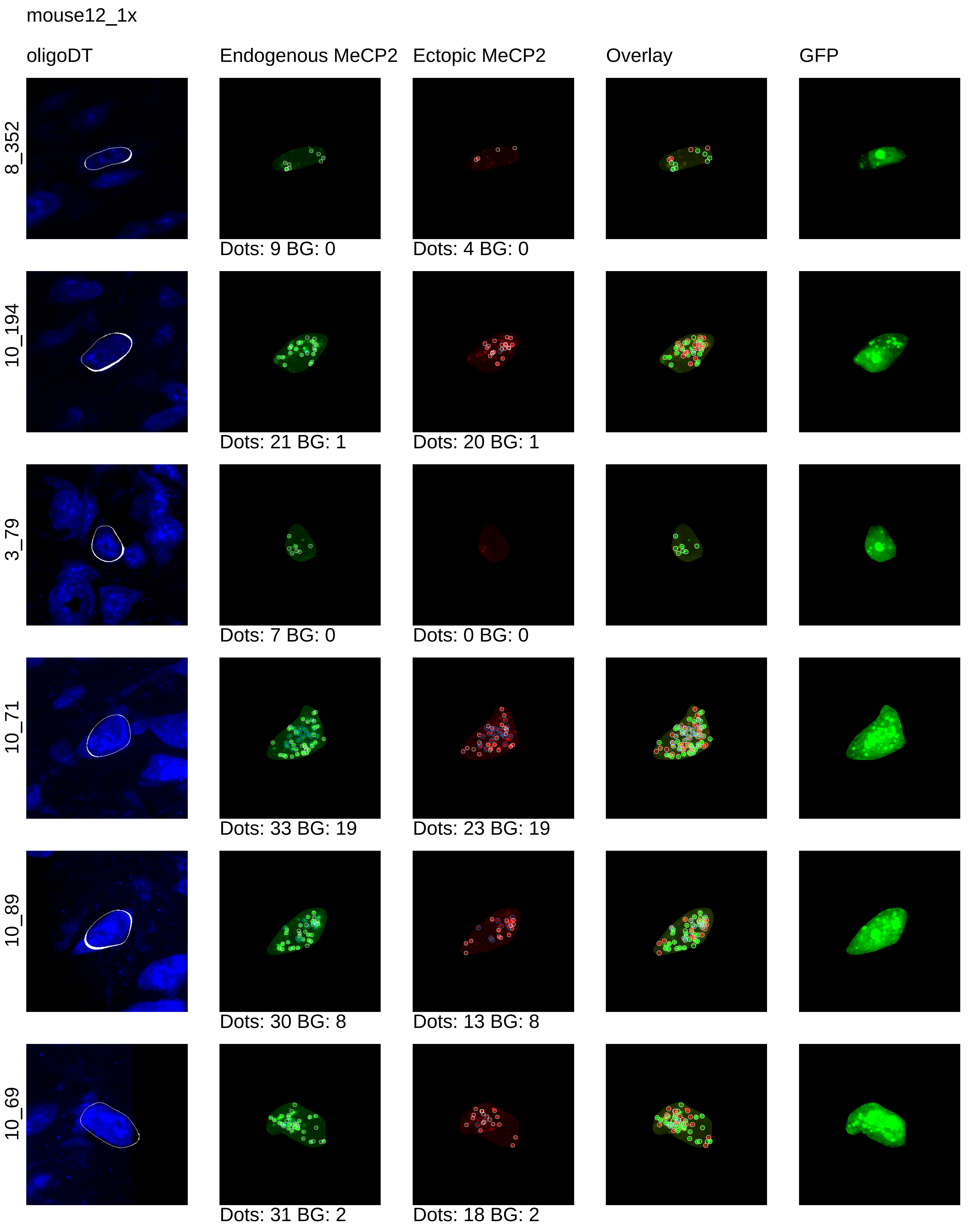
mouse12_1x
oligoDT
Endogenous MeCP2
Ectopic MeCP2
Overlay
GFP
8_352
Dots: 9 BG: 0
Dots: 4 BG: 0
10_194
Dots: 21 BG: 1
Dots: 20 BG: 1
3_79
Dots: 7 BG: 0
Dots: 0 BG: 0
10_71
Dots: 33 BG: 19
Dots: 23 BG: 19
10_89
Dots: 30 BG: 8
Dots: 13 BG: 8
10_69
Dots: 31 BG: 2
Dots: 18 BG: 2

mouse12_1x

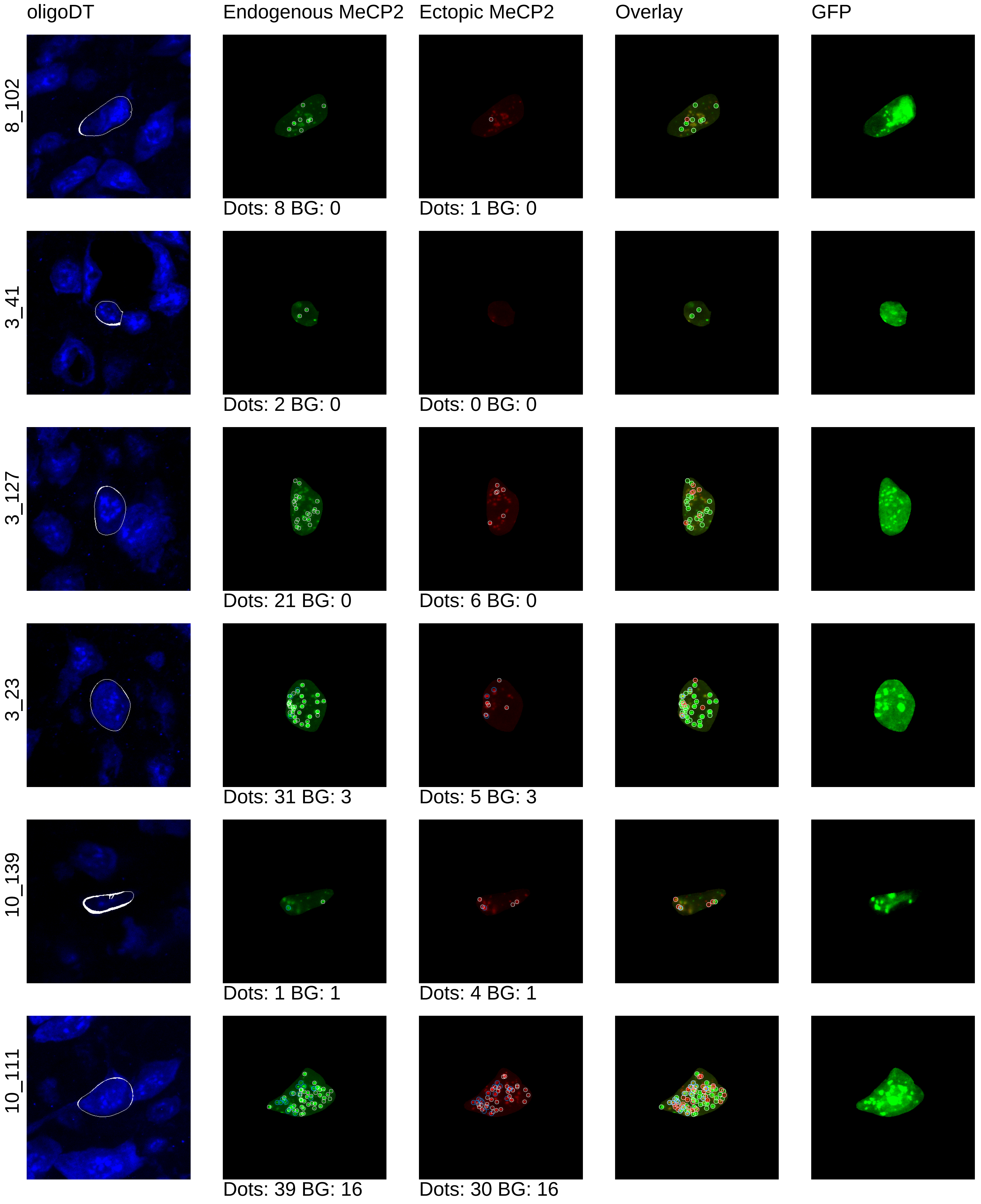
oligoDT
Endogenous MeCP2
Ectopic MeCP2
Overlay
GFP
8_102
Dots: 8 BG: 0
Dots: 1 BG: 0
3_41
Dots: 2 BG: 0
Dots: 0 BG: 0
3_127
Dots: 21 BG: 0
Dots: 6 BG: 0
3_23
Dots: 31 BG: 3
Dots: 5 BG: 3
10_139
Dots: 1 BG: 1
Dots: 4 BG: 1
10_111
Dots: 39 BG: 16
Dots: 30 BG: 16

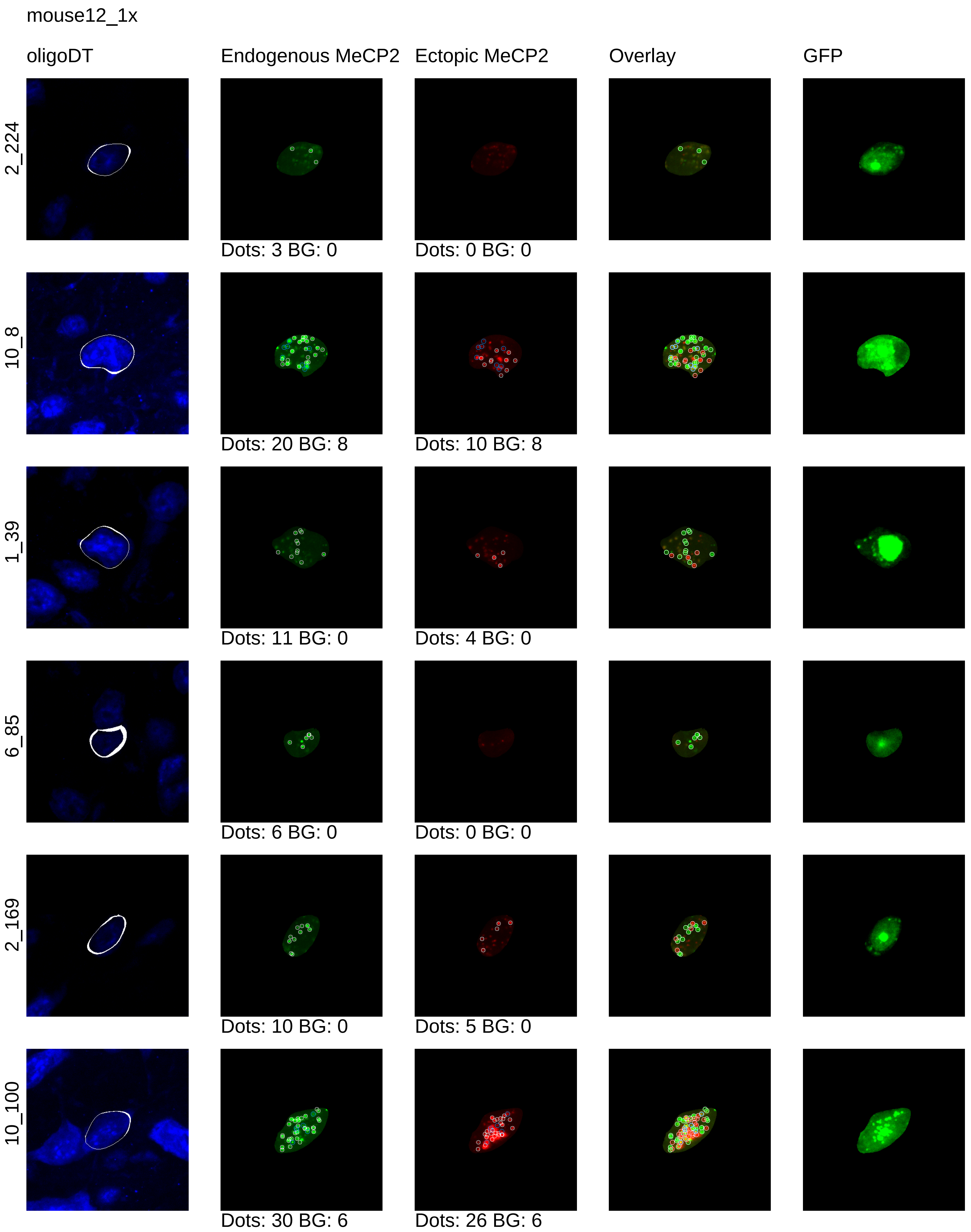
mouse12_1x
oligoDT
Endogenous MeCP2
Ectopic MeCP2
Overlay
GFP
2_224
Dots: 3 BG: 0
Dots: 0 BG: 0
10_8
Dots: 20 BG: 8
Dots: 10 BG: 8
1_39
Dots: 11 BG: 0
Dots: 4 BG: 0
6_85
Dots: 6 BG: 0
Dots: 0 BG: 0
2_169
Dots: 10 BG: 0
Dots: 5 BG: 0
10_100
Dots: 30 BG: 6
Dots: 26 BG: 6

mouse12_1x

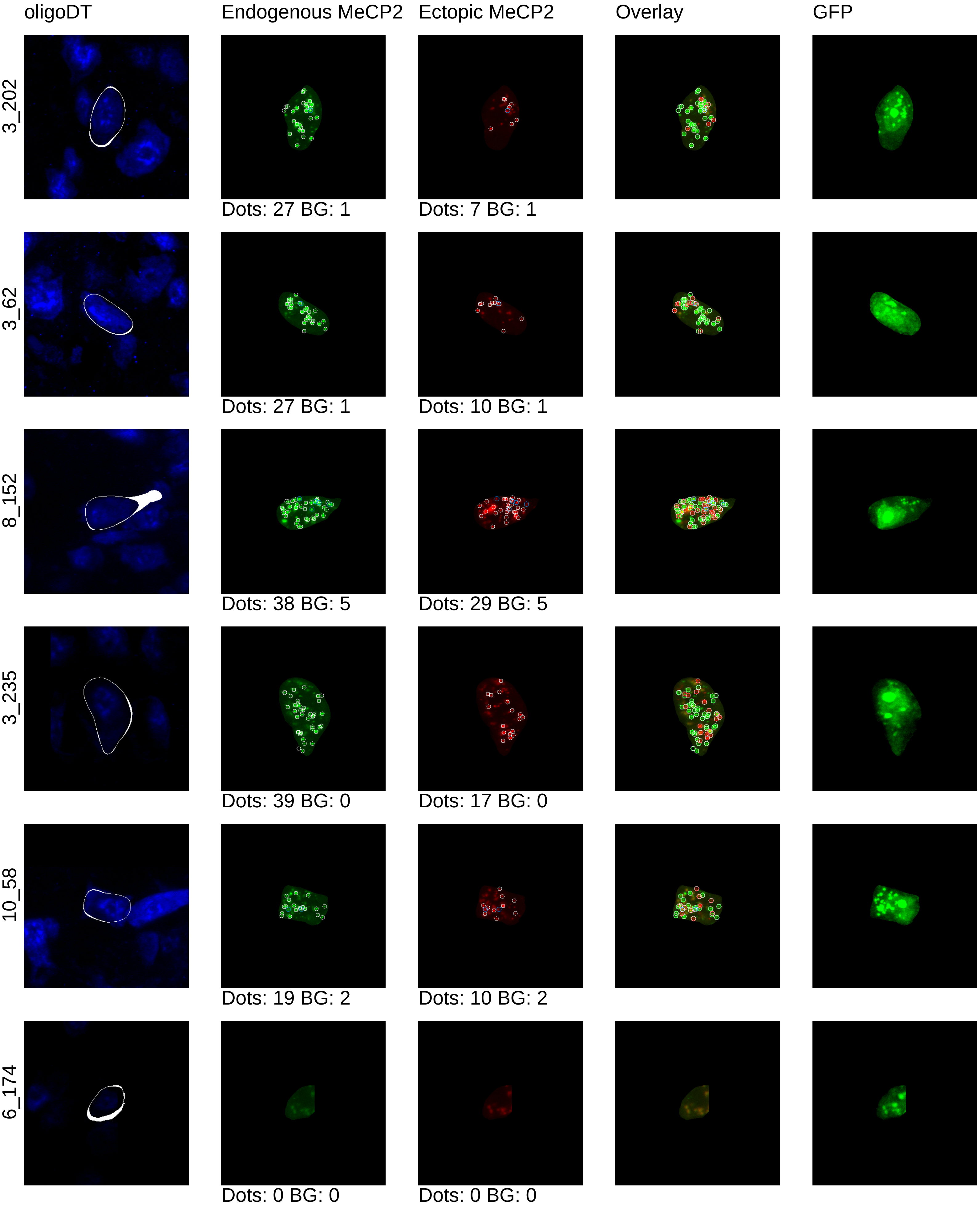
oligoDT
Endogenous MeCP2
Ectopic MeCP2
Overlay
GFP
3_202
Dots: 27 BG: 1
Dots: 7 BG: 1
3_62
Dots: 27 BG: 1
Dots: 10 BG: 1
8_152
Dots: 38 BG: 5
Dots: 29 BG: 5
3_235
Dots: 39 BG: 0
Dots: 17 BG: 0
10_58
Dots: 19 BG: 2
Dots: 10 BG: 2
6_174
Dots: 0 BG: 0
Dots: 0 BG: 0

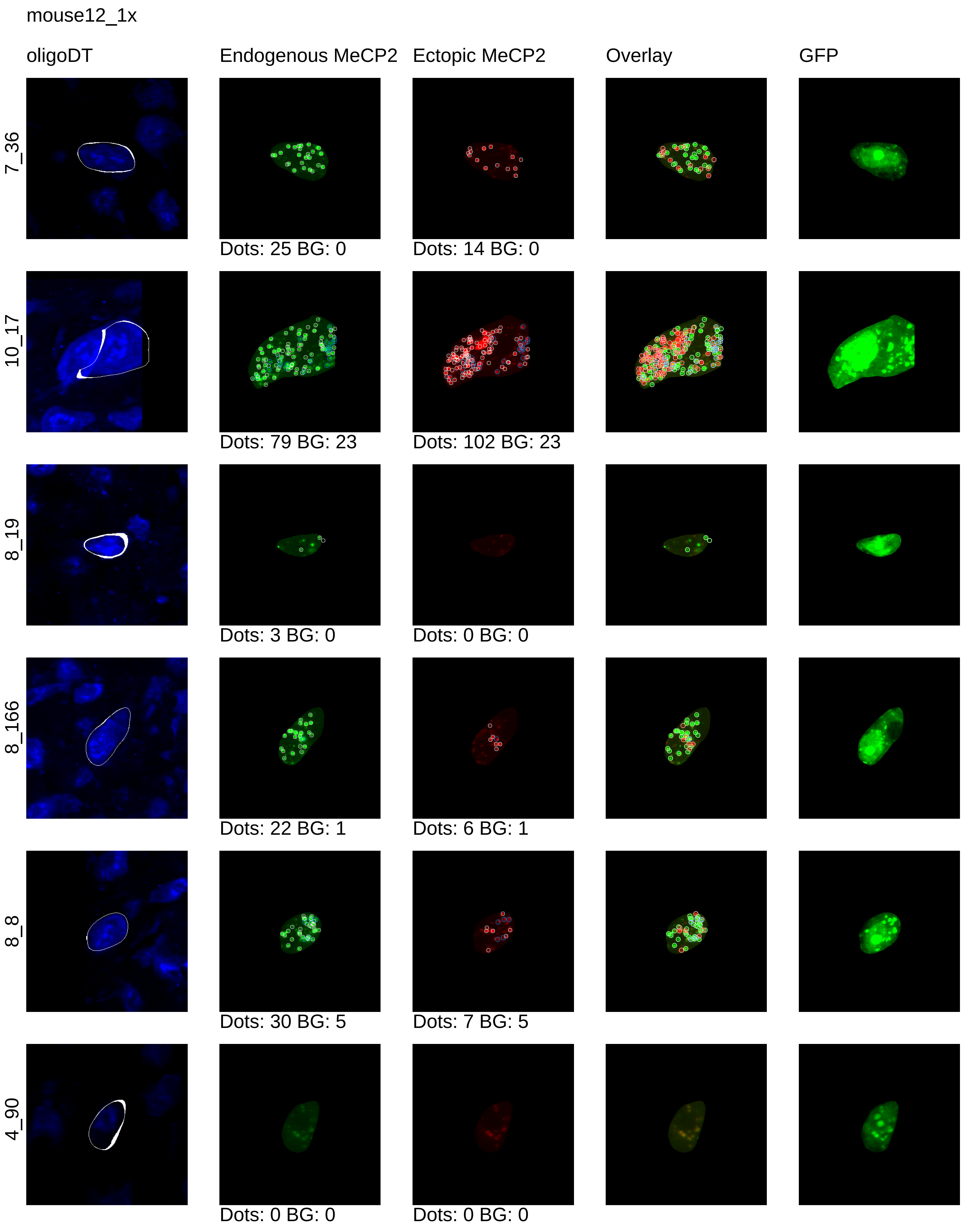
mouse12_1x
oligoDT
Endogenous MeCP2
Ectopic MeCP2
Overlay
GFP
7_36
Dots: 25 BG: 0
Dots: 14 BG: 0
10_17
Dots: 79 BG: 23
Dots: 102 BG: 23
8_19
Dots: 3 BG: 0
Dots: 0 BG: 0
8_166
Dots: 22 BG: 1
Dots: 6 BG: 1
8_8
Dots: 30 BG: 5
Dots: 7 BG: 5
4_90
Dots: 0 BG: 0
Dots: 0 BG: 0

mouse12_1x

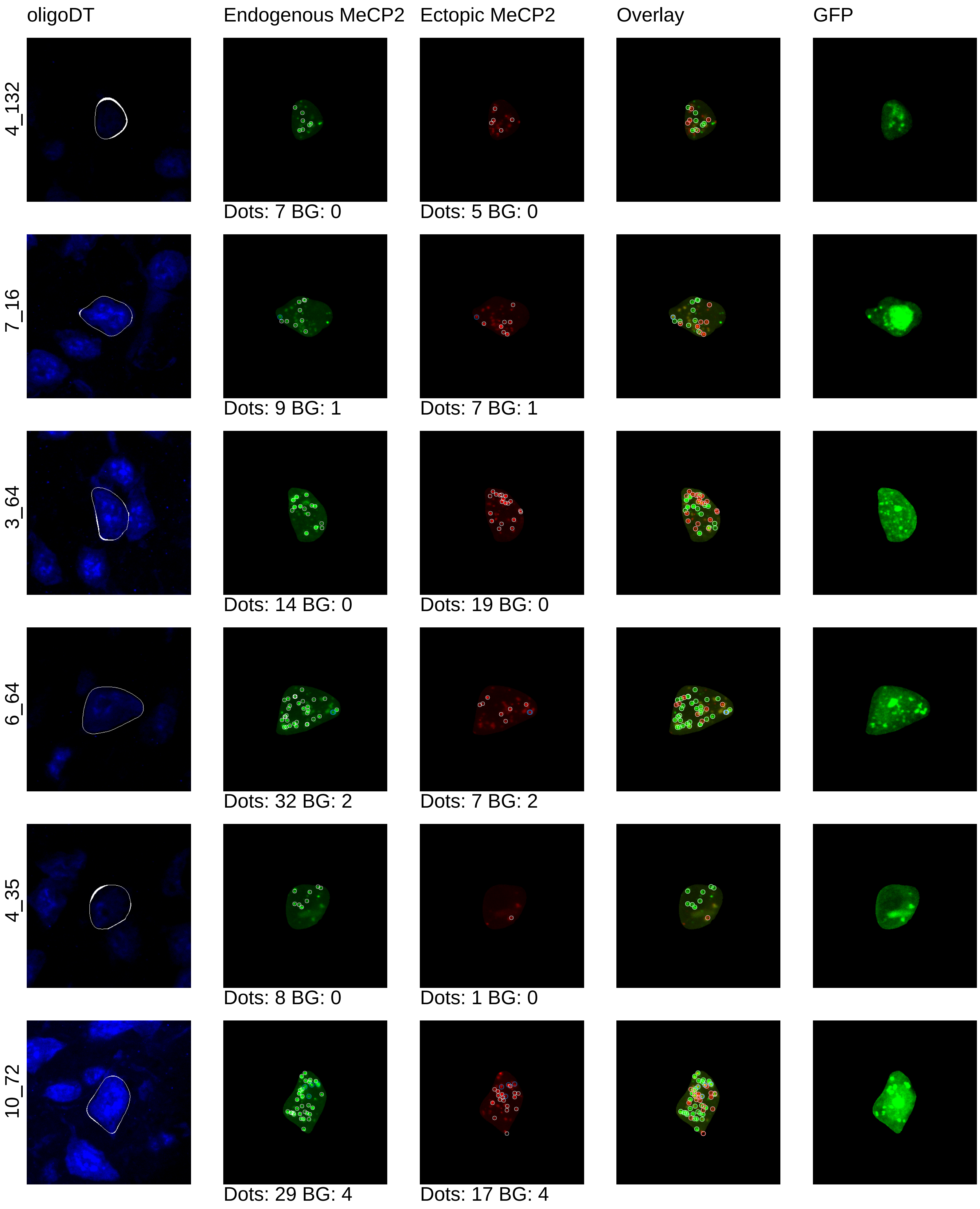
oligoDT
Endogenous MeCP2
Ectopic MeCP2
Overlay
GFP
4_132
Dots: 7 BG: 0
Dots: 5 BG: 0
7_16
Dots: 9 BG: 1
Dots: 7 BG: 1
3_64
Dots: 14 BG: 0
Dots: 19 BG: 0
6_64
Dots: 32 BG: 2
Dots: 7 BG: 2
4_35
Dots: 8 BG: 0
Dots: 1 BG: 0
10_72
Dots: 29 BG: 4
Dots: 17 BG: 4

mouse12_1x

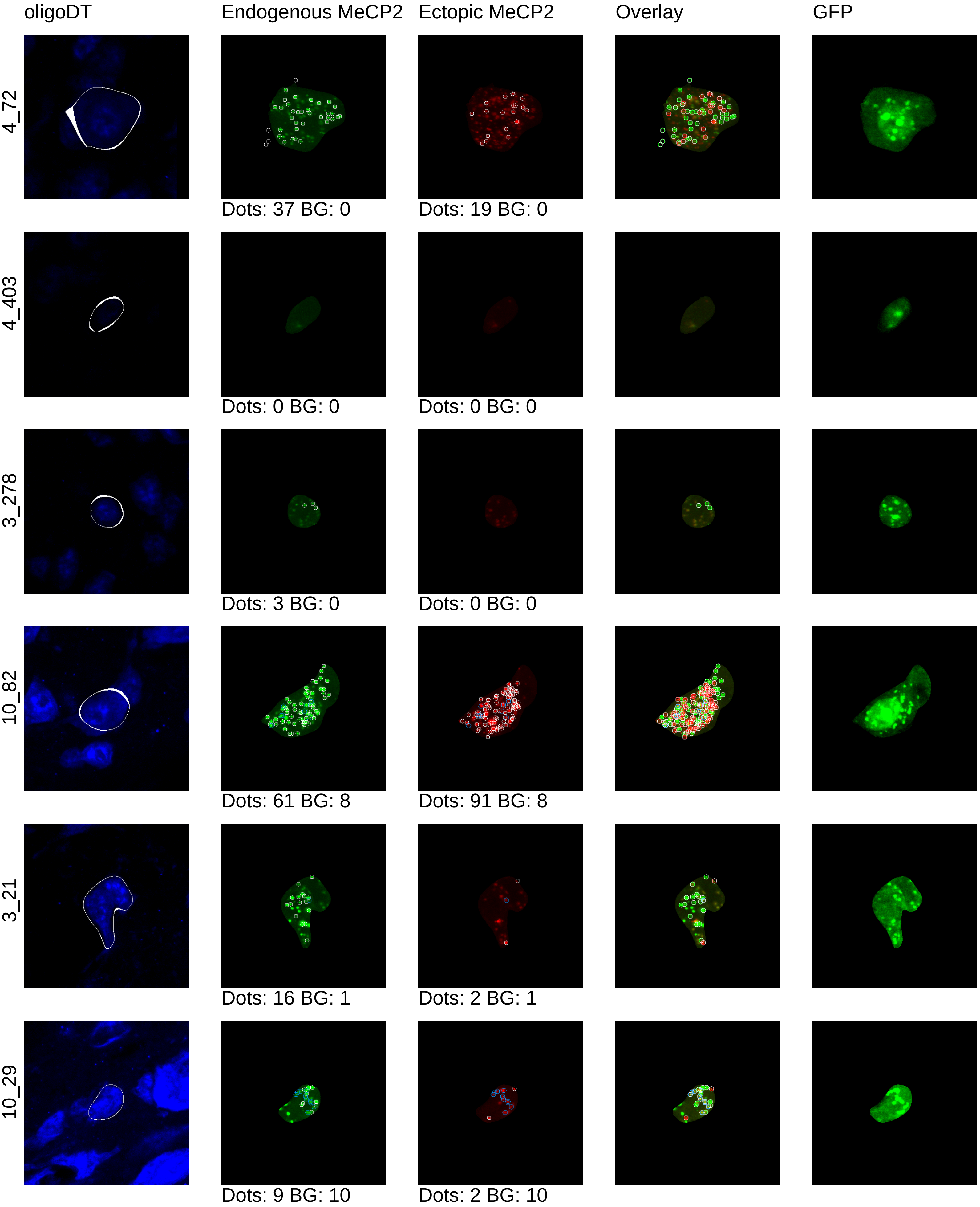

oligoDT
Endogenous MeCP2
Ectopic MeCP2
Overlay
GFP
4_72
Dots: 37 BG: 0
Dots: 19 BG: 0
4_403
Dots: 0 BG: 0
Dots: 0 BG: 0
3_278
Dots: 3 BG: 0
Dots: 0 BG: 0
10_82
Dots: 61 BG: 8
Dots: 91 BG: 8
3_21
Dots: 16 BG: 1
Dots: 2 BG: 1
10_29
Dots: 9 BG: 10
Dots: 2 BG: 10

mouse12_1x

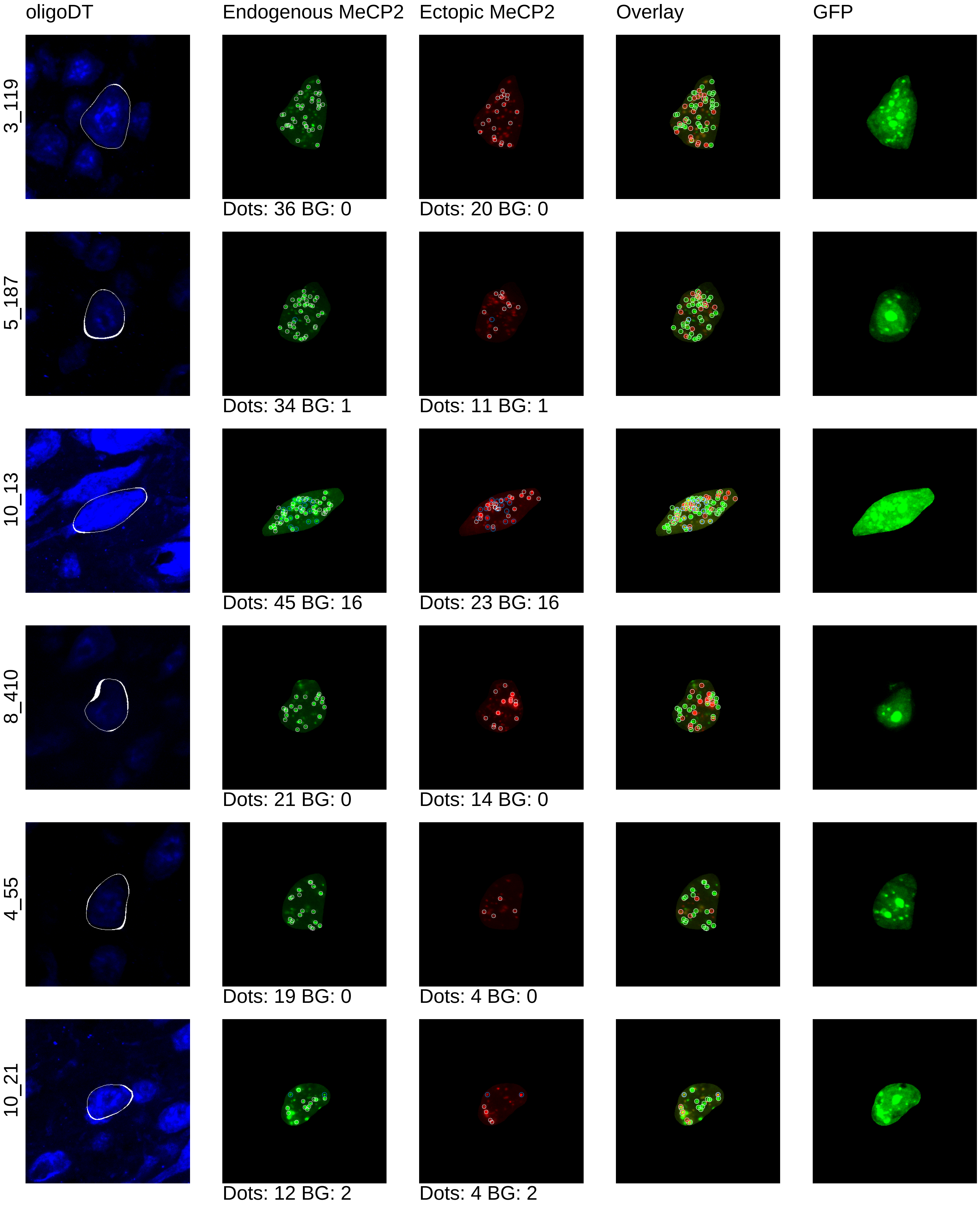
oligoDT
Endogenous MeCP2
Ectopic MeCP2
Overlay
GFP
3_119
Dots: 36 BG: 0
Dots: 20 BG: 0
5_187
Dots: 34 BG: 1
Dots: 11 BG: 1
10_13
Dots: 45 BG: 16
Dots: 23 BG: 16
8_410
Dots: 21 BG: 0
Dots: 14 BG: 0
4_55
Dots: 19 BG: 0
Dots: 4 BG: 0
10_21
Dots: 12 BG: 2
Dots: 4 BG: 2

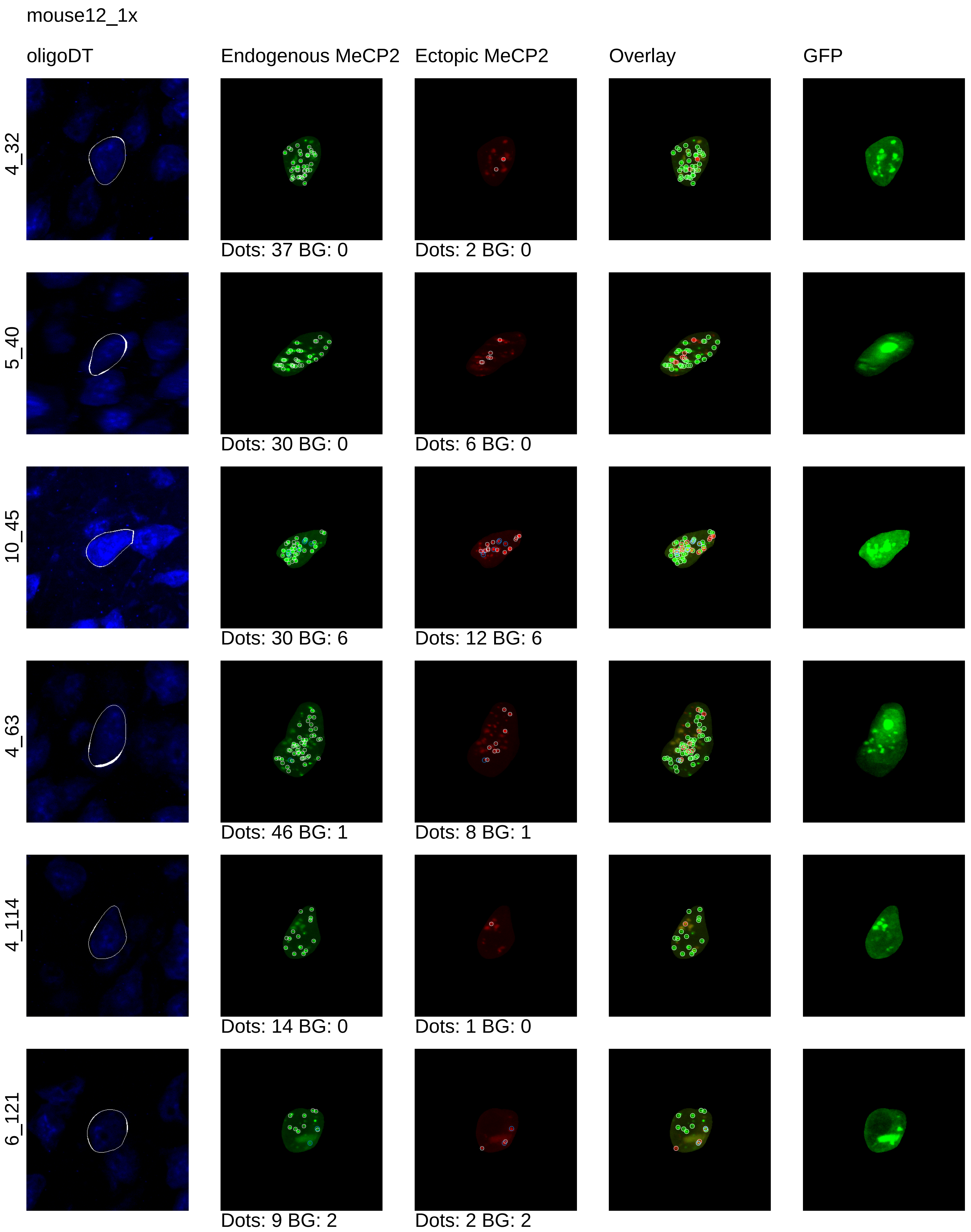
mouse12_1x
oligoDT
Endogenous MeCP2
Ectopic MeCP2
Overlay
GFP
4_32
Dots: 37 BG: 0
Dots: 2 BG: 0
5_40
Dots: 30 BG: 0
Dots: 6 BG: 0
10_45
Dots: 30 BG: 6
Dots: 12 BG: 6
4_63
Dots: 46 BG: 1
Dots: 8 BG: 1
4_114
Dots: 14 BG: 0
Dots: 1 BG: 0
6_121
Dots: 9 BG: 2
Dots: 2 BG: 2

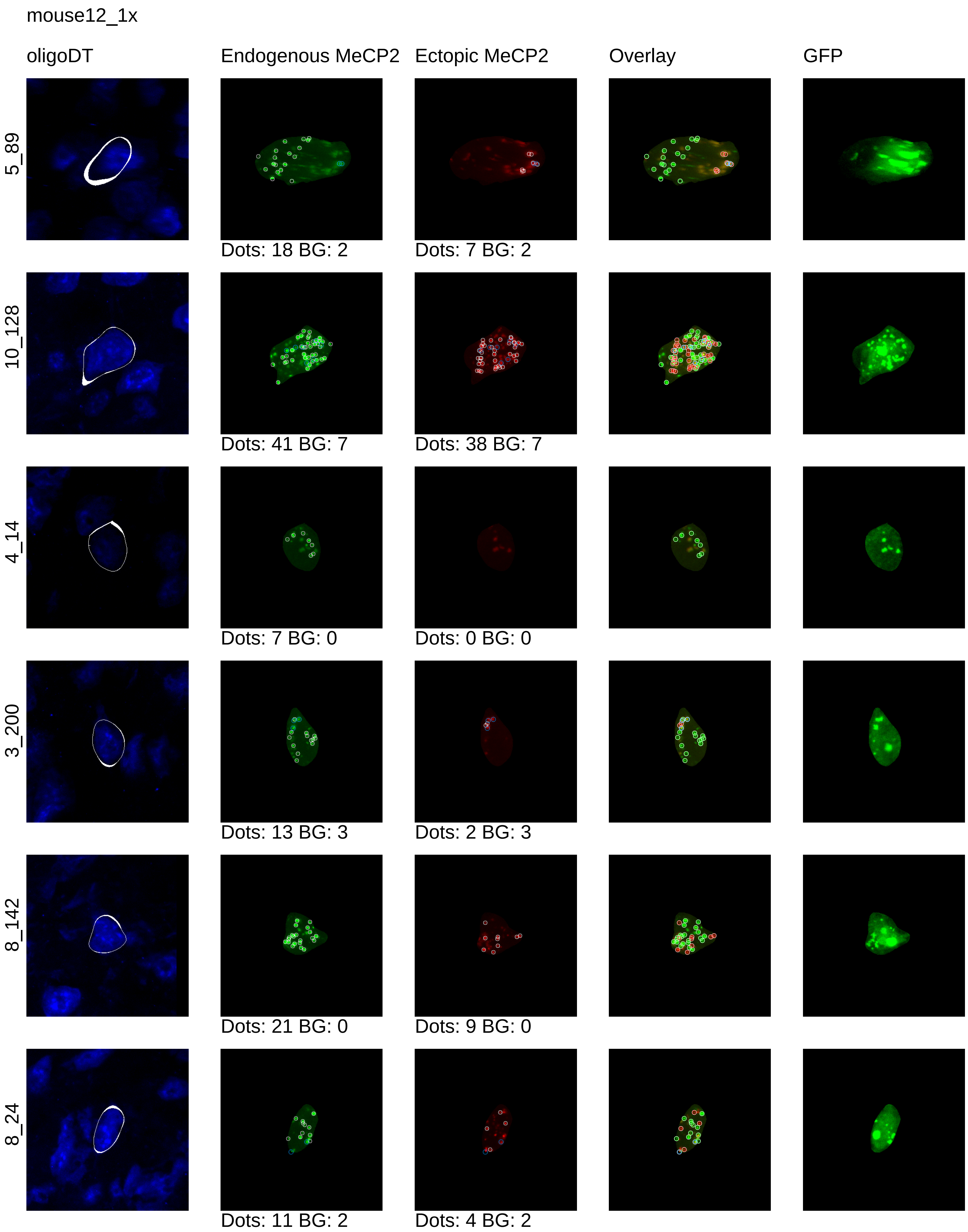
mouse12_1x
oligoDT
Endogenous MeCP2
Ectopic MeCP2
Overlay
GFP
5_89
Dots: 18 BG: 2
Dots: 7 BG: 2
10_128
Dots: 41 BG: 7
Dots: 38 BG: 7
4_14
Dots: 7 BG: 0
Dots: 0 BG: 0
3_200
Dots: 13 BG: 3
Dots: 2 BG: 3
8_142
Dots: 21 BG: 0
Dots: 9 BG: 0
8_24
Dots: 11 BG: 2
Dots: 4 BG: 2

mouse12_1x

oligoDT | Endogenous MeCP2 | Ectopic MeCP2 | Overlay | GFP

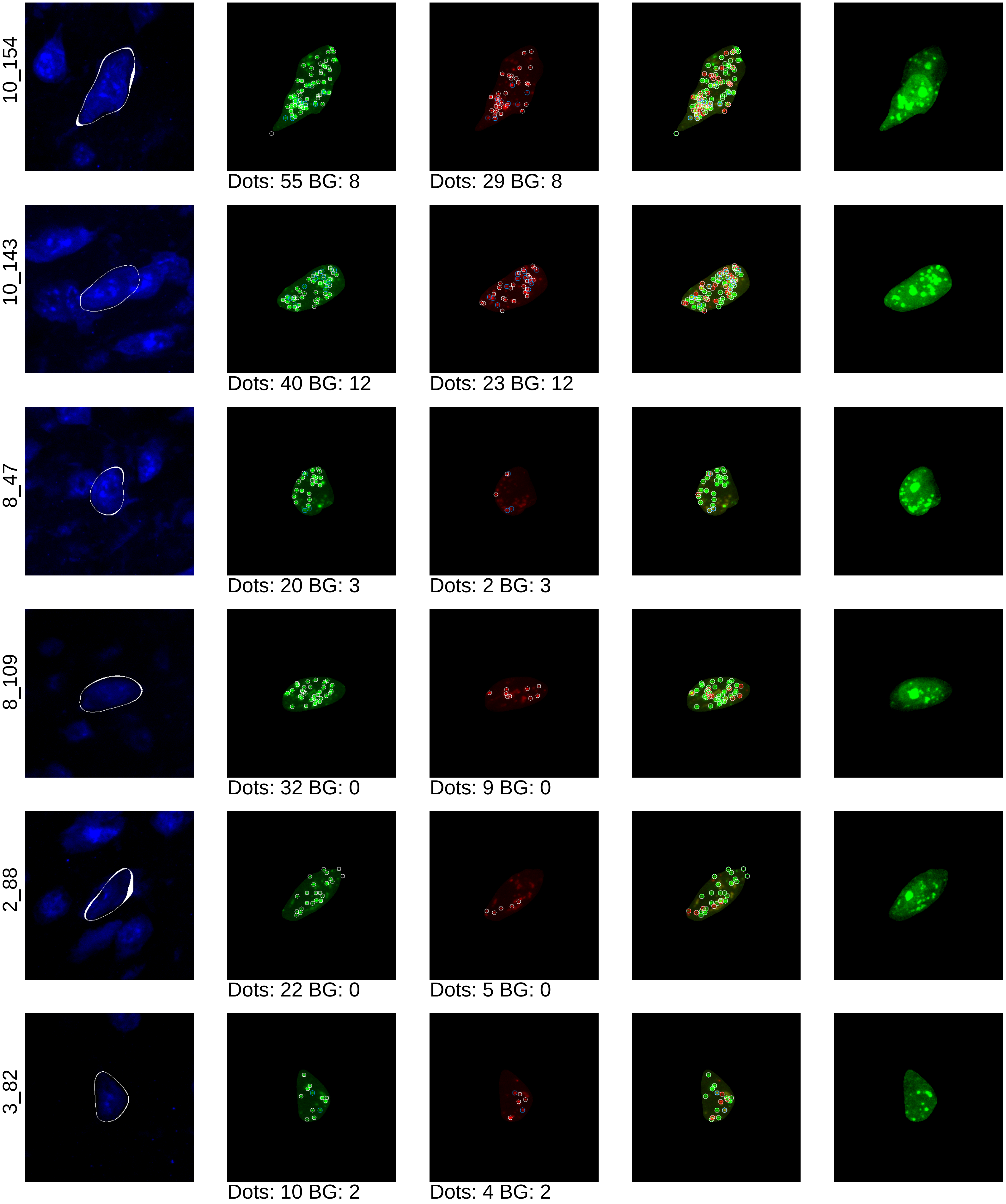

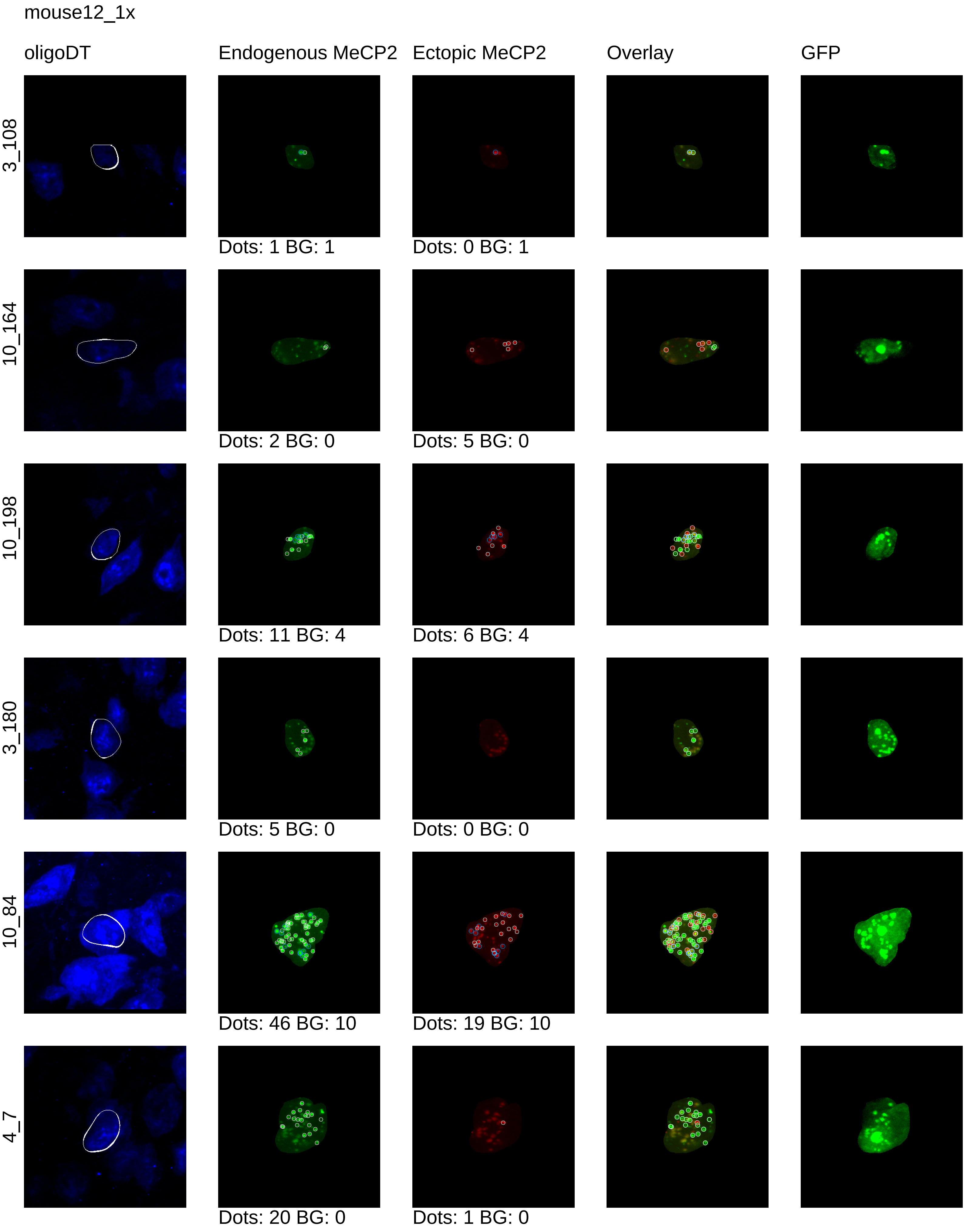
mouse12_1x
oligoDT
Endogenous MeCP2
Ectopic MeCP2
Overlay
GFP
3_108
Dots: 1 BG: 1
Dots: 0 BG: 1
10_164
Dots: 2 BG: 0
Dots: 5 BG: 0
10_198
Dots: 11 BG: 4
Dots: 6 BG: 4
3_180
Dots: 5 BG: 0
Dots: 0 BG: 0
10_84
Dots: 46 BG: 10
Dots: 19 BG: 10
4_7
Dots: 20 BG: 0
Dots: 1 BG: 0

mouse12_1x

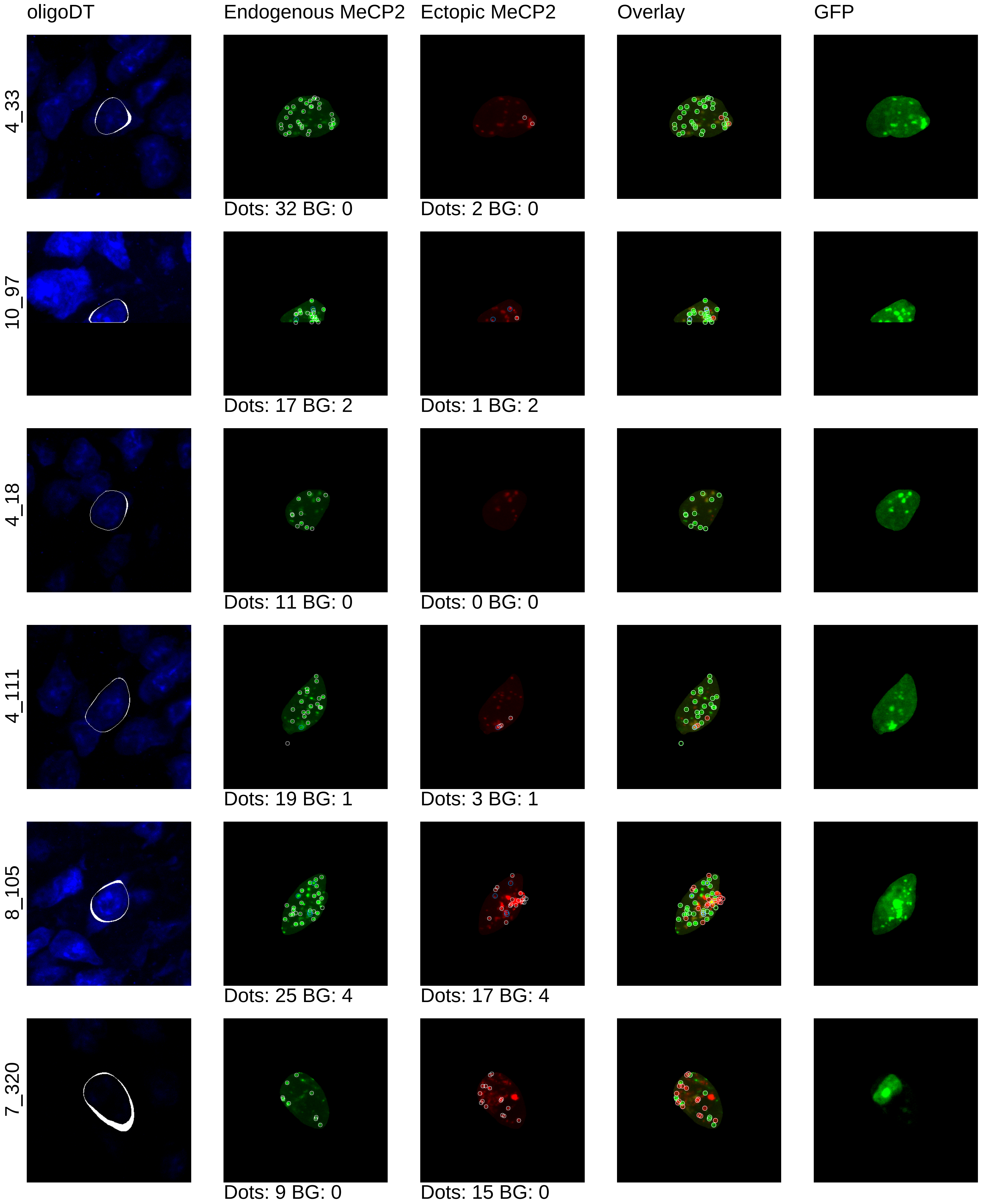
oligoDT
Endogenous MeCP2
Ectopic MeCP2
Overlay
GFP
4_33
Dots: 32 BG: 0
Dots: 2 BG: 0
10_97
Dots: 17 BG: 2
Dots: 1 BG: 2
4_18
Dots: 11 BG: 0
Dots: 0 BG: 0
4_111
Dots: 19 BG: 1
Dots: 3 BG: 1
8_105
Dots: 25 BG: 4
Dots: 17 BG: 4
7_320
Dots: 9 BG: 0
Dots: 15 BG: 0

mouse12_1x

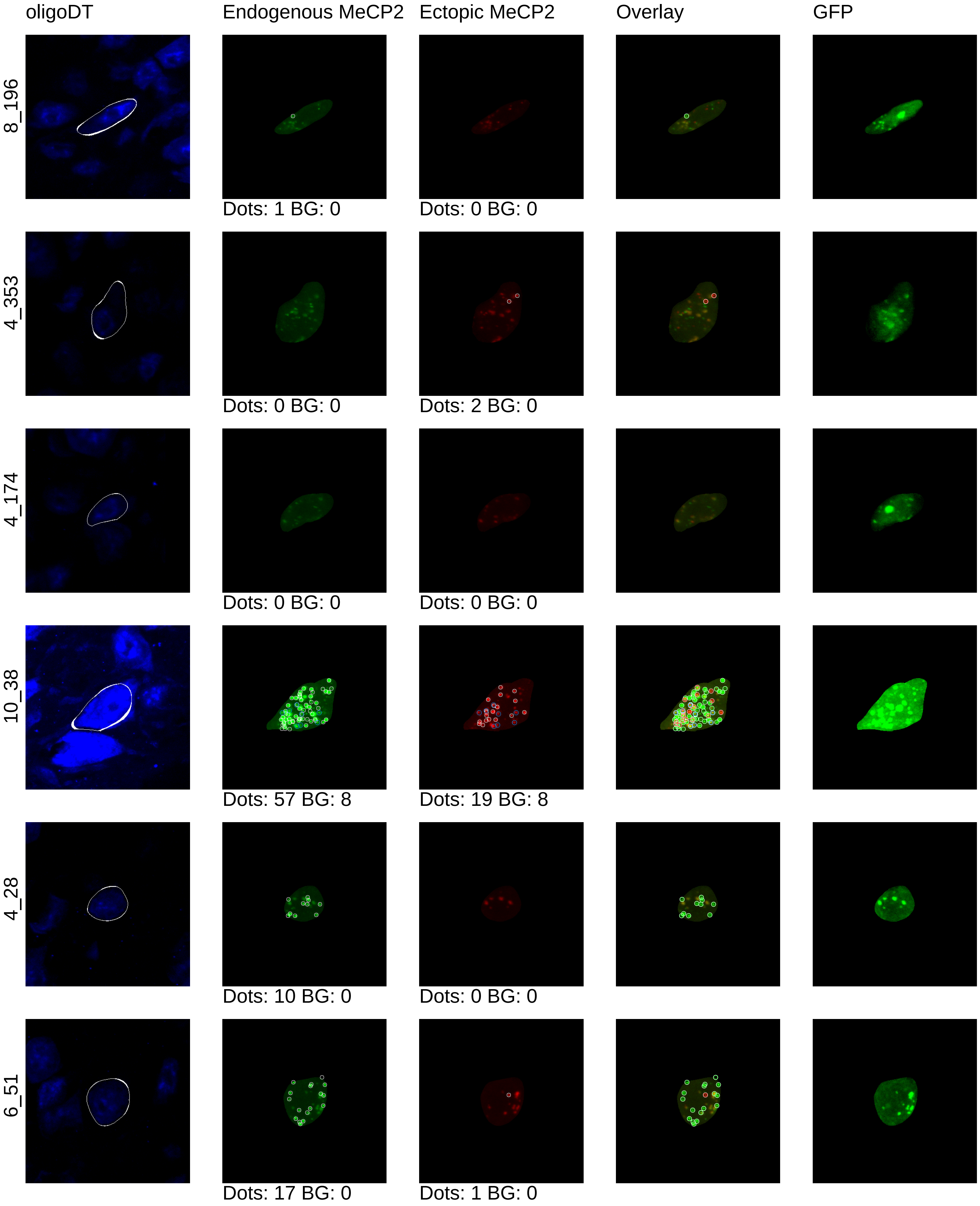

oligoDT
Endogenous MeCP2
Ectopic MeCP2
Overlay
GFP
8_196
Dots: 1 BG: 0
Dots: 0 BG: 0
4_353
Dots: 0 BG: 0
Dots: 2 BG: 0
4_174
Dots: 0 BG: 0
Dots: 0 BG: 0
10_38
Dots: 57 BG: 8
Dots: 19 BG: 8
4_28
Dots: 10 BG: 0
Dots: 0 BG: 0
6_51
Dots: 17 BG: 0
Dots: 1 BG: 0

mouse12_1x

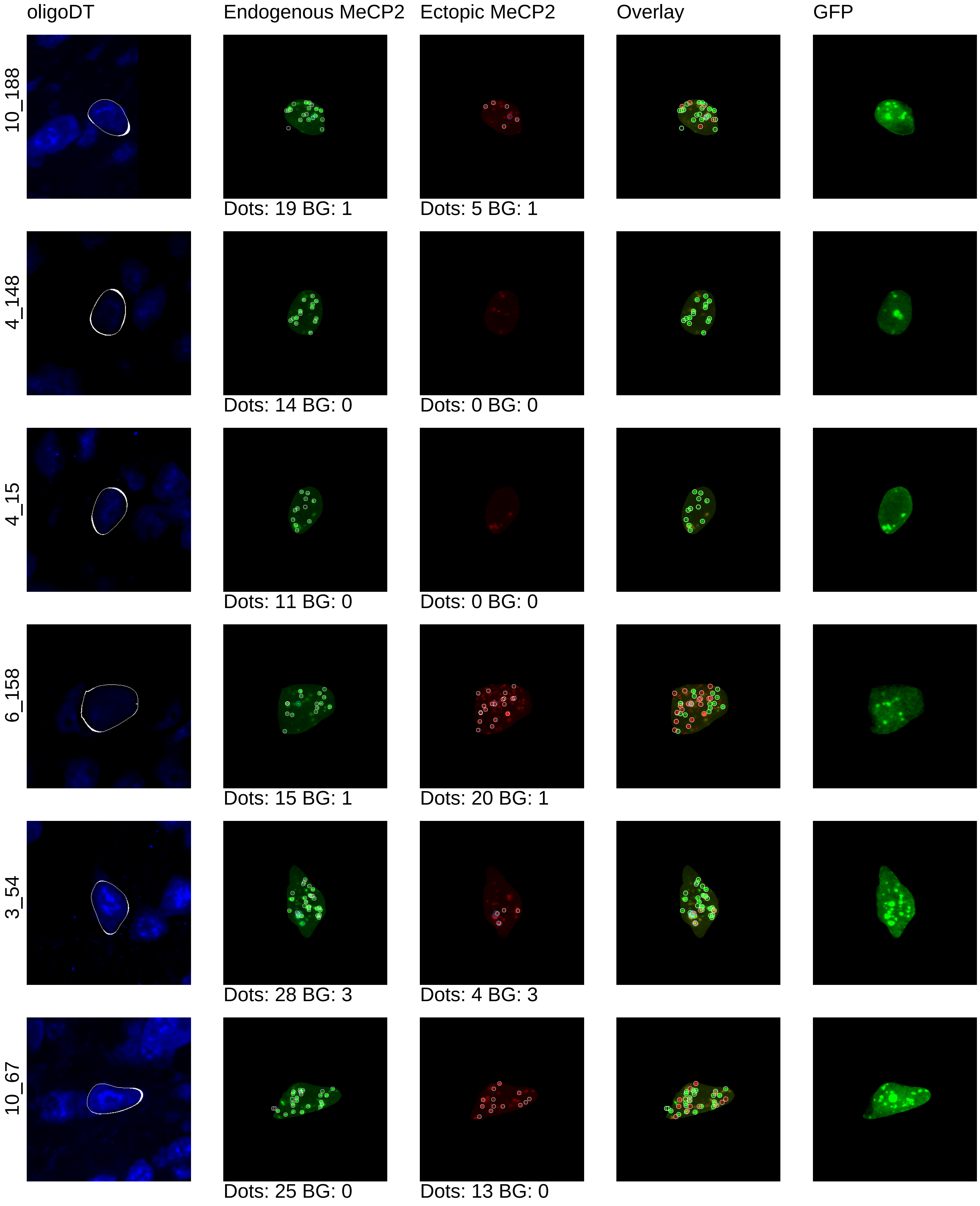
oligoDT
Endogenous MeCP2
Ectopic MeCP2
Overlay
GFP
10_188
Dots: 19 BG: 1
Dots: 5 BG: 1
4_148
Dots: 14 BG: 0
Dots: 0 BG: 0
4_15
Dots: 11 BG: 0
Dots: 0 BG: 0
6_158
Dots: 15 BG: 1
Dots: 20 BG: 1
3_54
Dots: 28 BG: 3
Dots: 4 BG: 3
10_67
Dots: 25 BG: 0
Dots: 13 BG: 0

mouse12_1x

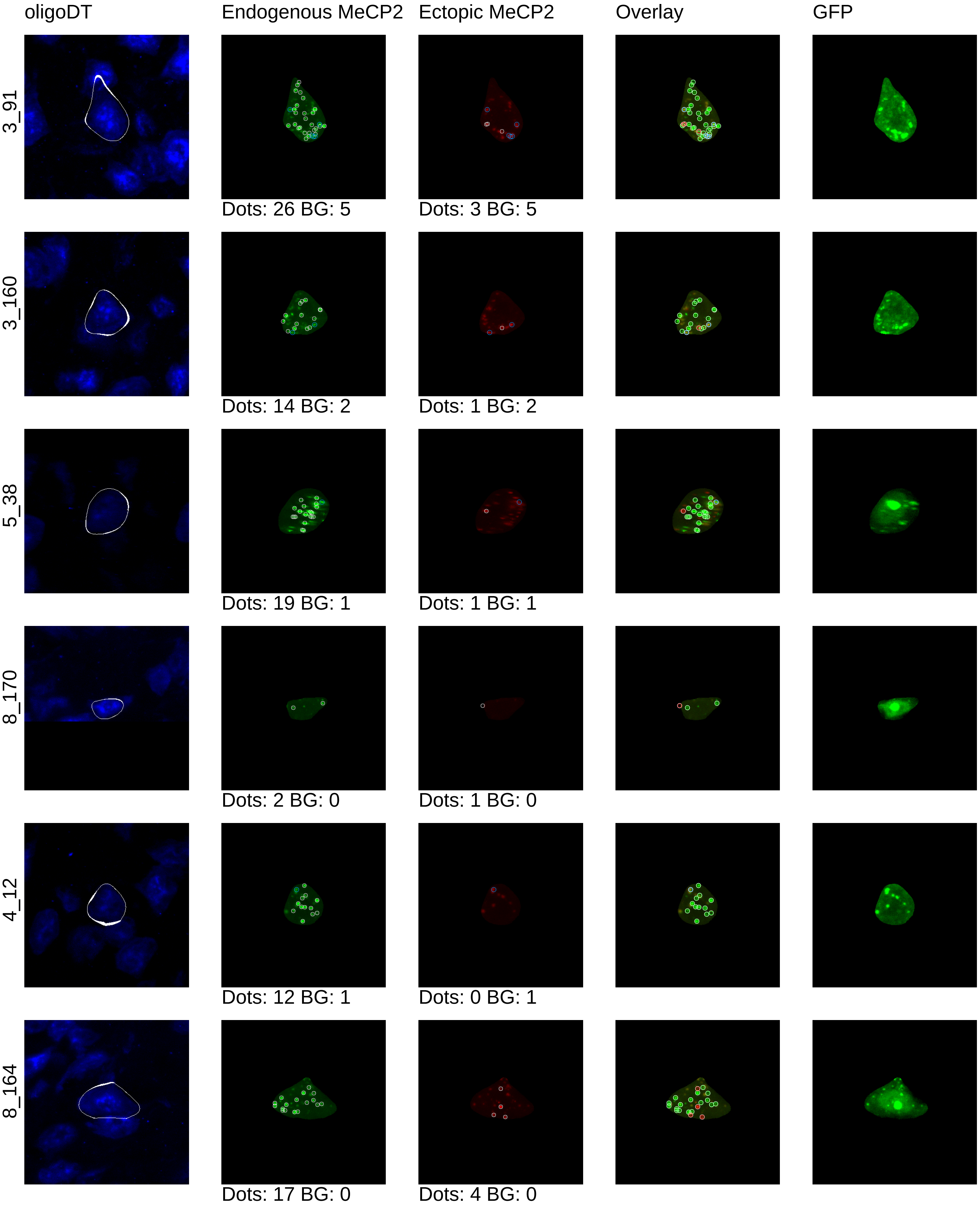
oligoDT
Endogenous MeCP2
Ectopic MeCP2
Overlay
GFP
3_91
Dots: 26 BG: 5
Dots: 3 BG: 5
3_160
Dots: 14 BG: 2
Dots: 1 BG: 2
5_38
Dots: 19 BG: 1
Dots: 1 BG: 1
8_170
Dots: 2 BG: 0
Dots: 1 BG: 0
4_12
Dots: 12 BG: 1
Dots: 0 BG: 1
8_164
Dots: 17 BG: 0
Dots: 4 BG: 0

mouse12_1x

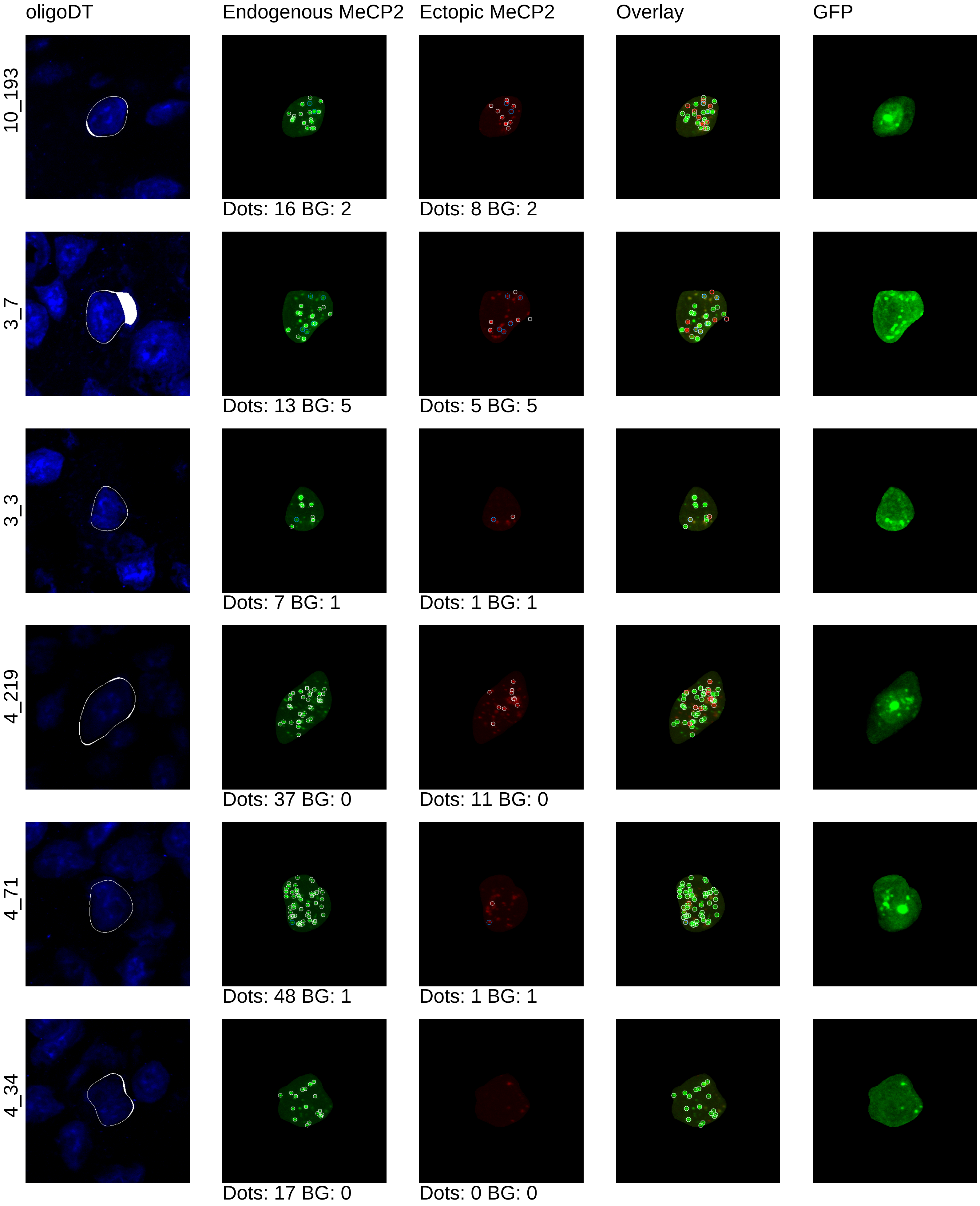
oligoDT
Endogenous MeCP2
Ectopic MeCP2
Overlay
GFP
10_193
Dots: 16 BG: 2
Dots: 8 BG: 2
3_7
Dots: 13 BG: 5
Dots: 5 BG: 5
3_3
Dots: 7 BG: 1
Dots: 1 BG: 1
4_219
Dots: 37 BG: 0
Dots: 11 BG: 0
4_71
Dots: 48 BG: 1
Dots: 1 BG: 1
4_34
Dots: 17 BG: 0
Dots: 0 BG: 0

mouse12_1x

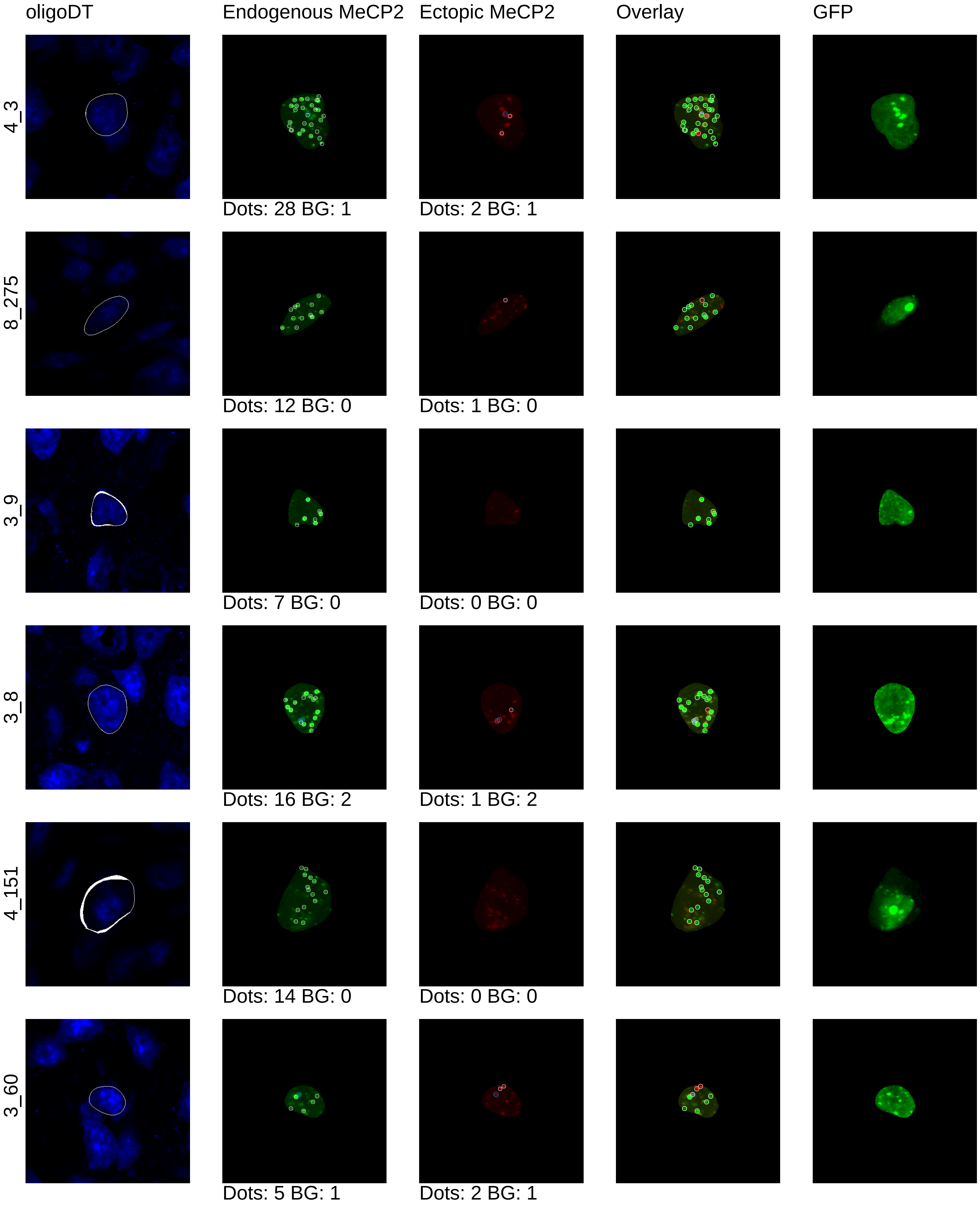

oligoDT
Endogenous MeCP2
Ectopic MeCP2
Overlay
GFP
4_3
Dots: 28 BG: 1
Dots: 2 BG: 1
8_275
Dots: 12 BG: 0
Dots: 1 BG: 0
3_9
Dots: 7 BG: 0
Dots: 0 BG: 0
3_8
Dots: 16 BG: 2
Dots: 1 BG: 2
4_151
Dots: 14 BG: 0
Dots: 0 BG: 0
3_60
Dots: 5 BG: 1
Dots: 2 BG: 1

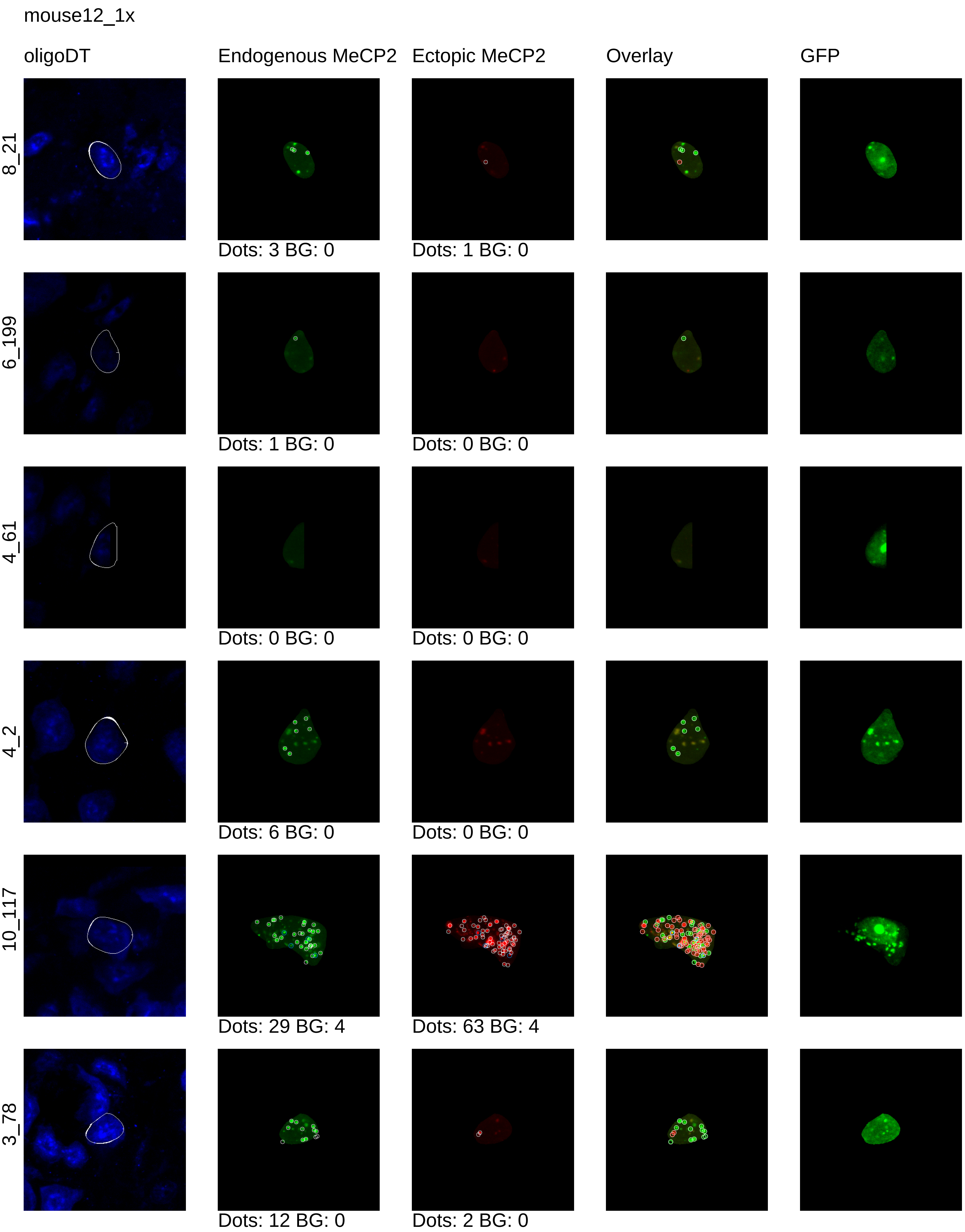
mouse12_1x
oligoDT
Endogenous MeCP2
Ectopic MeCP2
Overlay
GFP
8_21
Dots: 3 BG: 0
Dots: 1 BG: 0
6_199
Dots: 1 BG: 0
Dots: 0 BG: 0
4_61
Dots: 0 BG: 0
Dots: 0 BG: 0
4_2
Dots: 6 BG: 0
Dots: 0 BG: 0
10_117
Dots: 29 BG: 4
Dots: 63 BG: 4
3_78
Dots: 12 BG: 0
Dots: 2 BG: 0

mouse12_1x

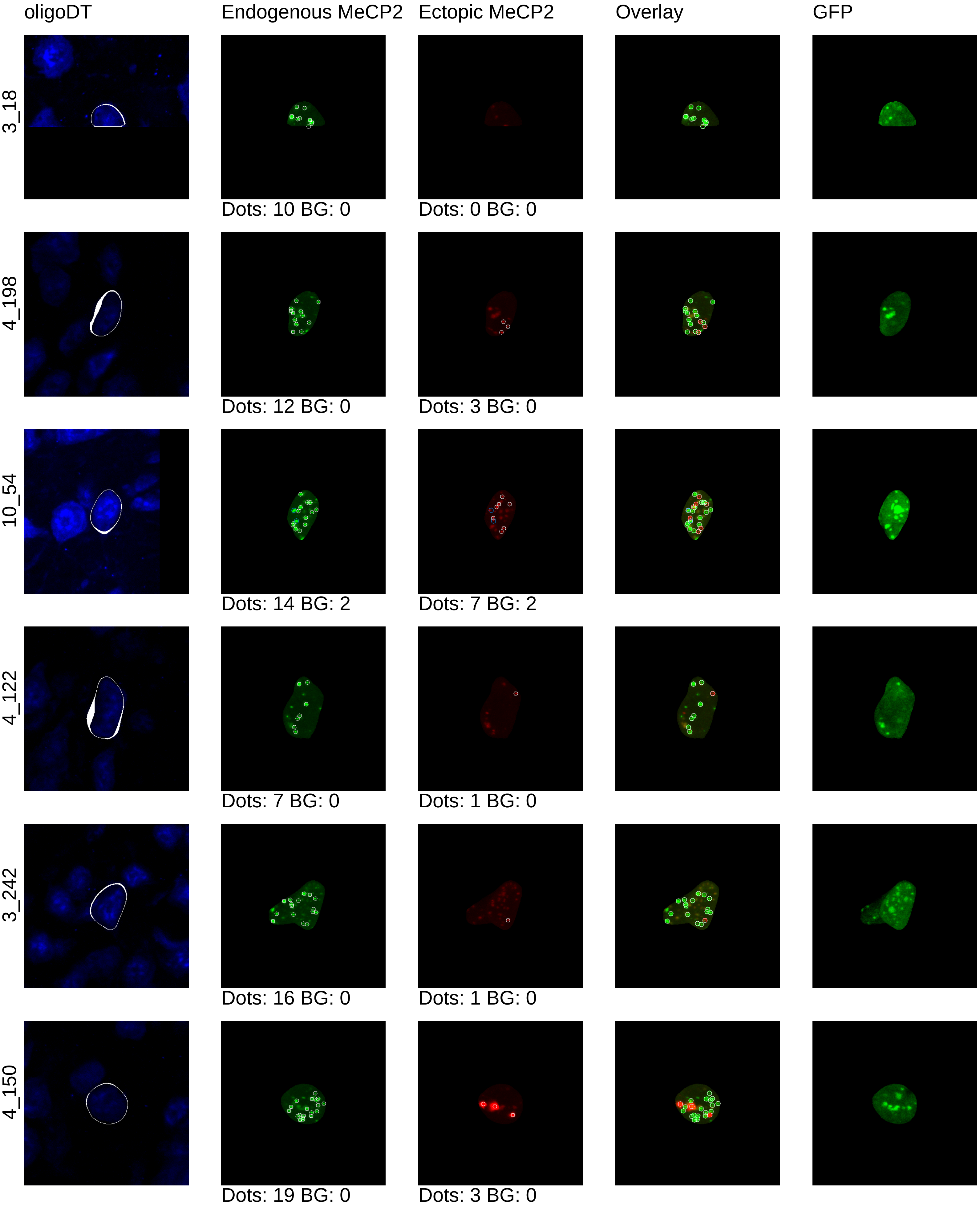
oligoDT
Endogenous MeCP2
Ectopic MeCP2
Overlay
GFP
3_18
Dots: 10 BG: 0
Dots: 0 BG: 0
4_198
Dots: 12 BG: 0
Dots: 3 BG: 0
10_54
Dots: 14 BG: 2
Dots: 7 BG: 2
4_122
Dots: 7 BG: 0
Dots: 1 BG: 0
3_242
Dots: 16 BG: 0
Dots: 1 BG: 0
4_150
Dots: 19 BG: 0
Dots: 3 BG: 0

mouse12_1x

oligoDT | Endogenous MeCP2 | Ectopic MeCP2 | Overlay | GFP

4_144

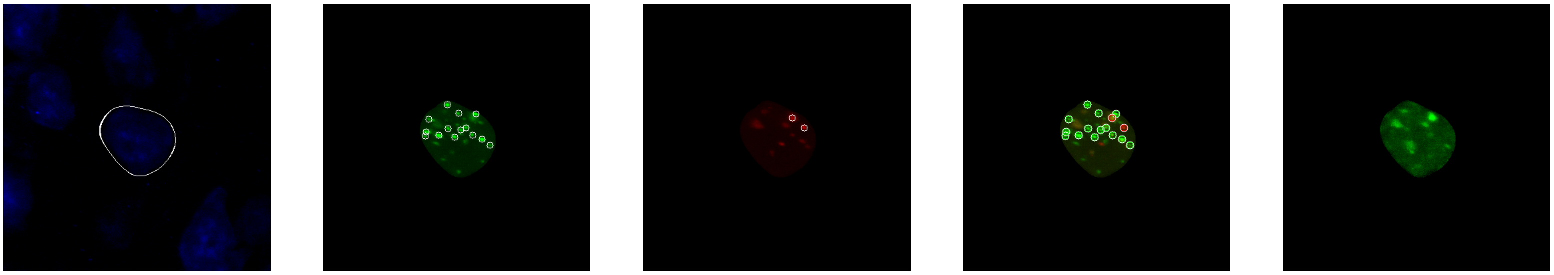

Dots: 14 BG: 0 | Dots: 2 BG: 0

8_117

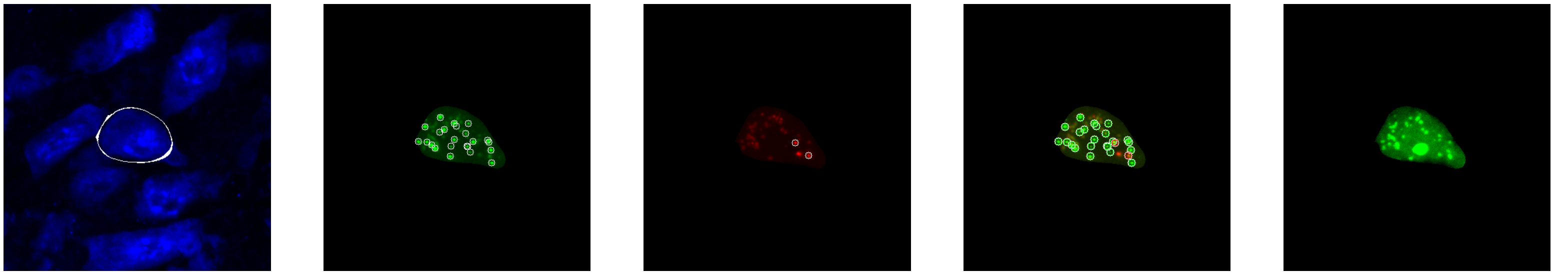

Dots: 23 BG: 0 | Dots: 2 BG: 0
