## Supplementary material for "Synthetic dosage-compensating miRNA circuits allow precision gene therapy for Rett syndrome": 4x_cell_segmentation

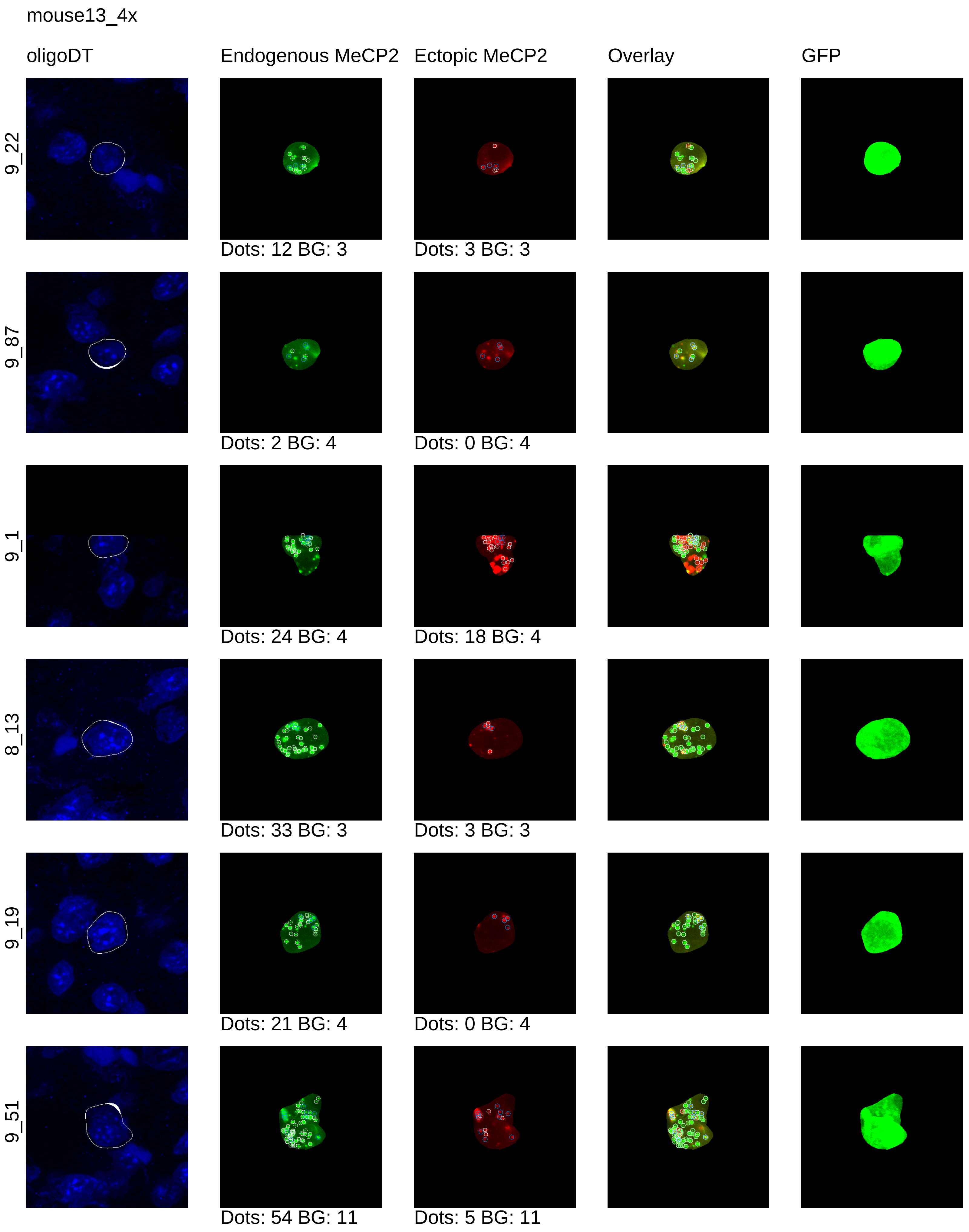
mouse13_4x
oligoDT
Endogenous MeCP2
Ectopic MeCP2
Overlay
GFP
9_22
Dots: 12 BG: 3
Dots: 3 BG: 3
9_87
Dots: 2 BG: 4
Dots: 0 BG: 4
9_1
Dots: 24 BG: 4
Dots: 18 BG: 4
8_13
Dots: 33 BG: 3
Dots: 3 BG: 3
9_19
Dots: 21 BG: 4
Dots: 0 BG: 4
9_51
Dots: 54 BG: 11
Dots: 5 BG: 11

mouse13_4x

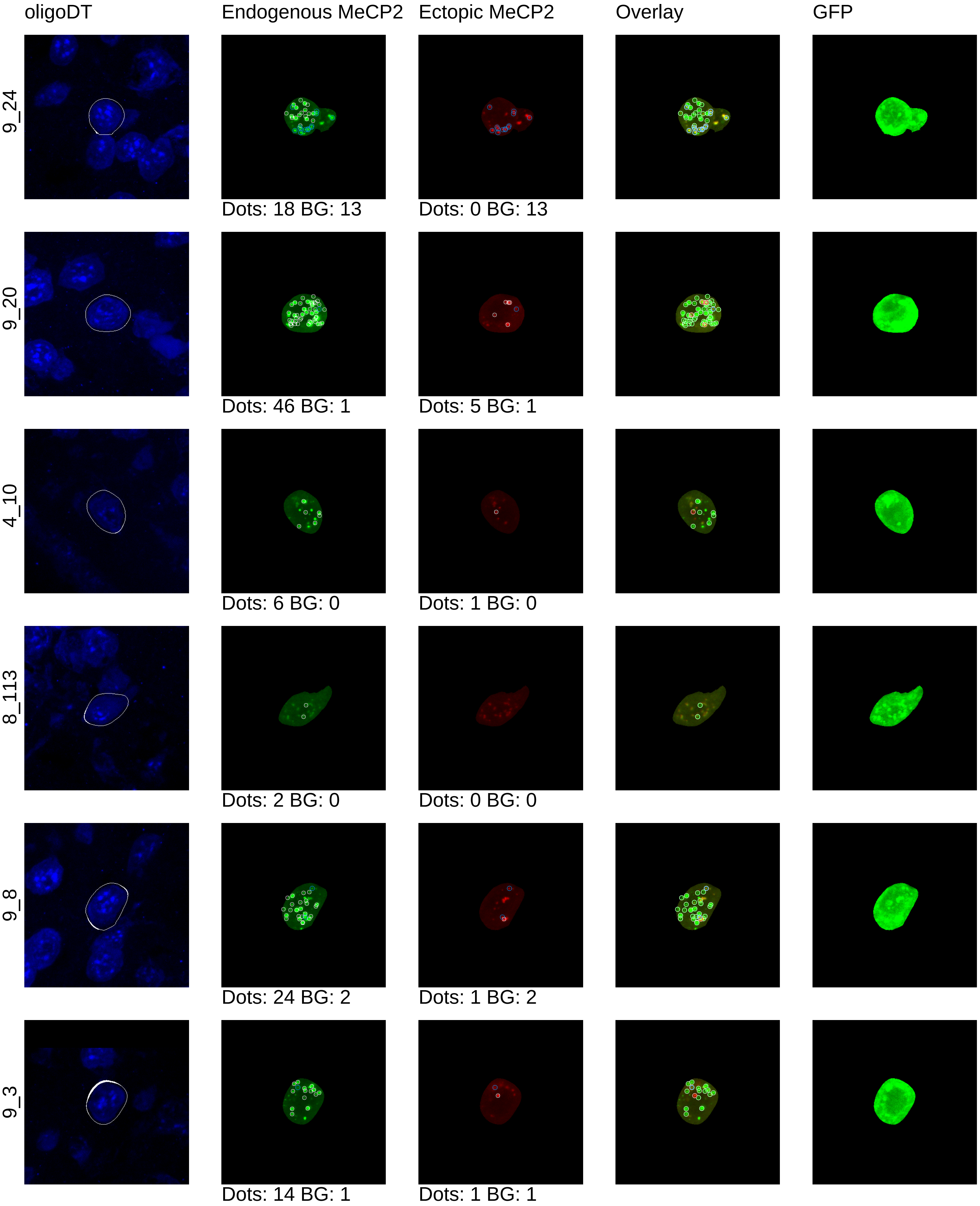

oligoDT
Endogenous MeCP2
Ectopic MeCP2
Overlay
GFP
9_24
Dots: 18 BG: 13
Dots: 0 BG: 13
9_20
Dots: 46 BG: 1
Dots: 5 BG: 1
4_10
Dots: 6 BG: 0
Dots: 1 BG: 0
8_113
Dots: 2 BG: 0
Dots: 0 BG: 0
9_8
Dots: 24 BG: 2
Dots: 1 BG: 2
9_3
Dots: 14 BG: 1
Dots: 1 BG: 1

mouse13_4x

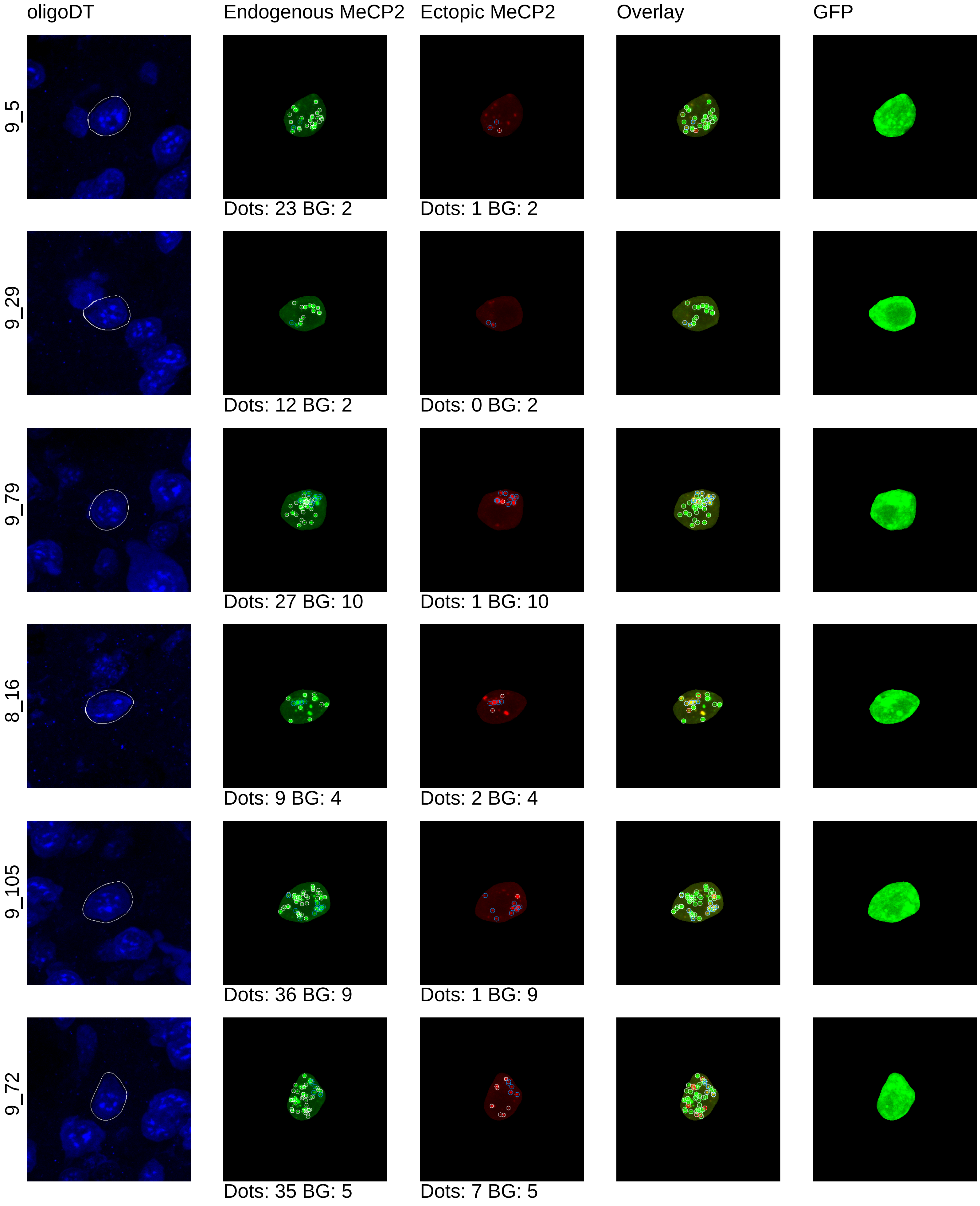
oligoDT
Endogenous MeCP2
Ectopic MeCP2
Overlay
GFP
9_5
Dots: 23 BG: 2
Dots: 1 BG: 2
9_29
Dots: 12 BG: 2
Dots: 0 BG: 2
9_79
Dots: 27 BG: 10
Dots: 1 BG: 10
8_16
Dots: 9 BG: 4
Dots: 2 BG: 4
9_105
Dots: 36 BG: 9
Dots: 1 BG: 9
9_72
Dots: 35 BG: 5
Dots: 7 BG: 5

mouse13_4x

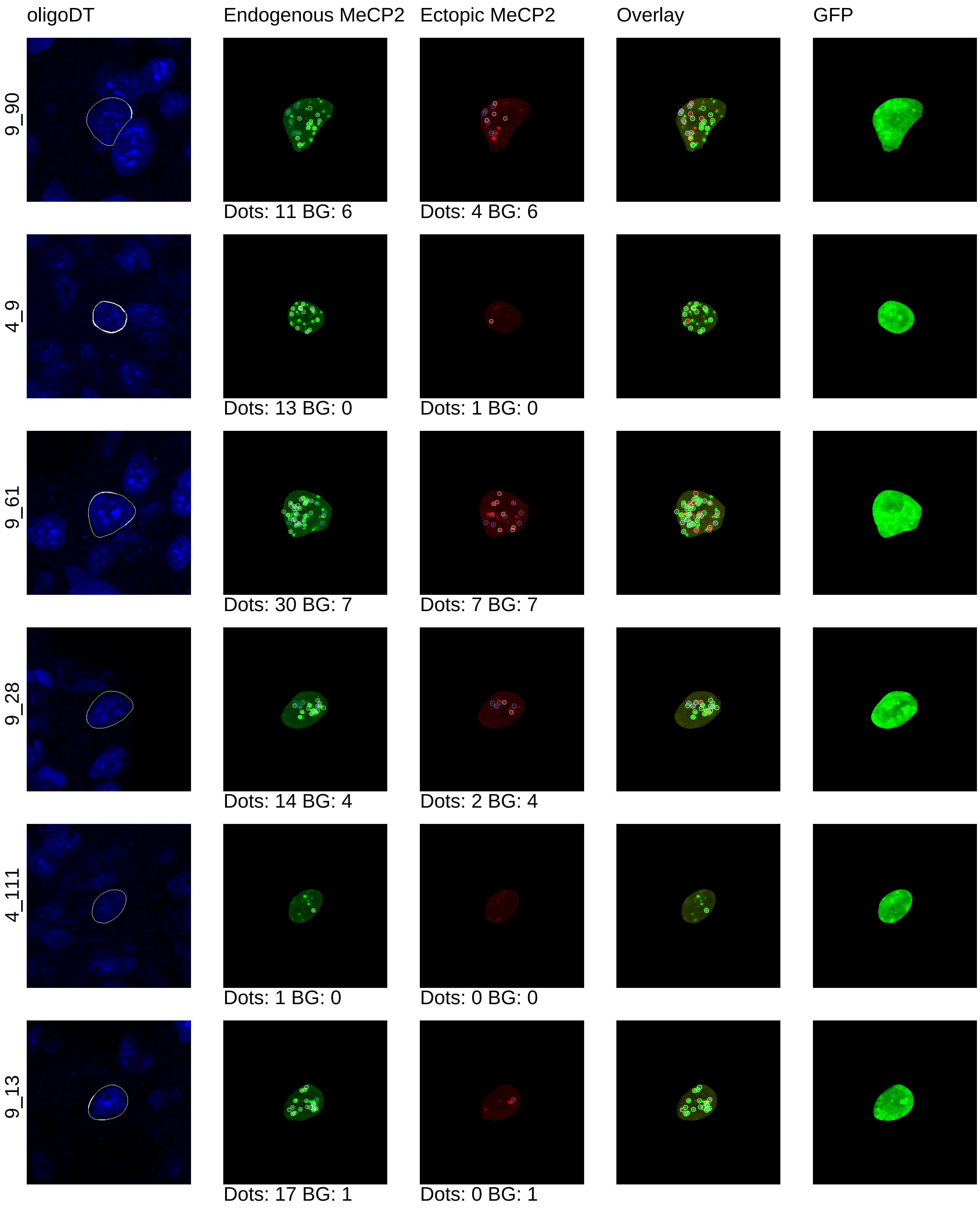
oligoDT
Endogenous MeCP2
Ectopic MeCP2
Overlay
GFP
9_90
Dots: 11 BG: 6
Dots: 4 BG: 6
4_9
Dots: 13 BG: 0
Dots: 1 BG: 0
9_61
Dots: 30 BG: 7
Dots: 7 BG: 7
9_28
Dots: 14 BG: 4
Dots: 2 BG: 4
4_111
Dots: 1 BG: 0
Dots: 0 BG: 0
9_13
Dots: 17 BG: 1
Dots: 0 BG: 1

mouse13_4x

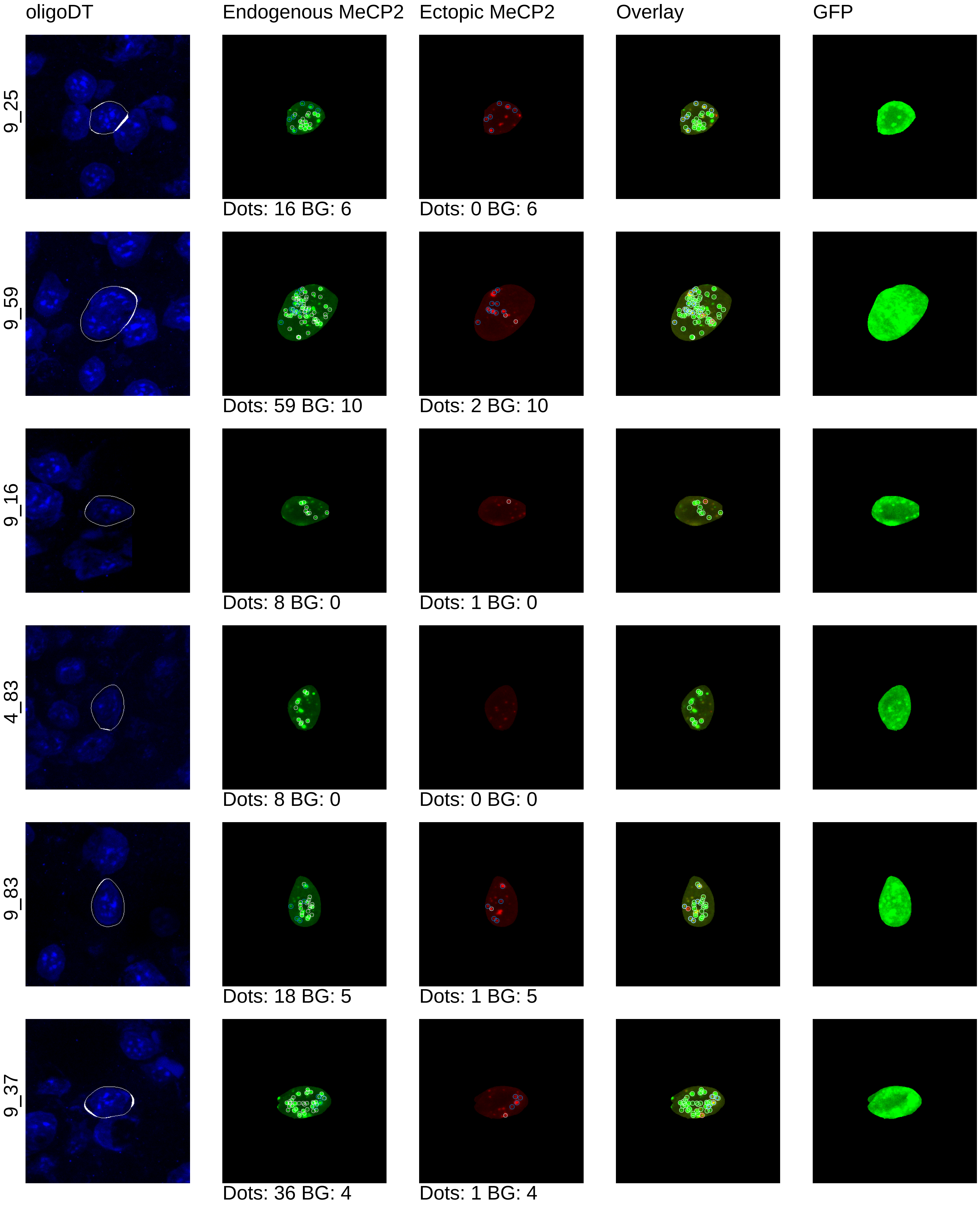
oligoDT
Endogenous MeCP2
Ectopic MeCP2
Overlay
GFP
9_25
Dots: 16 BG: 6
Dots: 0 BG: 6
9_59
Dots: 59 BG: 10
Dots: 2 BG: 10
9_16
Dots: 8 BG: 0
Dots: 1 BG: 0
4_83
Dots: 8 BG: 0
Dots: 0 BG: 0
9_83
Dots: 18 BG: 5
Dots: 1 BG: 5
9_37
Dots: 36 BG: 4
Dots: 1 BG: 4

mouse13_4x

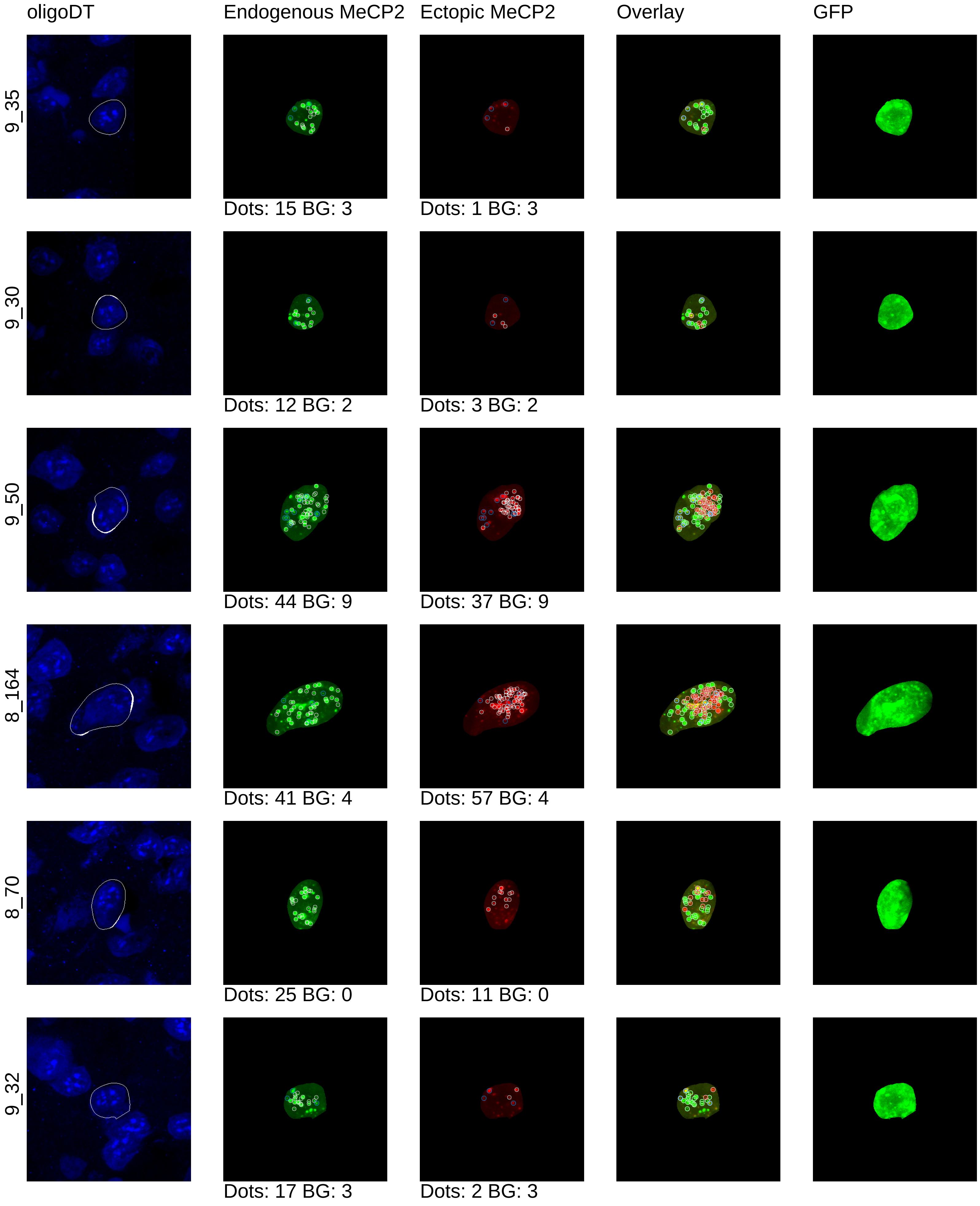
oligoDT
Endogenous MeCP2
Ectopic MeCP2
Overlay
GFP
9_35
Dots: 15 BG: 3
Dots: 1 BG: 3
9_30
Dots: 12 BG: 2
Dots: 3 BG: 2
9_50
Dots: 44 BG: 9
Dots: 37 BG: 9
8_164
Dots: 41 BG: 4
Dots: 57 BG: 4
8_70
Dots: 25 BG: 0
Dots: 11 BG: 0
9_32
Dots: 17 BG: 3
Dots: 2 BG: 3

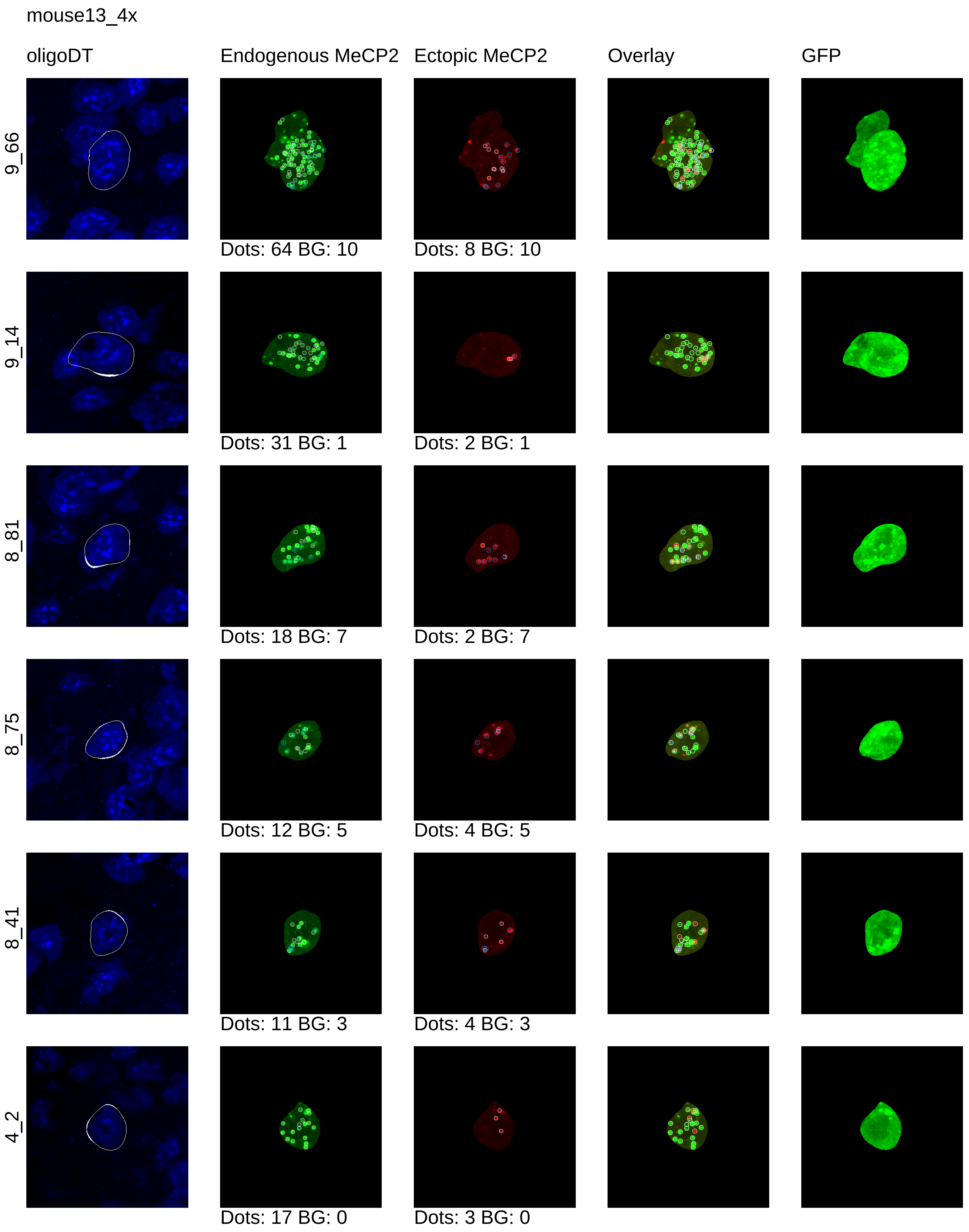
mouse13_4x
oligoDT
Endogenous MeCP2
Ectopic MeCP2
Overlay
GFP
9_66
Dots: 64 BG: 10
Dots: 8 BG: 10
9_14
Dots: 31 BG: 1
Dots: 2 BG: 1
8_81
Dots: 18 BG: 7
Dots: 2 BG: 7
8_75
Dots: 12 BG: 5
Dots: 4 BG: 5
8_41
Dots: 11 BG: 3
Dots: 4 BG: 3
4_2
Dots: 17 BG: 0
Dots: 3 BG: 0

mouse13_4x

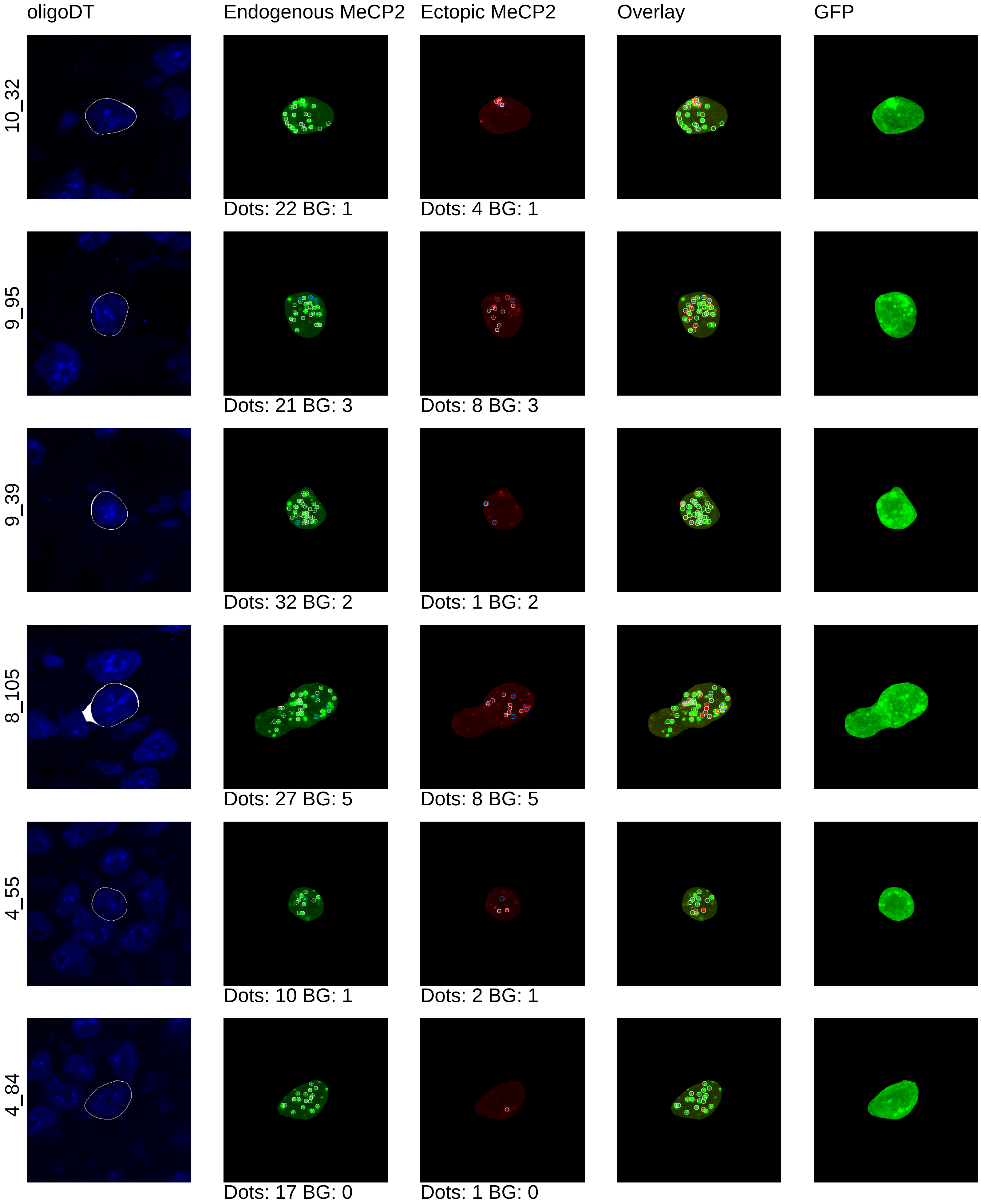
oligoDT
Endogenous MeCP2
Ectopic MeCP2
Overlay
GFP
10_32
Dots: 22 BG: 1
Dots: 4 BG: 1
9_95
Dots: 21 BG: 3
Dots: 8 BG: 3
9_39
Dots: 32 BG: 2
Dots: 1 BG: 2
8_105
Dots: 27 BG: 5
Dots: 8 BG: 5
4_55
Dots: 10 BG: 1
Dots: 2 BG: 1
4_84
Dots: 17 BG: 0
Dots: 1 BG: 0

mouse13_4x

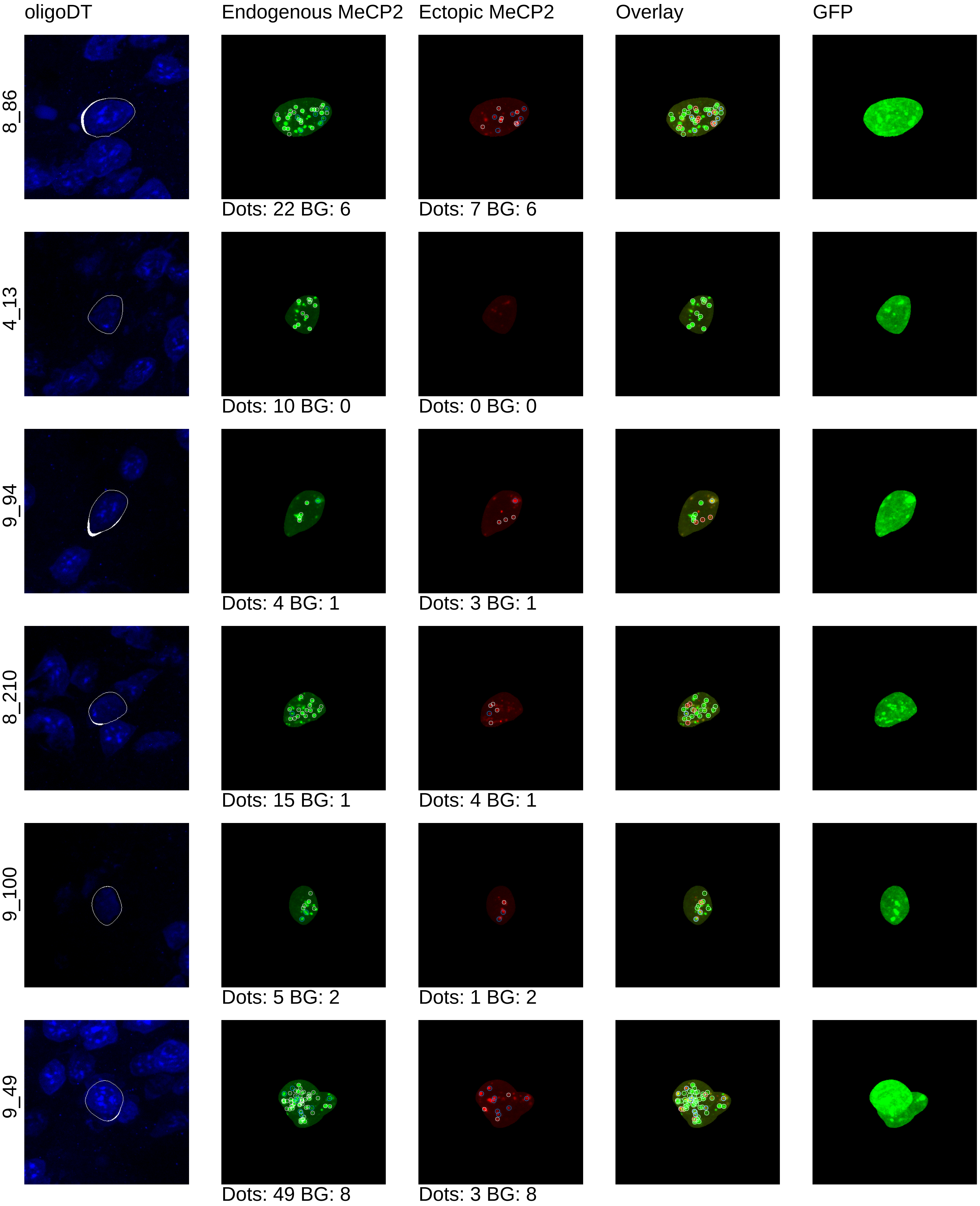
oligoDT
Endogenous MeCP2
Ectopic MeCP2
Overlay
GFP
8_86
Dots: 22 BG: 6
Dots: 7 BG: 6
4_13
Dots: 10 BG: 0
Dots: 0 BG: 0
9_94
Dots: 4 BG: 1
Dots: 3 BG: 1
8_210
Dots: 15 BG: 1
Dots: 4 BG: 1
9_100
Dots: 5 BG: 2
Dots: 1 BG: 2
9_49
Dots: 49 BG: 8
Dots: 3 BG: 8

mouse13_4x

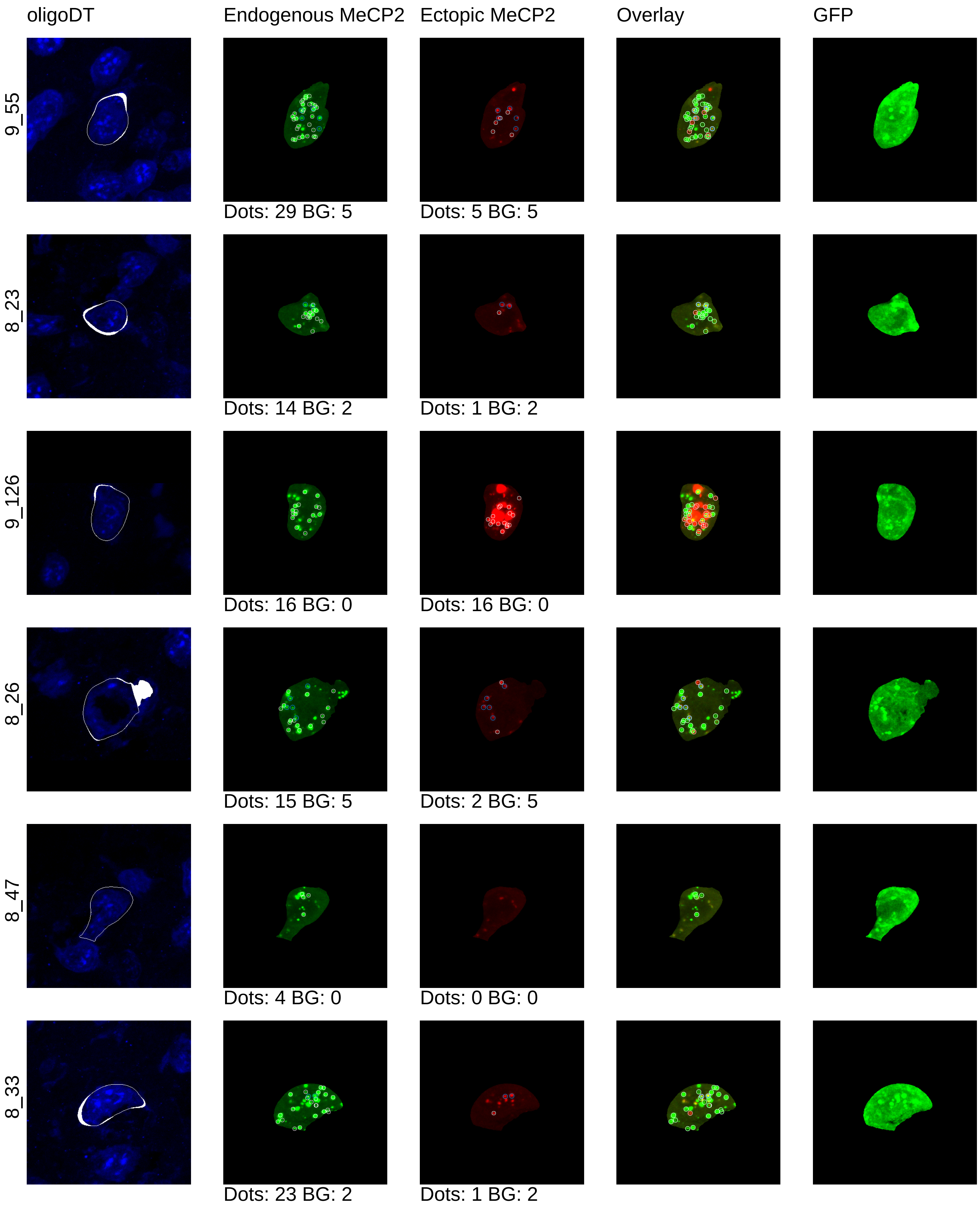

oligoDT
Endogenous MeCP2
Ectopic MeCP2
Overlay
GFP
9_55
Dots: 29 BG: 5
Dots: 5 BG: 5
8_23
Dots: 14 BG: 2
Dots: 1 BG: 2
9_126
Dots: 16 BG: 0
Dots: 16 BG: 0
8_26
Dots: 15 BG: 5
Dots: 2 BG: 5
8_47
Dots: 4 BG: 0
Dots: 0 BG: 0
8_33
Dots: 23 BG: 2
Dots: 1 BG: 2

mouse13_4x

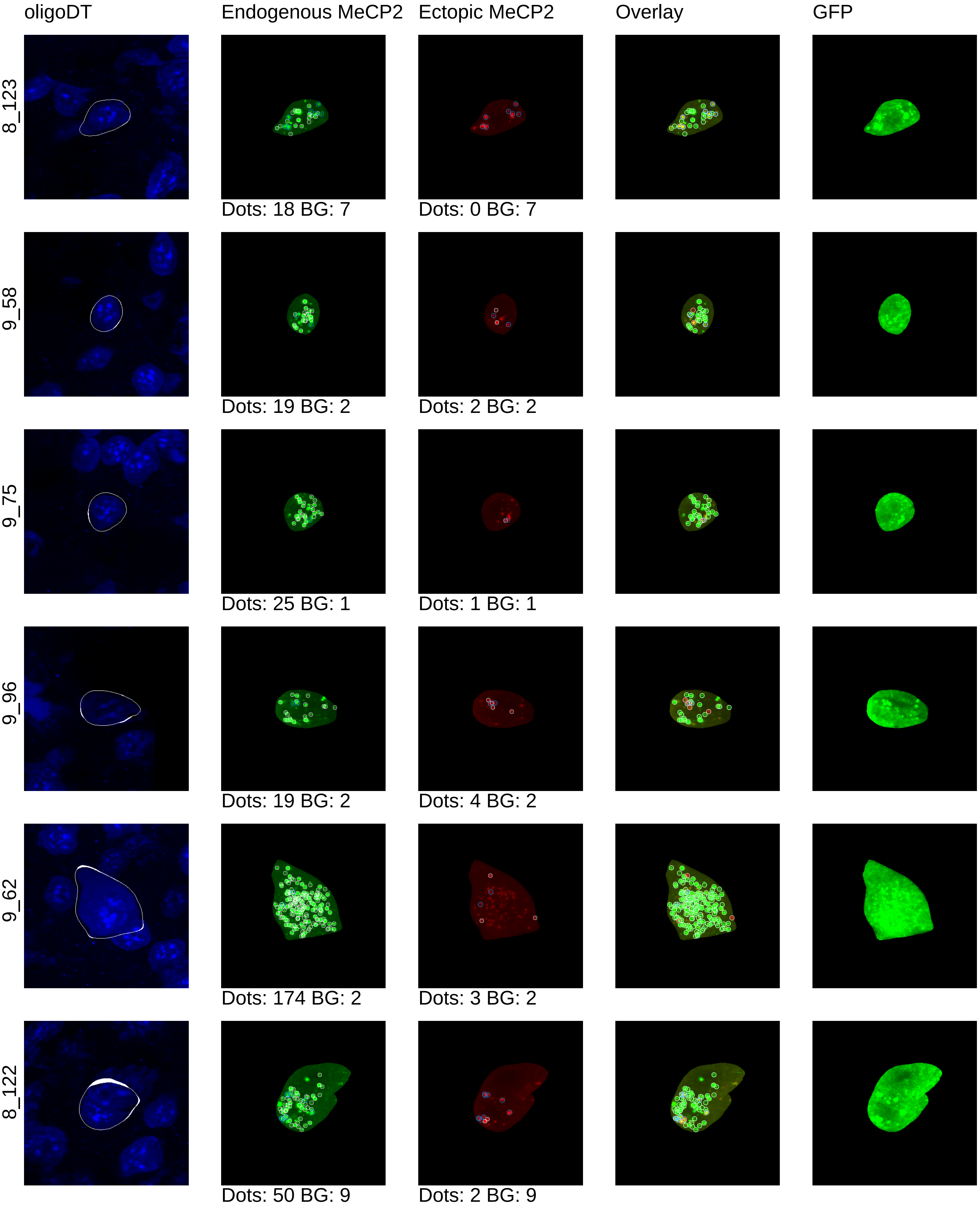
oligoDT
Endogenous MeCP2
Ectopic MeCP2
Overlay
GFP
8_123
Dots: 18 BG: 7
Dots: 0 BG: 7
9_58
Dots: 19 BG: 2
Dots: 2 BG: 2
9_75
Dots: 25 BG: 1
Dots: 1 BG: 1
9_96
Dots: 19 BG: 2
Dots: 4 BG: 2
9_62
Dots: 174 BG: 2
Dots: 3 BG: 2
8_122
Dots: 50 BG: 9
Dots: 2 BG: 9

mouse13_4x

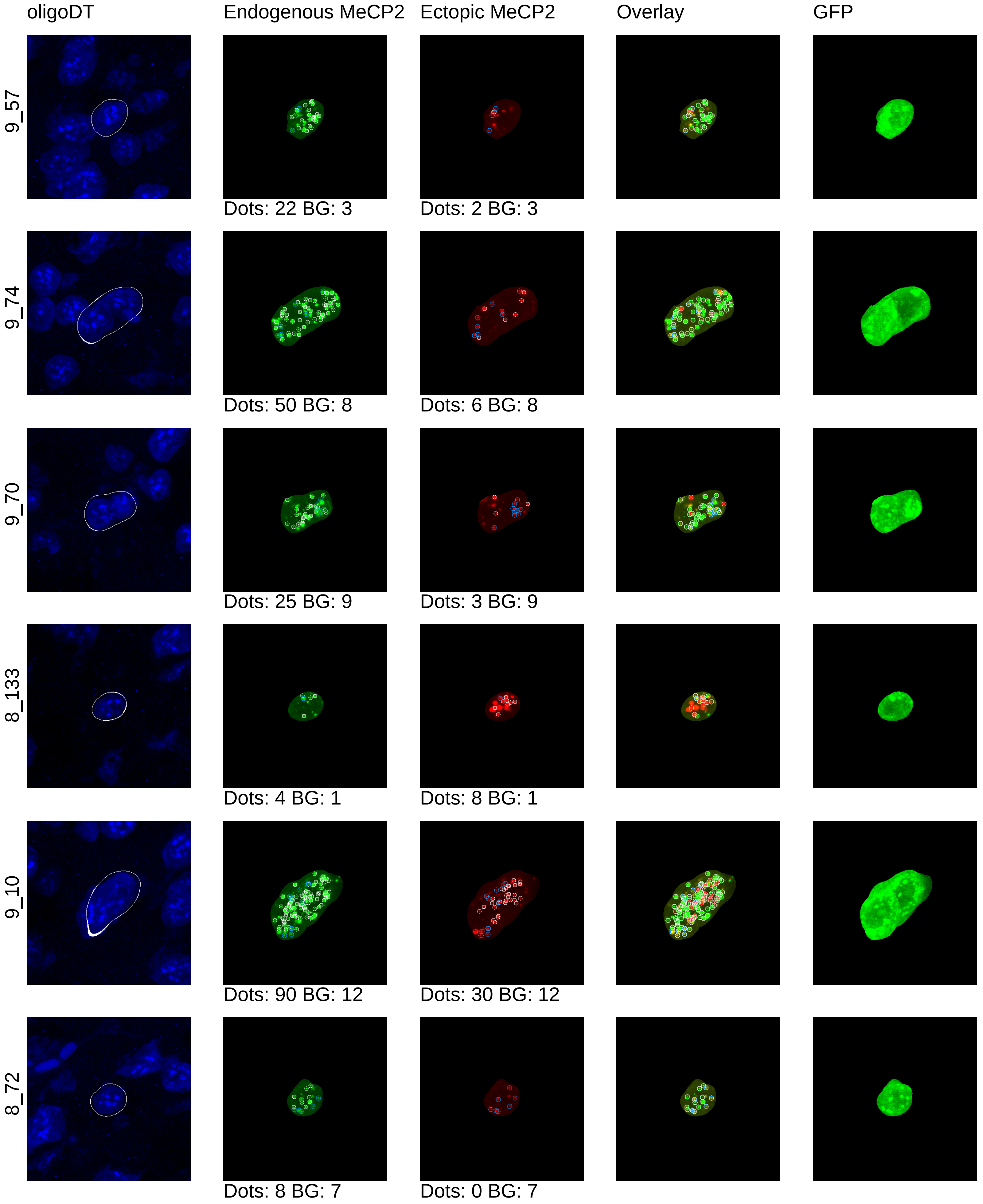

oligoDT
Endogenous MeCP2
Ectopic MeCP2
Overlay
GFP
9_57
Dots: 22 BG: 3
Dots: 2 BG: 3
9_74
Dots: 50 BG: 8
Dots: 6 BG: 8
9_70
Dots: 25 BG: 9
Dots: 3 BG: 9
8_133
Dots: 4 BG: 1
Dots: 8 BG: 1
9_10
Dots: 90 BG: 12
Dots: 30 BG: 12
8_72
Dots: 8 BG: 7
Dots: 0 BG: 7

mouse13_4x

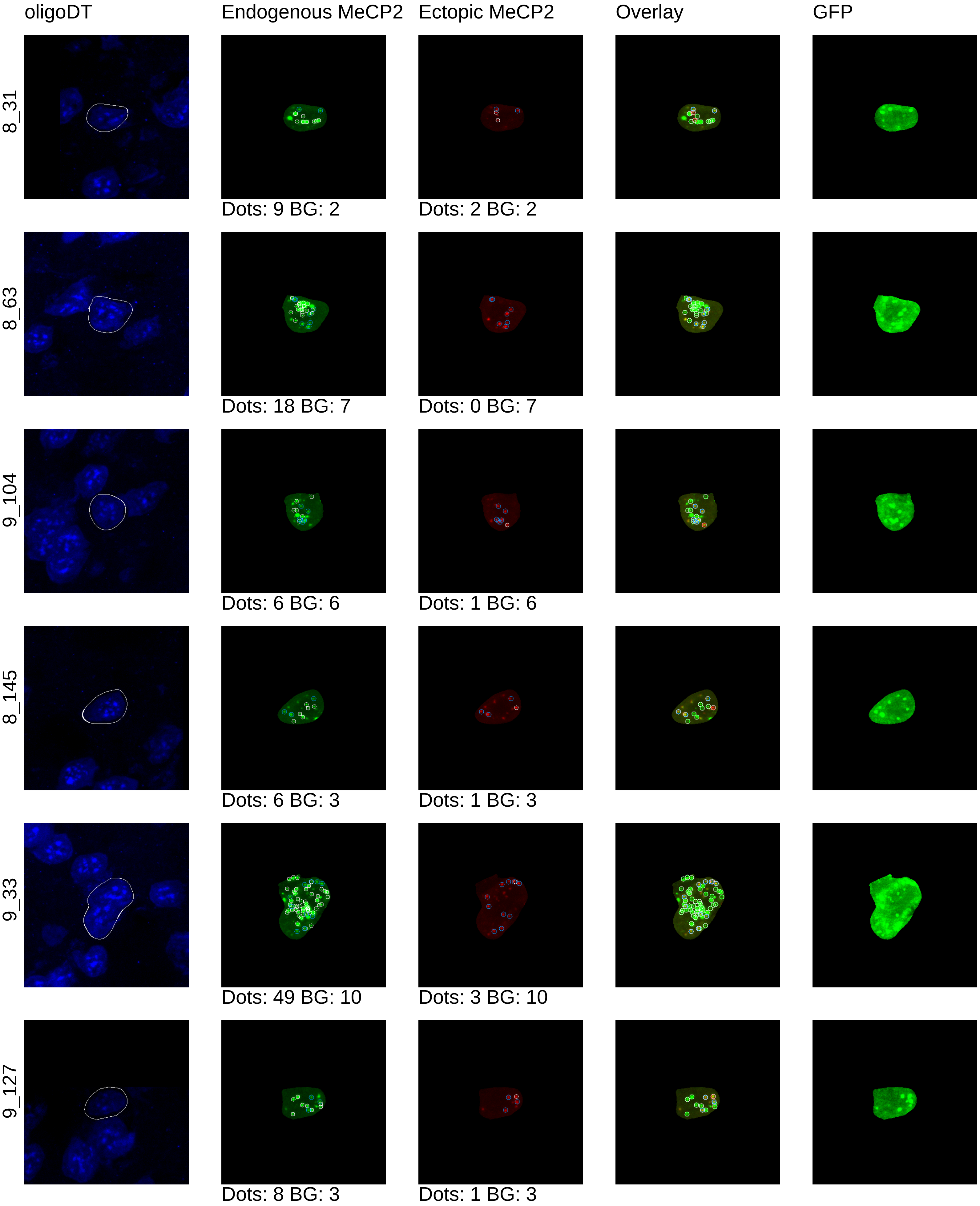

oligoDT
Endogenous MeCP2
Ectopic MeCP2
Overlay
GFP
8_31
Dots: 9 BG: 2
Dots: 2 BG: 2
8_63
Dots: 18 BG: 7
Dots: 0 BG: 7
9_104
Dots: 6 BG: 6
Dots: 1 BG: 6
8_145
Dots: 6 BG: 3
Dots: 1 BG: 3
9_33
Dots: 49 BG: 10
Dots: 3 BG: 10
9_127
Dots: 8 BG: 3
Dots: 1 BG: 3

mouse13_4x

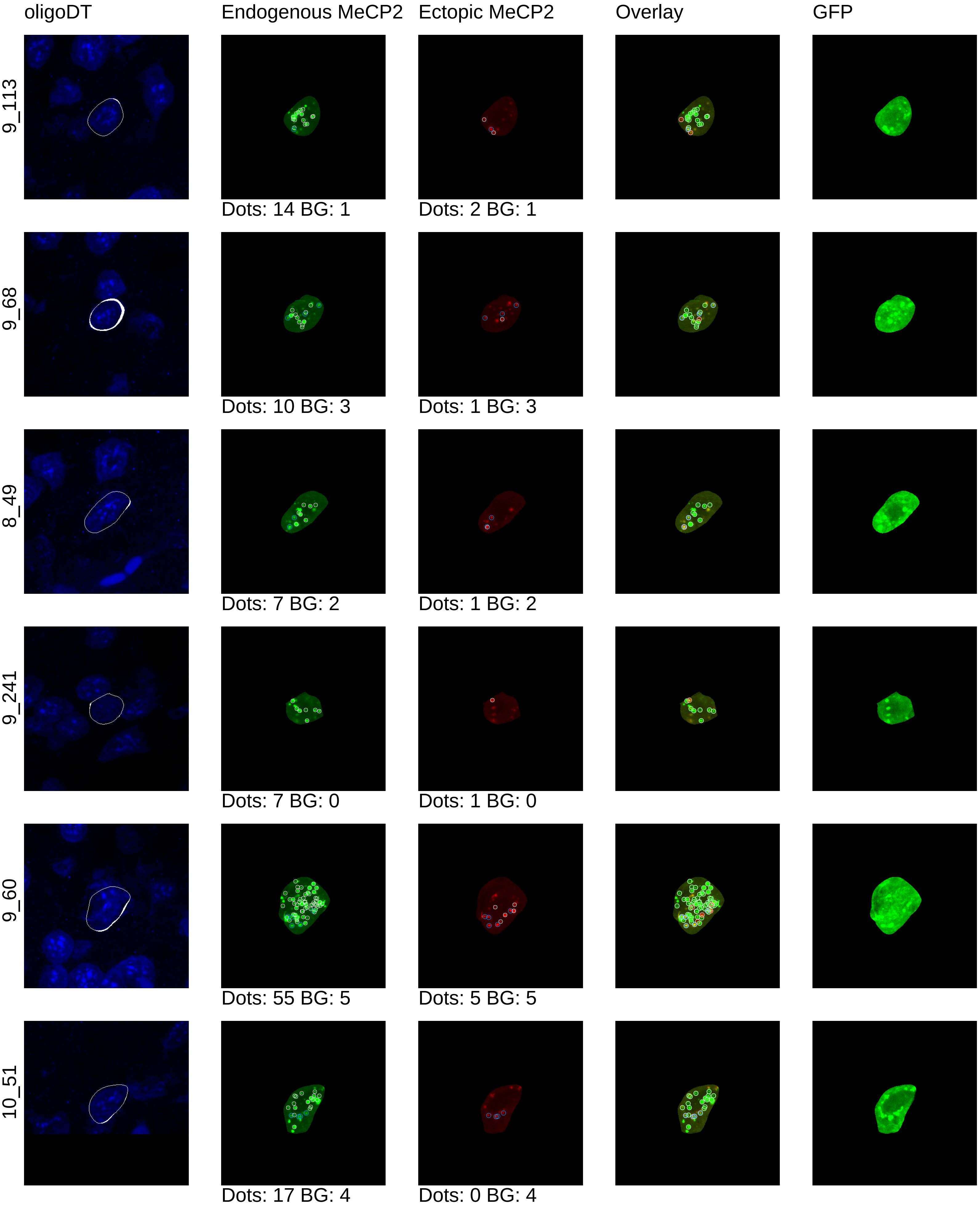
oligoDT
Endogenous MeCP2
Ectopic MeCP2
Overlay
GFP
9_113
Dots: 14 BG: 1
Dots: 2 BG: 1
9_68
Dots: 10 BG: 3
Dots: 1 BG: 3
8_49
Dots: 7 BG: 2
Dots: 1 BG: 2
9_241
Dots: 7 BG: 0
Dots: 1 BG: 0
9_60
Dots: 55 BG: 5
Dots: 5 BG: 5
10_51
Dots: 17 BG: 4
Dots: 0 BG: 4

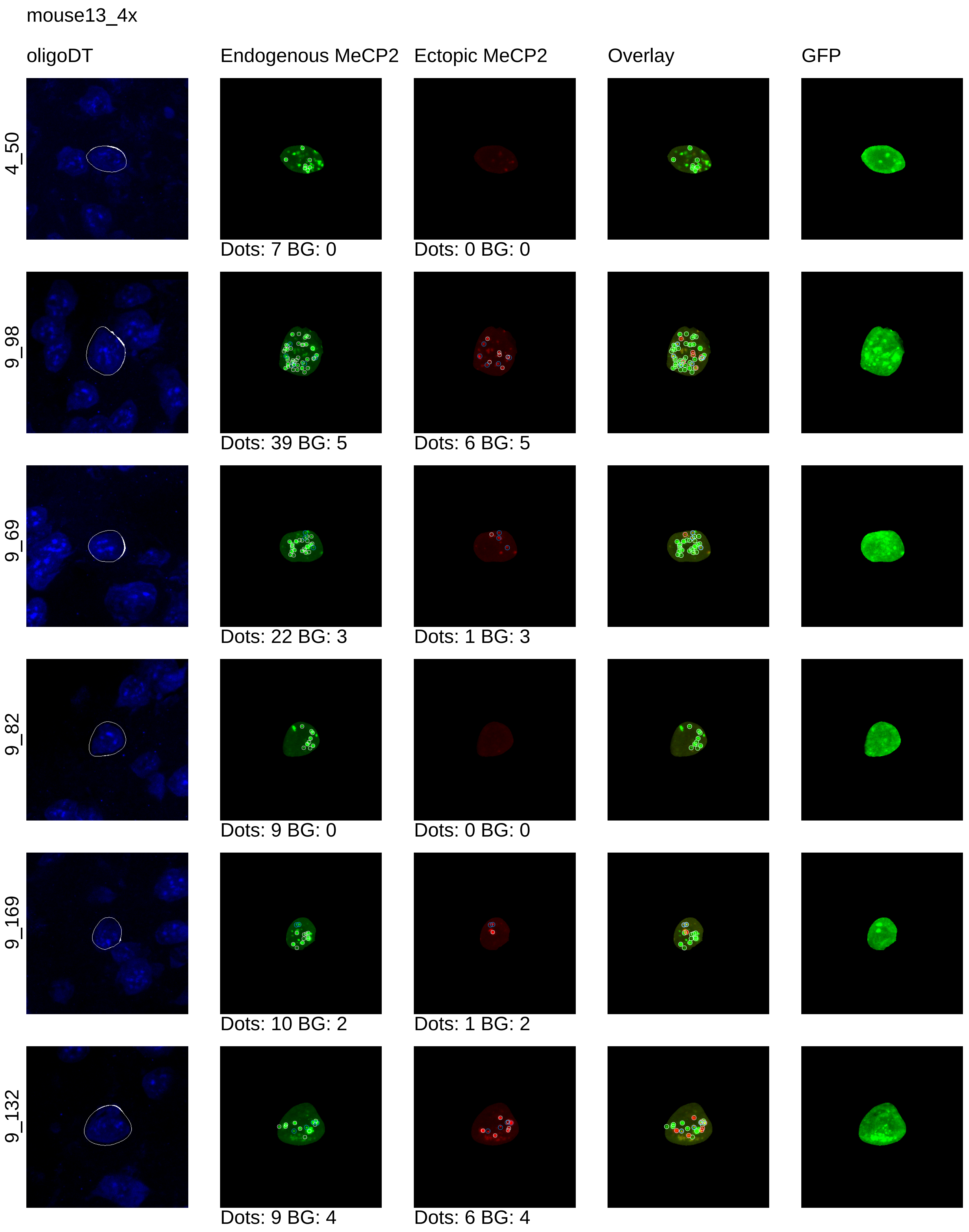
mouse13_4x
oligoDT
Endogenous MeCP2
Ectopic MeCP2
Overlay
GFP
4_50
Dots: 7 BG: 0
Dots: 0 BG: 0
9_98
Dots: 39 BG: 5
Dots: 6 BG: 5
9_69
Dots: 22 BG: 3
Dots: 1 BG: 3
9_82
Dots: 9 BG: 0
Dots: 0 BG: 0
9_169
Dots: 10 BG: 2
Dots: 1 BG: 2
9_132
Dots: 9 BG: 4
Dots: 6 BG: 4

mouse13_4x

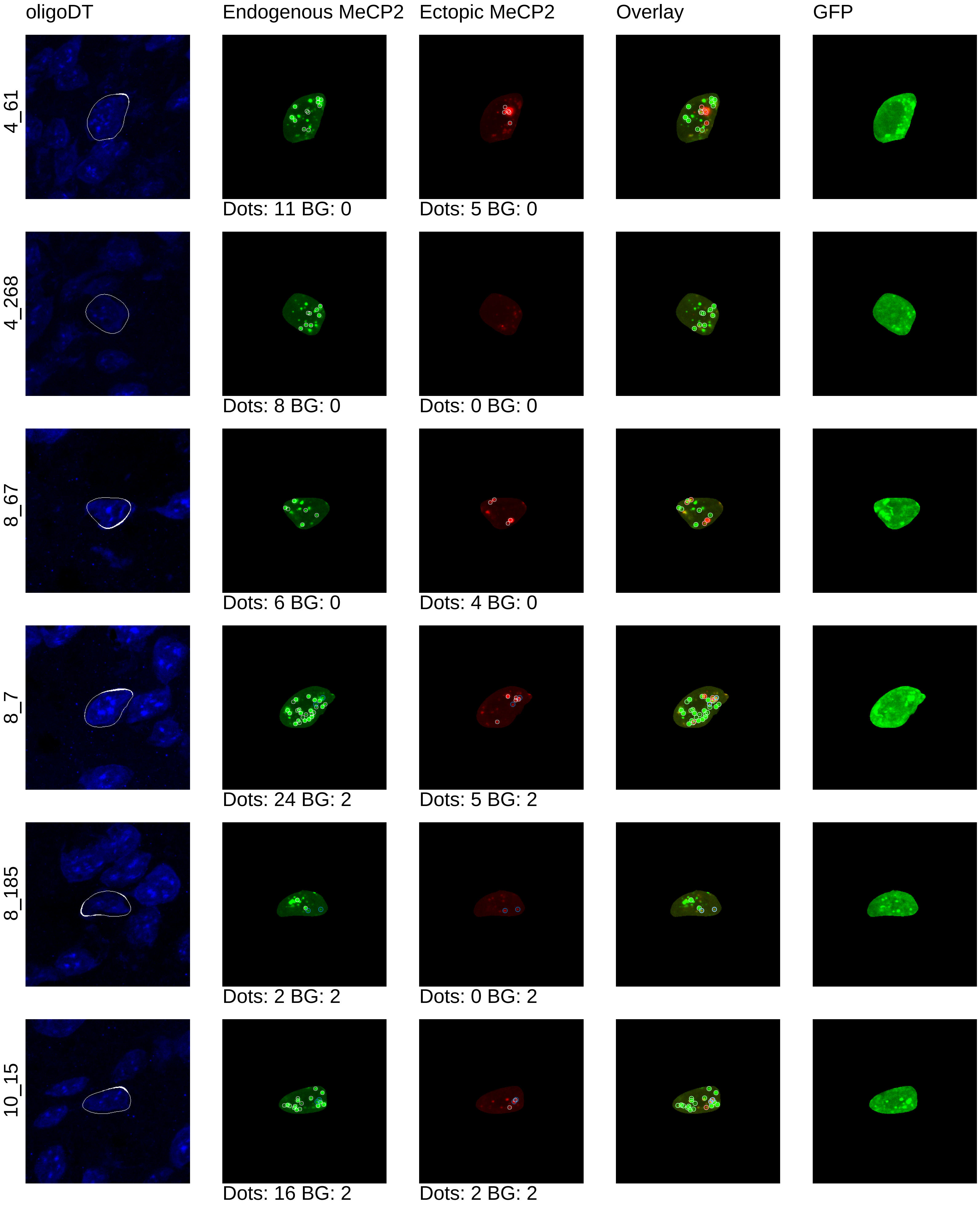
oligoDT
Endogenous MeCP2
Ectopic MeCP2
Overlay
GFP
4_61
Dots: 11 BG: 0
Dots: 5 BG: 0
4_268
Dots: 8 BG: 0
Dots: 0 BG: 0
8_67
Dots: 6 BG: 0
Dots: 4 BG: 0
8_7
Dots: 24 BG: 2
Dots: 5 BG: 2
8_185
Dots: 2 BG: 2
Dots: 0 BG: 2
10_15
Dots: 16 BG: 2
Dots: 2 BG: 2

mouse13_4x

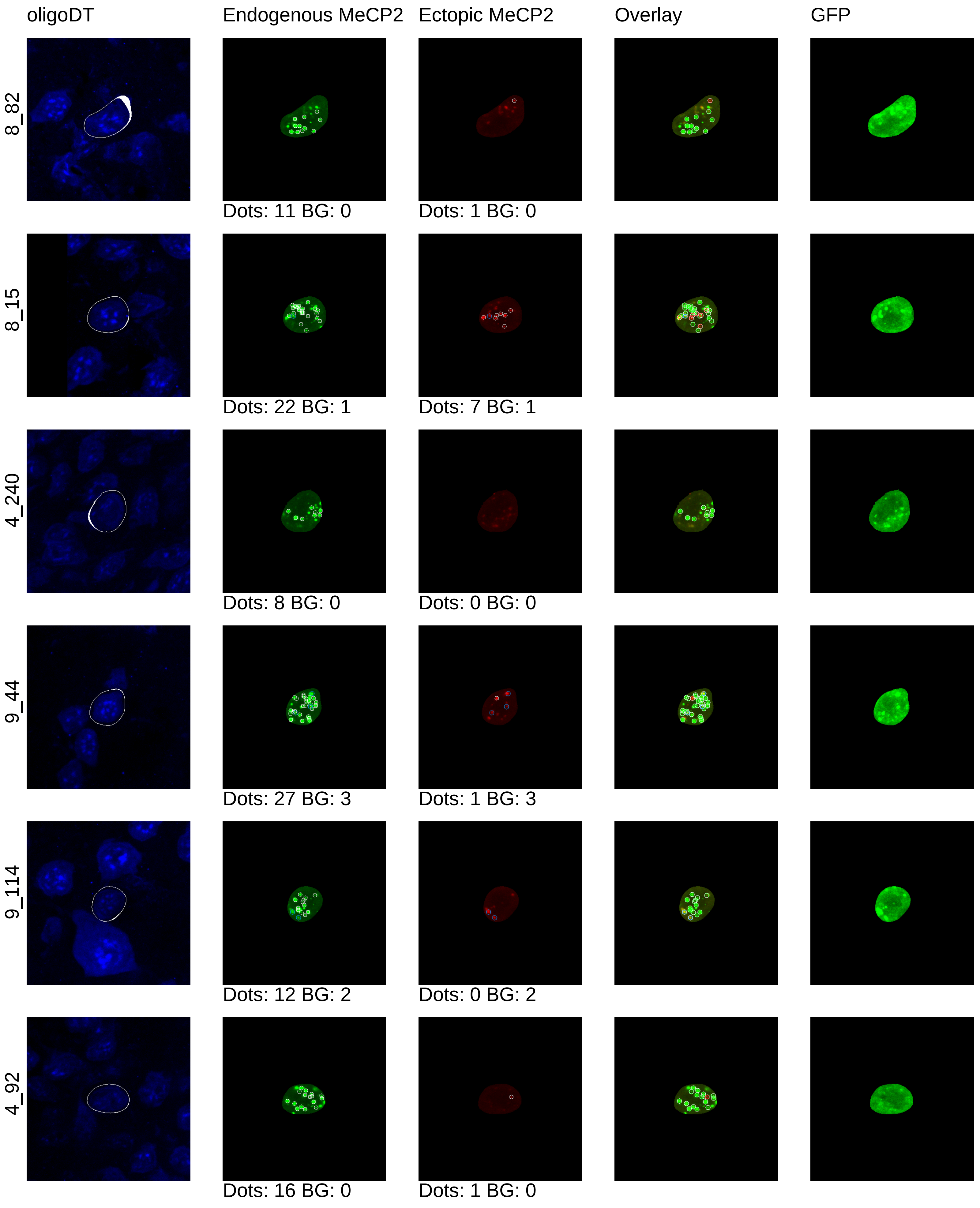
oligoDT
Endogenous MeCP2
Ectopic MeCP2
Overlay
GFP
8_82
Dots: 11 BG: 0
Dots: 1 BG: 0
8_15
Dots: 22 BG: 1
Dots: 7 BG: 1
4_240
Dots: 8 BG: 0
Dots: 0 BG: 0
9_44
Dots: 27 BG: 3
Dots: 1 BG: 3
9_114
Dots: 12 BG: 2
Dots: 0 BG: 2
4_92
Dots: 16 BG: 0
Dots: 1 BG: 0

mouse13_4x

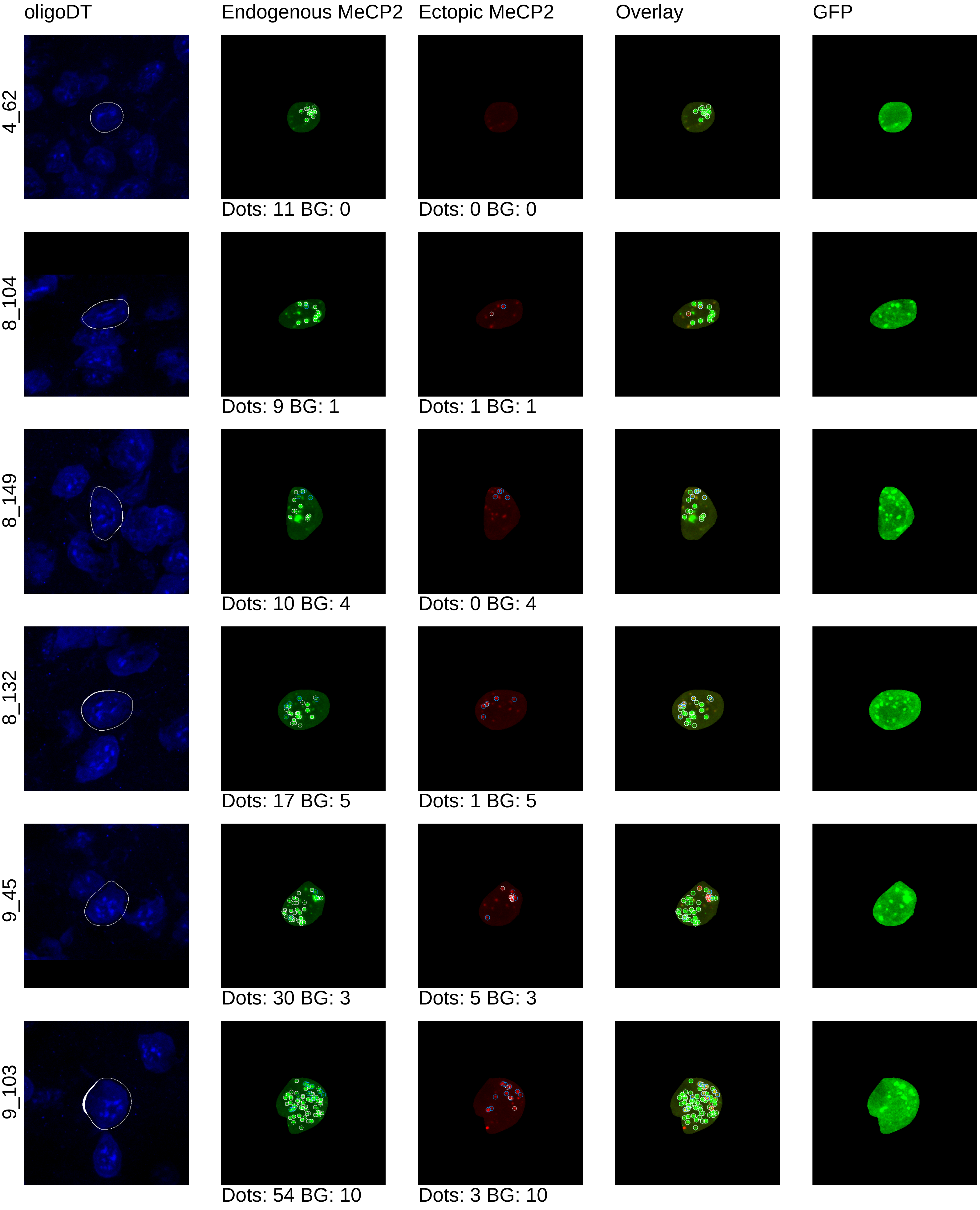
oligoDT
Endogenous MeCP2
Ectopic MeCP2
Overlay
GFP
4_62
Dots: 11 BG: 0
Dots: 0 BG: 0
8_104
Dots: 9 BG: 1
Dots: 1 BG: 1
8_149
Dots: 10 BG: 4
Dots: 0 BG: 4
8_132
Dots: 17 BG: 5
Dots: 1 BG: 5
9_45
Dots: 30 BG: 3
Dots: 5 BG: 3
9_103
Dots: 54 BG: 10
Dots: 3 BG: 10

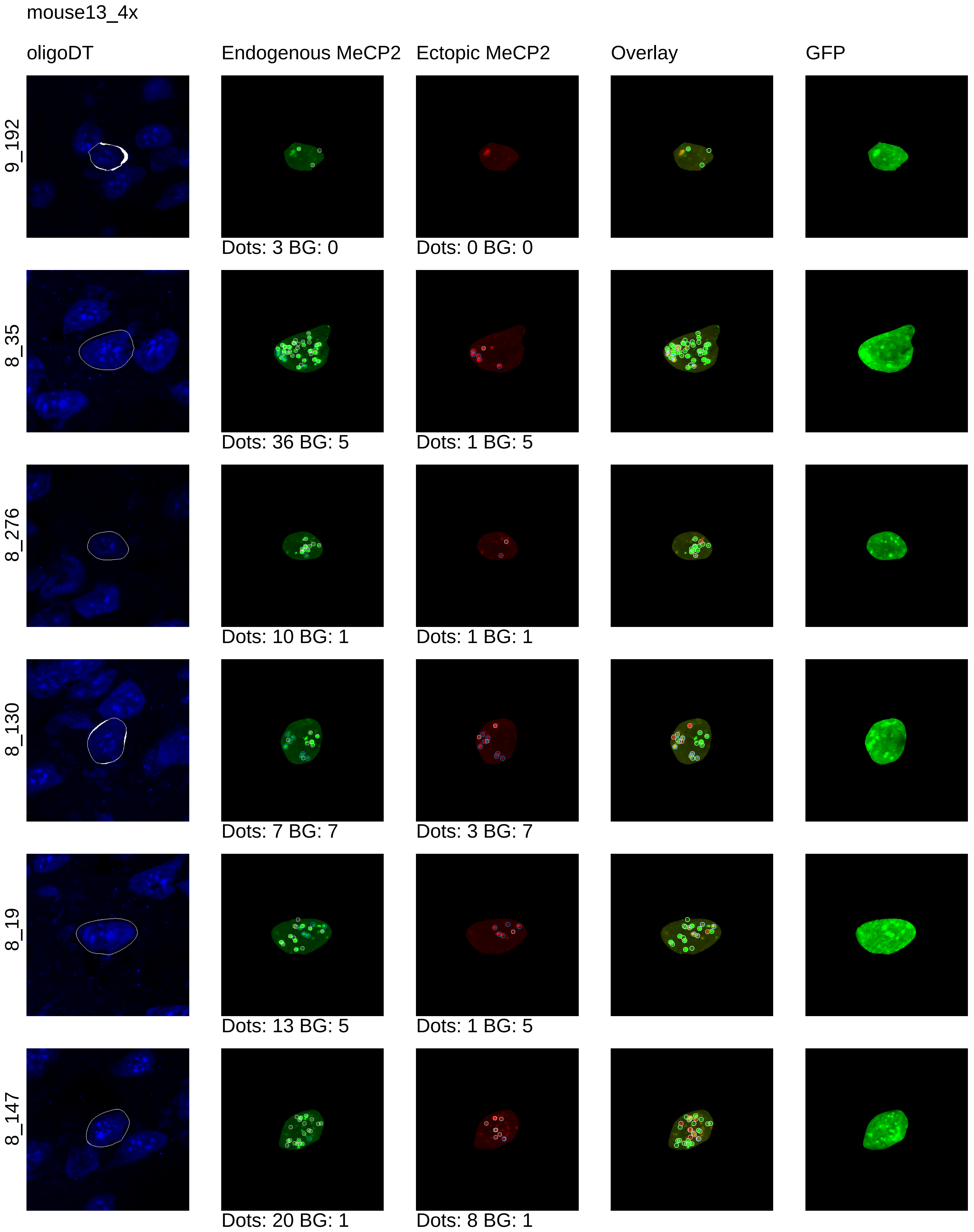
mouse13_4x
oligoDT
Endogenous MeCP2
Ectopic MeCP2
Overlay
GFP
9_192
Dots: 3 BG: 0
Dots: 0 BG: 0
8_35
Dots: 36 BG: 5
Dots: 1 BG: 5
8_276
Dots: 10 BG: 1
Dots: 1 BG: 1
8_130
Dots: 7 BG: 7
Dots: 3 BG: 7
8_19
Dots: 13 BG: 5
Dots: 1 BG: 5
8_147
Dots: 20 BG: 1
Dots: 8 BG: 1

mouse13_4x

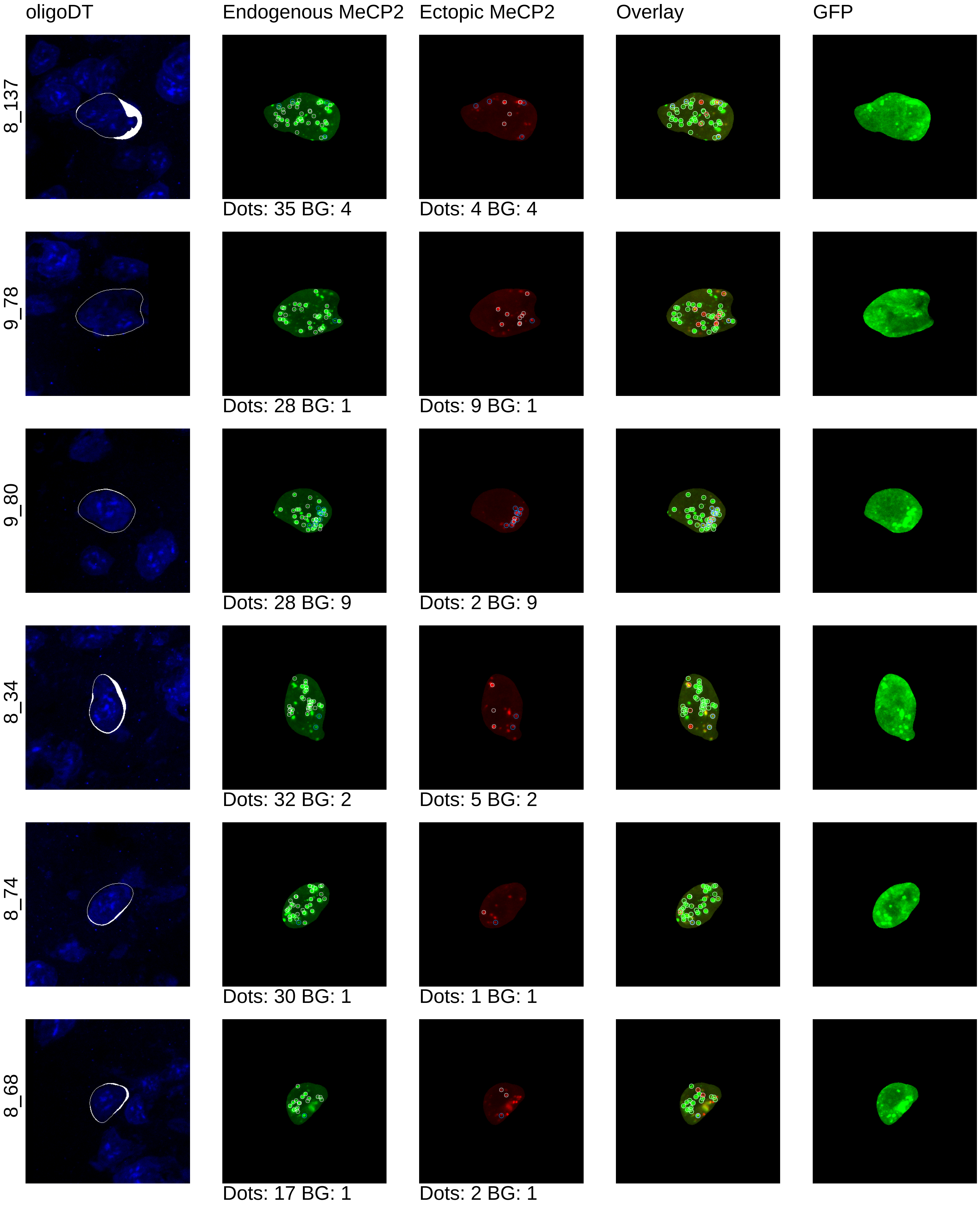
oligoDT
Endogenous MeCP2
Ectopic MeCP2
Overlay
GFP
8_137
Dots: 35 BG: 4
Dots: 4 BG: 4
9_78
Dots: 28 BG: 1
Dots: 9 BG: 1
9_80
Dots: 28 BG: 9
Dots: 2 BG: 9
8_34
Dots: 32 BG: 2
Dots: 5 BG: 2
8_74
Dots: 30 BG: 1
Dots: 1 BG: 1
8_68
Dots: 17 BG: 1
Dots: 2 BG: 1

mouse13_4x

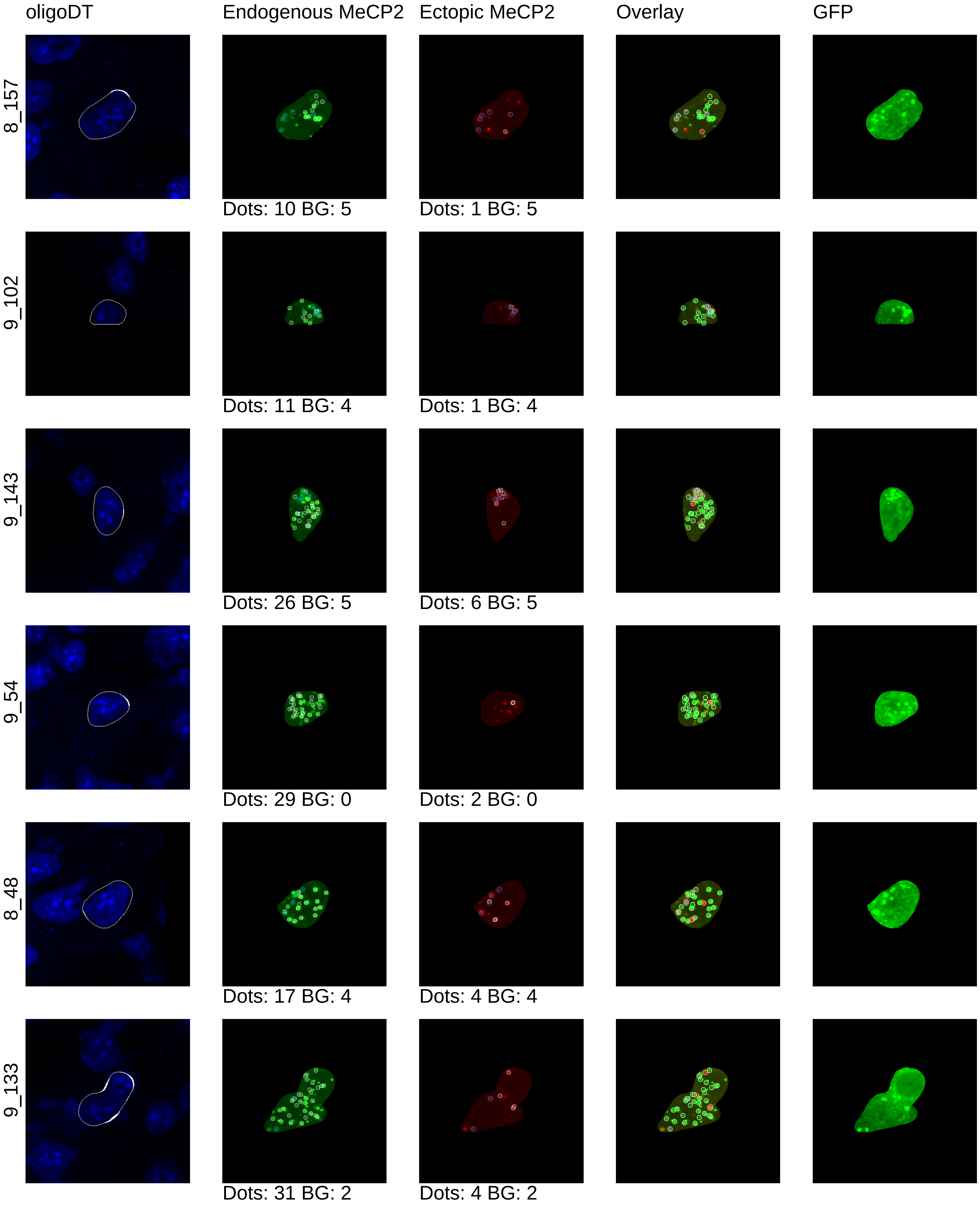
oligoDT
Endogenous MeCP2
Ectopic MeCP2
Overlay
GFP
8_157
Dots: 10 BG: 5
Dots: 1 BG: 5
9_102
Dots: 11 BG: 4
Dots: 1 BG: 4
9_143
Dots: 26 BG: 5
Dots: 6 BG: 5
9_54
Dots: 29 BG: 0
Dots: 2 BG: 0
8_48
Dots: 17 BG: 4
Dots: 4 BG: 4
9_133
Dots: 31 BG: 2
Dots: 4 BG: 2

mouse13_4x

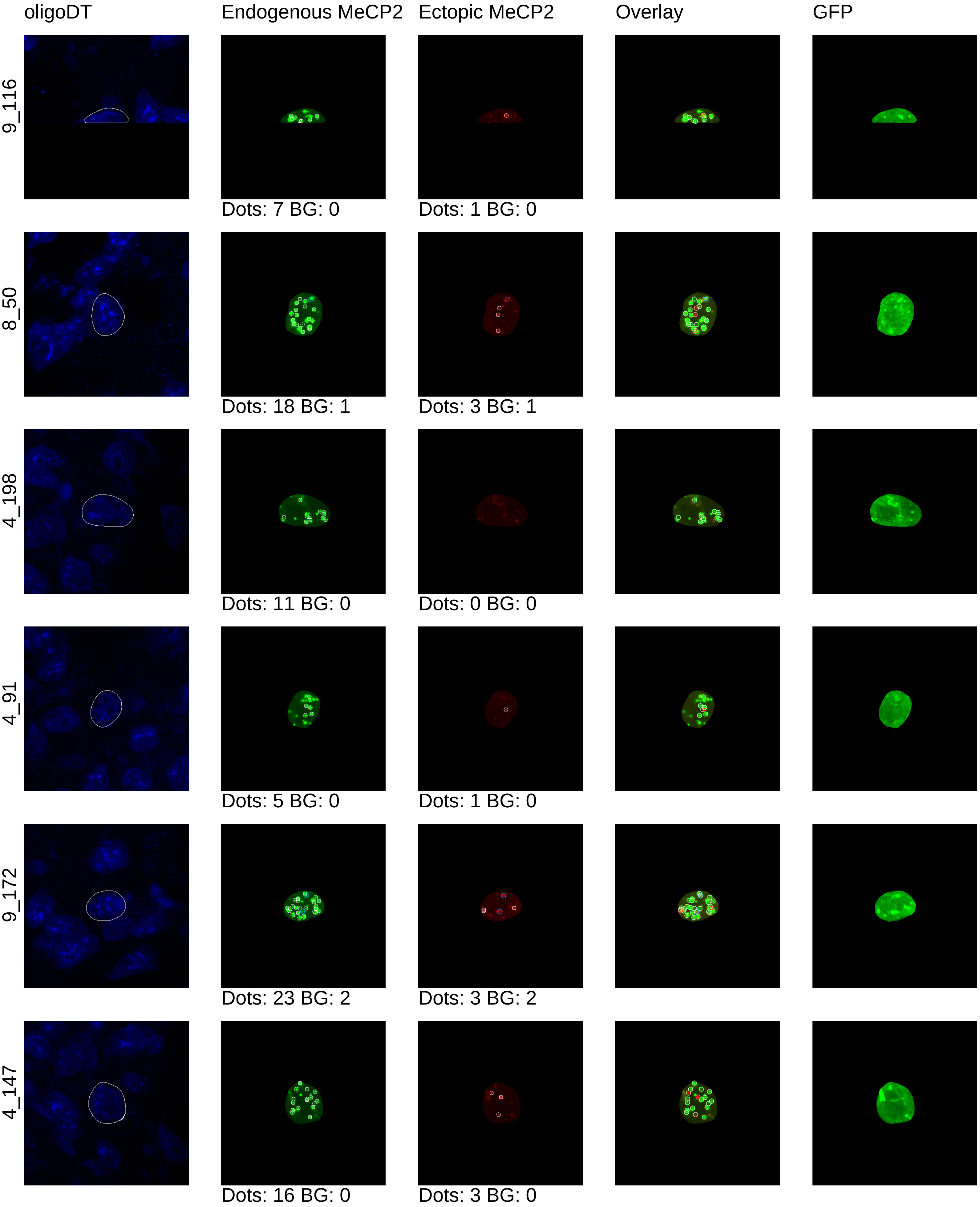
oligoDT
Endogenous MeCP2
Ectopic MeCP2
Overlay
GFP
9_116
Dots: 7 BG: 0
Dots: 1 BG: 0
8_50
Dots: 18 BG: 1
Dots: 3 BG: 1
4_198
Dots: 11 BG: 0
Dots: 0 BG: 0
4_91
Dots: 5 BG: 0
Dots: 1 BG: 0
9_172
Dots: 23 BG: 2
Dots: 3 BG: 2
4_147
Dots: 16 BG: 0
Dots: 3 BG: 0

mouse13_4x

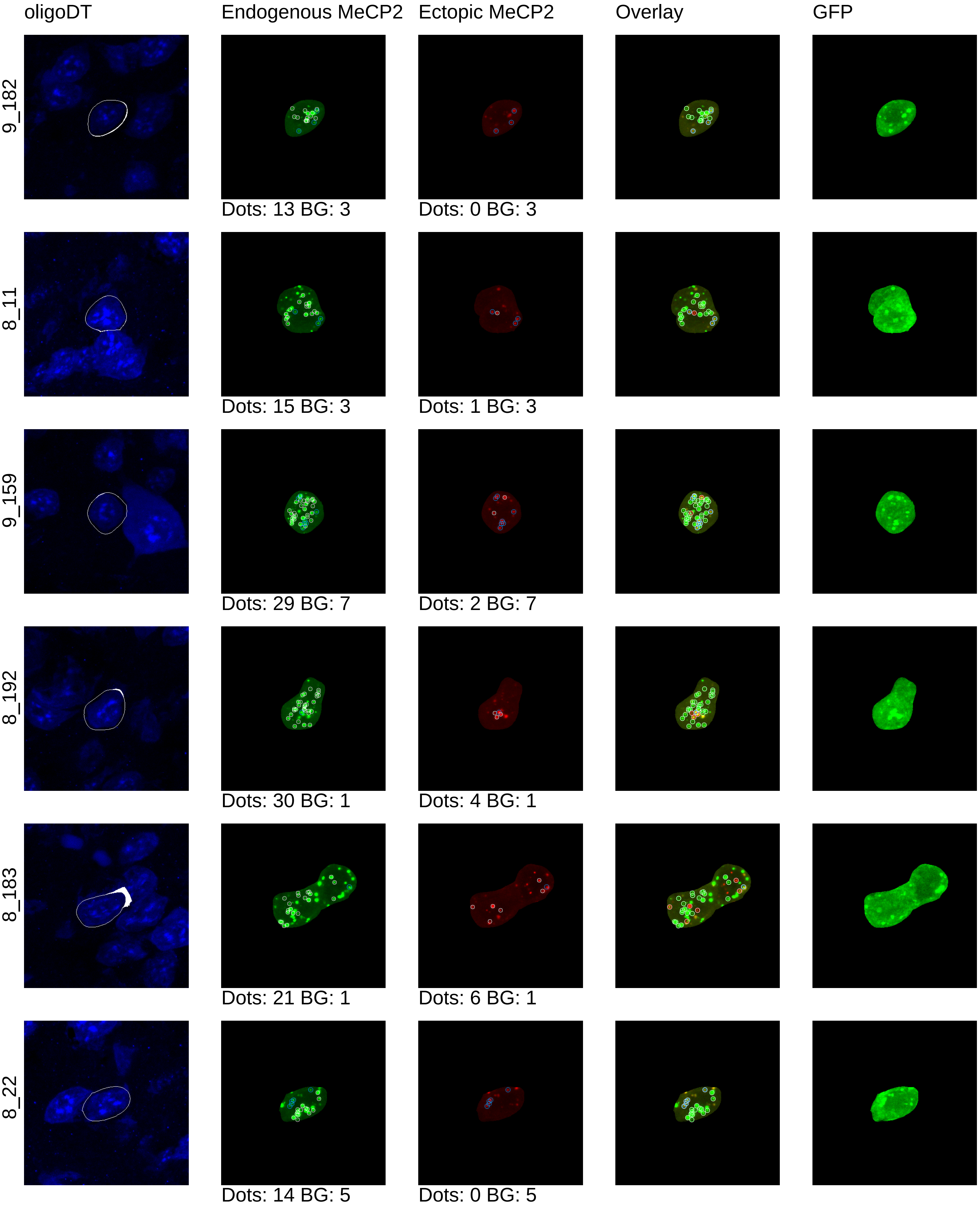
oligoDT
Endogenous MeCP2
Ectopic MeCP2
Overlay
GFP
9_182
Dots: 13 BG: 3
Dots: 0 BG: 3
8_11
Dots: 15 BG: 3
Dots: 1 BG: 3
9_159
Dots: 29 BG: 7
Dots: 2 BG: 7
8_192
Dots: 30 BG: 1
Dots: 4 BG: 1
8_183
Dots: 21 BG: 1
Dots: 6 BG: 1
8_22
Dots: 14 BG: 5
Dots: 0 BG: 5

mouse13_4x

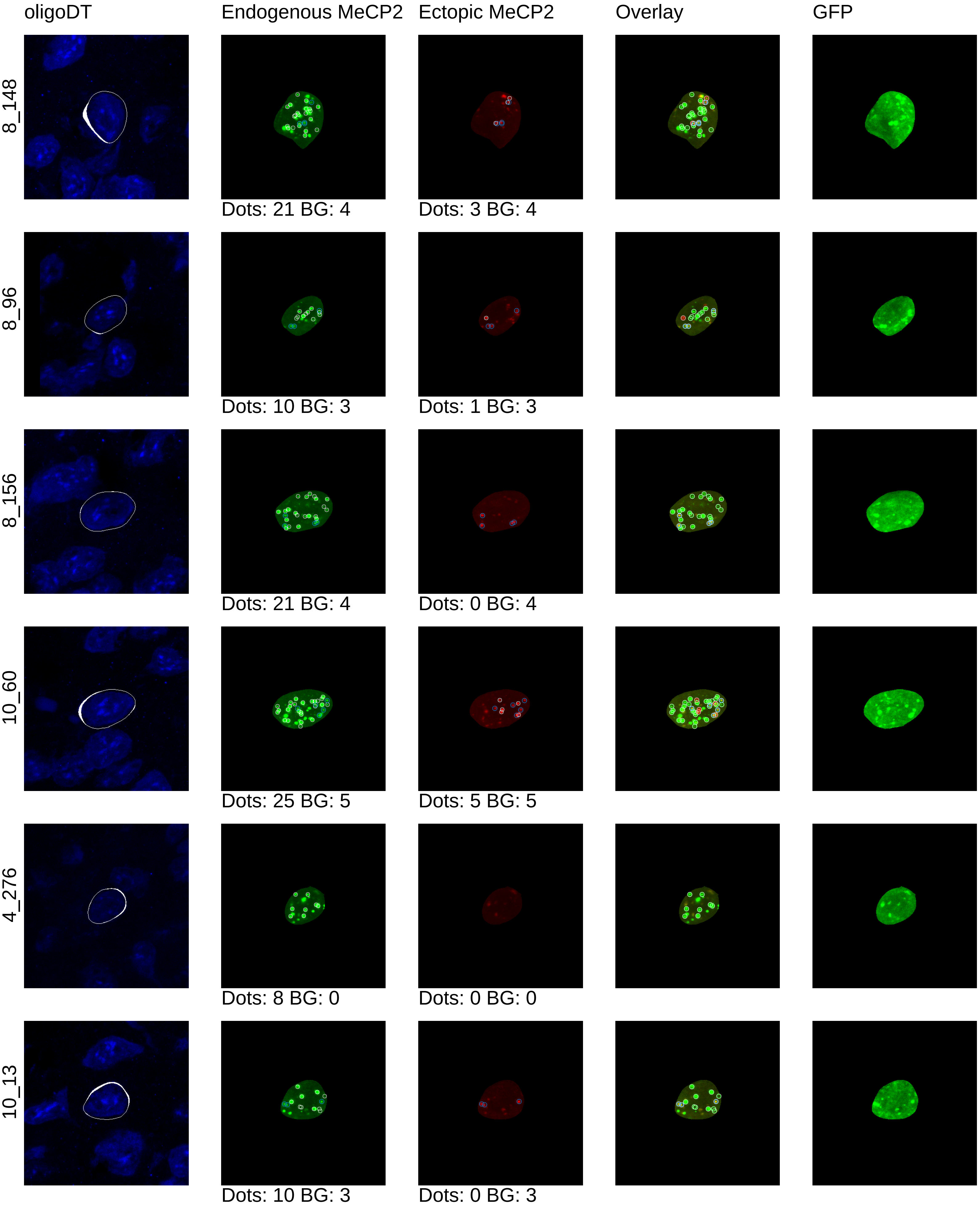
oligoDT
Endogenous MeCP2
Ectopic MeCP2
Overlay
GFP
8_148
Dots: 21 BG: 4
Dots: 3 BG: 4
8_96
Dots: 10 BG: 3
Dots: 1 BG: 3
8_156
Dots: 21 BG: 4
Dots: 0 BG: 4
10_60
Dots: 25 BG: 5
Dots: 5 BG: 5
4_276
Dots: 8 BG: 0
Dots: 0 BG: 0
10_13
Dots: 10 BG: 3
Dots: 0 BG: 3

mouse13_4x

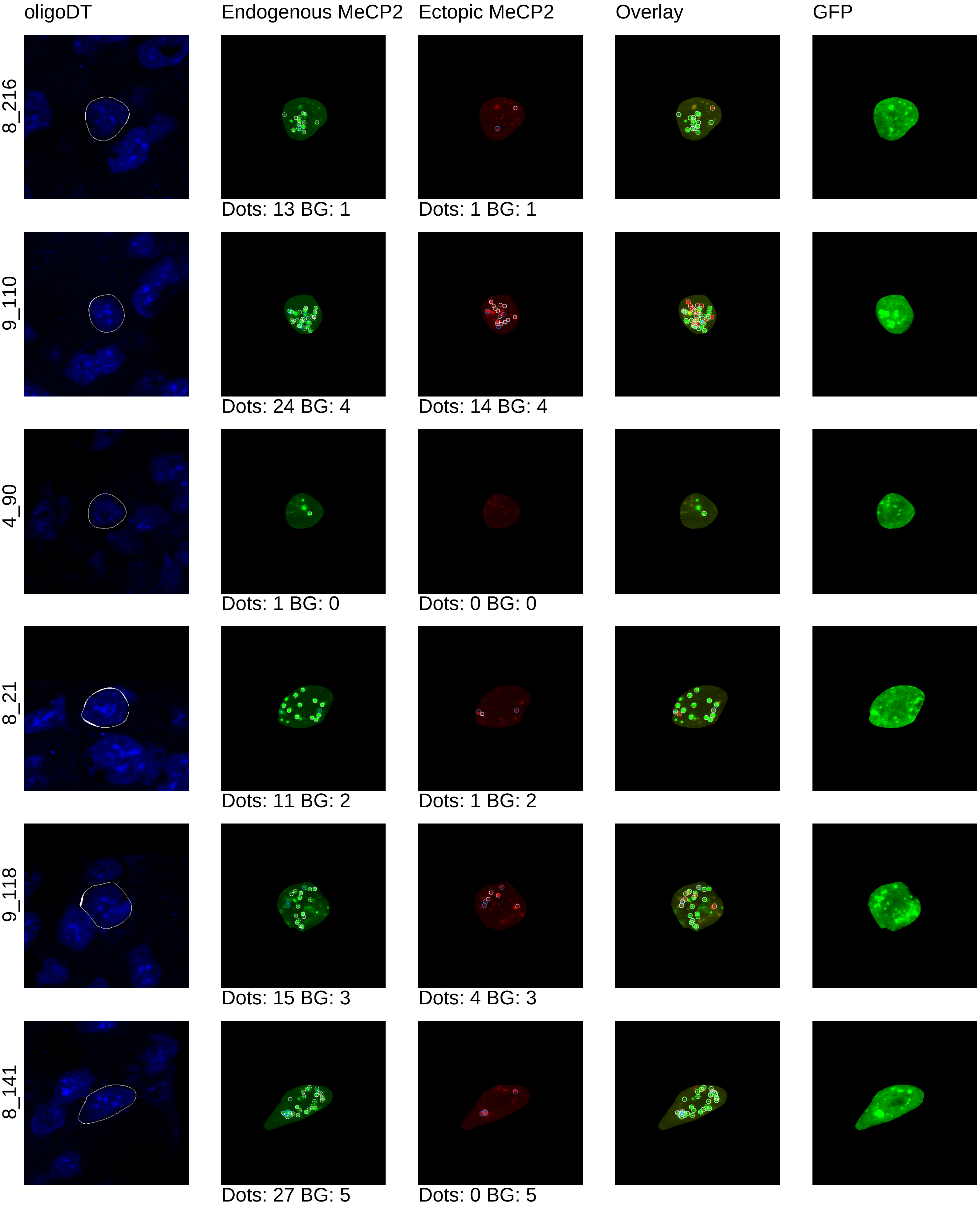
oligoDT
Endogenous MeCP2
Ectopic MeCP2
Overlay
GFP
8_216
Dots: 13 BG: 1
Dots: 1 BG: 1
9_110
Dots: 24 BG: 4
Dots: 14 BG: 4
4_90
Dots: 1 BG: 0
Dots: 0 BG: 0
8_21
Dots: 11 BG: 2
Dots: 1 BG: 2
9_118
Dots: 15 BG: 3
Dots: 4 BG: 3
8_141
Dots: 27 BG: 5
Dots: 0 BG: 5

mouse13_4x

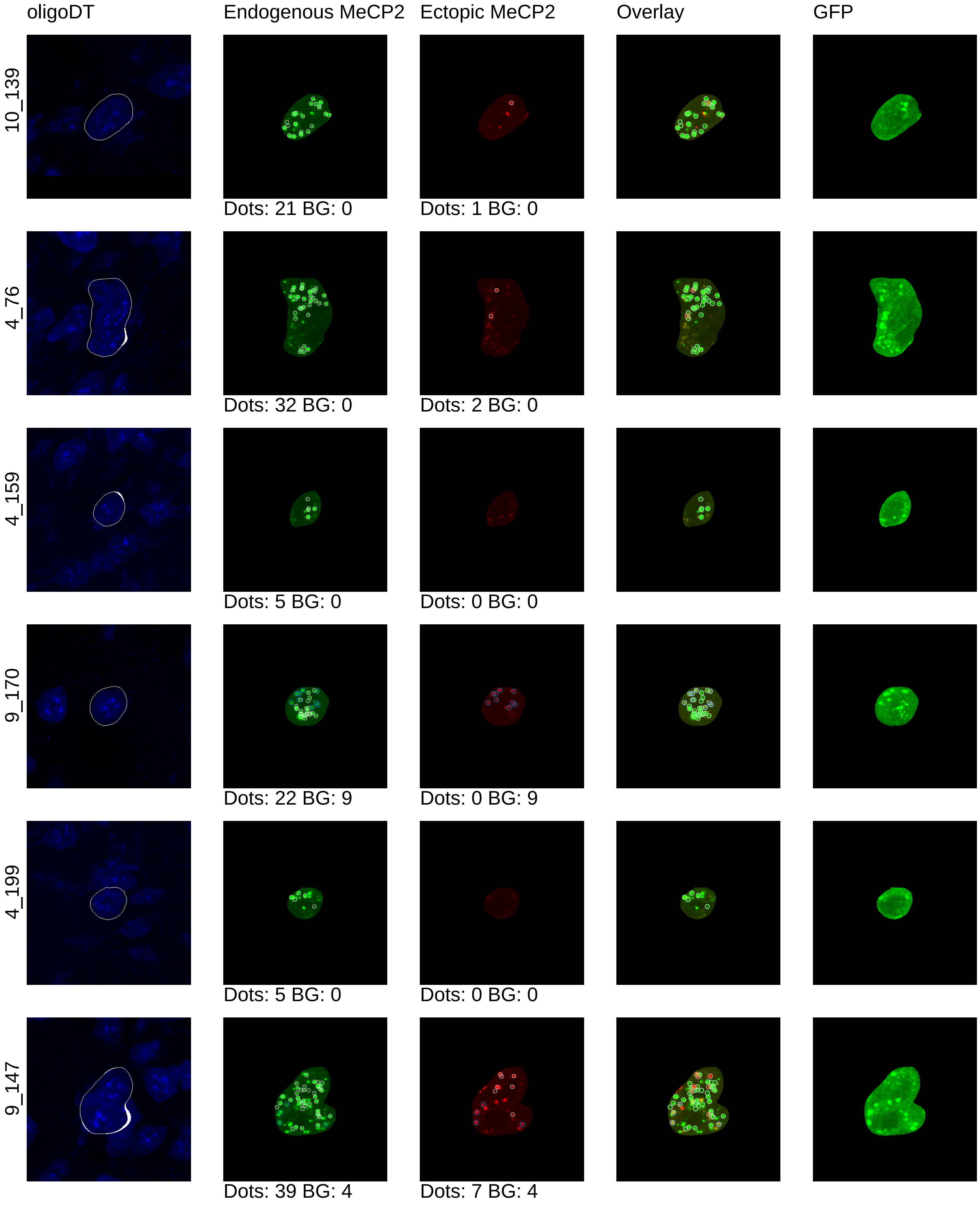
oligoDT
Endogenous MeCP2
Ectopic MeCP2
Overlay
GFP
10_139
Dots: 21 BG: 0
Dots: 1 BG: 0
4_76
Dots: 32 BG: 0
Dots: 2 BG: 0
4_159
Dots: 5 BG: 0
Dots: 0 BG: 0
9_170
Dots: 22 BG: 9
Dots: 0 BG: 9
4_199
Dots: 5 BG: 0
Dots: 0 BG: 0
9_147
Dots: 39 BG: 4
Dots: 7 BG: 4

mouse13_4x

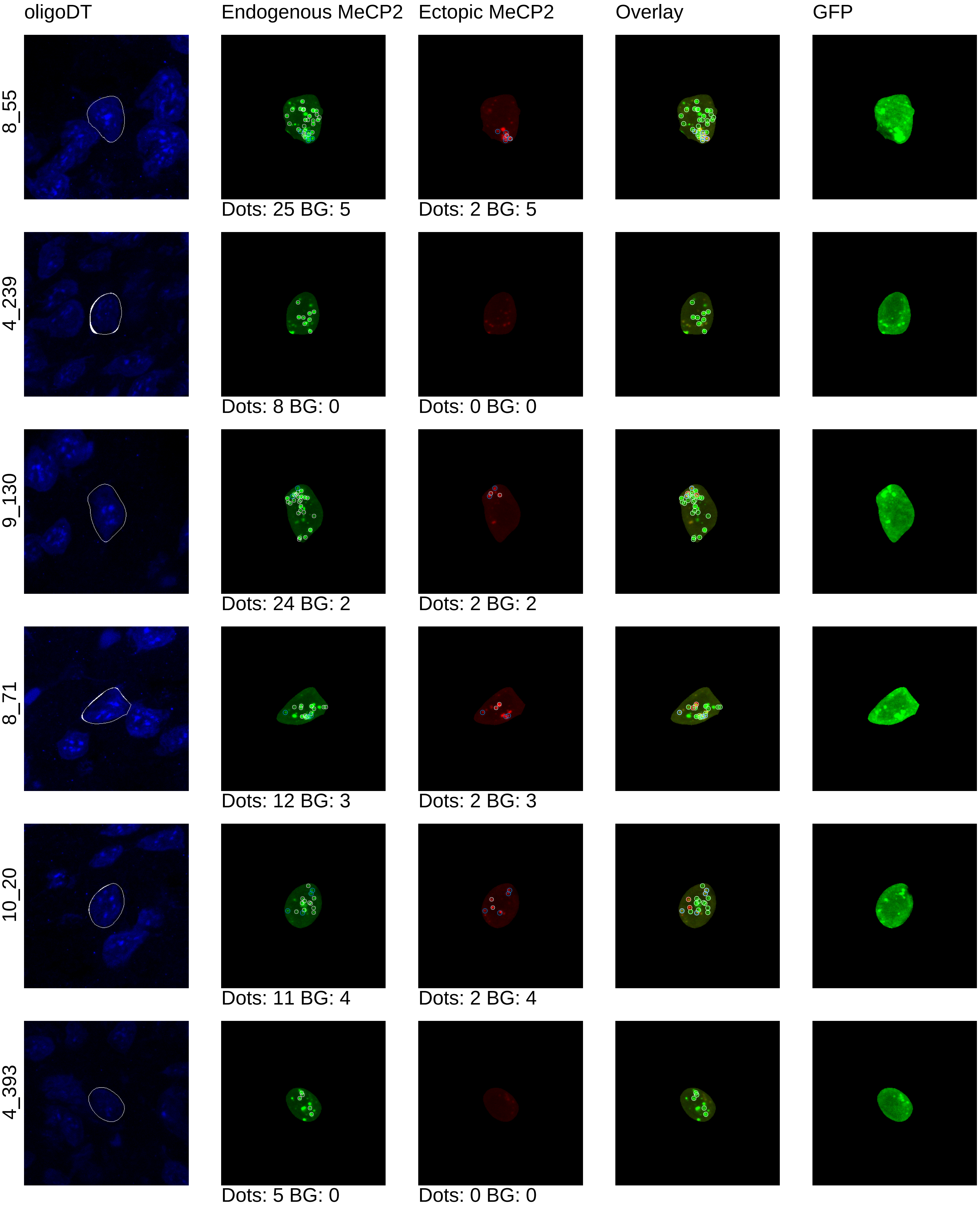

oligoDT
Endogenous MeCP2
Ectopic MeCP2
Overlay
GFP
8_55
Dots: 25 BG: 5
Dots: 2 BG: 5
4_239
Dots: 8 BG: 0
Dots: 0 BG: 0
9_130
Dots: 24 BG: 2
Dots: 2 BG: 2
8_71
Dots: 12 BG: 3
Dots: 2 BG: 3
10_20
Dots: 11 BG: 4
Dots: 2 BG: 4
4_393
Dots: 5 BG: 0
Dots: 0 BG: 0

mouse13_4x

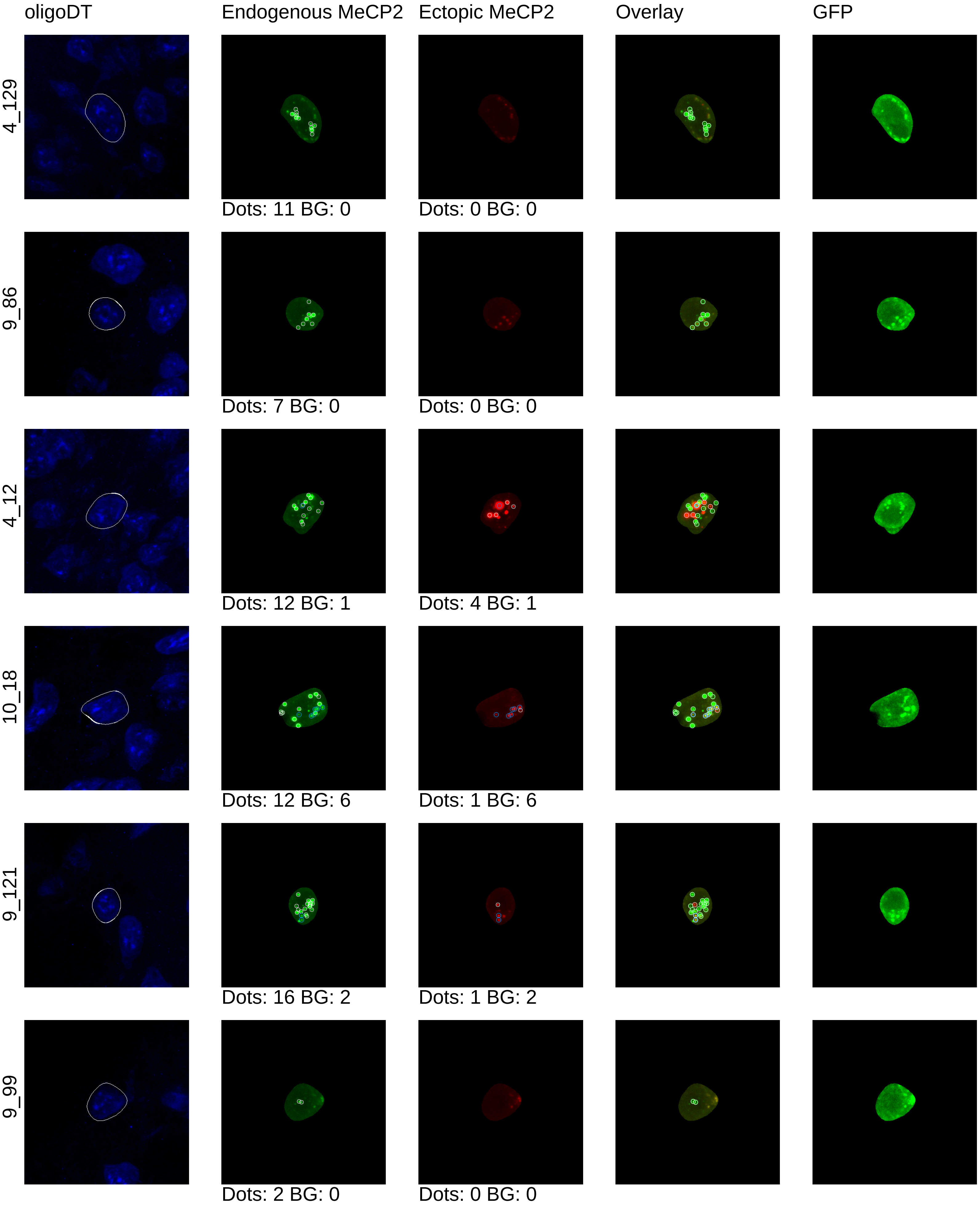
oligoDT
Endogenous MeCP2
Ectopic MeCP2
Overlay
GFP
4_129
Dots: 11 BG: 0
Dots: 0 BG: 0
9_86
Dots: 7 BG: 0
Dots: 0 BG: 0
4_12
Dots: 12 BG: 1
Dots: 4 BG: 1
10_18
Dots: 12 BG: 6
Dots: 1 BG: 6
9_121
Dots: 16 BG: 2
Dots: 1 BG: 2
9_99
Dots: 2 BG: 0
Dots: 0 BG: 0

### mouse13_4x

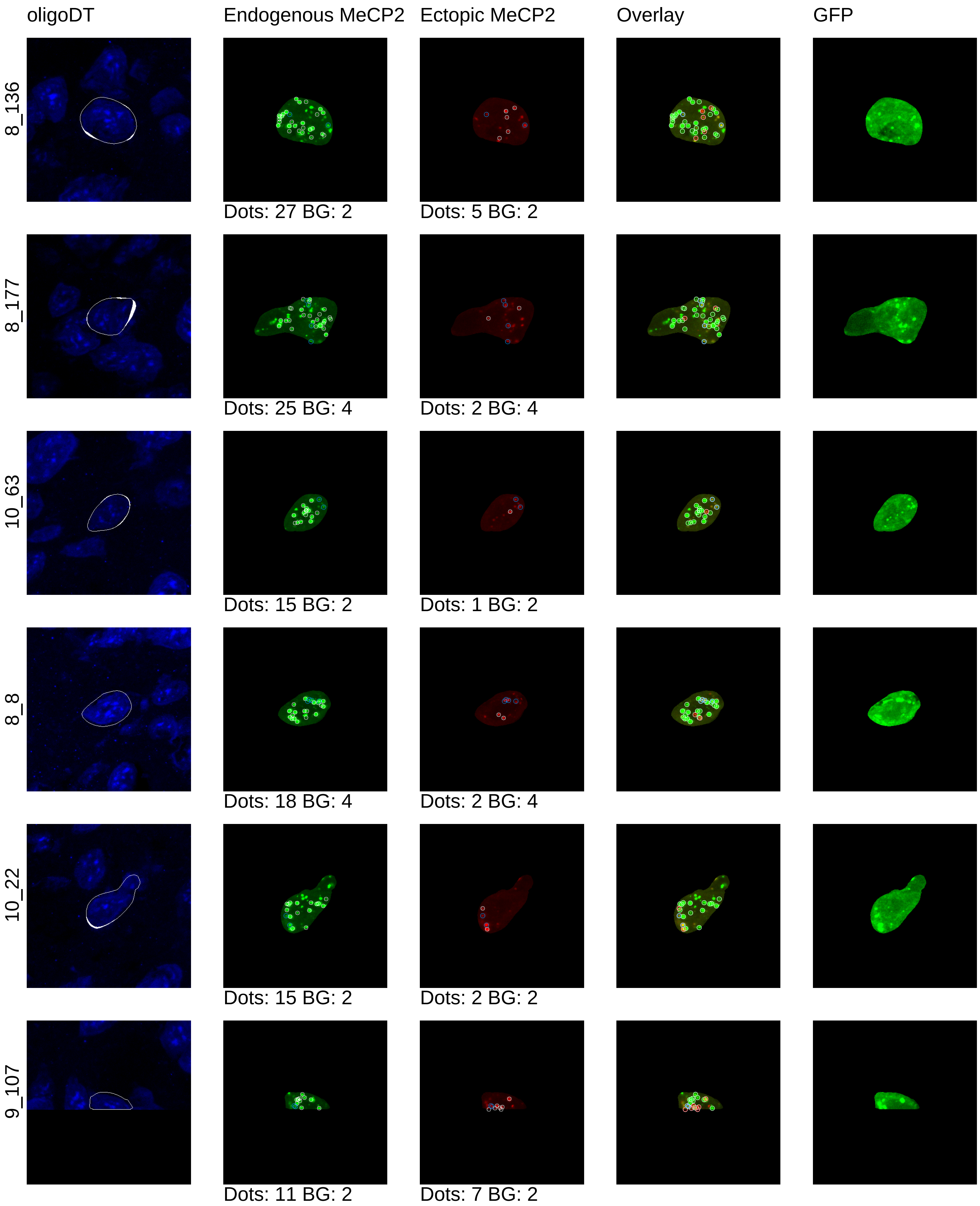
oligoDT
Endogenous MeCP2
Ectopic MeCP2
Overlay
GFP
8_136
Dots: 27 BG: 2
Dots: 5 BG: 2
8_177
Dots: 25 BG: 4
Dots: 2 BG: 4
10_63
Dots: 15 BG: 2
Dots: 1 BG: 2
8_8
Dots: 18 BG: 4
Dots: 2 BG: 4
10_22
Dots: 15 BG: 2
Dots: 2 BG: 2
9_107
Dots: 11 BG: 2
Dots: 7 BG: 2

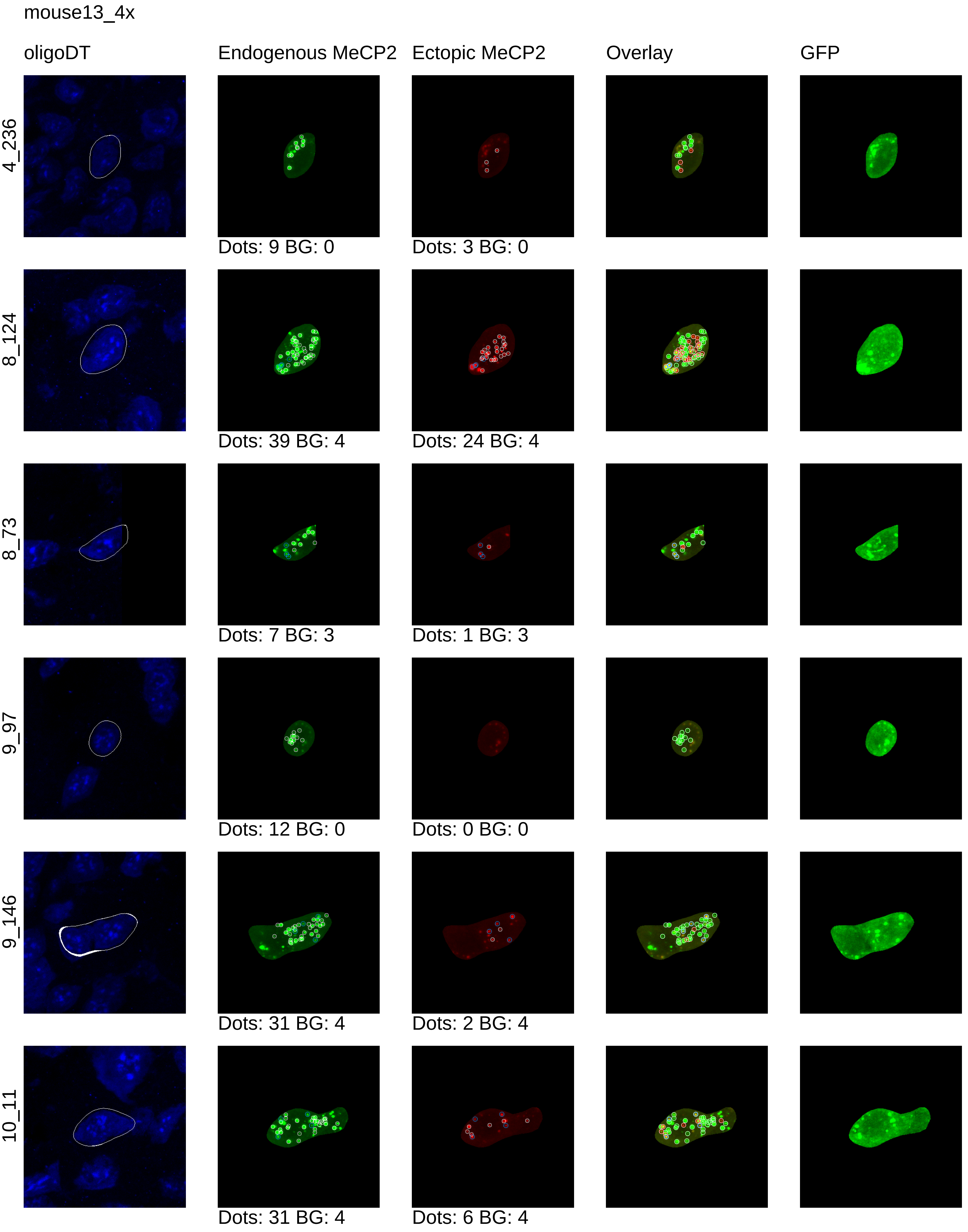
mouse13_4x
oligoDT
Endogenous MeCP2
Ectopic MeCP2
Overlay
GFP
4_236
Dots: 9 BG: 0
Dots: 3 BG: 0
8_124
Dots: 39 BG: 4
Dots: 24 BG: 4
8_73
Dots: 7 BG: 3
Dots: 1 BG: 3
9_97
Dots: 12 BG: 0
Dots: 0 BG: 0
9_146
Dots: 31 BG: 4
Dots: 2 BG: 4
10_11
Dots: 31 BG: 4
Dots: 6 BG: 4

mouse13_4x

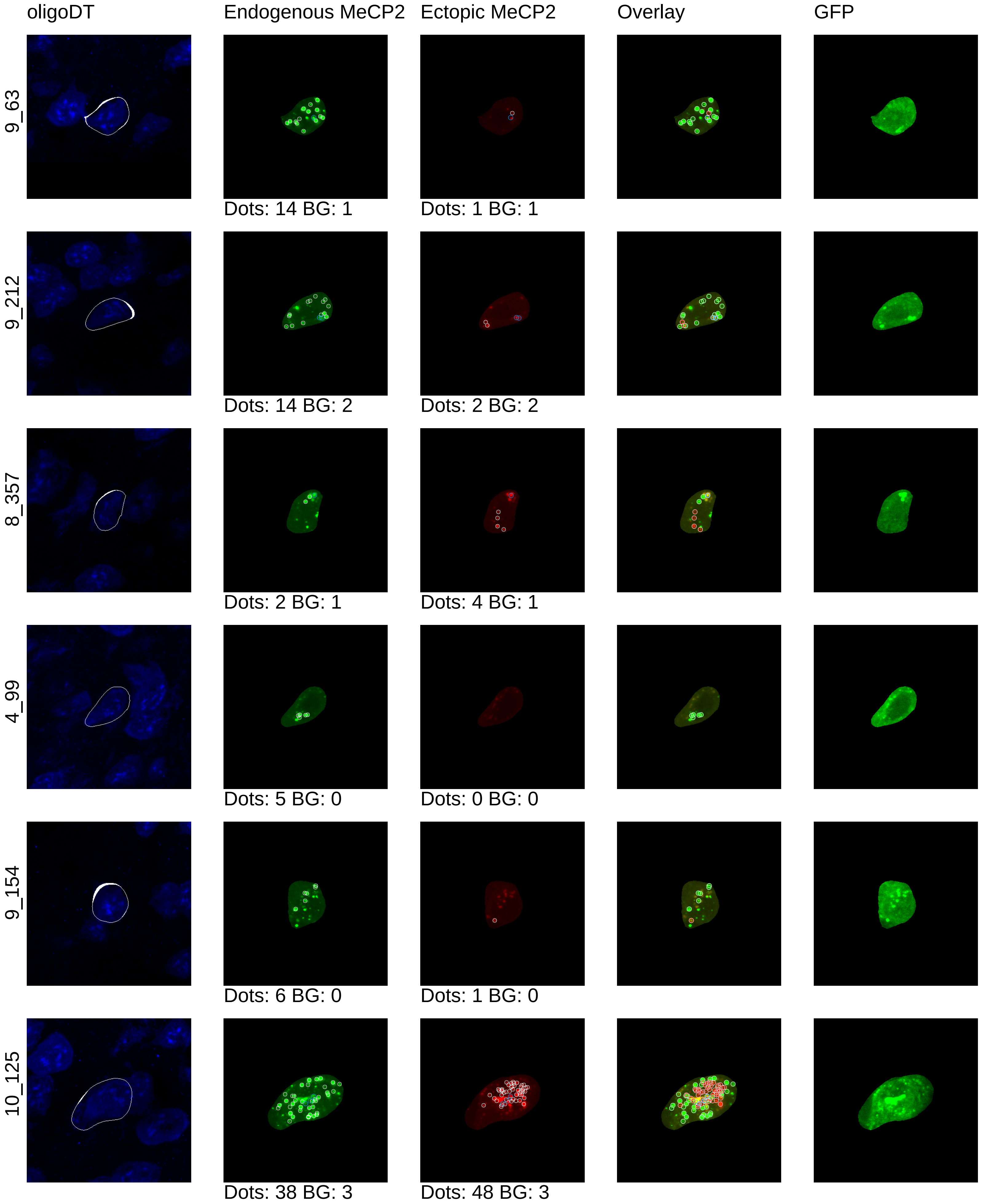

oligoDT
Endogenous MeCP2
Ectopic MeCP2
Overlay
GFP
9_63
Dots: 14 BG: 1
Dots: 1 BG: 1
9_212
Dots: 14 BG: 2
Dots: 2 BG: 2
8_357
Dots: 2 BG: 1
Dots: 4 BG: 1
4_99
Dots: 5 BG: 0
Dots: 0 BG: 0
9_154
Dots: 6 BG: 0
Dots: 1 BG: 0
10_125
Dots: 38 BG: 3
Dots: 48 BG: 3

mouse13_4x

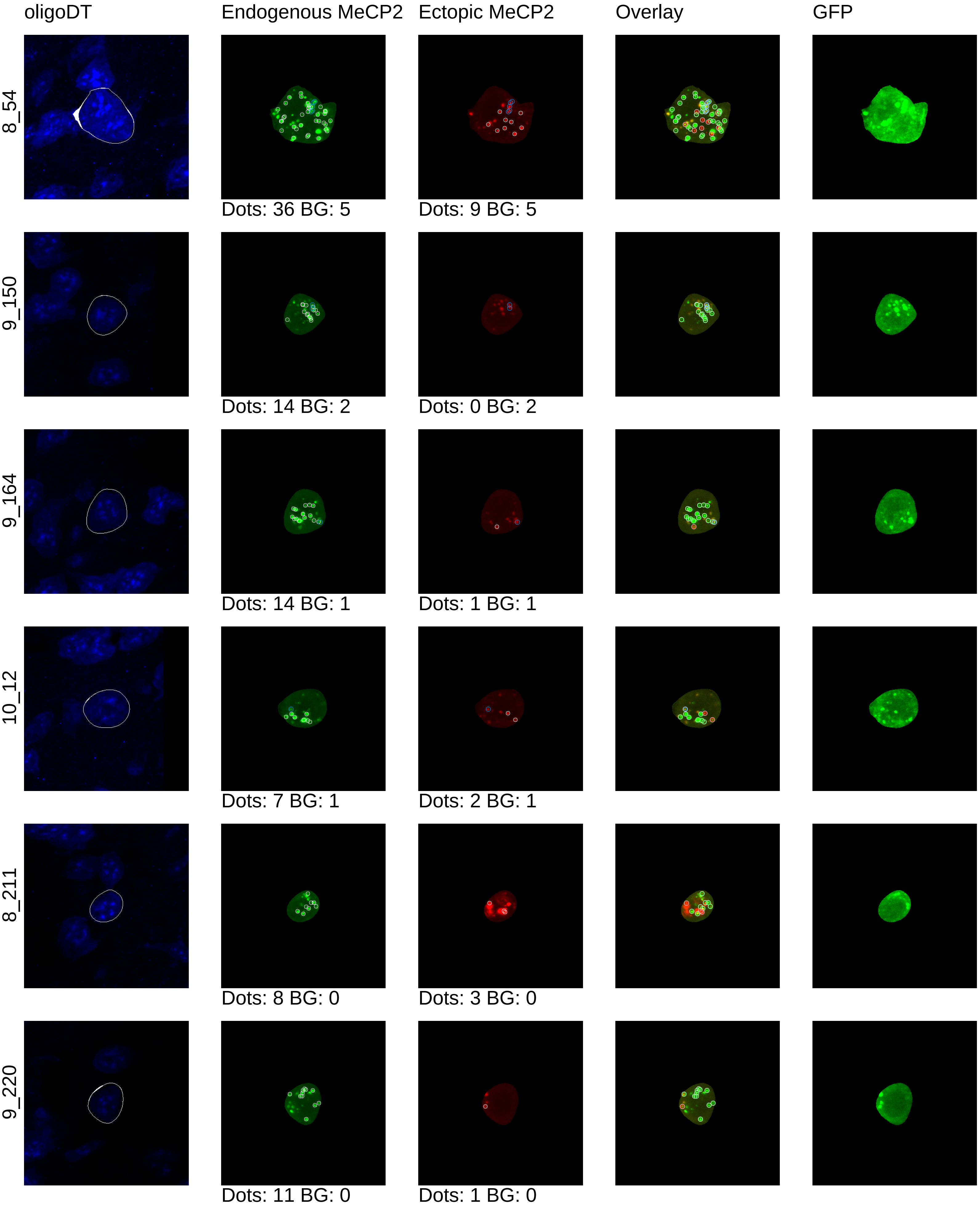
oligoDT
Endogenous MeCP2
Ectopic MeCP2
Overlay
GFP
8_54
Dots: 36 BG: 5
Dots: 9 BG: 5
9_150
Dots: 14 BG: 2
Dots: 0 BG: 2
9_164
Dots: 14 BG: 1
Dots: 1 BG: 1
10_12
Dots: 7 BG: 1
Dots: 2 BG: 1
8_211
Dots: 8 BG: 0
Dots: 3 BG: 0
9_220
Dots: 11 BG: 0
Dots: 1 BG: 0

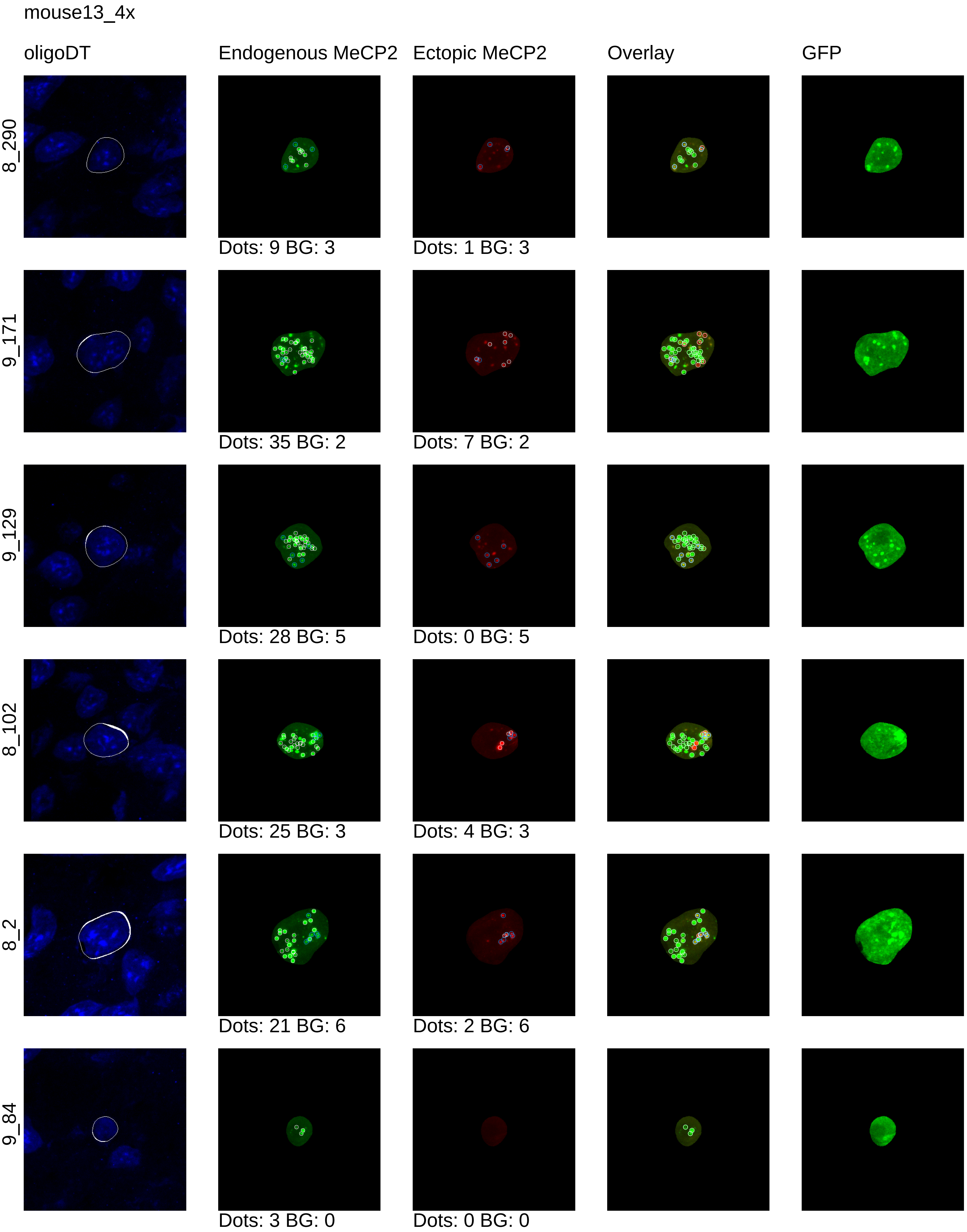
mouse13_4x
oligoDT
Endogenous MeCP2
Ectopic MeCP2
Overlay
GFP
8_290
Dots: 9 BG: 3
Dots: 1 BG: 3
9_171
Dots: 35 BG: 2
Dots: 7 BG: 2
9_129
Dots: 28 BG: 5
Dots: 0 BG: 5
8_102
Dots: 25 BG: 3
Dots: 4 BG: 3
8_2
Dots: 21 BG: 6
Dots: 2 BG: 6
9_84
Dots: 3 BG: 0
Dots: 0 BG: 0

mouse13_4x

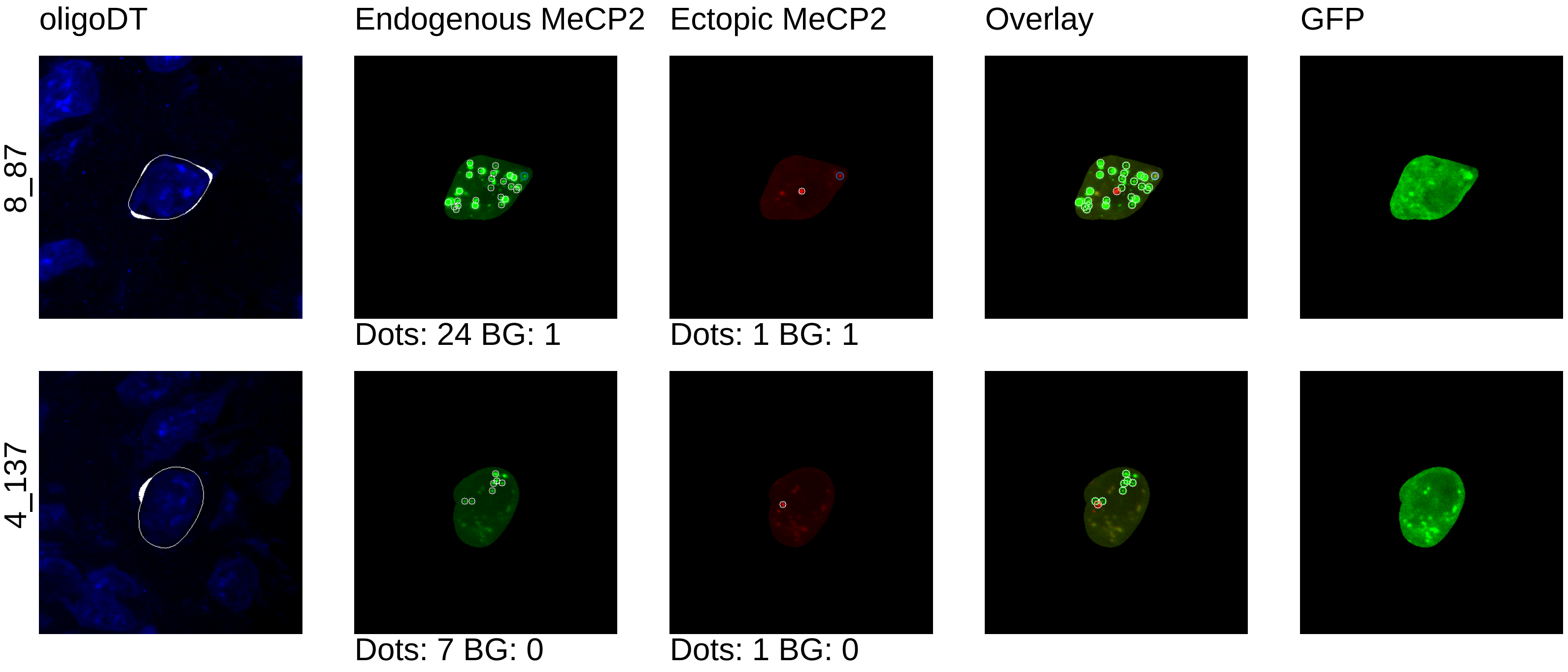
oligoDT
Endogenous MeCP2
Ectopic MeCP2
Overlay
GFP
8_87
Dots: 24 BG: 1
Dots: 1 BG: 1
4_137
Dots: 7 BG: 0
Dots: 1 BG: 0
